# High-throughput discovery of arginine-depleted peptides enables effective antisense delivery for Duchenne muscular dystrophy

**DOI:** 10.64898/2026.06.07.730741

**Authors:** Charlotte E. Farquhar, Nathan W. Dow, Carly K. Schissel, Anirban Bardhan, Alex J. Callahan, Christopher D. Greer, Alec M. Wright, Anindita Mitra, Kristin Ha, Perla Castaneda, Emily G. Thompson, Tushare Jinadasa, Ryan A. Oliver, Kathy Y. Morgan, Vincent Guerlavais, Bradley L. Pentelute

## Abstract

Phosphorodiamidate morpholino oligomers (PMOs) are approved exon-skipping antisense therapeutics for Duchenne muscular dystrophy (DMD), but their clinical utility is limited by poor uptake in muscle tissue, necessitating frequent high-dose administration. Cell-penetrating peptides (CPPs) can enhance intracellular delivery of PMOs, yet conventional arginine-rich CPPs often cause dose-limiting toxicity, including renal damage, which hinders their clinical translation. To address this challenge, we developed a high-throughput, charge-based chromatographic enrichment platform capable of screening over 15,000 synthetic peptides, including sequences with noncanonical (abiotic) amino acids. This approach enabled de novo discovery of arginine-depleted CPPs with improved delivery profiles. Four lead candidates demonstrated efficient nuclear PMO delivery with ~10-fold lower in vitro toxicity compared to standard CPPs such as penetratin. The top-performing peptide, CXP1, showed robust splice-switching activity and favorable tolerability in both cellular and animal models. In dystrophic mdx mice, CXP1–PMO conjugates achieved greater exon skipping compared to PMOs conjugated to R6G at equivalent doses. Tissue levels of CXP1–PMO correlated with exon-skipping efficacy, establishing a clear pharmacokinetic–pharmacodynamic relationship. These findings highlight a mechanistically novel and translationally relevant discovery strategy, demonstrating the potential of high-throughput platforms to generate more effective CPP-based delivery vehicles for antisense therapeutics in DMD and related neuromuscular disorders.

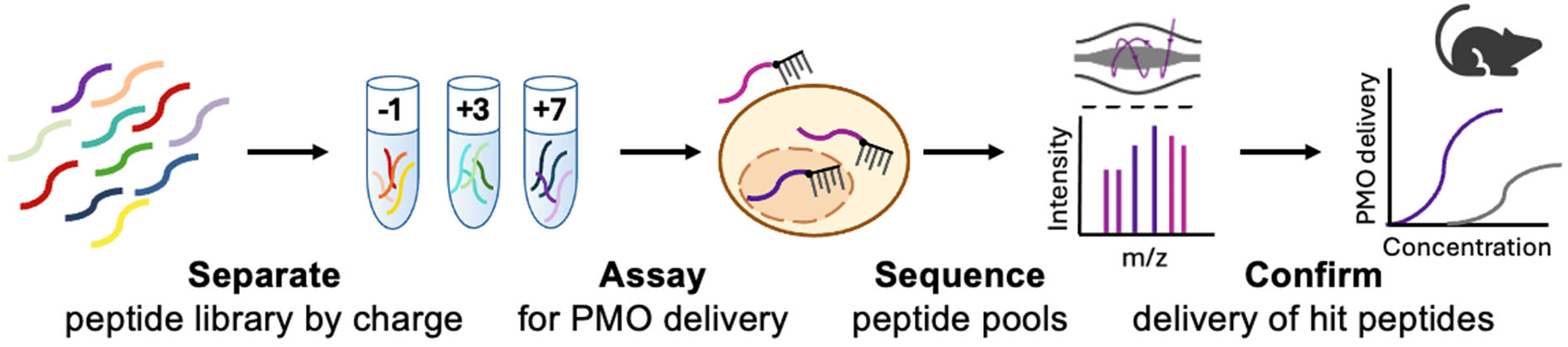

## Introduction

Antisense oligonucleotides (ASOs) are an expanding therapeutic option for sequence-specific gene regulation, altering gene expression to restore, knock-down, or modify the target protein.^1^ While approaches to delivery including polymer encapsulation, nanoparticles, and covalent modification are effective for delivering these ASOs into cells in vitro, strategies for effective in vivo delivery remain limited.^2,3^ Despite these limitations, a number of ASO therapies have achieved clinical application. Exon-skipping oligonucleotides have been developed for the treatment of Duchenne muscular dystrophy (DMD), a rare genetic neuromuscular disorder caused by mutations in the X-linked dystrophin gene.^4,5^ Exon-skipping ASO drugs can restore dystrophin function by binding a pre-mRNA transcript and blocking a splice site, correcting splicing to produce an in-frame shortened functional dystrophin.^6^ The clinically approved exon-skipping drugs for DMD have a nonnatural phosphorodiamidate morpholino oligomer (PMO) backbone, which hybridizes to the pre-mRNA. PMOs offer several distinct advantages over other ASO backbones which make them particularly attractive for therapeutic applications. Unlike phosphorothioates (PS), PMOs are uncharged and are composed of morpholine rings, which confers exceptional resistance to nucleases and proteases.^7^ Furthermore, PMOs have minimal nonspecific interactions, which reduces the off-target effects and immunogenicity, greatly improving their safety profile. Despite their clinical approval, The approved PMO therapies and the investigational PMO-based therapeutic approaches face the same delivery challenges as other ASOs.^7^ Animal and human models illustrate that PMO drugs suffer from poor cell permeability, therefore necessitating frequent, large doses to reach effective concentrations in muscle tissues.^8^ This delivery challenge driven by poor cell penetration prevents more widespread use of PMO and other antisense oligonucleotide drugs.

Cell-penetrating peptides (CPPs) have been widely studied for enhancing the intracellular delivery of covalently conjugated oligonucleotides, both in vitro and in vivo.^9–10^ Penetratin, one of the earliest and most extensively studied CPPs, was identified through investigations of the intracellular activity of homeoproteins.^11^ Subsequent experiments revealed that this peptide could efficiently deliver various payloads to cells, providing the foundation for developing additional cell-penetrating peptides. Over the years, more than 1,000 CPP sequences have been discovered and applied to the effective delivery of cargoes including fluorophores, proteins, and ASOs.^12,13^ CPP-mediated delivery is cargo-dependent, making existing peptides not always effective at delivering a variety of molecules with different molecular weights or biophysical properties.^14,15^

CPPs have traditionally incorporated arginine residues to improve delivery efficiency, owing to arginine’s unique chemical and biophysical properties. The guanidinium group in the arginine enables hydrogen bonding and electrostatic interaction with negatively charged components of the cell membrane, facilitating the binding and internalization of CPP-oligonucleotide conjugates.^16–19^ Oligoarginine-based CPPs, such as R12, R6G, and the B-peptide (also termed Bpep, RXRRβRRXRRβR, where X is aminohexanoic acid and β is β-alanine), have been widely studied in the past to improve, with increased numbers of arginine moieties shown to improve cellular uptake.^20–22^ However, while higher arginine content has been demonstrated to increase intracellular delivery, they are also associated with increased cytotoxicity and reduced in vivo tolerability.^18, 23–27^

While the highly cationic peptides continue to show promise as effective vectors to deliver PMO and other antisense oligonucleotides into cells, charged guanidinium-rich sequences such as polyarginines, are not sufficient for safe therapeutic delivery. The peptide-PMO conjugate SRP-5051 (vesleteplirsen) demonstrated more effective delivery and dystrophin synthesis compared to eteplirsen, although clinical development was discontinued due to undesired hypomagnesemia.^10,26^ Therefore, more effective means of discovering peptide sequences that are capable of enhanced cargo delivery with minimal toxicity is paramount.^28,29^

Previous work in the discovery of CPPs for oligonucleotide delivery have ranged from testing known CPPs from the literature to rationally or computationally designing novel sequences.Click or tap here to enter text.^16–19,30–32^ While individually synthesizing and testing new CPP-oligonucleotide conjugates has produced positive outcomes, the process can be time-consuming and limits the overall number of peptide sequences that can be generated. These studies have uncovered general trends in improving CPPs for PMO delivery, including the importance of positive charge, peptide length, arginine content, macrocyclization and secondary structure.^30,33–36^

These previous results underscore the need for discovering more diverse sequences, including those that contain amino acids not found in nature. Noncanonical amino acids are found in many of the most effective CPPs for PMO delivery developed so far,^19,26,30^ and they offer increased protease stability and chemical diversity compared to canonical residues.^27,37^ Computational models trained on existing datasets have generated additional novel CPPs, but their sequences remain limited by the predominantly natural amino acids and structures found in the training data.^15,30^ Additional machine-learning methods trained on larger datasets are often no longer cargo-specific, leading to significant variability in the ability to deliver different classes of ASOs.^38,39^ Methods that allow the rapid generation and screening of thousands to millions of peptides include display methodologies such as mRNA display and phage display, and one-bead-one-compound screening.^40–48^ While these methods offer facile workflows to test peptide libraries up to 10^13^ members, many also include bulky encoding tags and it is difficult to incorporate an ASO during the screening or peptide generation process. Thus, there remains a need for the implementation of additional platforms to discover novel, abiotic CPPs tailored towards the delivery of therapeutic ASO cargoes with low arginine content and toxicity.

To further push the boundaries of de novo CPP discovery, we developed a chromatographic resolution assay platform for the discovery of novel CPPs from an 18,000-member library of synthetic, randomized peptide sequences. Previous work has demonstrated that chromatographic methods confer reduced experimental error and improved handling of complex mixtures in the context of discovering membrane-permeable peptides.^49^ While Koch et al.^49^ used hydrophobicity as a correlating property for passive membrane diffusion, positive charge has been found to be a stronger driver of cell uptake efficiency with CPP-cargo conjugates.^20,50,51^ We hypothesized that library selection by charge would permit the discovery of novel CPPs, allowing us to combine both high-throughput discovery and design-based optimization in a single platform.

Herein, we report a pool-testing strategy that resolves a large and chemically diverse peptide library based on charge into smaller pools that can be sequenced and evaluated for their PMO delivery efficacy (**Figure 1**). This methodology utilizes existing principles of CPP design, incorporates nonnatural residues into the generated peptides, and includes the cargo of interest (PMO) during the screening process. The peptide sequences recovered from the library pools revealed a clear trend between more positively charged peptides and increased PMO delivery. The chromatographic enrichment platform produced four new abiotic CPPs with high propensity for delivering PMOs, measured in terms of exon skipping efficacy, and low in vitro toxicity. The most efficacious sequence discovered, CXP1, demonstrated high PMO delivery and tolerability both in vitro and in vivo with multiple PMO sequences. Despite containing minimal arginine residues, the CXP1 peptide outperformed the traditional arginine-rich B-peptide and R_6_G vectors in PMO delivery in both healthy and diseased mouse models.

**Figure 1:**
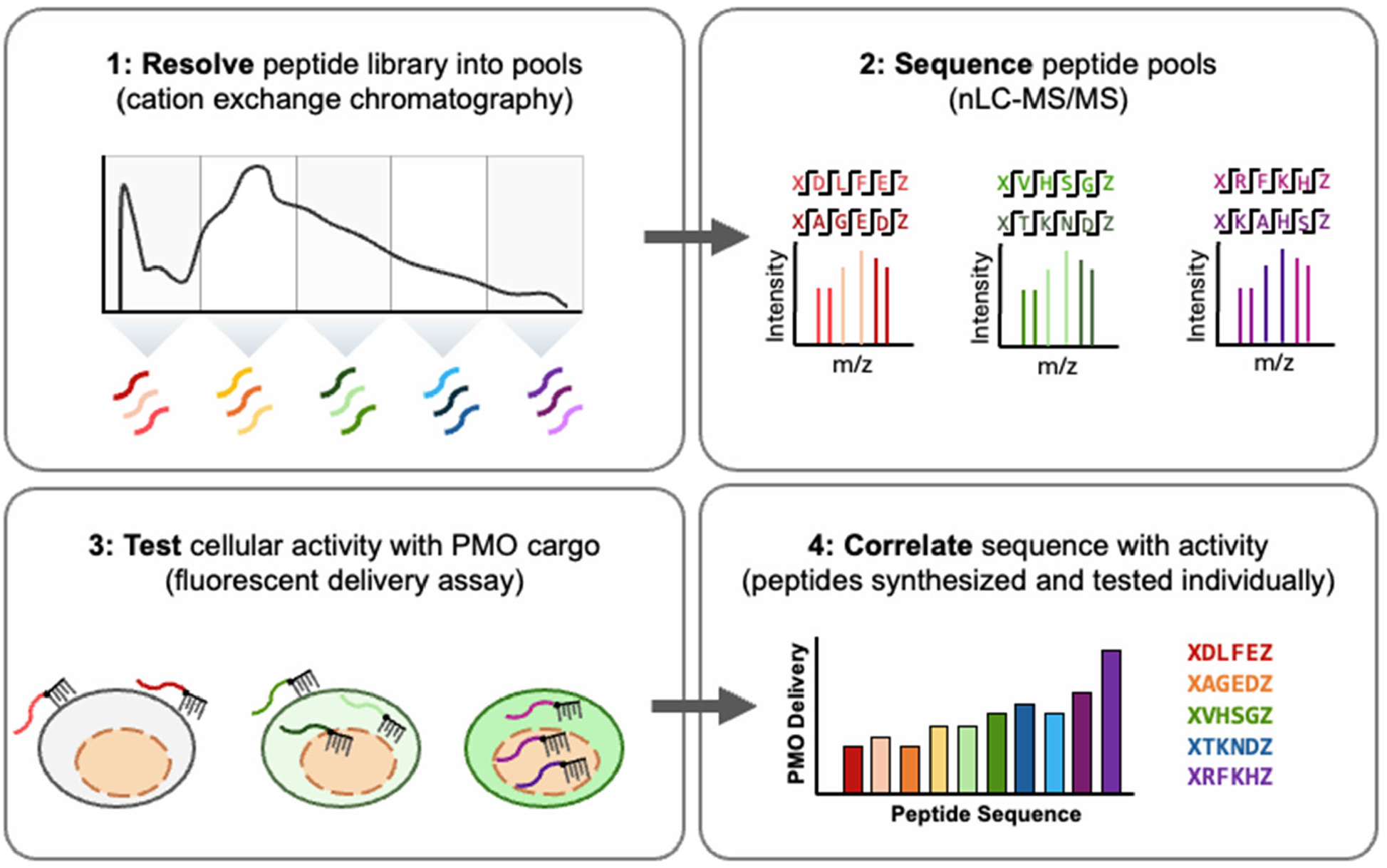
Overview of the chromatographic resolution platform for discovering novel peptides for PMO delivery. **(1)** A diverse and random peptide library is generated through split-and-pool SPPS. The library is then resolved into smaller pools through cation exchange chromatography, using an ammonium acetate gradient to elute more positively charged peptides over time. The eluted peptides are then combined into pools containing sequences of similar charge. **(2)** The peptide pools are sequenced through MS/MS to reveal the peptide sequences within each pool. **(3)** The pools are desalted and conjugated to the DBCO-labelled PMO cargo through strain-promoted azide-alkyne cycloaddition (SPAAC) chemistry. Each pool is tested for delivery in HeLa 654 cells, which exhibit fluorescence upon nuclear PMO delivery of the anti-IVS2-654 antisense cargo. **(4)** Peptide sequences from top-performing pools are resynthesized and tested for delivery individually. Activity is then correlated with individual peptide sequences.

## Results

### Chromatographic resolution of a peptide library demonstrates a positive trend between peptide charge and PMO delivery into cells

We designed and synthesized an abiotic library containing a fixed “CPP-like” motif to facilitate chromatographic resolution and subsequent sequencing of library fractions. The C-terminal four amino acid motif, KWKK, derived from the CPP penetratin, improves binding to a cation exchange column and has been previously demonstrated to increase the CPP-like character of a peptide library.^48,11^ The peptide library was designed with a focus on increasing chemical diversity and stability with the incorporation of canonical and noncanonical amino acids within the ten residues of the variable region. 16 natural amino acids were used along with 6 additional noncanonical amino acids (theoretical diversity = 2.7 × 10^13^) (**Figure 2**). The overall construct size, variable region length, and amino acids chosen were optimized for peptide sequencing based on our previous work.^52^ The N-terminal 5-azidopentanoic acid permits highly efficient coupling to the PMO cargo through strain-promoted azide-alkyne cycloaddition (SPAAC). The library was generated through split-pool solid-phase peptide synthesis on 180 µm monosized TentaGel resin (0.64 g; 5 × 10^5^ beads). 12,000-member and 3,000-member portions of the library were cleaved off resin and analyzed by Orbitrap nano-liquid chromatography–tandem mass spectrometry (nLC– MS/MS), confirming effective incorporation of the canonical and noncanonical residues across the library constructs. Canonical and noncanonical residues were uniformly incorporated across the variable region, except for the guanidyl and pyridyl-alanine residues, which exhibited reduced incorporation (**Supplementary Information Section 4**), following known trends.^53^

**Figure 2:**
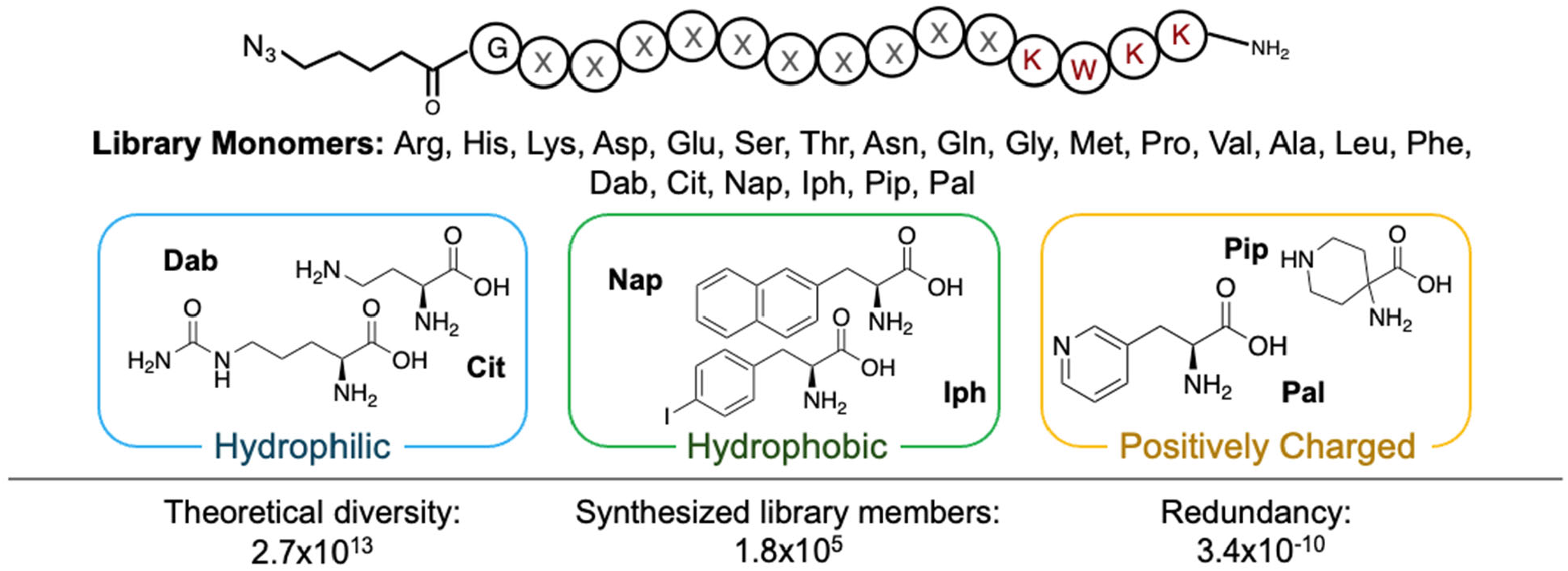
CPP library design. The library construct (top) and variable region monomers (bottom). The sequence in red is derived from the existing CPP penetratin, and the remaining library (represented by “X”) consists of a variable region containing any of the 22 monomers. The glycine at the N-terminus separates the N-terminal azide from the variable region. The library is amidated at the C-terminus. “Dab” represents L-2,4-diaminobutyric acid, “Cit” represents L-citrulline, “Nap” represents 3-(2-naphthyl)-L-alanine, “Iph” represents 4-iodo-L-phenylalaine, “Pip” represents 4-aminopiperidine-4-carboxylic acid, and “Pal” represents 3-(4’-pyridyl)-L-alanine.

The peptide library was resolved chromatographically into ten smaller pools containing peptides with varying, gradually increasing average charge. After cleavage from resin, 3,000 library peptides were separated through cation exchange chromatography based on their charge-mediated interactions with the column. Peptides were collected into small library pools based on the UV chromatogram to produce ten pools of ~300 individual peptide sequences (**Figure 3a**). UV-Vis quantification of each pool at 280 nm was performed to ensure that each pool contained equal concentration of peptide. MS/MS sequencing of the peptides within each pool demonstrated that the ten library pools differ from each other based on the net charge of their peptides (**Figure 3b**). These smaller library pools, numbered 01-10 based on their cation exchange elution time, were then desalted, conjugated to the PMO cargo through the dibenzocyclooctyne (DBCO) tag on the 5’-end of the PMO, and dried via lyophilization.

**Figure 3:**
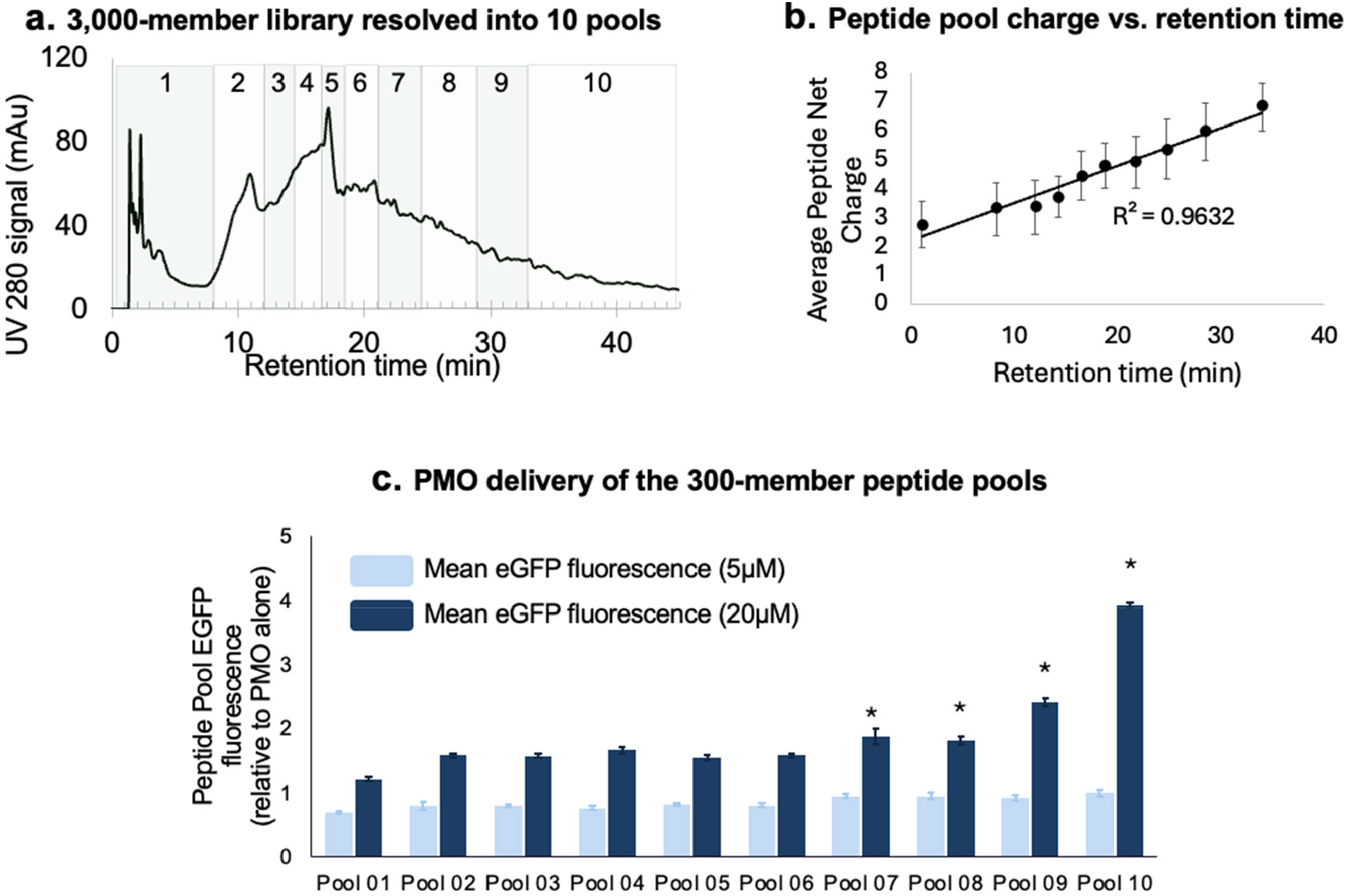
EGFP fluorescence of HeLa 654 cells treated with 5 μM or 20 μM of total PMO-peptide from the resolved library fractions demonstrates cationic pools with appreciable PMO delivery. **(a)** The 3,000-member library was resolved through cation exchange chromatography in ammonium acetate 10% acetonitrile (pH 5) buffer with a 20-500 mM gradient over 50 minutes. All material was collected and pooled based on UV signal at 280 nm. The retention time windows of each pool are shown in green. **(b)** Average net charge of the peptides found within each pool plotted against the mean pool retention time. Each pool was sequenced through nLC-MS/MS and the net charge of each peptide was found. Pool 3 is not shown due to incomplete sequencing coverage. **(c)** EGFP fluorescence of the 10 pools from 3,000-member library resolution. HeLa 654 cells were treated with 5 or 20 µM PMO-peptide for 22 h prior to flow cytometry. Results are given as the mean EGFP fluorescence of cells treated with PMO-peptide relative to the fluorescence of cells treated with vehicle only. Bars represent mean ± SD, N = 3. PMO-peptide fractions 7 through 10 improved PMO delivery significantly over the PMO-654 alone at 20 µM (* = p<0.05 by unpaired *t* test). The resolution of an additional 3,000-member library and a 12,000-member library are shown in **Figure S1**.

The ten library pools demonstrated differential PMO delivery into cells which correlated to the net charge of each pool. These peptide pools which were conjugated to an anti-IVS2-654 PMO cargo were evaluated in a functional cellular assay (HeLA-654). Briefly, the PMO delivery to the nucleus results in corrective splicing of a gene coding for EGFP and is quantified by flow cytometry.^54^ The PMO delivery efficacy of the ten library pools is shown in **Figure 3c** relative to unconjugated PMO. The four most cationic peptide pools (with average charge greater than +5) showed significant delivery efficacy of the PMO cargo, with pool 10 showing the most efficient delivery. Additionally, these peptide pools did not show disruption of the cell membrane up to 20 µM in a lactate dehydrogenase leakage assay (**Figure S1**), demonstrating cell penetration without corresponding membrane lysis or toxicity at the tested concentrations.

The trend between the number of positive charges of the cationic peptides and higher PMO delivery efficacy is reproducible across different library sizes, with smaller libraries showing a higher sequencing efficiency as expected.^52^ An additional 3,000-member peptide library and a 12,000-member peptide library were also resolved into 10 pools using the same cation exchange chromatographic resolution workflow. Again, the pools containing more cationic peptide sequences consistently showed more efficacious PMO delivery into cells (**Figure S2**). With two replicates of this 3,000-member library resolution and MS/MS sequencing, we identified 52 unique peptide sequences with high sequencing fidelity from the highest performing and most cationic pool 10.

### Individual library peptides from the most cationic peptide pools show high PMO delivery

Peptide sequences from pool-10 of the 3,000-member libraries showed high PMO delivery when tested individually. These peptides were named CXP, to designate that they came from cation-exchange pooling of a library. From the sequenced pool-10 peptides, we chose nine peptide sequences (CXP1-CXP9) with the highest sequencing fidelity and lowest arginine content, to avoid arginine-derived peptide toxicity (**Figure 4a**).^18^ In addition, we chose one more peptide (CXP0) from the lower-performing pool 9. The ten peptides were generated via solid-phase peptide synthesis, conjugated to the PMO cargo, and tested for nuclear PMO delivery into HeLa-654 cells using the HeLa 654 splice-correction assay. All ten peptide-PMO conjugates showed enhanced delivery, as measured through GFP expression, relative to unconjugated PMO at 20 µM (**Figure 4b**). The representative peptide from the less effective pool 9, CXP0, did not show PMO delivery on-par with most peptides selected from pool 10, although delivery was still more than 2-fold higher than the unconjugated PMO cargo. In addition, CXP1-CXP4 were also analyzed on cation exchange, which confirmed that their retention time matches the retention window of pool 10 (**Figure S3**). Both the retention time and PMO delivery of the tested peptides confirm that these peptide sequences were among those contributing to the high PMO delivery of the most cationic library pool 10.

**Figure 4:**
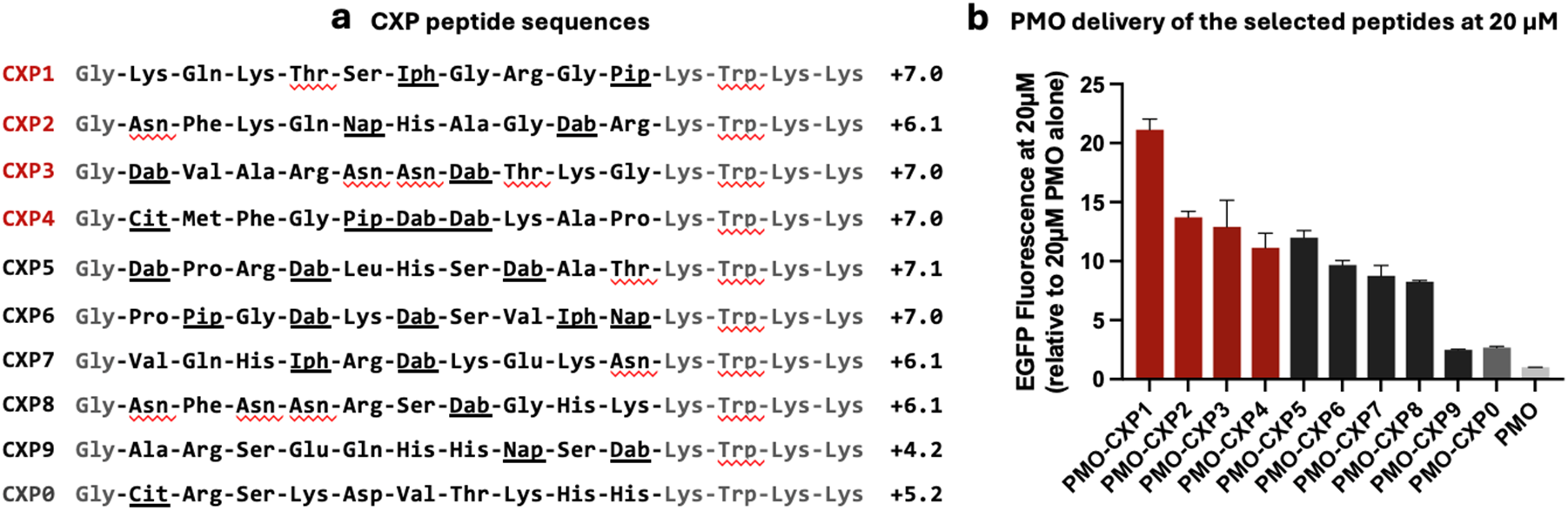
The PMO delivery of the ten peptide sequences chosen from pool 10. **(a)** The sequences of the eleven peptides, wherein the abbreviations and structures of the noncanonical amino acids correspond to those shown in Figure 2. Peptides were conjugated to PMO-DBCO via N-terminal azidopentanoic acid. **(b)** The EGFP fluorescence of each peptide-PMO relative to the fluorescence of 20 μM of PMO 654 alone determined via flow cytometry. HeLa 654 cells were treated with 5 or 20 µM PMO-peptide for 22 h prior to flow cytometry. Results are given as the mean EGFP fluorescence of cells treated with PMO-peptide relative to the fluorescence of cells treated with vehicle only. Bars represent mean ± SD, N = 3. All PMO-peptides, including CXP9 and CXP0, improved PMO delivery significantly over the PMO-654 alone at 20 µM (p<0.05). The PMO delivery of the CXP peptides at 5 µM is shown in **Figure S4**. The membrane toxicity of the CXP peptides at 5 and 20 µM is shown in **Figure S4**.

Four selected peptides from the CXP set (**CXP1-4**) were brought forward for a more in-depth evaluation, due to their high levels of PMO delivery at 5 and 20 µM. The peptides showed dose-dependent delivery of PMO (**Figure 5a**) with no significant membrane toxicity within the tested concentration range, as measured by the release of cytosolic lactate dehydrogenase (LDH) into the surrounding cell media (**Figure 5b**). Because PMO and PMO-peptide conjugates can demonstrate renal toxicity, as kidney is the primary off-target organ for this class of ASO drugs, the PMO-CXP compounds were also tested for membrane permeabilization and acute toxicity in TH-1 human renal proximal tubule epithelial cells.^11,55–56^ Together, these data generated the efficacy and toxicity windows in **Figure S5**. In all cases, the peptides hit their maximum efficacy in delivering PMOs in HeLa 654 cells before they caused appreciable toxicity to the cell membrane in TH-1 cells at the same concentration range. Our results stand in contrast to the higher toxicity of the literature CPP penetratin (LC_50_ = 30 µM), from which the KWKK motif was selected (**Figure 5c**) – thereby validating the favorable toxicity profile of arginine-depleted peptide sequences.

**Figure 5:**
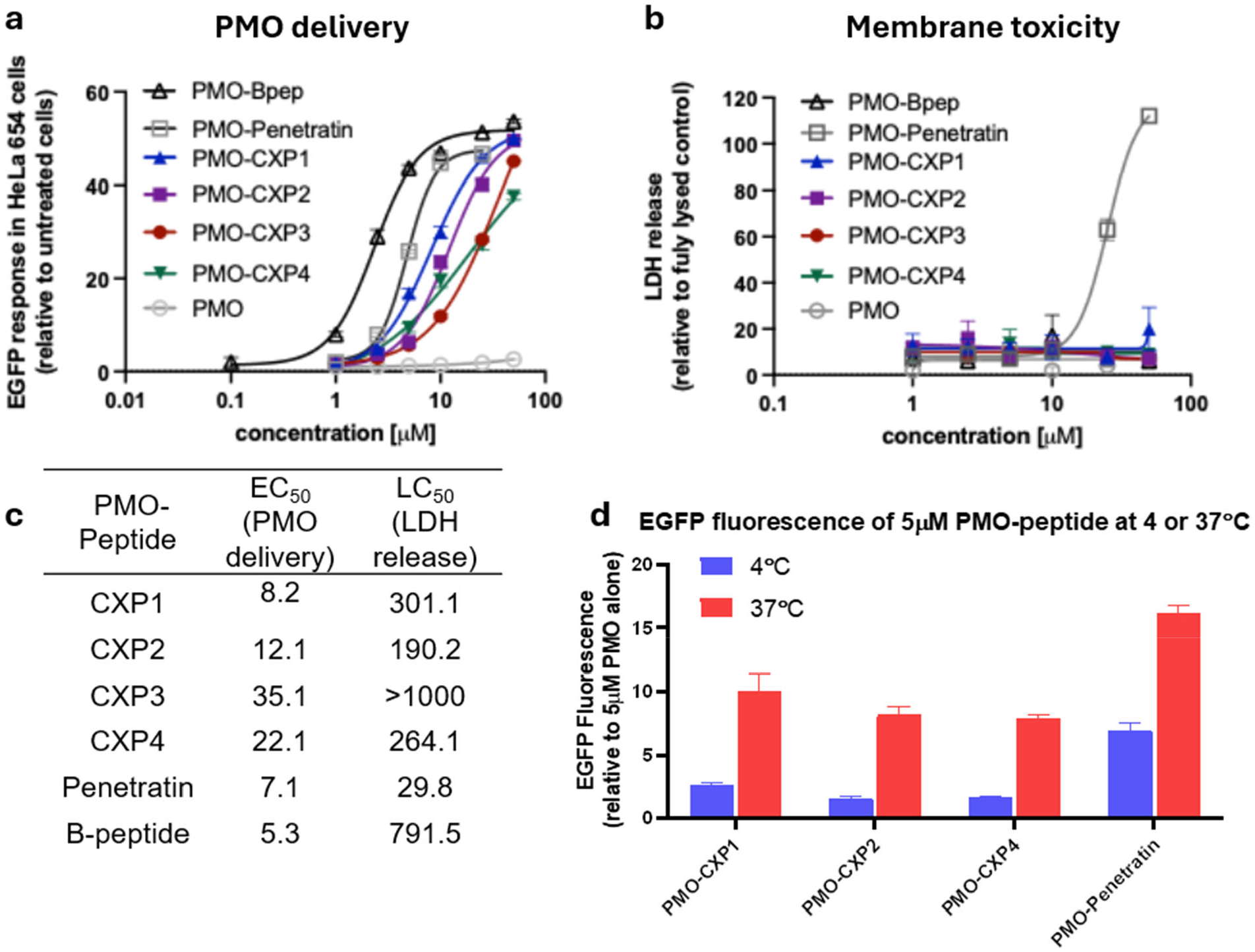
The PMO delivery and membrane toxicity of the 4 strongest CXP peptides at 37 °C and 4 °C. **(a)** HeLa 654 cells were treated with 1, 2.5, 5, 10, 25, or 50 µM PMO-CPP for 22 h prior to flow cytometry. Results are shown as the mean EGFP fluorescence of cells treated with PMO-peptide relative to the fluorescence of cells treated with media only. Bars represent mean ± SD, N = 3. **(b)** Cell supernatant from (a) was tested for LDH release as a measure of membrane permeability. Results are given as percent LDH release above vehicle relative to fully lysed cells. No compound showed statistically significant LDH release above untreated cells. **(c)** Calculated EC_50_ and LC_50_ for each of the PMO-peptides tested. Full dose-response and toxicity graphs are shown in **Figure S5. (d)** The CXP peptides show higher PMO delivery at 37 °C than 4 °C. HeLa 654 cells were treated with 20 µM of PMO-CPP for 2 h at 4 °C or 37 °C. Cells were then washed with PBS and 0.1 mg/mL heparin sulfate, before returning to 37 °C for 20 h prior to flow cytometry. Results are given as the mean EGFP fluorescence of cells treated with PMO-peptide relative to the fluorescence of cells treated with 20 µM of PMO only. Bars represent mean ± SD, N = 3.

We hypothesized that the CXP peptides enter the cell through endocytosis, an energy-dependent mechanism of cellular uptake, in a similar manner to previously studied PMO-peptide conjugates.^48,54^ The hit PMO-CXP peptides were tested for their cell entry mechanism through incubation with HeLa654 cells at either the standard 37 °C or 4 °C, which does not allow for energy-dependent mechanisms of uptake. After incubation for two hours, the cells were washed with PBS and heparin sulfate to remove external PMO-CPP constructs and then returned to the 37 °C incubator to allow for production of the EGFP protein. The 10 library pools from the 12,000-member library demonstrated a marked difference in cell entry between 4 °C and 37 °C in pools 5-10 (**Figure S6a**). In addition, all four CXP peptides show low activity at 4 °C, consistent with an energy-dependent cell uptake mechanism (**Figure 5d, Figure S6b, S6c**).

### Structure-activity relationships demonstrate that peptide sequence is important for PMO delivery while peptide chirality is not

While the chromatographic resolution platform relies on positive charge as a strategy to resolve and enrich peptide libraries, the disparate activities of CXP1-CXP9 demonstrate that positive charge alone is not sufficient for PMO delivery. The sequence of the selected CXP peptides plays an important role for PMO delivery, while peptide chirality does not. To demonstrate this, we synthesized scrambled versions of four CXP peptides. These scrambled peptides contain the same amino acids as the parent sequence, but in a random order. This arrangement breaks up both the penetratin-specific “KWKK” motif and the specific charge patterning across the peptide. Across all four peptides, the scrambled versions were significantly less efficacious for PMO delivery (**Figure 6a**). In contrast, the mirror-image peptides, which have the same peptide sequence but opposite chirality, showed no significant decrease in EGFP fluorescence relative to the native peptide (**Figure 6b**). While the scrambled peptides demonstrate that the peptide sequence is necessary for PMO delivery, chirality does not hold the same significance. This underscores the importance of not only the positive charges in these peptides, but also their specific patterning across each sequence. The enrichment platform allows for the discovery of peptides with high positive charge without restricting the charged positions or otherwise limiting the sequence diversity outside of the fixed motif, as demonstrated by the scrambled and D-peptides.

**Figure 6:**
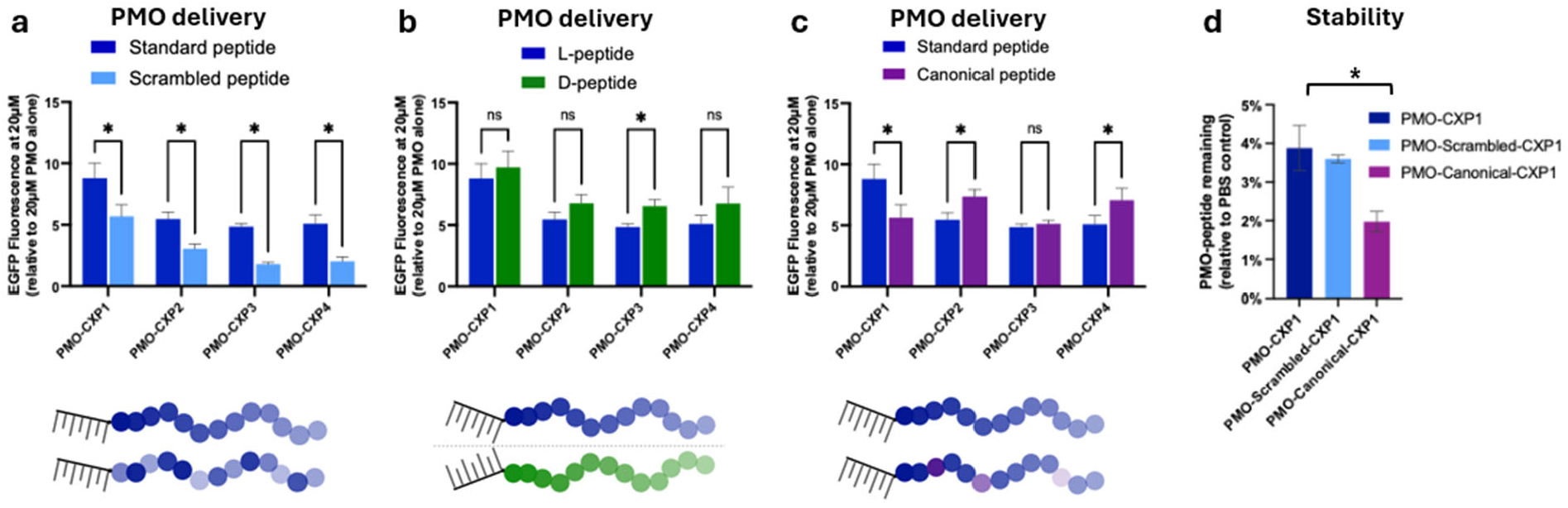
The PMO delivery and stability of scrambled, D-, or canonical variants of the CXP peptides. HeLa 654 cells were treated with 20 µM of PMO-CPP for 22 h prior to flow cytometry. Results are given as the mean EGFP fluorescence of cells treated with PMO-peptide relative to the fluorescence of cells treated with 20 µM of PMO only. Bars represent mean ± SD, N = 3. **(a)** The PMO delivery of PMO-CXP peptides and their scrambled variants. (**b)** The PMO delivery of PMO-CXP peptides and their mirror-image D-variants. **(c)** The PMO delivery of PMO-CXP peptides and their canonical variants. **(d)** The mouse serum stability of PMO-CXP1 and variants. PMO-peptides were incubated in 25% mouse serum in PBS or 100% PBS for 6 h. Remaining peptide was measured by LC-MS and quantified relative to the PBS-incubated samples. Data for remaining timepoints (0, 3, and 12 h) are shown in **Figure S8** (* = p<0.05, ** = p<0.01 ***=p<0.001 by unpaired *t* test).

The noncanonical amino acids in the peptide libraries have varying effects on cell penetration compared to their canonical analogs. To understand the role of these residues, canonical versions of the four hit CXP peptides were also synthesized. These canonical variants contain canonical amino acids in place of the parent sequences’ abiotic residues. For example, 4-iodophenylalanine was replaced with phenylalanine and 2,4-diaminobutyric acid was replaced with a lysine. While in some cases these substitutions have little effect on the PMO delivery, for CXP1 there is a statistically significant decrease in EFGP fluorescence after the noncanonical amino acids were substituted (**Figure 6c**). In contrast, CXP2 and CXP4 experienced enhanced PMO delivery at 20 µM after the noncanonical residues were replaced. The non-natural amino acids in the library design, therefore, affect not only the chemical diversity, but also PMO delivery of the peptide sequences.

The noncanonical residues in CXP1 conferred both cell penetration and additional proteolytic stability when compared to the canonical version of the peptide. The canonical and parent variants of PMO-CXP1 were incubated in mouse serum for 24 h. By 6 h, there was significantly less (~50% less) canonical CXP1 remaining in solution compared to either the parent peptide or the scrambled sequence (**Figure 6d, Figure S7**). This outcome suggests that the noncanonical amino acids also impart a degree of protease stability to the peptide constructs, which has also been previously demonstrated.^27,37^

### CXP-1 to CXP-4 retain high PMO delivery and wide tolerability in vivo across multiple PMO cargoes

To validate our in vitro findings, we sought to evaluate the efficacy of CXP1-CXP4 in vivo. The EGFP-654 murine model, described previously,^57^ which encodes enhanced green fluorescence protein (EGFP) that is interrupted by an aberrantly spliced mutant intron of the human beta-globin gene, was utilized to assess splice correction, similar to the HeLa-654 splice-correction assay. The uptake of the splice-switching PMO corrects the aberrant splicing, producing EGFP protein, and thus EGFP fluorescence can be used as a proxy for exon skipping. We conjugated the anti-IVS2-654 PMO to CXP1, B-peptide and R_6_G to assess and compare the efficacy of these PMO-CPP constructs. This assessment compares CPPs that differ significantly in the total number of arginine residues despite similar charge. While the CXP1 peptide is depleted in arginine content (#R = 1, net charge = +7), both B-peptide (#R = 8, net charge = +8) and R_6_G (#R = 6, net charge = +6) are enriched in arginine residues.^31,58,59^

Mice were administered the PMO-CPP compounds intravenously at doses ranging from 1 to 40 mg/kg for CXP1, 2.5 to 30 mg/kg for B-peptide, 3 to 160 mg/kg for R_6_G, and EGFP expression was measured in tissues (quadriceps, heart and kidney) at seven days post-dosing. We observed that PMO-CXP1 resulted in similar efficacy to PMO-B-peptide in both quadriceps and heart and was significantly higher than that of the PMO-R_6_G counterpart (**Figure 7a, 7b**). Notably, PMO-CXP1 exhibited superior efficacy, with higher EGFP fluorescence at lower doses (≤ 10 mg/kg), while B-peptide only approached comparable efficacy at the 20 and 30 mg/kg doses. PMO-CXP1 had an ED_50_ of 12.5 mg/kg in the quadriceps, while that of B-peptide was 14.6 mg/kg and PMO-R_6_G had a much higher ED_50_ of 80.6 mg/kg (**Figure 7a**). A similar trend in efficacy was observed in the heart muscles (**Figure 7b**). These findings suggest both PMO-CXP1 and PMO-B-peptide exhibit a similar efficacy profile with both significantly more potent than PMO-R_6_G. Additionally, PMO-CXP1 outperforms PMO-B-peptide at lower doses despite having fewer arginine residues and lower net charge.

**Figure 7:**
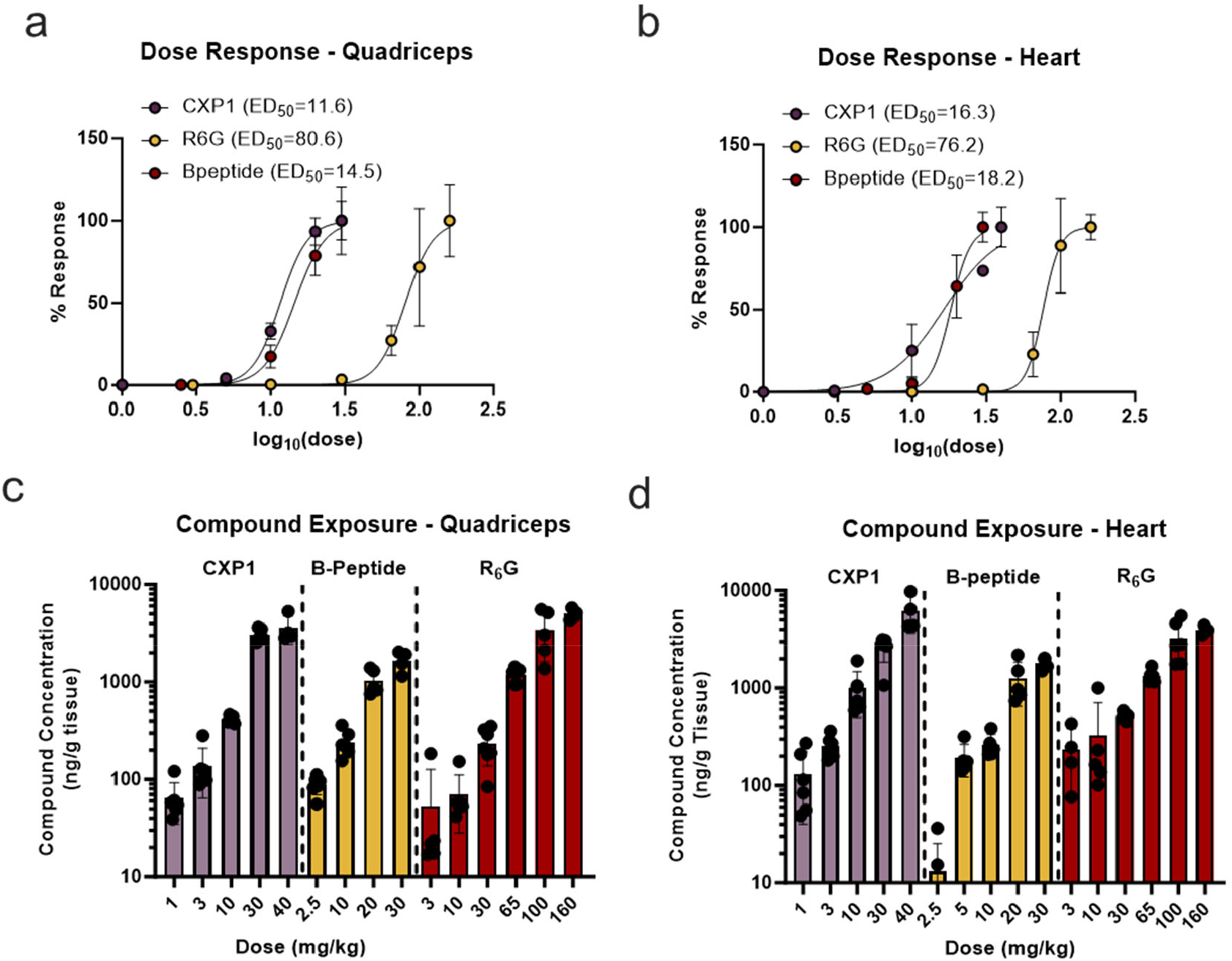
Evaluation of efficacy and exposure of CXP1 versus arginine rich peptides. EGFP-654 transgenic mice were treated with various doses of PMO-CPP for 7 days prior to tissue harvesting. The dose response in quadricep (**a)** and heart **(b)** for PMO-CXP1, PMO-R6G and PMO-B-peptide is shown as the log_10_ dose versus the normalized percent response with a variable slope nonlinear regression to determine the ED_50_ value. Compound exposure for quadricep **(c)** and heart **(d)**. Results are expressed as the mean ± SD.

Next, we measured tissue exposure of our PMO-CPPs compounds using an Enzyme-Linked Oligonucleotide Hybridization Assay (ELOHA). This assay detects and quantifies specific oligonucleotides using an enzyme-conjugated complementary probe that hybridizes with the nucleic acid sequence. The bound probes are detected in the assay via an enzyme-catalyzed reaction, producing a measurable fluorescence signal. Using an IVS2-654 PMO probe, we measured the concentration of PMOs in quadricep and heart. It was observed that conjugation of the PMO with CXP1 resulted in increased delivery to striated muscles tissues compared to B-peptide and R_6_G at all comparable doses (**Figure 7c, 7d**).^60,61^ Overall, our findings from the EGFP-654 transgenic mice suggest that arginine-depleted peptides, such as CXP1, can serve as efficient and well-tolerated payload carriers achieving high therapeutic efficacy at relatively low doses.

Next, we sought to assess the efficacy of CXP1-CXP4 conjugated PMOs in the same transgenic mice model. EGFP-654 mice were dosed intravenously at two doses: 12 mg/kg, the ED_50_ of PMO-CXP1, and at 35 mg/kg, a high dose where maximal efficacy is observed. Assessment of efficacy at 7 days post-dosing, showed that PMO-CXP4 resulted in a significant increase in EGFP fluorescence in the quadriceps at 12 mg/kg while all four PMO-CPPs demonstrated similar efficacy in the heart at the same dose (**Figure 8a, 8b**). At the 35 mg/kg dose, no significant difference in EGFP expression was observed among the four PMO-CPPs in both skeletal and cardiac muscles tissues. (**Figure 8c, 8d**). Considering the overall results, no clear trend was observed among the 4 PMO-CPP conjugates.

**Figure 8:**
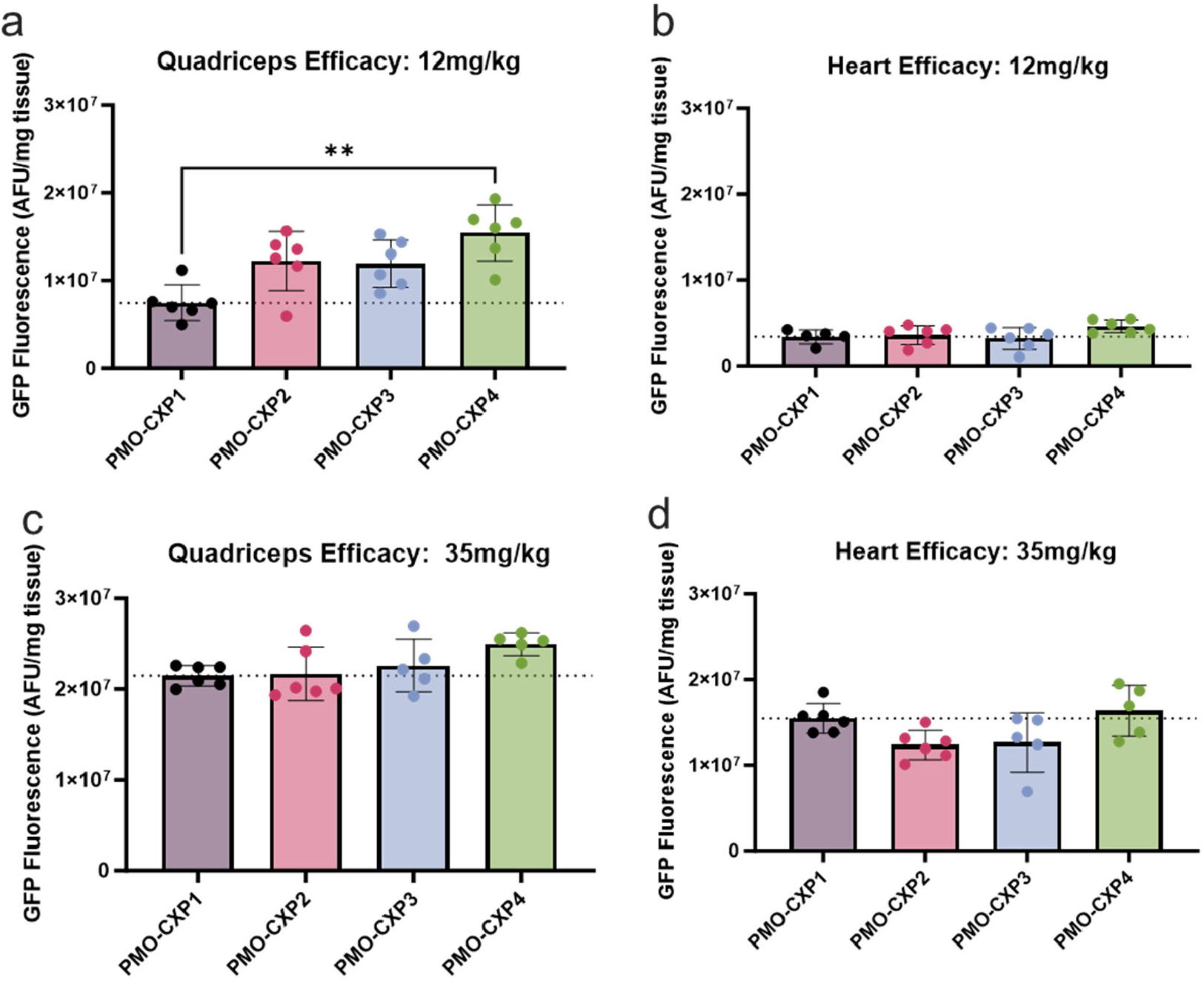
Evaluation of CXP peptides in EGFP-654 transgenic mice. Efficacy of PMO-CXP1, PMO-CXP2, PMO-CXP3 and PMO-CXP4 were measured 7 days post-dosing either at 12 mg/kg or 35 mg/kg. Efficacy results from the quadriceps **(a)-(b)** and heart **(c)-(d)** are shown. Data is represented as EGFP fluorescence normalized to tissue weight. Results are expressed as the mean with error bars (SD). The Brown-Forsythe and Welch ANOVA tests were used to compare data sets. **P<0.001.

Lastly, we assessed the efficacy of CXP1 compared to R_6_G in a DMD-relevant disease mouse model, the *mdx* mouse. The *mdx* mouse model contains a point mutation in exon 23 of the dystrophin gene, which introduces a premature stop codon and causes a dystrophic phenotype.^62–63^

We observed that PMO-CXP1 demonstrated greater exon skipping efficacy than the R_6_G counterpart in both the quadriceps and the heart muscles (**Figure 9a, 9b**), a trend that matches the results from the EGFP-654 transgenic mouse model. The PMO-R_6_G demonstrated a higher ED_50_(~4x) value than that of PMO-CXP1 indicating its lower potency. The slightly enhanced efficacy of the PMO-CPPs in this mouse model, compared to the non-diseased EGFP-654 mouse model, can be attributed to membrane instability due to constant degeneration/regeneration of muscle fibers in the atrophic state.^64–65^ In terms of compound accumulation, we observed a similar trend where PMO-CXP1 had slightly higher tissue exposure in both striated muscles as determined using ELOHA (**Figure 9c-d**).

**Figure 9:**
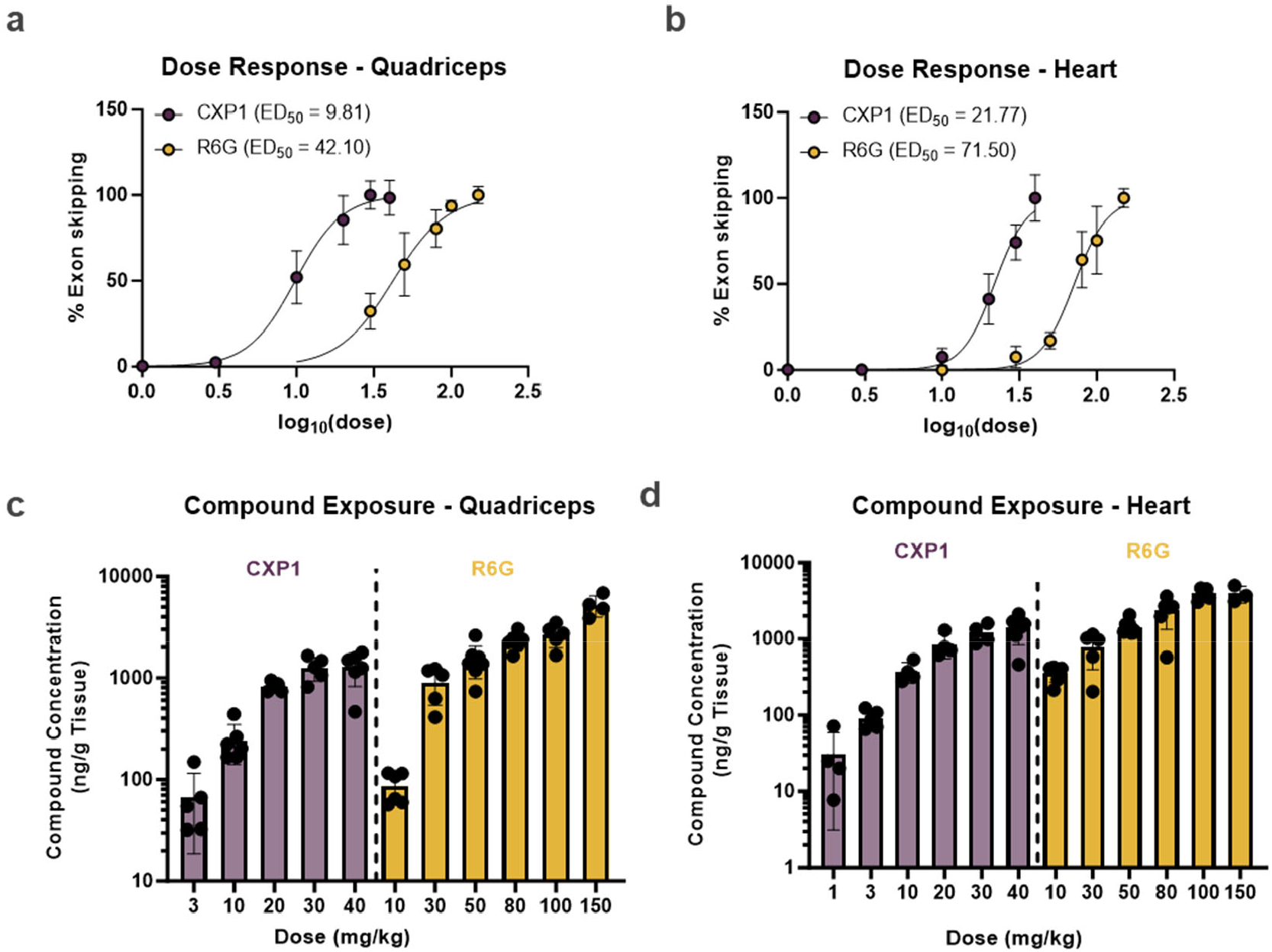
Comparison of efficacy and exposure of CXP1 versus R6G in *mdx* mice. *mdx* mice were treated with various doses of PMO-CPP for 7 days prior to tissue harvesting. The dose response in quadricep (**a)** and heart **(b)** for PMO-CXP1 and PMO-R6G is shown as the log_10_ dose versus the normalized percent exon skipping with a variable slope nonlinear regression to determine the ED_50_ value. Compound exposure for quadricep **(c)** and heart **(d)**. Results are expressed as the mean with error bars (SD).

Additionally, we collected terminal serum from the *mdx* mice at 7 days post-dosing to assess muscle damage biomarkers. While traditionally utilized as serum biomarkers for acute liver injury, aspartate transaminase (AST) and alanine transaminase (ALT) are also robustly expressed in skeletal muscle and elevated in the serum of DMD patients.^66,67^ Along with creatine kinase (CK) and lactate dehydrogenase (LDH), these enzymes serve as additional markers for assessing skeletal muscle damage. Blood Urea Nitrogen (BUN) is a nitrogenous end-product of metabolism and when normalized to urinary creatinine to control for hydration status, it serves as an indicator of renal health.^68^

Levels of creatine kinase and LDH exhibited a dose-dependent reduction with levels of both biomarkers recapitulating the levels of the wild-type control mice after a single 20 mg/kg dose (**Figures 10a-10c**). This indicates a reversal in tissue damage upon treatment with PMO-CPPs. Similarly, the liver biomarkers AST and ALT all showed dose dependent reductions returning to normal levels after a single 10 mg/kg dose (**Figures 10d, 10e**). The BUN/creatinine ratio did not show any significant changes after dosing with PMO-CXP1 in any of the dose groups when compared to the PBS treated group, indicating no kidney associated adverse events occur due to treatment with either of the PMO-CPPs (**Figure 10f**).

**Figure 10:**
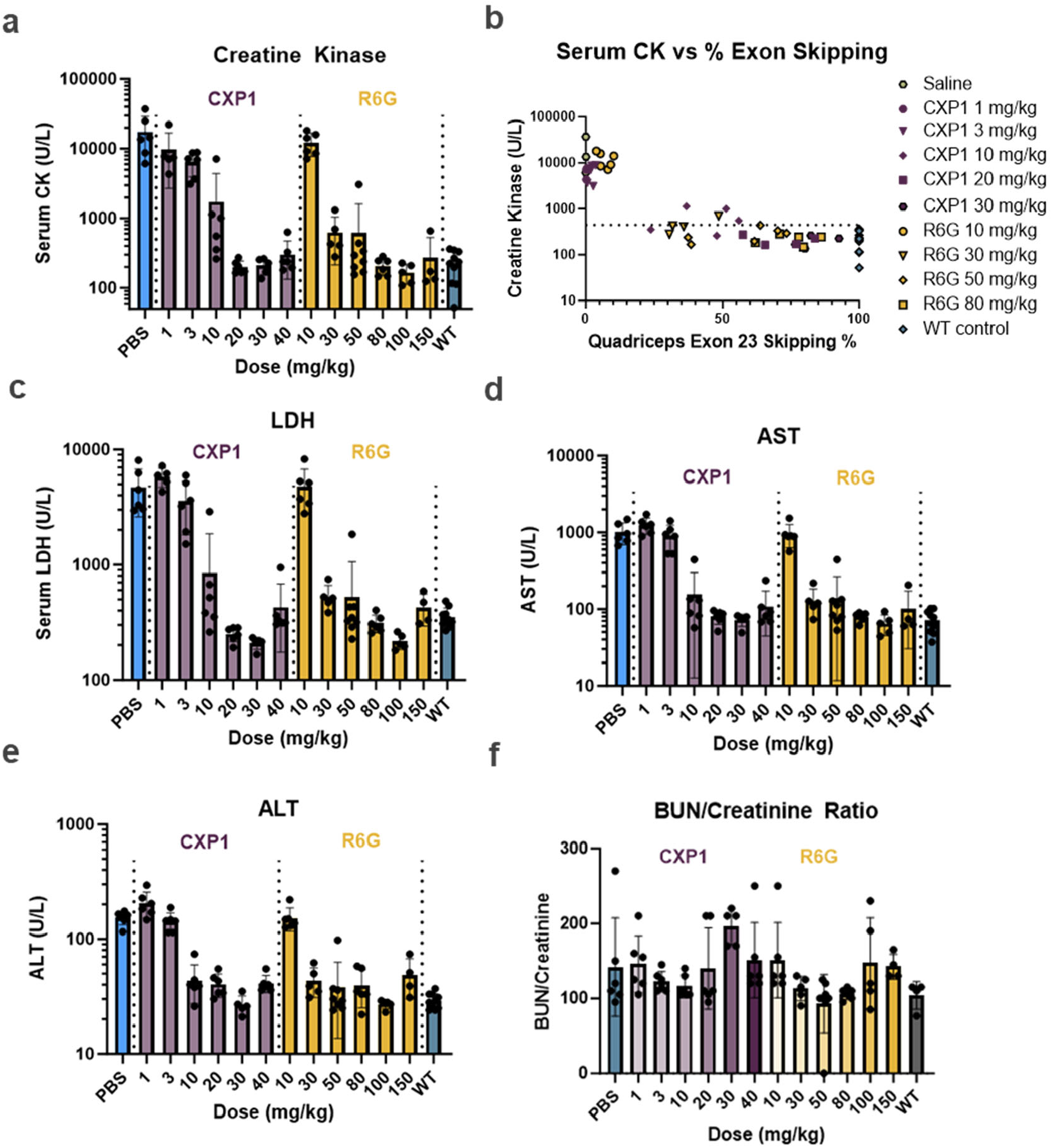
Evaluation of serum biomarkers of DMD pathology. Terminal serum from *mdx* mice was collected by cardiac puncture at 7 days post injection and biomarkers of skeletal muscle damage were quantified. Creatine kinase levels **(a)**, a representative correlation between creatine kinase versus exon skipping **(b)**, LDH **(c)**, AST **(d)**, ALT **(e)**, and BUN/Creatinine ratio Creatine Kinase levels **(f)** for *mdx* mice treated with PMO-CXP1, and PMO-R6G. PBS-treated and WT animals were included as controls. Results are expressed as the mean with error bars (SD).

## Discussion

Novel CPPs for PMO delivery have historically been discovered through careful optimization of known peptides or rational design, processes which can be time-consuming or cumbersome.^15–19, 28–30^ In the case of abiotic peptides, the incorporation of noncanonical amino acids exponentially expands the library combinatorial diversity, making brute-force screening impractical without a tailored strategy. We have previously reported a library screening method to discover novel, effective PMO-CPPs directly from the cell cytosol.^48^ Here we describe an orthogonal methodology that allows for selection and in-cell enrichment of PMO-CPPs from an even larger and higher-diversity library without additional tags between the peptide and PMO cargo. Resolving these large libraries by charge and subsequent evaluation in a standardized functional readout assay allows us to enrich for more cationic and highly cell-permeable peptides, demonstrated by the high PMO delivery from the most cationic peptide pools across three trials with different library sizes. Tellingly, the screen did not simply rediscover known CPPs—the classical peptide penetratin, for example, was not identified among the top hits despite its partial motif being present in the library design. This outcome indicates that previously unexplored sequence space was mined successfully, yielding entirely new CPP candidates rather than variations of known archetypes. This method presents a flexible alternative to more focused, fixed, or directed library designs and individual peptide synthesis.

The chromatographic resolution method demonstrated a clear correlation between more positively charged peptide pools and higher PMO delivery efficacy. This correlation has been previously demonstrated in the literature and shown here to be present when the peptide pools average a charge of +5 or above.^30,34–36^ Mechanistically, positively charged amino acid residues have been associated with both triggering endocytosis and peptide endosomal escape,^69–70^ consistent with the mechanism of entry demonstrated for both the CXP peptides and the overall PMO-peptide library. The CXP peptides are also enriched with lysine residues, a positively charged amino acid that, unlike arginine, lacks a guanidinium group. Lysine is not a trivial substitute for arginine in cationic peptide design, as its side chain chemistry and biological interactions differ profoundly. This structural difference results in a lower p*K*_a_ and reduced basicity, limiting the ability of lysine to engage in strong electrostatic interactions with phospholipid bilayers.^71–73^ In contrast, arginine residues can form bidentate hydrogen bonds and salts bridges with negatively charged membrane components, whereas lysine typically forms only monodentate interactions.^71–74^ These biophysical distinctions are widely recognized as key factors underlying the reduced membrane translocation efficiency observed when lysine substitutes for arginine in arginine-rich CPPs. We hypothesize that this mode of cellular entry resulting from the lysine enrichment in CXP peptides, may contribute to a subtly distinct uptake mechanism that reduces their toxicity profile by minimizing membrane disruption relative to direct translocation mechanisms.^75^

The chromatographic enrichment platform produced a set of four PMO-CPP constructs with high PMO delivery and broad tolerability. The noncanonical residues contained within these peptides enhanced their sequence diversity and stability to serum proteases. The mirror-image D-peptides retained their PMO delivery, while scrambled versions of the CXP peptides showed lower cell penetration. These scrambled peptides retain high net charge (+7) but lost both the specific charge patterning and their PMO delivery activity. The additional serum-stability and efficacy conferred by the noncanonical amino acids in the CXP-1 peptide further underscore the utility of this and other abiotic peptides such as the Mach and P6 peptides previously developed in our laboratory as in vivo delivery vehicles.^30,76^

Taken together, these variants demonstrate both the number and the position of charged residues within the sequence matters. The low impact of chirality on PMO delivery corroborates our previous studies with a set of PMO conjugates with established L- and D-CPPs, in which we demonstrated that for most CPPs tested, chirality had little impact on in-vitro cell penetration.^77^ The importance of charge spacing has also been discovered empirically through the rational design of peptides like Pip-6 and B-Peptide, which were specifically developed to space positive charges with more hydrophobic regions.^21–23^ The chromatographic resolution platform not only recapitulated these previous results but also demonstrated new sequences and patterns that can be used for charge spacing for future CPP developments.

B-peptide has long served as a benchmark for PMO delivery efficacy, in large part due to its arginine-rich nature, which facilities efficient cellular delivery and robust exon-skipping activity but also triggers dose-limiting side effects in vivo.^31^ In contract, the emergence of CXP1 as a delivery vector is therefore notable, it matches or even surpasses the potency of B-Peptide in certain conditions, despite depleted numbers of arginine residues, indicating that potent PMO delivery can be achieved without guanidinium-facilitated cellular internalization. In direct comparative studies, PMO-CXP1 modestly outperforms the arginine-rich B-Peptide at lower doses in teams of efficacy and tissue exposure. The enhanced delivery efficiency of CXP1 has meaningful implications, since more effective uptake at lower doses suggests the potential for decreased overall drug burden, thereby reducing the likelihood of dose-dependent tolerability concerns and enables lower and less frequent dosing regimens. Prior clinical experiences with CPP-conjugated PMOs support this concept, since CPP-mediated delivery of PMOs, achieved robust and durable exon skipping with a reduced dosing frequency compared to an unconjugated PMO drug. CXP1 extends this by yielding a 4–7-fold increase in activity. Treatment of *mdx* mice with PMO-CXP1 reduced biomarkers of tissue stress and pathology, supporting that effective exon skipping can be achieved with a more favorable dosing profile. ^10,26^

The PMO-CXP1 conjugate, unlike many other ASO delivery strategies,^3^ showed strong translational ability from in vitro to in vivo experiments. It additionally highlights that an arginine depleted peptide is effective in delivering different PMO payloads in both healthy (*EGFP-654*) and diseased (*mdx*) mouse models. We additionally evaluated the top four peptide (CXP1-4) in the transgenic EGFP-654 model where we did not observe a significant difference at two doses (12 and 35 mg/kg) in both skeletal and cardiac muscles except for CXP4 slightly outperforming the rest at 12 mg/kg in quadriceps.

Overall, this chromatographic enrichment platform can enable broader applications across the field of CPP discovery. In this work, we not only discovered a set of new peptide sequences but also gained a better understanding of the relationship between positive charge and delivery of a PMO cargo. By choosing additional, orthogonal methods of fractionation in the future, we can understand new trends between the biophysical character of a peptide and its ability to deliver the cargo of choice. Current investigations in our lab are focusing on applying this platform to additional therapeutically relevant cargoes and discovering new biophysical properties important for cell penetration.

## Acknowledgements

This research was funded by Sarepta Therapeutics. C.E.F. (4000057441) and C.K.S. (4000057398) acknowledge the National Science Foundation Graduate Research Fellowship (NSF Grant No. 1122374) for research support. We acknowledge support from the Swanson Biotechnology Center Flow Cytometry Facility at the Koch Institute for Integrative Cancer Research at MIT through the use of their flow cytometers (NCI Cancer Center Support Grant P30 CA014051).

## Competing Interests

The authors declare the following competing financial interest(s): B.L.P. is a co-founder and/or member of the scientific advisory board of several companies focusing on the development of protein and peptide therapeutics. MIT and Sarepta Therapeutics filed a joint patent application on the products described in this study. A.B., C.D.G., A.M.W., E.G.T., K.Y.M. and V.G. are currently employed by Sarepta Therapeutics.

## Data and materials availability

All data generated during this study are available either in the main text or supplementary materials.

## References

(1) Lauffer, M. C.; van Roon-Mom, W.; Aartsma-Rus, A. Possibilities and Limitations of Antisense Oligonucleotide Therapies for the Treatment of Monogenic Disorders. Communications Medicine 2024, 4, 1–11.

(2) Belgrad, J.; Fakih, H. H.; Khvorova A. Nucleic Acid Therapeutics: Successes, Milestones, and Upcoming Innovation. Nucleic Acid Ther. 2024, 34, 52–72.

(3) Anand, P.; Zhang, Y.; Patil, S.; Kaur, K. Metabolic Stability and Targeted Delivery of Oligonucleotides: Advancing RNA Therapeutics Beyond The liver. J. Med. Chem. 2025, 68, 6870–6896.

(4) Falzarano, M. S.; Scotton, C.; Passarelli, C.; Ferlini, A. Duchenne Muscular Dystrophy: From Diagnosis to Therapy. Molecules 2015, 20, 18168–18184.

(5) Findlay, A. R.; Wein, N.; Kaminoh, Y.; Taylor, L. E.; Dunn, D. M.; Mendell, J. R.; King, W. M.; Pestronk, A.; Florence, J. M.; Mathews, K. D.; Finkel, R. S.; Swoboda, K. J.; Howard, M. T.; Day, J. W.; McDonald, C.; Nicolas, A.; Le Rumeur, E.; Weiss, R. B.; Flanigan, K. M. Clinical Phenotypes as Predictors of the Outcome of Skipping around *DMD* Exon 45. Ann Neurol 2015, 77, 668–674.

(6) Charleston, J. S.; Schnell, F. J.; Dworzak, J.; Donoghue, C.; Lewis, S.; Chen, L.; Young, G. D.; Milici, A. J.; Voss, J.; DeAlwis, U.; Wentworth, B.; Rodino-Klapac, L. R.; Sahenk, Z.; Frank, D.; Mendell, J. R. Eteplirsen Treatment for Duchenne Muscular Dystrophy: Exon Skipping and Dystrophin Production. Neurology 2018, 90, e2146–e2154.

(7) Gupta, S.; Sharma, S. N.; Kundu, J.; Pattanayak, S.; Sinha, S. Morpholino Oligonucleotide-Mediated Exon Skipping for DMD Treatment: Past Insights, Present Challenges and Future Perspectives. J Biosci 2023, 48, 1–17.

(8) Moulton, H. M.; Moulton, J. D. Morpholinos and Their Peptide Conjugates: Therapeutic Promise and Challenge for Duchenne Muscular Dystrophy. Biochimica et Biophysica Acta - Biomembranes. Elsevier December 1, 2010, pp 2296–2303.

(9) McClorey, G.; Banerjee, S. Cell-Penetrating Peptides to Enhance Delivery of Oligonucleotide-Based Therapeutics. Biomedicines 2018, 6, 51.

(10) Sheikh, O.; Yokota, T. Pharmacology and Toxicology of Eteplirsen and SRP-5051 for DMD Exon 51 Skipping: An Update. Arch Toxicol 2022, 96, 1–9.

(11) Derossi, D.; Joliot, A. H.; Chassaing, G.; Prochiantz, A. The Third Helix of the Antennapedia Homeodomain Translocates through Biological Membranes. J Biol Chem 1994, 269, 10444–10450.

(12) Bajiya, N.; Najrin, S; Kumar, P.; Choudhury, S.; Tomer, R.; Raghava, G. P. S. CPPsite3: An updated large repository of experimentally validated cell-penetrating peptides. Drug Discov Today 2025, 30, 104421.

(13) Preto, A. J.; Caniceiro, A. B.; Duarte, F.; Fernandes, H.; Ferreira, L.; Mourão, J.; Moreira, I. S. POSEIDON: Peptidic Objects Sequence-Based Interaction with Cellular Domains: A New Database and Predictor. J Cheminform 2024, 16, 1–13.

(14) El-Andaloussi, S.; Järver, P.; Johansson, H. J.; Langel, Ü. Cargo-Dependent Cytotoxicity and Delivery Efficacy of Cell-Penetrating Peptides: A Comparative Study. Biochemical Journal 2007, 407, 285–292.

(15) Wolfe, J. M.; Fadzen, C. M.; Choo, Z. N.; Holden, R. L.; Yao, M.; Hanson, G. J.; Pentelute, B. L. Machine Learning to Predict Cell-Penetrating Peptides for Antisense Delivery. ACS Cent Sci 2018, 4, 512–520.

(16) Kauffman, W. B.; Guha, S.; Wimley, W. C. Synthetic Molecular Evolution of Hybrid Cell Penetrating Peptides. Nat Commun 2018, 9, 2568.

(17) Dastpeyman, M.; Sharifi, R.; Amin, A.; Karas, J. A.; Cuic, B.; Pan, Y.; Nicolazzo, J. A.; Turner, B. J.; Shabanpoor, F. Endosomal Escape Cell-Penetrating Peptides Significantly Enhance Pharmacological Effectiveness and CNS Activity of Systemically Administered Antisense Oligonucleotides. Int J Pharm 2021, 599, 120398.

(18) Tünnemann, G.; Ter-Avetisyan, G.; Martin, R. M.; Stöckl, M.; Herrmann, A.; Cardoso, M. C. Live-cell analysis of cell penetration ability and toxicity of oligo-arginines. J Pept Sci. 2008, 14, 469–76.

(19) Abes, R.; Moulton, H. M.; Clair, P.; Yang, S. T.; Abes, S.; Melikov, K.; Prevot, P.; Youngblood, D. S.; Iversen, P. L.; Chernomordik, L. V.; Lebleu, B. Delivery of Steric Block Morpholino Oligomers by (R-X-R)4 Peptides: Structure–Activity Studies. Nucleic Acids Res 2008, 36, 6343–6354.

(20) Hong M. Moulton, Michelle H. Nelson, Susie A. Hatlevig, Muralimohan T. Reddy, and Patrick L. Iversen. Bioconjugate Chemistry 2004, 15, 290–299.

(21) Wu B, Moulton HM, Iversen PL, Jiang J, Li J, Li J, Spurney CF, Sali A, Guerron AD, Nagaraju K, Doran T, Lu P, Xiao X, Lu QL. Effective rescue of dystrophin improves cardiac function in dystrophin-deficient mice by a modified morpholino oligomer. Proc Natl Acad Sci U. S. A. 2008, 105, 14814–9.

(22) Jearawiriyapaisarn N, Moulton HM, Sazani P, Kole R, Willis MS. Long-term improvement in mdx cardiomyopathy after therapy with peptide-conjugated morpholino oligomers. Cardiovasc Res. 2010, 85, 444–53.

(23) Li, Q.; Xu, M.; Cui, Y.; Huang, C.; Sun, M. Arginine-Rich Membrane-Permeable Peptides Are Seriously Toxic. Pharmacol Res Perspect 2017, 5, e00334.

(24) Saar, K.; Lindgren, M.; Hansen, M.; Eiríksdóttir, E.; Jiang, Y.; Rosenthal-Aizman, K.; Sassian, M.; Langel, Ü. Cell-Penetrating Peptides: A Comparative Membrane Toxicity Study. Anal Biochem 2005, 345, 55–65.

(25) Herce, H. D.; Garcia, A. E.; Litt, J.; Kane, R. S.; Martin, P.; Enrique, N.; Rebolledo, A.; Milesi, V. Arginine-Rich Peptides Destabilize the Plasma Membrane, Consistent with a Pore Formation Translocation Mechanism of Cell-Penetrating Peptides. Biophys J 2009, 97, 1917.

(26) Haque, S. U.; Kohut, M.; Yokota, T. Comprehensive review of adverse reactions and toxicology in ASO-based therapies for Duchenne Muscular Dystrophy: From FDA-approved drugs to peptide-conjugated ASO. Curr. Res. Toxicol. 2024, 7, 100182.

(27) Ding, Y.; Ting, J. P.; Liu, J.; Al-Azzam, S.; Pandya, P.; Afshar, S. Impact of Non-Proteinogenic Amino Acids in the Discovery and Development of Peptide Therapeutics. Amino Acids 2020, 52, 1207.

(28) Porosk, L.; Gaidutsik, I.; Langel, U. Approaches for the discovery of new cell-penetrating peptides. Expert Opin Drug Discov. 2021, 16, 553–565.

(29) Porosk, L.; Langel, U. Approaches for evaluation of novel CPP-based cargo delivery sytems. Front. Pharmacol. 2022, 13, 1056467.

(30) Schissel, C. K.; Mohapatra, S.; Wolfe, J. M.; Fadzen, C. M.; Bellovoda, K.; Wu, C. L.; Wood, J. A.; Malmberg, A. B.; Loas, A.; Gómez-Bombarelli, R.; Pentelute, B. L. Deep Learning to Design Nuclear-Targeting Abiotic Miniproteins. Nat Chem 2021, 13, 992–1000.

(31) Jearawiriyapaisarn, N.; Moulton, H. M.; Buckley, B.; Roberts, J.; Sazani, P.; Fucharoen, S.; Iversen, P. L.; Kole, R. Sustained Dystrophin Expression Induced by Peptide-Conjugated Morpholino Oligomers in the Muscles of Mdx Mice. Mol Ther 2008, 16, 1624.

(32) Klein, A. F.; Varela, M. A.; Arandel, L.; Holland, A.; Naouar, N.; Arzumanov, A.; Seoane, D.; Revillod, L.; Bassez, G.; Ferry, A.; Jauvin, D.; Gourdon, G.; Puymirat, J.; Gait, M. J.; Furling, D.; Wood, M. J. A. Peptide-conjugated oligonucleotides evoke long lasting myotonic dystrophy correction in patient derived cells and mice. J Clin Invest. 2019, 129, 4739–4744.

(33) Gait, M. J.; Arzumanov, A. A.; McClorey G.; Godfrey, C.; Betts, C.; Hammond, S.; Wood, M. J. A. Cell-penetrating Peptide Conjugate of Steric Blocking Oligonucleotides as Therapeutics for Neuromuscular Diseases from a Historical Perspective to Current Prospects of Treatment. Nucleic Acid Ther. 2019, 29, 1–12.

(34) Dastpeyman, M.; Sharifi, R.; Amin, A.; Karas, J. A.; Cuic, B.; Pan, Y.; Nicolazzo, J. A.; Turner, B. J.; Shabanpoor, F. Endosomal Escape Cell-Penetrating Peptides Significantly Enhance Pharmacological Effectiveness and CNS Activity of Systemically Administered Antisense Oligonucleotides. Int J Pharm 2021, 599, 120398.

(35) Takada, H.; Tsuchiya, K.; Demizu, Y. Helix-Stabilized Cell-Penetrating Peptides for Deliveryof Antisense Morpholino Oligomers: Relationships among Helicity, CellularUptake, and Antisense Activity. Bioconjug Chem 2022, 33, 1311–1318.

(36) Li, X.; Kheirabadi, M.; Dougherty, P. G.; Kamer, K. J.; Shen, X.; Estrella, N. L.; Peddigari, S.; Pathak, A.; Blake, S. L.; Sizensky, E.; Del Genio, C.; Gaur, A. B.; Dhanabal, M.; Girgenrath, M.; Sethuraman, N.; Qian, Z. The endosomal escape vehicle platform enhances delivery of oligonucleotides in preclinical models of neuromuscular disorders. Mol Ther Nucleic Acids 2023, 33, 273–285.

(37) Knudsen, L. B.; Lau, J. The Discovery and Development of Liraglutide and Semaglutide. Front Endocrinol 2019, 10, 155.

(38) Park, H.; Park, J. H.; Kim, M. S.; Cho, K.; Shin, J. M. In Silico Screening and Optimization of Cell-Penetrating Peptides Using Deep Learning Methods. Biomolecules 2023, 13, 522.

(39) Shi, K.; Xiong, Y.; Wang, Y.; Deng, Y.; Wang, W.; Jing, B.; Gao, X. PractiCPP: A Deep Learning Approach Tailored for Extremely Imbalanced Datasets in Cell-Penetrating Peptide Prediction. Bioinformatics 2024, 40, btae058.

(40) Hoffmann, K.; Milech, N.; Juraja, S. M.; Cunningham, P. T.; Stone, S. R.; Francis, R. W.; Anastasas, M.; Hall, C. M.; Heinrich, T.; Bogdawa, H. M.; Winslow, S.; Scobie, M. N.; Dewhurst, R. E.; Florez, L.; Ong, F.; Kerfoot, M.; Champain, D.; Adams, A. M.; Fletcher, S.; Viola, H. M.; Hool, L. C.; Connor, T.; Longville, B. A. C.; Tan, Y. F.; Kroeger, K.; Morath, V.; Weiss, G. A.; Skerra, A.; Hopkins, R. M.; Watt, P. M. A Platform for Discovery of Functional Cell-Penetrating Peptides for Efficient Multi-Cargo Intracellular Delivery. Scientific Reports 2018, 8, 12538.

(41) Gao, S.; Simon, M. J.; Hue, C. D.; Morrison, B.; Banta, S. An Unusual Cell Penetrating Peptide Identified Using a Plasmid Display-Based Functional Selection Platform. ACS Chem Biol 2011, 6, 484–491.

(42) Carney, R. P.; Thillier, Y.; Kiss, Z.; Sahabi, A.; Heleno Campos, J. C.; Knudson, A.; Liu, R.; Olivos, D.; Saunders, M.; Tian, L.; Lam, K. S. Combinatorial Library Screening with Liposomes for Discovery of Membrane Active Peptides. ACS Comb Sci 2017, 19, 299–307.

(43) Gao, C.; Mao, S.; Ditzel, H. J.; Farnaes, L.; Wirsching, P.; Lerner, R. A.; Janda, K. D. A Cell-Penetrating Peptide from a Novel PVII–PIX Phage-Displayed Random Peptide Library. Bioorg Med Chem 2002, 10, 4057–4065.

(44) Lee, J. H.; Song, H. S.; Lee, S. G.; Park, T. H.; Kim, B. G. Screening of Cell-Penetrating Peptides Using MRNA Display. Biotechnol J 2012, 7, 387–396.

(45) Hoffmann, K.; Milech, N.; Juraja, S. M.; Cunningham, P. T.; Stone, S. R.; Francis, R. W.; Anastasas, M.; Hall, C. M.; Heinrich, T.; Bogdawa, H. M.; Winslow, S.; Scobie, M. N.; Dewhurst, R. E.; Florez, L.; Ong, F.; Kerfoot, M.; Champain, D.; Adams, A. M.; Fletcher, S.; Viola, H. M.; Hool, L. C.; Connor, T.; Longville, B. A. C.; Tan, Y. F.; Kroeger, K.; Morath, V.; Weiss, G. A.; Skerra, A.; Hopkins, R. M.; Watt, P. M. A Platform for Discovery of Functional Cell-Penetrating Peptides for Efficient Multi-Cargo Intracellular Delivery. Scientific Reports 2018, 8, 1–16.

(46) Jirka, S. M. G.; ‘t Hoen, P. A. C.; Diaz Parillas, V.; Tanganyika-de Winter, C. L.; Verheul, R. C.; Aguilera, B.; de Visser, P. C.; Aartsma-Rus, A. M. Cyclic Peptides to Improve Delivery and Exon Skipping of Antisense Oligonucleotides in a Mouse Model for Duchenne Muscular Dystrophy. Molecular Therapy 2018, 26, 132–147.

(47) Bowen, J.; Schloop, A. E.; Reeves, G. T.; Menegatti, S.; Rao, B. M. Discovery of Membrane-Permeating Cyclic Peptides via MRNA Display. Bioconjug Chem 2020, 31, 2325–2338.

(48) Schissel, C. K.; Farquhar, C. E.; Loas, A.; Malmberg, A. B.; Pentelute, B. L. In-Cell Penetration Selection-Mass Spectrometry Produces Noncanonical Peptides for Antisense Delivery. ACS Chem Biol. 2023, 18, 615–628.

(49) Koch, G.; Engstrom, A.; Taechalertpaisarn, J.; Faris, J.; Ono, S.; Naylor, M. R.; Lokey, R. S. Chromatographic Determination of Permeability-Relevant Lipophilicity Facilitates Rapid Analysis of Macrocyclic Peptide Scaffolds. J Med Chem 2024, 67, 19612.

(50) Ramaker, K.; Henkel, M.; Krause, T.; Röckendorf, N.; Frey, A. Cell Penetrating Peptides: A Comparative Transport Analysis for 474 Sequence Motifs. Drug Deliv 2018, 25, 928–937.

(51) Najjar, K.; Erazo-Oliveras, A.; Mosior, J. W.; Whitlock, M. J.; Rostane, I.; Cinclair, J. M.; Pellois, J. P. Unlocking Endosomal Entrapment with Supercharged Arginine-Rich Peptides. Bioconjug Chem 2017, 28, 2932–2941.

(52) Quartararo, A. J.; Gates, Z. P.; Somsen, B. A.; Hartrampf, N.; Ye, X.; Shimada, A.; Kajihara, Y.; Ottmann, C.; Pentelute, B. L. Ultra-Large Chemical Libraries for the Discovery of High-Affinity Peptide Binders. Nature Communications 2020, 11, 1–11.

(53) Mohapatra, S.; Hartrampf, N.; Poskus, M.; Loas, A.; Gómez-Bombarelli, R.; Pentelute, B. L. Deep Learning for Prediction and Optimization of Fast-Flow Peptide Synthesis. ACS Cent Sci 2020, 6, 2277–2286.

(54) Fadzen, C. M.; Holden, R. L.; Wolfe, J. M.; Choo, Z. N.; Schissel, C. K.; Yao, M.; Hanson, G. J.; Pentelute, B. L. Chimeras of Cell-Penetrating Peptides Demonstrate Synergistic Improvement in Antisense Efficacy. Biochemistry 2019, 58, 3980–3989.

(55) Sazani, P.; Ness, K. P. V.; Weller, D. L.; Poage, D. W.; Palyada, K.; Shrewsbury, S. B. Repeat-Dose Toxicology Evaluation in Cynomolgus Monkeys of AVI-4658, a Phosphorodiamidate Morpholino Oligomer (PMO) Drug for the Treatment of Duchenne Muscular Dystrophy. Int J Toxicol 2011, 30, 313–321.

(56) Amantana, A.; Moulton, H. M.; Cate, M. L.; Reddy, M. T.; Whitehead, T.; Hassinger, J. N.; Youngblood, D. S.; Iversen, P. L. Pharmacokinetics, Biodistribution, Stability and Toxicity of a Cell-Penetrating Peptide - Morpholino Oligomer Conjugate. Bioconjug Chem 2007, 18, 1325–1331.

(57) Sazani, P.; Gemignani, F.; Kang, S. H.; Maier, M. A.; Manoharan, M.; Persmark, M.; Bortner, D.; Kole, R. Systemically delivered antisense oligomers upregulate gene expression in mouse tissues. Nat Biotechnol. 2002, 20, 1228–33.

(58) Gan, L.; Wu, L.C.L., Wood, J.A.; Yao, M., Treleaven, C.M.; Estrella, N.L.; Wentworth, B.M.; Hanson, G.J.; Passini, M.A. A cell-penetrating peptide enhances delivery and efficacy of phosphorodiamidate morpholino oligomers in mdx mice. Mol. Ther. Nucleic Acids 2022, 30, 17–27.

(59) Oliver, R. A.; Ahern, M. E.; Castaneda, P. G.; Jinadasa, T.; Bardhan, A.; Morgan, K. Y.; Ha, K.; Adhikari, K.; Jungels, N.; Liberman, N.; Mitra, A.; Greer, C. D.; Wright, A. M.; Thompson, E. G.; Garcia, S.; Copson, E.; Allu, S.; Tan, X.; Callahan, A. J.; Cai, B. Z.; Guerlavais, V.; Kim, K. J.; Malmberg, A. B. Splicing correction by peptide-conjugated morpholinos as a novel treatment for late-onset Pompe disease. Mol Ther Nucleic Acids. 2025, 36, 102524.

(60) Takakusa, H.; Iwazaki, N.; Nishikawa, M.; Yoshida, T.; Obika, S.; Inoue, T. Drug metabolism and pharmacokinetics of antisense oligonucleotide therapeutics: Typical profiles, evaluation approaches, and points to consider compared with small molecule drugs. Nucleic Acid Ther. 2023, 33, 83–94.

(61) Bäckström, E.; Bonetti, A.; Johnsson, P.; Öhlin, S.; Dahlén, A.; Andersson, P.; Andersson, S.; Gennemark, P. Tissue pharmacokinetics of antisense oligonucleotides. Mol Ther Nucleic Acids 2024, 35, 102133.

(62) Bulfield, G.; Siller, W. G.; Wight, P. A.; & Moore, K. J. X chromosome-linked muscular dystrophy (mdx) in the mouse. Proc Natl Acad Sci U. S. A. 1984, 81, 1189–1192.

(63) Stedman, H.; Sweeney, H.; Shrager, J. et al. The mdx mouse diaphragm reproduces the degenerative changes of Duchenne muscular dystrophy. Nature 1991, 352, 536–539.

(64) Sicinski, P.; Geng, Y.; Ryder-Cook, A.; Barnard, E.; Darlinson, M.; Barnard, P. The Molecular Basis of Muscular Dystrophy in the *mdx* Mouse: a Point Mutation. Science 1989, 244, 1578–1580.

(65) Chamberlain, J.S.; Metzger, J.; Reyes, M.; Townsend, D.; Faulkner, J.A. Dystrophin-deficient mdx mice display a reduced life span and are susceptible to spontaneous rhabdomyosarcoma. FASEB J. 2007, 21, 2195–2204.

(66) Zhu, Y., Zhang, H., Sun, Y., Li, Y., Deng, L., Wen, X., Wang, H., & Zhang, C. Serum Enzyme Profiles Differentiate Five Types of Muscular Dystrophy. Disease Markers 2015, 543282.

(67) McMillan, H. J.; Gregas, M.; Darras, B. T., & Kang, P. B. Serum transaminase levels in boys with Duchenne and Becker muscular dystrophy. Pediatrics 2011, 127, e132–e136.

(68) Lin, W. Clinical Significance of Serum Creatinine, Urea Nitrogen and Uric Acid Levels in Patients with Chronic Renal Failure. IJBLS 2023, 3, 19–26.

(69) Åmand, H. L.; Fant, K.; Nordén, B.; Esbjörner, E. K. Stimulated Endocytosis in Penetratin Uptake: Effect of Arginine and Lysine. Biochem Biophys Res Commun 2008, 371, 621–625.

(70) LeCher, J. C.; Nowak, S. J.; McMurry, J. L. Breaking in and Busting out: Cell-Penetrating Peptides and the Endosomal Escape Problem. Biomol Concepts 2017, 8, 131.

(71) Su, Y.; Doherty, T.; Waring, A. J.; Ruchala, P.; Hong, M. Roles of Arginine and Lysine Residues in the Translocation of a Cell-Penetrating Peptide from 13C, 31P and 19F Solid-State NMR. Biochemistry 2009, 48, 4587–4595.

(72) Li, L.; Vorobyov, I.; Allen, T. W. The Different Interactions of Lysine and Arginine Side Chains with Lipid Membranes. J Phys Chem B. 2013, 117, 11906–11920.

(73) Robison, A. D.; Sun, S.; Poyton, M. F.; Johnson, G. A.; Pellois, J.-P.; Jungwirth, P.l; Vazdar, M.; Cremer, P. S. Polyarginine Interacts More Strongly and Cooperatively than Polylysine with Phospholipid Bilayers. J. Phys. Chem. B 2016, 120, 9287−9296.

(74) Choe, S. Insights into Translocation of Arginine-Rich Cell-Penetrating Peptides across a Model Membrane. J Phys Chem B. 2024, 128, 10894–10903.

(75) Walrant, A.; Cardon, S.; Burlina, F.; Sagan, S. Membrane Crossing and Membranotropic Activity of Cell-Penetrating Peptides: Dangerous Liaisons? Acc Chem Res 2017, 50, 2968–2975.

(76) López-Vidal, E. M.; Schissel, C. K.; Mohapatra, S.; Bellovoda, K.; Wu, C. L.; Wood, J. A.; Malmberg, A. B.; Loas, A.; Gómez-Bombarelli, R.; Pentelute, B. L. Deep Learning Enables Discovery of a Short Nuclear Targeting Peptide for Efficient Delivery of Antisense Oligomers. JACS Au 2021, 1, 2009–2020.

(77) Schissel, C. K.; Farquhar, C. E.; Malmberg, A. B.; Loas, A.; Pentelute, B. L. Cell-Penetrating d-Peptides Retain Antisense Morpholino Oligomer Delivery Activity. ACS Bio & Med Chem Au 2022, 2, 150–160.

