## Supplementary Information for "High-throughput discovery of arginine-depleted peptides enables effective antisense delivery for Duchenne muscular dystrophy"

†Author passed away in May 2024

§Affiliation at the time of study

### 1. Materials and General Methods

#### 1.1 Reagents and Solvents

H-Rink Amide-ChemMatrix 0.45 mmol/g loading resin was obtained from PCAS BioMatrix Inc. (St-Jean-sur-Richelieu, Quebec, Canada) and 180 µm monosized TentaGel 0.28 mmol/g loading resin was obtained from Rapp Polymere (Tuebingen, Germany). 1-[Bis(dimethylamino)methylene]-1*H*-1,2,3-triazolo[4,5-b]pyridinium-3-oxid-hexafluorophosphate (HATU), *N*α-Fmoc-*N*γ-Boc-L-2,4-diaminobutyric acid, Fmoc-L-Iodophenylalanine, Fmoc-3-(1-naphthyl)-D-alanine, 1-Boc-piperidine-4-Fmoc-amino-4-carboxylic acid, Fmoc-L-citrulline, Fmoc-3-(4'-pyridyl)-L-alanine, and 5-azidopentanoic acid were purchased from Chem-Impex International (Wood Dale, IL). (7-Azabenzotriazol-1-yloxy)tripyrrolidinophosphonium hexafluorophosphate (PyAOP, ≥97.0%) was purchased from P3 Biosystems. Fmoc-protected L-amino acids (Fmoc-Arg(Pbf)-OH; Fmoc-Asn(Trt)-OH; Fmoc-Asp-(O*t*-Bu)-OH; Fmoc-Gln(Trt)-OH; Fmoc-Glu(O*t*-Bu)-OH; Fmoc-Gly-OH; Fmoc-His(Trt)-OH; Fmoc-Lys(Boc)-OH; Fmoc-Phe-OH; Fmoc-Pro-OH; Fmoc-Ser(But)-OH; Fmoc-Thr(*t*-Bu)-OH; Fmoc-Trp(Boc)-OH), were purchased from the Novabiochem-line from MilliporeSigma. Dibenzocyclooctyne acid was purchased from Click Chemistry Tools (Scottsdale, AZ). Peptide synthesis-grade *N,N*-dimethylformamide (DMF, ≥99.9%, HiPerSolv CHROMANORM® for HPLC), $CH_2Cl_2$ (DCM), diethyl ether, and HPLC-grade acetonitrile were obtained from VWR International (Radnor, PA). Diisopropylethylamine (DIEA; 99.5%, biotech grade), piperidine (ACS reagent, ≥99.0%), trifluoroacetic acid (TFA, HPLC grade, ≥99.0%), and triisopropylsilane (≥98.0%) were purchased from Sigma-Aldrich. The LDH Assay kit was purchased from Promega (Madison, WI). All other reagents were purchased from Sigma-Aldrich (St. Louis, MO). Milli-Q water was used exclusively for all experiments and obtained from a Milli-Q Reference water purification system (Millipore).

#### 1.2 General peptide preparation

<u>Fast-flow Peptide Synthesis:</u>
Peptides were synthesized on a 0.1 mmol scale using a semi-automated fast-flow peptide synthesizer as previously reported.[1] 1 mmol of amino acid was combined with 2.5 mL of 0.4 M HATU and 500 µL of DIEA and mixed before being delivered to the reactor containing resin via syringe pump at 6 mL/min. The reactor was submerged in a water bath heated to 70 ºC. An HPLC pump delivered either DMF (20 mL) for washing or 20% piperidine/DMF (6.7 mL) for Fmoc deprotection, at 20 mL/min.

<u>Peptide cleavage and deprotection:</u>
Peptide chains were cleaved off the solid-phase resin and deprotected as previously reported.[1] Each peptide was subjected to simultaneous global side-chain deprotection and cleavage from resin by treatment with 5 mL of 94% trifluoroacetic acid (TFA), 2.5% thioanisole, 2.5% water, and 1% triisopropylsilane (TIPS) (v/v) at room temperature for 2 to 4 h. The cleavage cocktail was first concentrated by bubbling $N_2$ through the mixture, and cleaved peptide was precipitated and triturated with 40 mL of cold ether (chilled in dry ice). The crude product was pelleted by centrifugation for three minutes at 4,000 rpm and the ether was decanted. This wash step was repeated two more times. After the third wash, the pellet was dissolved in 50% water and 50% acetonitrile (v/v) containing 0.1% TFA, filtered through a fritted syringe to remove the resin and lyophilized.

<u>Peptide Purification:</u>
The peptides were dissolved in water and acetonitrile containing 0.1% TFA, filtered through a 0.22 µm nylon filter and purified by mass-directed semi-preparative reversed-phase HPLC. Solvent A

was water with 0.1% TFA additive and Solvent B was acetonitrile with 0.1% TFA additive. A linear gradient from 5 to 45% B that changed at a rate of 0.5% B/min was used. The peptides were purified on an Agilent Zorbax SB C18 column: 9.4 x 250 mm, 5 µm. Based on target ion mass data recorded for each fraction, only pure fractions were lyophilized. The purity of each fraction was confirmed by LC-MS and pure fractions were combined after lyophilization.

Preparation of peptide library:
Split-and-pool synthesis was carried out on 180 µm TentaGel resin (0.28 mmol/g) for a 50,000-member library. Splits were performed by suspending the resin in DCM and dividing it evenly (via pipetting) among 22 plastic fritted syringes on a vacuum manifold. Couplings were carried out as follows: solutions of Fmoc-protected amino acids (10 eq. relative to the resin loading), PyAOP (0.38 M in DMF; 0.95 eq. relative to amino acid), and DIEA (1.1 eq. for histidine; 3 eq. for all other amino acids) were each added to individual portions of resin. Couplings were allowed to proceed for 60 min. Resin portions were recombined and washed with DCM and DMF. Fmoc removal was carried out by treatment of the resin with 20% piperidine in DMF (1x flow wash; 2 x 10 min batch treatments). Resin was washed again with DMF and DCM before the next split. After synthesis, the library was separated into 3,000-member portions by mass. Each portion was conjugated to 5-azidopentanoic acid through a coupling with PyAOP as described above. Library peptides were then cleaved as described above and lyophilized to generate libraries of 3,000 peptide sequences. To create the 12,000-member libraries, four 3,000-member peptide libraries were combined into one.

Preparation of PMO-DBCO:
PMO-DBCO was generated as previously reported.[2] PMO IVS2-654 (50 mg, 8 µmol) obtained from Sarepta Therapeutics was dissolved in 150 µL DMSO. To the solution was added a solution containing 2 eq. of dibenzocyclooctyne acid (5.3 mg, 16 µmol) activated with HBTU (37.5 µL of 0.4 M HBTU in DMF, 15 µmol) and DIEA (2.8 µL, 16 µmol) in 40 µL DMF (final reaction volume = 0.23 mL). The reaction proceeded for 25 min before being quenched with 1 mL of water and 2 mL of ammonium hydroxide. The ammonium hydroxide hydrolyzed any ester formed during the course of the reaction. After 1 hour, the solution was diluted to 40 mL in water/acetonitrile and purified using reverse-phase HPLC (Agilent Zorbax SB C3 column: 21.2 x 100 mm, 5 µm) and a linear gradient from 2 to 60% B (solvent A: water; solvent B: acetonitrile) over 58 min (1% B / min). Using mass data about each fraction from the instrument, only pure fractions were pooled and lyophilized. The purity of the fraction pool was confirmed by LC-MS.

Conjugation of PMO to peptides:
PMO-DBCO (1 eq., 5 mM, water) was conjugated to azido-peptides (1 eq., 5 mM, water) or azido-peptide library (1 eq., 1 mM, water) at room temperature for 2 h, or 12 h for peptide library. Reaction progress was monitored by LC-MS and additional stock of azido-peptide was added until all PMO-DBCO was consumed. The purity of the final construct was confirmed by LC-MS to be >95%.

#### 1.3 Liquid chromatography—mass spectrometry

LC-MS analyses:
Analysis was performed on an Agilent 6550 iFunnel Q-TOF LC-MS system (abbreviated as 6550) coupled to an Agilent 1290 Infinity HPLC system. Mobile phases were: 0.1% formic acid in water (solvent A) and 0.1% formic acid in acetonitrile (solvent B). The following LC-MS method was used for characterization:

1-61% B over 6 min, Zorbax C3 column (6550)
LC: Agilent Zorbax C3 column RRHD column: 2.1 × 50 mm, 1.8 µm, column temperature: 40 °C, gradient: 0-1 min 1% B, 1-6 min, 1-61% B, 6-7 min, 91% B, 7-8 min, 1% B; flow rate: 0.5 mL/min.

MS: Positive electrospray ionization (ESI) extended dynamic range mode in mass range 300–3000 m/z. MS is on from 1 to 6 min.
All data were processed using the Agilent MassHunter Quantitatvie analysis software package. Y-axis in all chromatograms shown represents total ion current (TIC) unless noted.

Orbitrap LC-MS/MS:
Analysis was performed on an EASY-nLC 1200 (Thermo Fisher Scientific) nano-liquid chromatography handling system connected to an Orbitrap Fusion Lumos Tribrid Mass Spectrometer (Thermo Fisher Scientific). Samples were run on a PepMap RSLC 03-18 column (2 µm particle size, 15 cm × 50 µm ID; Thermo Fisher Scientific, P/N ES901). A nanoViper Trap Column (03-18, 3 µm particle size, 100 Å pore size, 20 mm × 75 µm ID; Thermo Fisher Scientific, P/N 164946) was used for desalting. The standard nano-LC method was run at 40 °C and a flow rate of 300 nL/min with the following gradient: 1% solvent B in solvent A ramping linearly to 41% B in A over 55 min, where solvent A = water (0.1% FA), and solvent B = 80% acetonitrile, 20% water (0.1% FA). Positive ion spray voltage was set to 2200 V. Orbitrap detection was used for primary MS, with the following parameters: resolution = 120,000; quadrupole isolation; scan range = 150–1200 m/z; RF lens = 30%; AGC target = 250%; maximum injection time = 100 ms; 1 microscan. Acquisition of secondary MS spectra was done in a data-dependent manner: dynamic exclusion was employed such that a precursor was excluded for 30 s if it was detected four or more times within 30 s (mass tolerance: 10.00 ppm); monoisotopic precursor selection used to select for peptides; intensity threshold was set to $2 \times 10^4$; charge states 2–10 were selected; and precursor selection range was set to 200–1400 m/z. The top 15 most intense precursors that met the preceding criteria were subjected to subsequent fragmentation. Two fragmentation modes— higher-energy collisional dissociation (HCD), and electron-transfer/higher-energy collisional dissociation (EThcD)—were used for acquisition of secondary MS spectra. Detection was performed in the Orbitrap (resolution = 30,000; quadrupole isolation; isolation window = 1.3 m/z; AGC target = $2 \times 10^4$; maximum injection time = 100 ms; 1 microscan). For HCD, a stepped collision energy of 3, 5, or 7% was used. For EThcD, a supplemental activation collision energy of 25% was used.

De novo peptide sequencing and filtering:
De novo peptide sequencing of the acquired data was performed in PEAKS 8 (BioInformatics Solutions Inc.). Using PEAKS, spectra were prefiltered to remove noise, and sequenced. All non-canonical amino acids were sequenced as post-translational modifications based on the canonical amino acid most closely matching their molecular mass. Twenty candidate sequence assignments were created for each secondary scan. Post-de novo data analysis was performed as previously described.[3]

Charge assignment:
To determine the average charge of each peptide pool, all positively charged residues (Lys, Arg, His, Pip, Dab, and Pal) were assigned +1, and negatively charged residues (Glu, Asp) were assigned -1. A net charge was assigned to each peptide sequence, which were then averaged to generate net charge of the peptide pools.

#### 1.4. Library fractionation and pooling

Library separation:
The 3,000-member peptide library was dissolved in loading buffer (10 mM ammonium acetate, pH 5, 20% acetonitrile). The peptides were then resolved on a 250 mm x 4 mm Propac SCX-10 column (Thermo Fisher). Solvent A was 10 mM ammonium acetate with 10% acetonitrile, pH 5, and Solvent B was 1 M ammonium acetate with 10% acetonitrile, pH 5. A linear gradient from 1 to 75% B over 75 min was used, with a 10 min hold at 1% B to allow compound to load onto the column. Fractions eluting off the column were collected every minute, with a total volume of 1 mL.

Library desalting:

All fractions were lyophilized overnight, and then re-dissolved in 1 mL of water. Fractions were then frozen and re-lyophilized to remove remaining ammonium acetate buffer. This process was repeated for at least 3 lyophilization cycles to fully remove the volatile buffer. Once the ammonium acetate was removed, appearance of peptide samples changed from crystalline solid to a fluffy powder.

Library pooling:

After desalting, the peptide concentration on each fraction was measured spectroscopically at 280 nM, based on the absorbance of the single tryptophan residue in each peptide library member. Neighboring peptide fractions were combined based on peptide concentration to generate 10 pools with approximately equal peptide concentration. For the 3,000-member libraries, this resulted in pools containing about 300 peptide sequences. For the 12,000-member libraries, this resulted in pools containing about 1,200 different peptide sequences.

#### 1.5. In-vitro evaluation of PMO-peptides

EGFP Assay:

The delivery of PMO to the cell nucleus was measured in a functional assay using HeLa 654 cells as previously reported.[2] HeLa 654 cells obtained from the University of North Carolina Tissue Culture Core facility were maintained in DMEM supplemented with 10% (v/v) fetal bovine serum (FBS) and 1% (v/v) penicillin-streptomycin at 37 °C and 5% $CO_2$. 18 h prior to treatment, the cells were plated at a density of 5,000 cells per well in a 96-well plate in DMEM supplemented with 10% FBS and 1% penicillin-streptomycin.

For individual peptide testing, PMO-peptides were dissolved in PBS without $Ca^{2+}$ or $Mg^{2+}$ at a concentration of 1 mM (determined by UV absorbance of the PMO) before being diluted in DMEM. Cells were incubated at the designated concentrations in triplicate for 22 h at 37 °C and 5% $CO_2$. Next, the treatment media was removed, and the cells were washed once before being incubated with 0.25 % Trypsin-EDTA for 15 min at 37 °C and 5% $CO_2$. Lifted cells were transferred to a V-bottom 96-well plate and washed once with PBS, before being resuspended in PBS containing 2% FBS and 2 μg/mL propidium iodide (PI). Flow cytometry analysis was carried out on a BD LSRII flow cytometer. Gates were applied to the data to ensure that cells that were positive for propidium iodide or had forward/side scatter readings that were sufficiently different from the main cell population were excluded. Each sample was capped at 5,000 gated events.

Analysis was conducted using GraphPad Prism 7 and FlowJo. For each sample, the mean fluorescence intensity (MFI) and the number of gated cells was measured. To report activity, triplicate MFI values were averaged and normalized to the PMO alone condition.

LDH assay:

Cytotoxicity assays were performed in HeLa 654 cells or TH-1 renal cell as previously reported.[2] TH-1 cells were maintained in DMEM supplemented with 10% (v/v) fetal bovine serum (FBS) and 1% (v/v) penicillin-streptomycin at 37 °C and 5% $CO_2$ Cells were plated at 5,000 cells/well in a 96-well plate 18 h prior to treatment and then treated with PMO-peptide compounds as described above. HeLa cell LDH assays were performed concurrently on the cells treated for flow cytometry, so all plating and treatment protocols match those described above.

17 hours after cell treatment, the Promega CytoTox lysis solution was added to 3 wells of each plate as a fully lysed control (100% LDH-release). 18 hours after treatment, cell supernatant was transferred to a new 96-well plate for analysis of LDH release. To each well of the 96-well plate containing supernatant was added CytoTox 96 Reagent (Promega). The plate was shielded from light and incubated at room

temperature for 30 min. Equal volume of Stop Solution was added to each well, mixed, and the absorbance of each well was measured at 490 nm. The measurement of vehicle-treated cells was subtracted from each measurement, and % LDH release was calculated as % cytotoxicity = 100 × Experimental LDH Release (OD490) / Maximum LDH Release (OD490).

Serum stability assay:

Normal mouse serum was purchased from ThermoFisher Scientific. Serum stability was assessed through incubating the PMO–peptides in 25% mouse serum or PBS at 50 μM at 0, 3, 6, 12, and 24 h in duplicate. At each timepoint peptide was dissociated from serum proteins with 4 M guanidine hydrochloride. To remove guanidine hydrochloride and serum proteins, 1/3 of the sample from each time point (10 μL) was purified via solid-phase extraction with Millipore ZTC18S008 10 μL ZipTips. Samples containing the PMO-peptide were eluted with 10 μL of 70% acetonitrile in water containing 0.1% TFA into an LC-MS vial containing 20 μL of water with 0.1% formic acid additive and analyzed via LC-MS. The amount of intact compound was determined through quantifying the ion count in the extracted ion chromatogram (EIC) using the MassHunter software. Each timepoint was performed in duplicate. Data is given for each timepoint relative to the same sample incubated in PBS at that timepoint.

#### 1.6. In-vivo evaluation of PMO-peptides

**Animals**

All animal studies were conducted with the approval of the Institutional Animal Care and Use Committee at Sarepta and performed according to their guidelines.

**EGFP Fluorescence Quantification (EGFP-654 mouse model)**

60-90 mg of tissue was homogenized in 20 μL ELOHA lysis buffer (10 mM Tris-HCl pH 7.5, 0.5% IGEPAL,100 mM NaCl, 5 mM EDTA) per mg tissue. Tissues were homogenized in individual 2 mL tubes containing metal beads using a SPEX SAMPLEPREP Geno/Grinder 2010 (5 cycles, 1,600 RPM). Homogenized lysate was centrifuged at max speed for 10 min at 4 °C. 100 μL of the supernatant/lysate was transferred to a Corning black bottom, 96 well plate and EGFP fluorescence was read using a SpectraMax plate reader (Ex485nm, Em511nm).

**RNA Isolation and Exon 23 Skipping ddPCR (*mdx* mouse model)**

Frozen tissue samples were cut on dry ice and homogenized in RLT buffer via FastPrep-24 bead-based homogenization system. After homogenization, samples were centrifuged, and supernatant was transferred to RNA isolation tubes. RNA isolation was completed via Zymo Quick-RNA Miniprep Kit (Zymo Cat# R1054). Once RNA concentration was measured via nanodrop, RNA samples were normalized and loaded into a ddPCR plate. Exon skipping was measured via droplet digital polymerase chain reaction (ddPCR) using One-Step RT-ddPCR Advanced Kit for Probes (Cat# 1864022) and the following primers and probes: Forward (5'-CAATGCGCTATCAGGAGACA-3'), Reverse (5'-ACATCAACTTCAGCCATCCA-3'), Unskipped (5'-/56-FAM/AGAAATATC/ZEN/TGTCAGAATTTGAAGAGATTGAGG/3IABkFQ/-3') and Skipped (5'-/5HEX/TGGAGGAGA/ZEN/GACTCGGGAAATTACAGAATCACATAAAAAC/3IABkFQ/-3') Droplets were generated using the Bio-Rad Automated Droplet Generator (Cat# 1864101). Droplets were amplified by thermocycling at 43C for 60 minutes, 95C for 10 minutes, 40 cycles of (95C for 30 seconds, 58C for 60 seconds, 72C for 2 minutes), followed by 10 minutes at 98C. Samples were analyzed using a Bio-Rad QX200 Droplet Reader. Percent exon skipping was calculated as a ratio of skipped copies to total transcripts (skipped+unskipped copies).

**Enzyme-Linked Oligonucleotide Hybridization Assay (ELOHA)**

The concentration of PMO in tissue was determined by Enzyme-Linked Oligonucleotide Hybridization Assay (ELOHA). Oligonucleotide capture probes were designed to hybridize to the 5' end of the PMO sequence. Coating solution (500 nM capture probe, 2.5% sodium bicarbonate solution) was incubated for 1 hour at 37°C to ensure the capture probes were covalently attached to Pierce Maleic Anhydride Activated Plates (ThermoFisher 15110). Plates were then washed and blocked overnight at 4°C with 10% milk in PBS-Tween20 (PBST). Tissue samples were lysed in ELOHA lysis buffer (10 mM Tris-HCl pH 7.5, 0.5% IGEPAL, 100 mM NaCl, 5 mM EDTA) using 20 µL of buffer per mg of tissue and then digested with Proteinase K ( 1mg/mL, Qiagen 19134) for 30 minutes at 60°C. 30 µL of the digested lysate was then incubated with 120 µL Detection Probe Solution (333 nM biotinylated oligonucleotide detection probe targeting the 3' sequence of the PMO, 500 mM guanidine thiocyanate, 0.04% lauryl sarcosine, 3 mM sodium citrate, and 1.25 mM DTT in PBST) and incubated overnight at 4°C on the blocked, capture probe coated maleic anhydride plate. The plates were washed using PBST and incubated for one hour with streptavidin-AP conjugate (Sigma-Aldrich 11093266910). Another wash-step was performed, and AttoPhos® AP Fluorescent Substrate System (Promega S1000) was added for 10 minutes at room temperature followed by 10 µL of EDTA (0.5M Solution/pH 8.0) (Fisher Scientific 8BP2482100) to stop the reaction. Using a Spectramax i3 reader (Ex440, Em555), fluorescence was measured. Each sample was interpolated to a standard curve prepared for each compound with a Sigmoidal, four-parameter logistic (4PL) model to determine ng PMO per µL of tissue lysate. The final tissue concentration value was reported as ng PMO per gram of tissue (ng/g).

**Serum Biomarker Evaluation**

Terminal serum from *mdx* mice was collected by cardiac puncture at 7 days post injection and biomarkers of skeletal muscle damage were analyzed by IDEXX BioAnalytics (North Grafton, MA) using a custom panel measuring the following markers: Creatine Kinase (CK), Alanine Aminotransferase (ALT), Aspartate Aminotransferase (AST), blood urea nitrogen (BUN)/Creatinine Ratio, and Lactate Dehydrogenase (LDH).

#### 2. Supplementary Figures

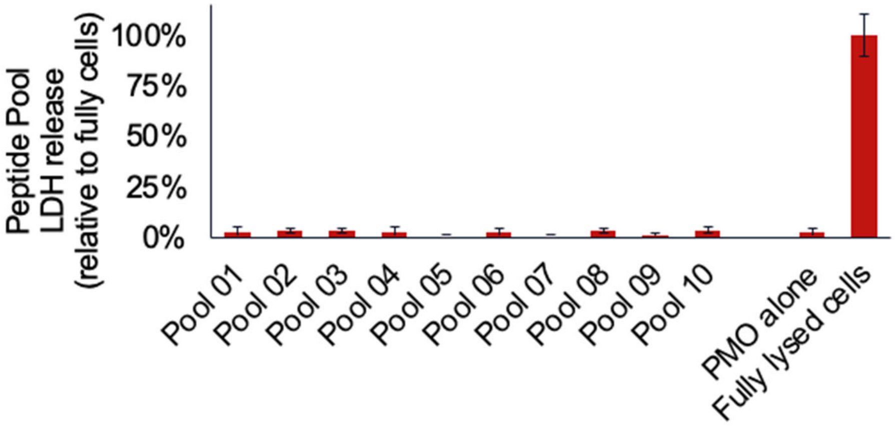

**Supplementary Figure 1. Cell membrane toxicity data from the 3,000-member peptide library divided into 10 pools.** Cell supernatant from Fig. 3a was tested for LDH release into the cell culture media at 20µM. Results are given as percent LDH release above untreated cells relative to fully lysed cells. No library pools demonstrated significant toxicity relative to untreated cells.

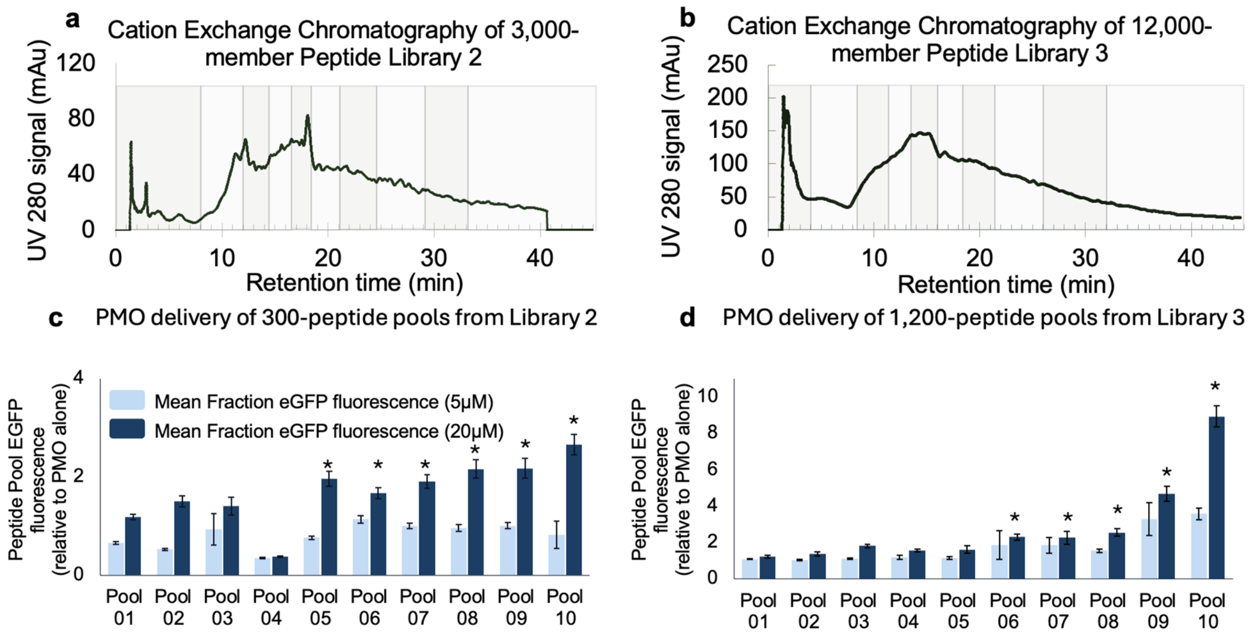

**Supplementary Figure 2. UV chromatogram traces and PMO delivery of two 3,000-member peptide libraries resolved into ten 300-peptide pools.** **(a)** The 3,000-member library was resolved through cation exchange chromatography in ammonium acetate 10% acetonitrile (pH 5) buffer with a 20-500 mM gradient over 50 min. All material was collected and pooled based on UV signal at 280 nm. Material was combined into 10 pools of approximately equal peptide content. **(b)** A 12,000-member library containing unique peptide sequences was resolved through cation exchange chromatography in ammonium acetate 10% acetonitrile (pH 5) buffer with a 20-500 mM gradient over 50 min. All material was collected and pooled based on UV signal at 280 nm. **c)** EGFP fluorescence of the 10 pools from the 3,000-member library resolution. HeLa 654 cells were treated with 5 or 20 μM PMO-peptide for 22 h prior to flow cytometry. Results are given as the mean EGFP fluorescence of cells treated with PMO-peptide relative to the fluorescence of cells treated with vehicle only. Bars represent mean ± SD, N = 3. PMO-peptide fractions 6 through 10 improved PMO delivery significantly over the PMO-654 alone at 20 μM ($p<0.01$). (**d)** EGFP fluorescence of the 10 pools from the second 12,000-member library resolution. HeLa 654 cells were treated with 5 or 20 μM PMO-peptide for 22 h prior to flow cytometry. Results are given as the mean EGFP fluorescence of cells treated with PMO-peptide relative to the fluorescence of cells treated with vehicle only. Bars represent mean ± SD, N = 3. PMO-peptide fractions 6 through 10 improved PMO delivery significantly over the PMO-654 alone at 20 μM ($p<0.01$).

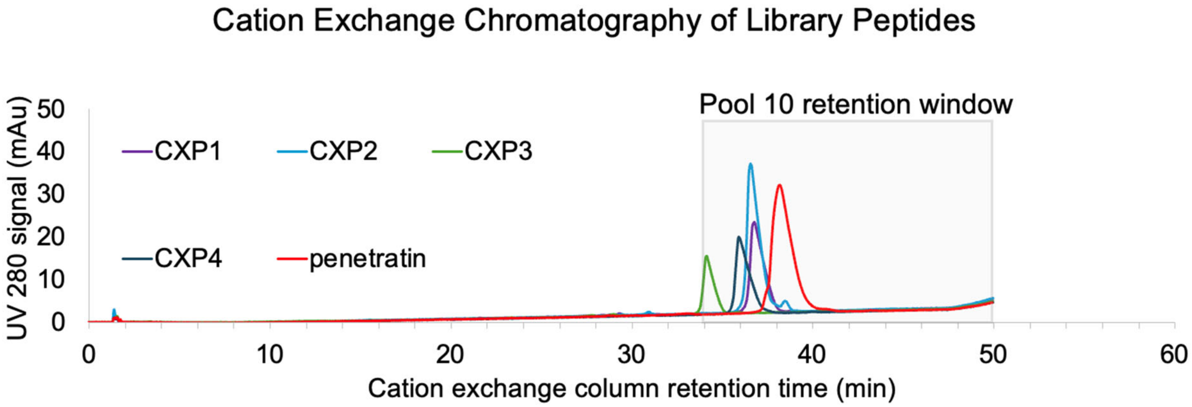

**Supplementary Figure 3. UV chromatogram traces of CXP peptides and penetratin. (a)** The 3,000-member library was resolved through cation exchange chromatography in ammonium acetate 10% acetonitrile (pH 5) buffer with a 20-500mM gradient over 50 minutes. All material was collected and pooled based on UV signal at 280nm. **(b)** a second 3,000-member library containing unique peptide sequences was resolved through cation exchange chromatography in ammonium acetate 10% acetonitrile (pH 5) buffer with a 20-500mM gradient over 50 minutes. All material was collected and pooled based on UV signal at 280nm.

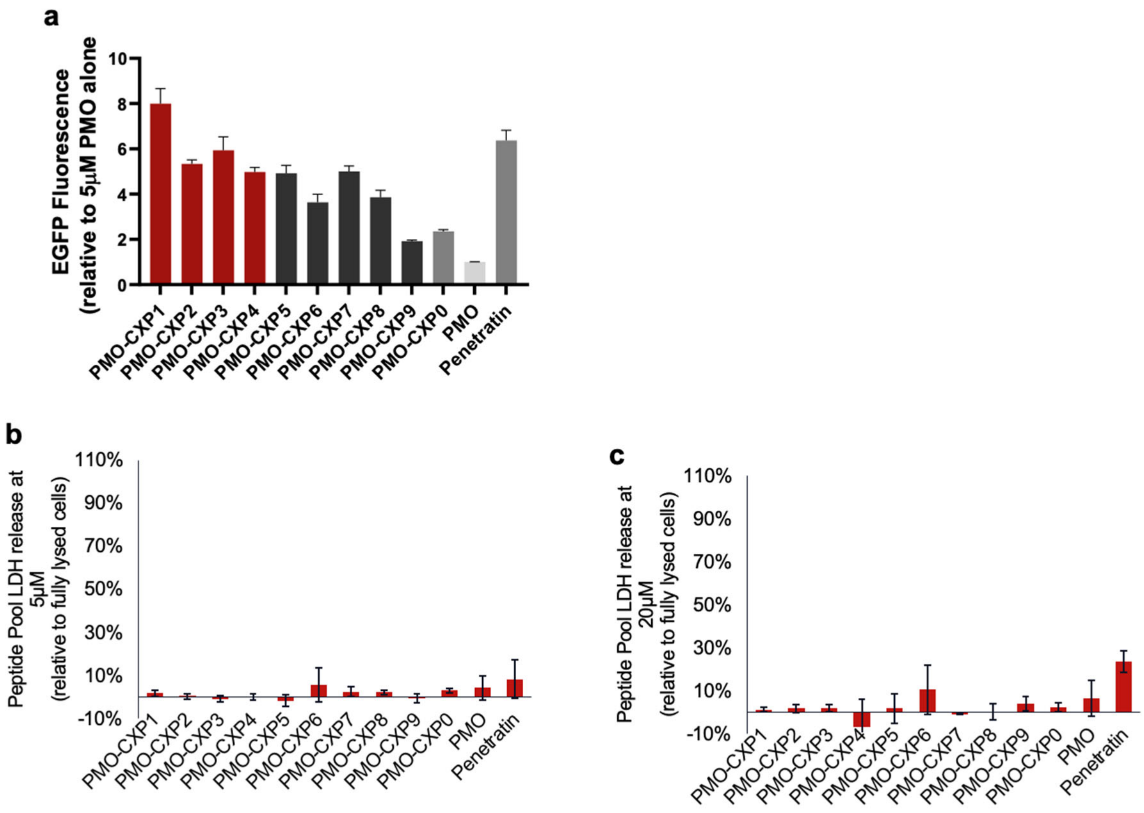

**Supplementary Figure 4: The PMO delivery and membrane toxicity of the ten peptide sequences chosen from pool 10.** **(a)** The EGFP fluorescence of each peptide-PMO relative to the fluorescence of 5 µM of PMO 654 alone determined via flow cytometry. HeLa 654 cells were treated with 5 or 20 µM PMO-peptide for 22 h prior to flow cytometry. Results are given as the mean EGFP fluorescence of cells treated with PMO-peptide relative to the fluorescence of cells treated with vehicle only. Bars represent mean ± SD, N = 3. **(b)** Cell supernatant from (a) was tested for LDH release into the treatment media. Results are given as percent LDH release above vehicle relative to fully lysed cells. (c) Cell supernatant from main text Figure 3(b) was tested for LDH release into the treatment media. Results are given as percent LDH release above vehicle relative to fully lysed cells. Only PMO-Penetratin at 20 µM showed significant toxicity relative to untreated cells.

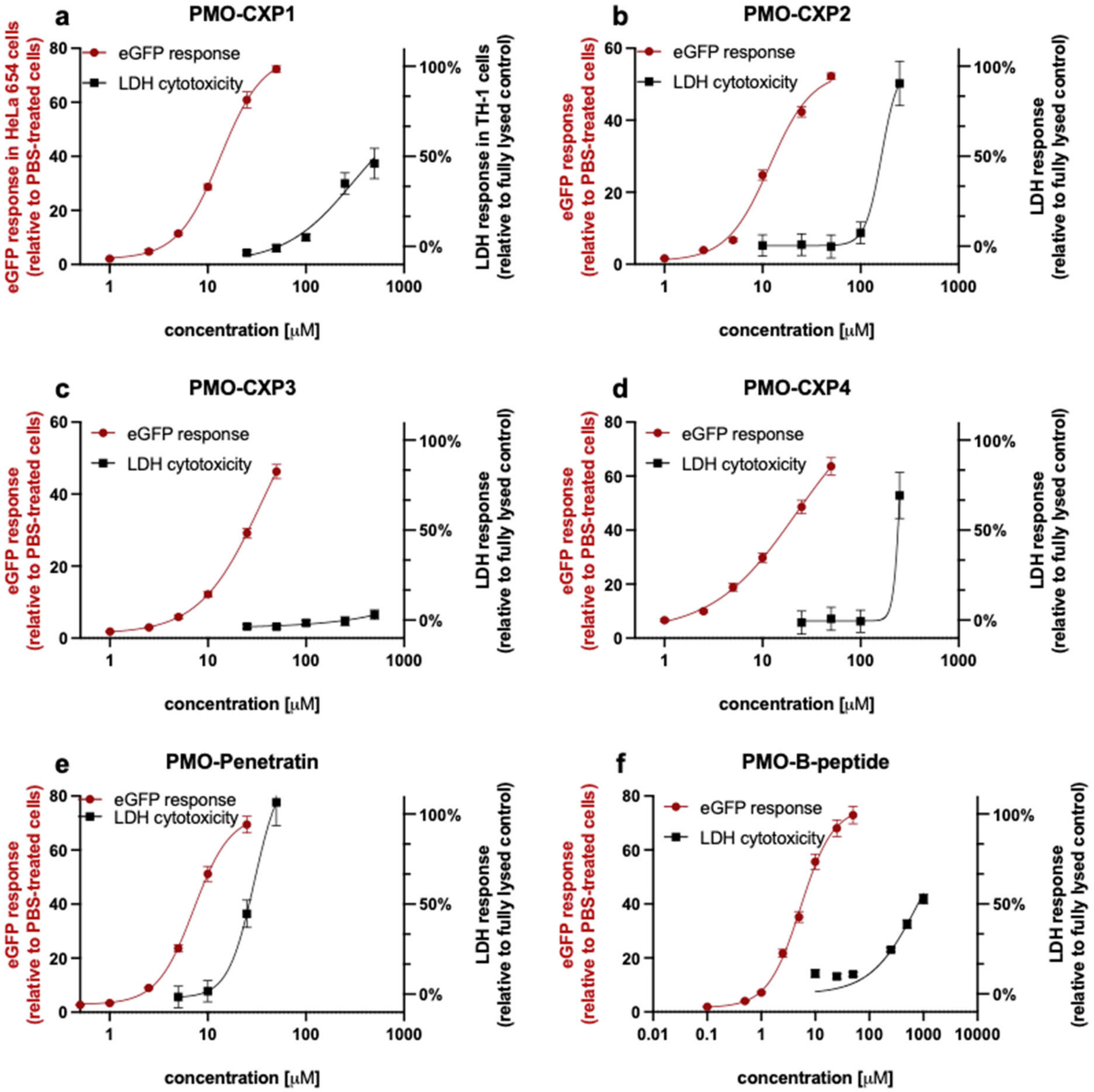

**Supplementary Figure 5: Efficacy (nuclear PMO delivery) in HeLa 654 cells and membrane toxicity (LDH release) in TH-1 renal cells of the CXP compounds and penetratin.**

**Red:** HeLa 654 cells were treated with 1, 2.5, 5, 10, 25, or 50 μM PMO-CPP for 22 h prior to flow cytometry. Results are given as the mean EGFP fluorescence of cells treated with PMO-peptide relative to the fluorescence of cells treated with vehicle only. Bars represent mean ± SD, N = 3. **Black:** TH-1 renal epithelial tubules were treated with 5, 10, 25, 50, 100, 250, or 500 μM PMO-CPP for 22 h was tested for LDH release as a measure of membrane permeability. Results are given as percent LDH release above vehicle relative to fully lysed cells. No compound showed significant LDH release above untreated cells. (a) PMO-CXP1, (b) PMO-CXP2, (c) PMO-CXP3, (d) PMO-CXP4, (e) PMO-Penetratin.

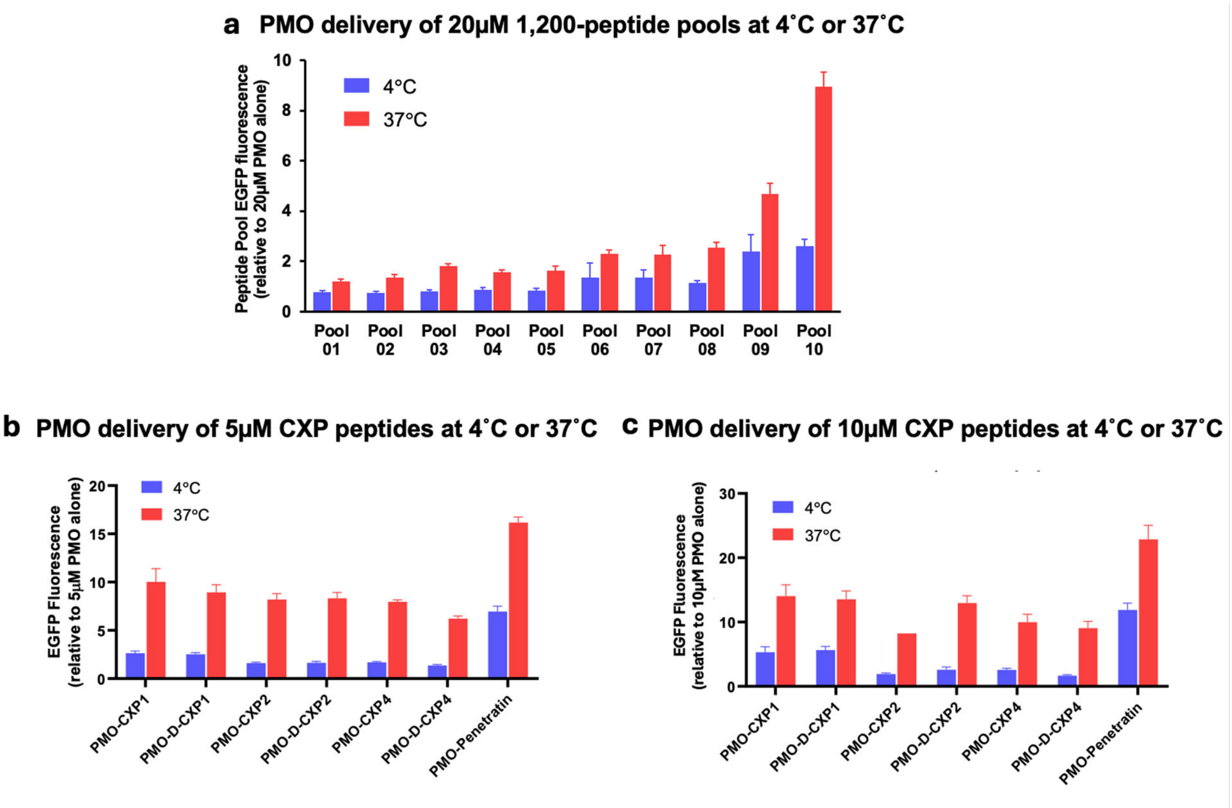

**Supplementary Figure 6. PMO delivery of 12,000-member peptide libraries resolved into 10 1,200-peptide pools at 37 ˚C and 4 ˚C.** HeLa 654 cells were treated with 5, 10, or 20 µM PMO-CPP for 2 hours at 4 ˚C or 37 ˚C. Cells were then washed with PBS and 0.1 mg/mL heparin, before returning to 37 ˚C for 20 h prior to flow cytometry. Results are given as the mean EGFP fluorescence of cells treated with PMO-peptide relative to the fluorescence of cells treated with 20µM PMO only. Bars represent mean ± SD, N = 3. **(a)** PMO-delivery of the 12,000-member library was divided into ten 1,200-peptide pools as shown in Figure 3. **(b)** PMO delivery of the D- and L-CXP peptides at 5 µM. **(c)** PMO delivery of the D- and L-CXP peptides at 5 µM.

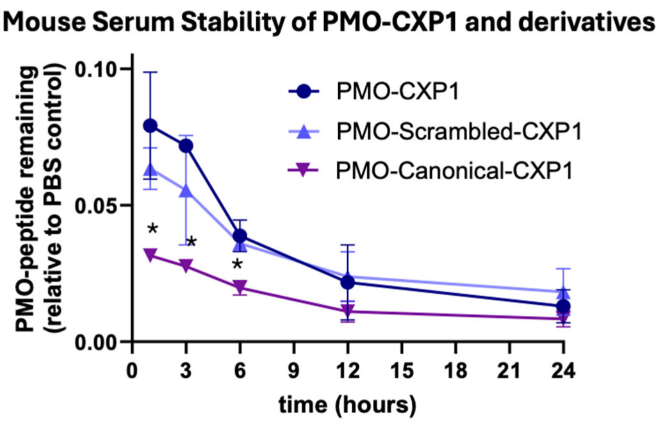

**Supplementary Figure 7: The mouse serum stability of PMO-CXP1 and variants.** PMO-peptides were incubated in 25% mouse serum in PBS or 100% PBS for 6 hours. Remaining peptide was measured by LCMS and quantified relative to the PBS-incubated samples. Bars represent mean ± SD, N = 2

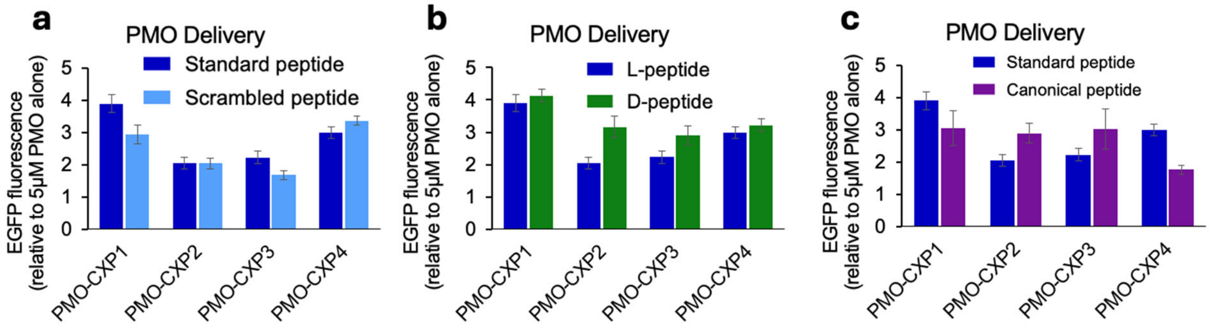

**Supplementary Figure 8: The PMO delivery and stability of scrambled, D-, or canonical variants of the CXP peptides.** HeLa 654 cells were treated with 5 μM PMO-CPP for 22 h prior to flow cytometry. Results are given as the mean EGFP fluorescence of cells treated with PMO-peptide relative to the fluorescence of cells treated with 5 μM PMO only. Bars represent mean ± SD, N = 3. **(a)** The PMO delivery of PMO-CXP peptides and their scrambled variants. (**b)** The PMO delivery of PMO-CXP peptides and their mirror-image D-variants. **(c)** The PMO delivery of PMO-CXP peptides and their canonical variants.

#### 3. LC-MS Characterization of PMO-peptides

Analysis was performed on an Agilent 6550 iFunnel Q-TOF LC-MS system (abbreviated as 6550) coupled to an Agilent 1290 Infinity HPLC system. Mobile phases were: 0.1% formic acid in water (solvent A) and 0.1% formic acid in acetonitrile (solvent B). The following LC-MS method was used for characterization:

1-61% B over 6 min, Zorbax C3 column (6550)

LC: Agilent Zorbax C3 column RRHD column: 2.1 × 50 mm, 1.8 µm, column temperature: 40 °C, gradient: 0-1 min 1% B, 1-6 min, 1-61% B, 6-7 min, 91% B, 7-8 min, 1% B; flow rate: 0.5 mL/min.

MS: Positive electrospray ionization (ESI) extended dynamic range mode in mass range 300–3000 m/z. MS is on from 1 to 6 min.

All data were processed using the Agilent MassHunter Quantitative analysis software package. Y-axis in all chromatograms shown represents total ion current (TIC) unless noted.

The figures below show the LCMS trace of the final PMO-peptide constructs, with the mass spectrum of the key peak spectral window shown below.

**PMO-DBCO**
Mass Expected: 6500.0
Mass Observed: 6499.9

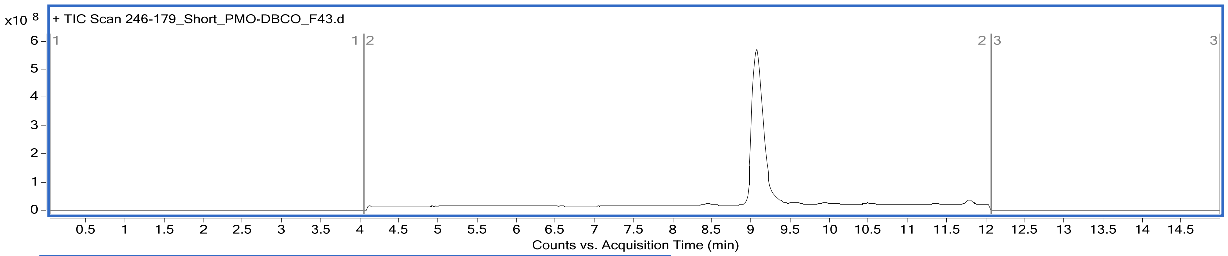

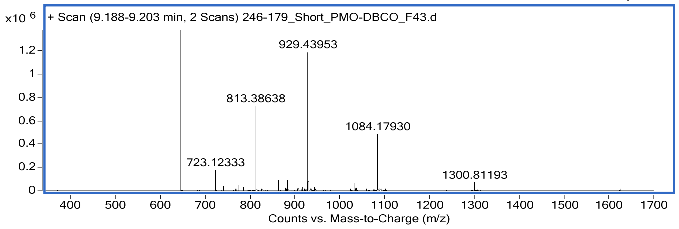

Spectral window of 9.19-9.20 min, observed mass calculated from computational deconvolution and confirmed by the $[M+7H]^{7+}$ ion, 929.4395

**PMO-CXP1**
Mass Expected: 8512.10
Mass Observed: 8512.06
Peptide sequence: 5azido-Gly-Lys-Gln-Lys-Thr-Ser-<u>Iph</u>-Gly-Arg-Gly-<u>Pip</u>-Lys-Trp-Lys-Lys

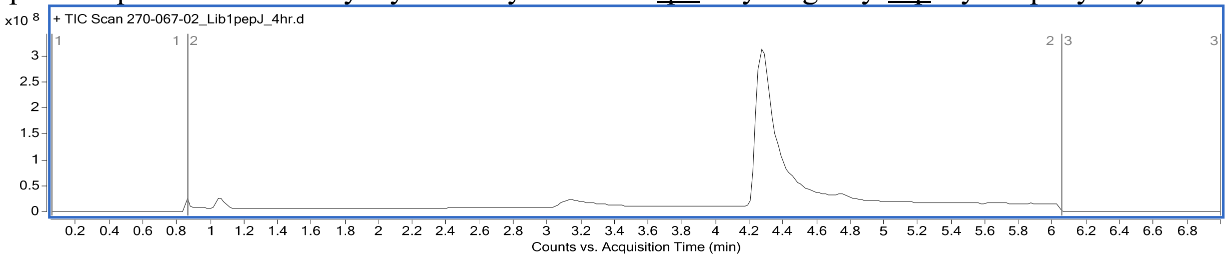

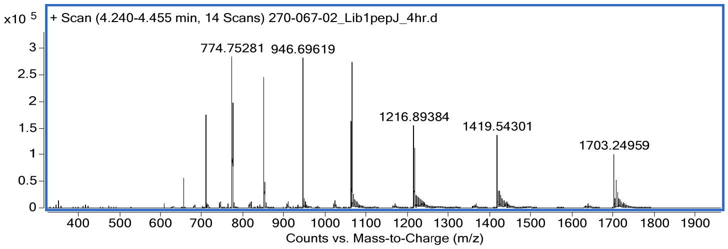

Spectral window of 4.24-4.46 min, observed mass calculated from computational deconvolution and confirmed by the $[M+9H]^{9+}$ ion, 946.6962

**PMO-CXP2**

Mass Expected: 8506.32

Mass Observed: 8506.35

Peptide sequence: 5azido-Gly-Asn-Phe-Lys-Gln-Nap-His-Ala-Gly-Dab-Arg-Lys-Trp-Lys-Lys

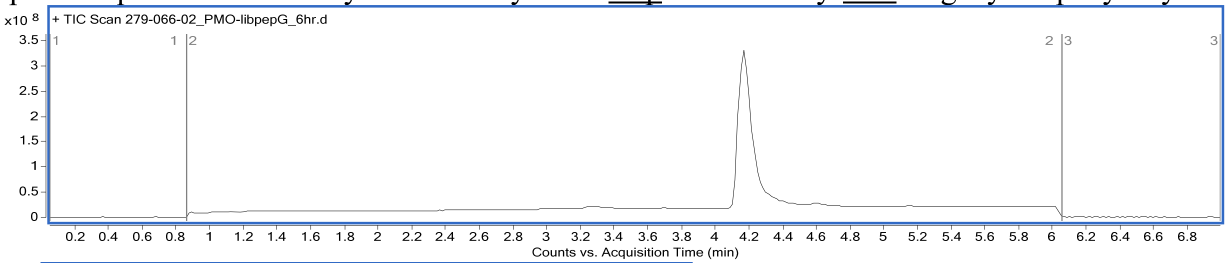

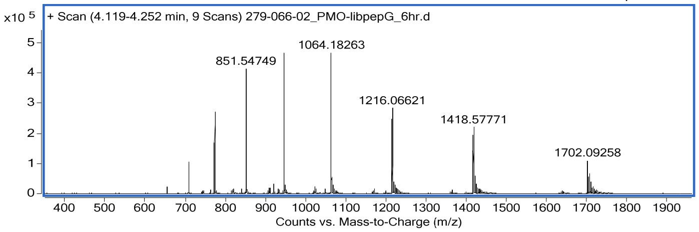

Spectral window of 4.25 min, observed mass calculated from computational deconvolution and confirmed by the $[M+8H]^{8+}$ ion, 1064.1826

**PMO-CXP3**

Mass Expected: 8311.09

Mass Observed: 8311.25

Peptide sequence: 5azido-Gly-Dab-Val-Ala-Arg-Asn-Asn-Dab-Thr-Lys-Gly-Lys-Trp-Lys-Lys

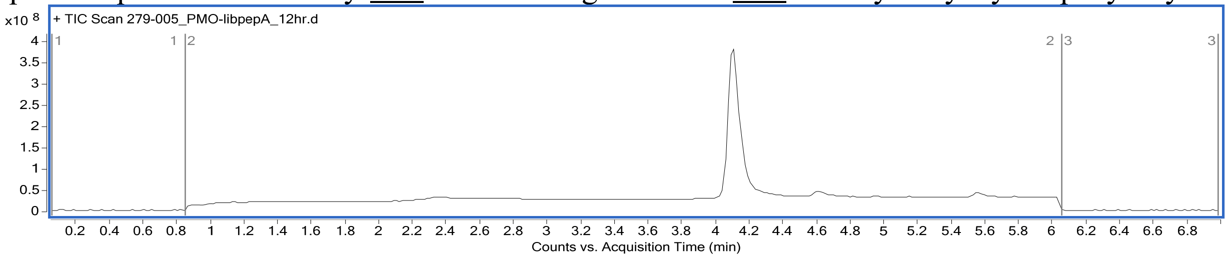

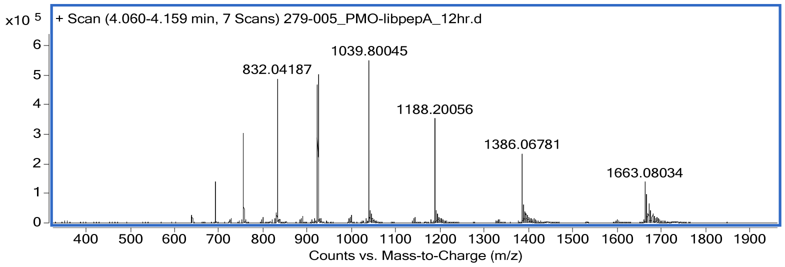

Spectral window of 4.06-4.16 min, observed mass calculated from computational deconvolution and confirmed by the $[M+8H]^{8+}$ ion, 1039.8005

**PMO-CXP4**
Mass Expected: 8385.29
Mass Observed: 8385.33
Peptide sequence: 5azido-Gly-Cit-Met-Phe-Gly-Pip-Dab-Dab-Lys-Ala-Pro-Lys-Trp-Lys-Lys

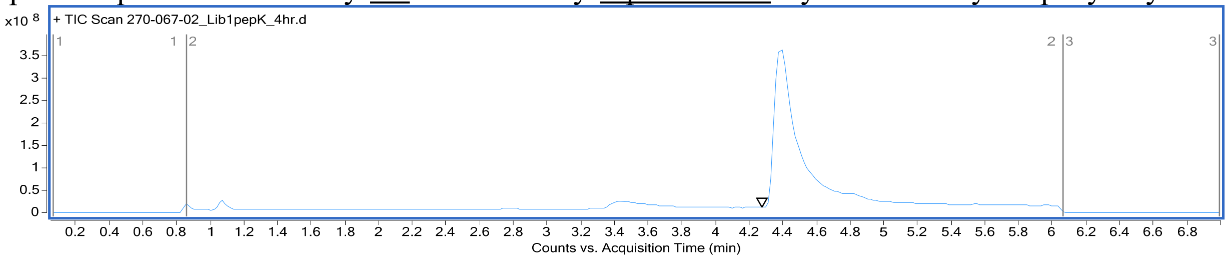

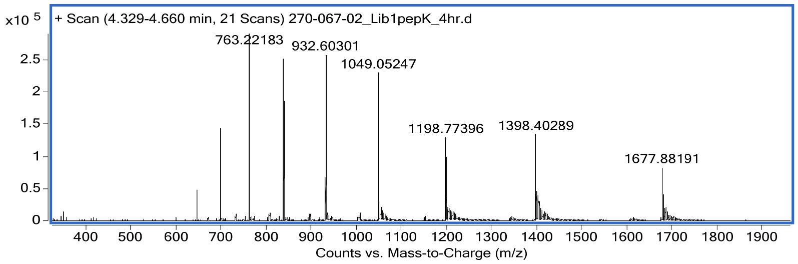

Spectral window of 4.33-4.66 min, observed mass calculated from computational deconvolution and confirmed by the $[M+8H]^{8+}$ ion, 1049.0525

**PMO-CXP5**
Mass Expected: 8333.13
Mass Observed: 8333.10
Peptide sequence: 5azido-Gly-Dab-Pro-Arg-Dab-Leu-His-Ser-Dab-Ala-Thr-Lys-Trp-Lys-Lys

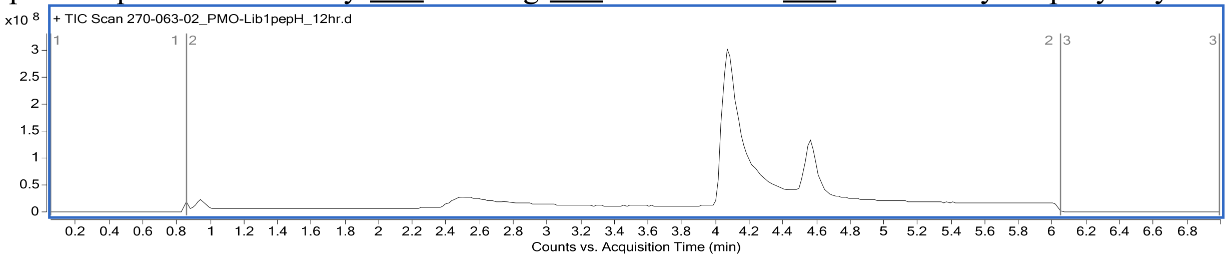

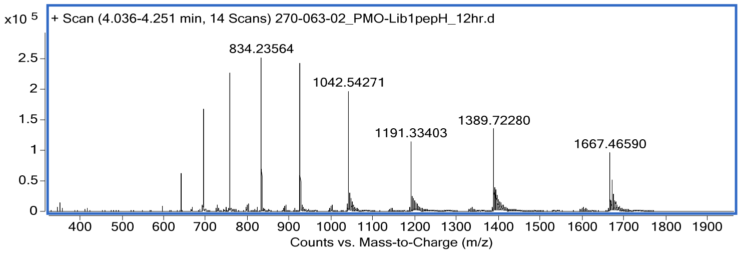

Spectral window of 4.04-4.25 min, observed mass calculated from computational deconvolution and confirmed by the $[M+8H]^{8+}$ ion, 1042.5427

**PMO-CXP6**
Mass Expected: 8535.16
Mass Observed: 8535.16
Peptide sequence: 5azido-Gly-Pro-Pip-Gly-Dab-Lys-Dab-Ser-Val-Iph-Nap-Lys-Trp-Lys-Lys

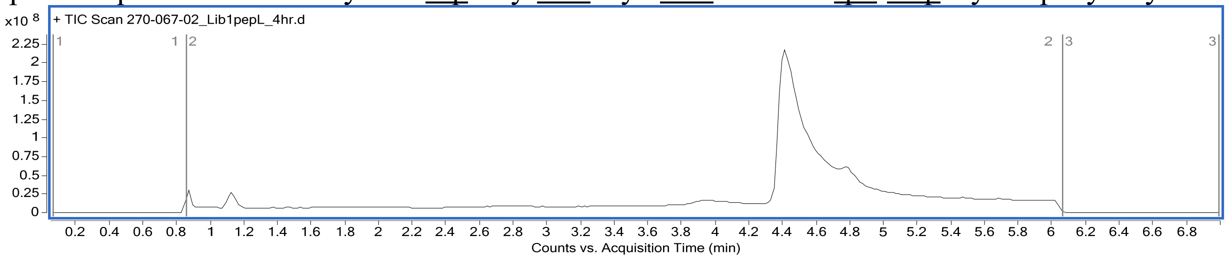

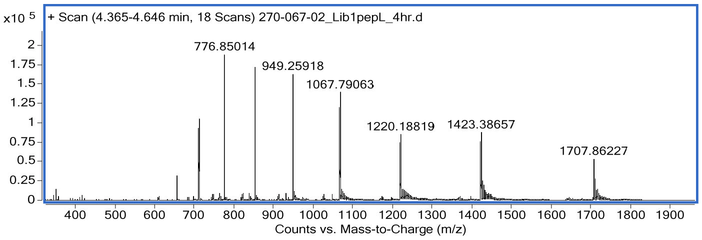

Spectral window of 4.37-4.65 min, observed mass calculated computational deconvolution and confirmed by from the $[M+8H]^{8+}$ ion, 1067.7906

**PMO-CXP7**
Mass Expected: 8663.27
Mass Observed: 8663.24
Peptide sequence: 5azido-Gly-Val-Gln-His-Iph-Arg-Dab-Lys-Glu-Lys-Asn-Lys-Trp-Lys-Lys

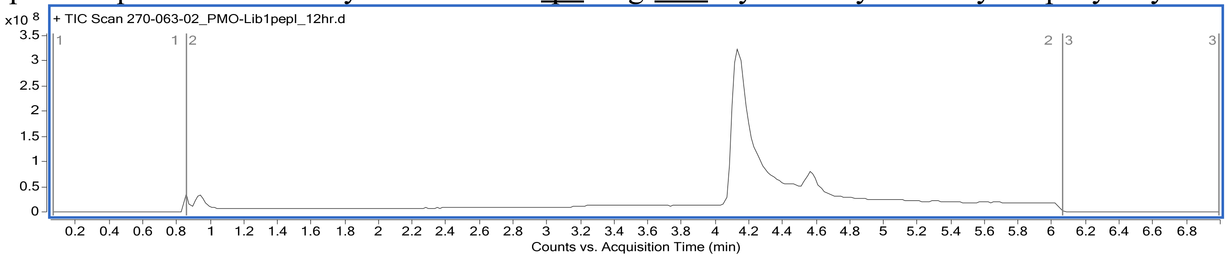

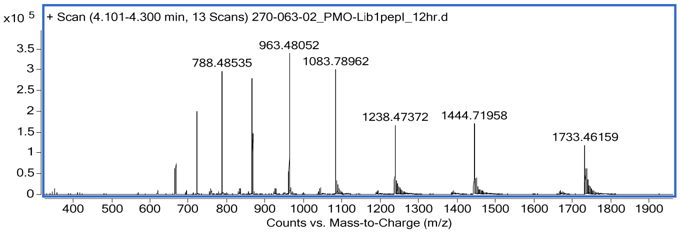

Spectral window of 4.10-4.30 min, observed mass calculated from computational deconvolution and confirmed by the $[M+9H]^{9+}$ ion, 963.4805

**PMO-CXP8**
Mass Expected: 8425.17
Mass Observed: 8425.31
Peptide sequence: 5azido-Gly-Asn-Phe-Asn-Asn-Arg-Ser-Dab-Gly-His-Lys-Lys-Trp-Lys-Lys

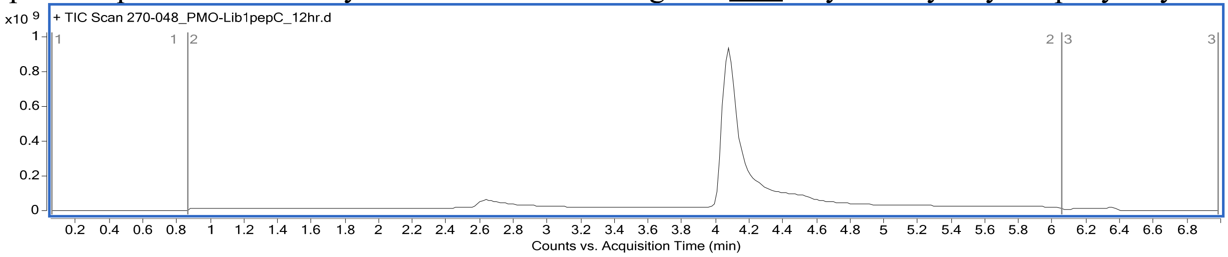

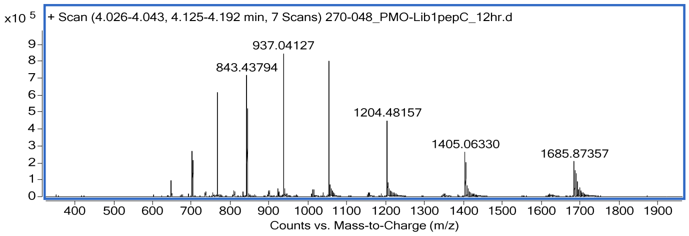

Spectral window of 4.03-4.13 min, observed mass calculated from computational deconvolution and confirmed by the $[M+9H]^{9+}$ ion, 937.04127

**PMO-CXP9**
Mass Expected: 8500.23
Mass Observed: 8500.38
Peptide sequence: 5azido-Gly-Ala-Arg-Ser-Glu-Gln-His-His-Nap-Ser-Dab-Lys-Trp-Lys-Lys

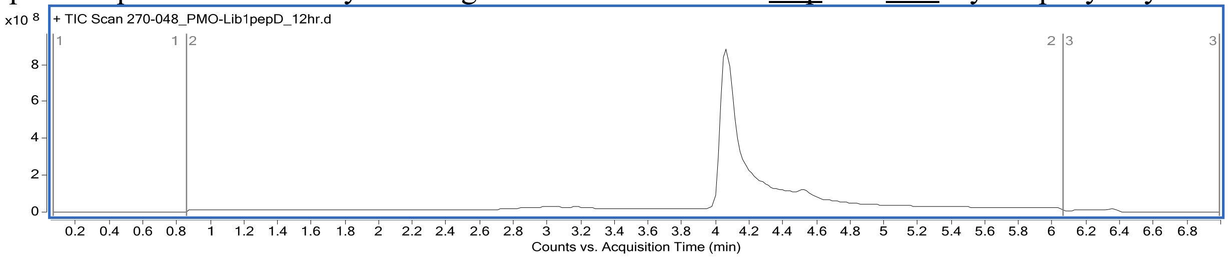

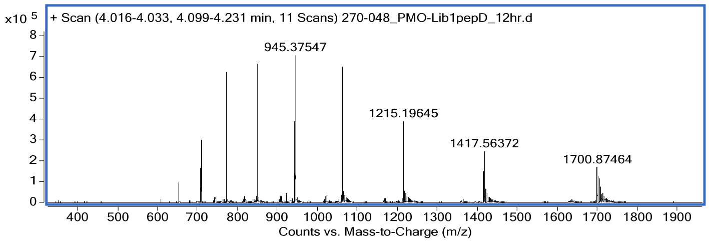

Spectral window of 4.02-4.10 min, observed mass calculated from computational deconvolution and confirmed by the $[M+9H]^{9+}$ ion, 945.3754

**PMO-CXP0**
Mass Expected: 8516.35
Mass Observed: 8516.41
Peptide sequence: 5azido-Gly-<u>Cit</u>-Arg-Ser-Lys-Asp-Val-Thr-His-His-Lys-Trp-Lys-Lys

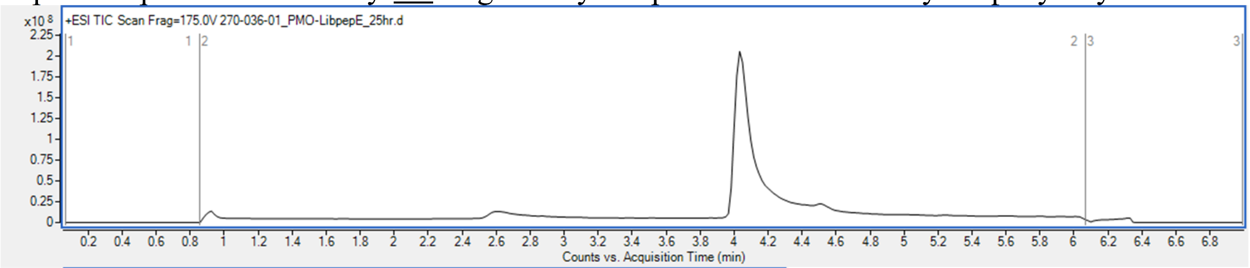

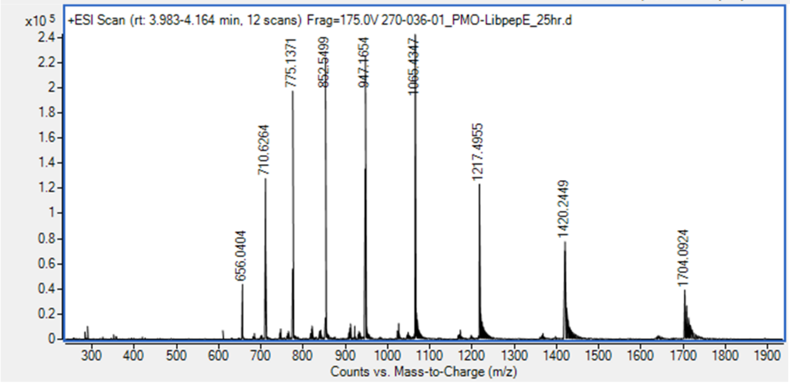

Spectral window of 3.98-4.16 min, observed mass calculated from computational deconvolution and confirmed by the $[M+9H]^{9+}$ ion, 947.1654

**PMO-D-CXP1**
Mass Expected: 8512.10
Mass Observed: 8511.99
Peptide sequence: 5azido-Gly-lys-gln-lys-thr-ser-<u>iph</u>-Gly-arg-Gly-<u>pip</u>-lys-trp-lys-lys

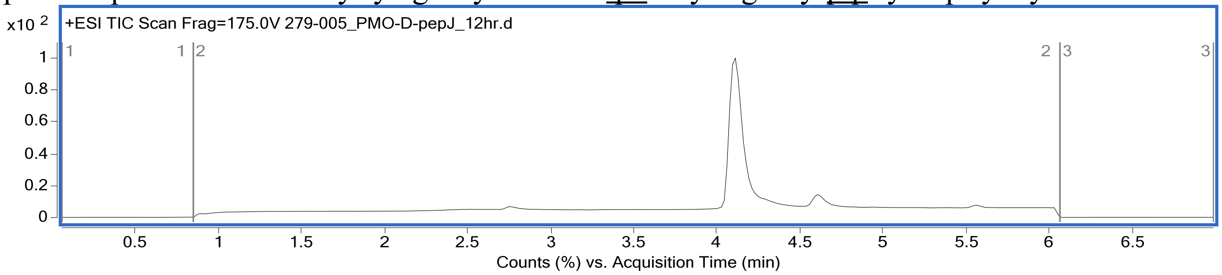

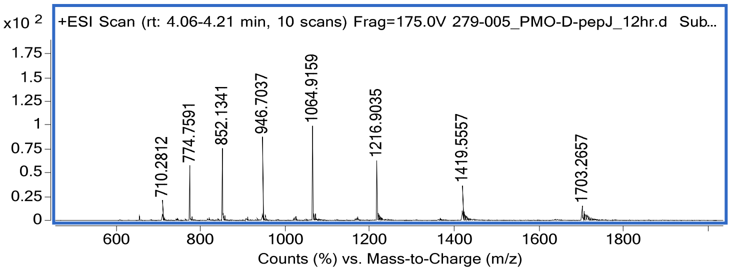

Spectral window of 4.06-4.21 min, observed mass calculated from computational deconvolution and confirmed by the $[M+8H]^{8+}$ ion, 1064.9159

**PMO-Scrambled-CXP1**
Mass Expected: 8512.10
Mass Observed: 8511.14
Peptide sequence: 5azido-Thr-Lys-Gly-Pip-Lys-Gly-Trp-Arg-Iph-Lys-Ser-Lys-Gly-Gln-Lys

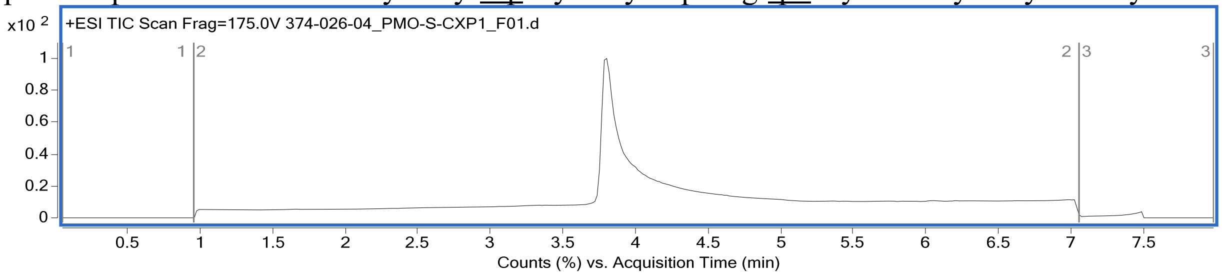

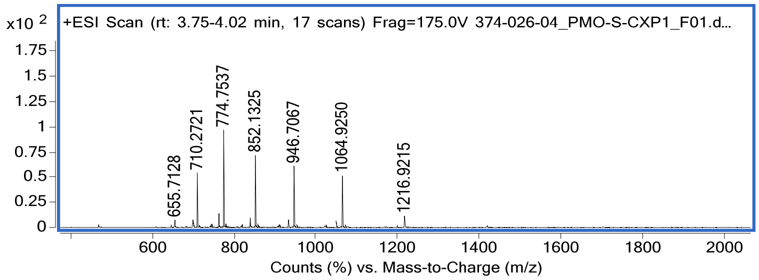

Spectral window of 3.75-4.02 min, observed mass calculated from computational deconvolution and confirmed by the $[M+8H]^{8+}$ ion, 1064.9250

**PMO-Canonical-CXP1**
Mass Expected: 8388.26
Mass Observed: 8388.23
Peptide sequence: 5azido-Gly-Lys-Gln-Lys-Thr-Ser-Phe-Gly-Arg-Gly-Lys-Lys-Trp-Lys-Lys

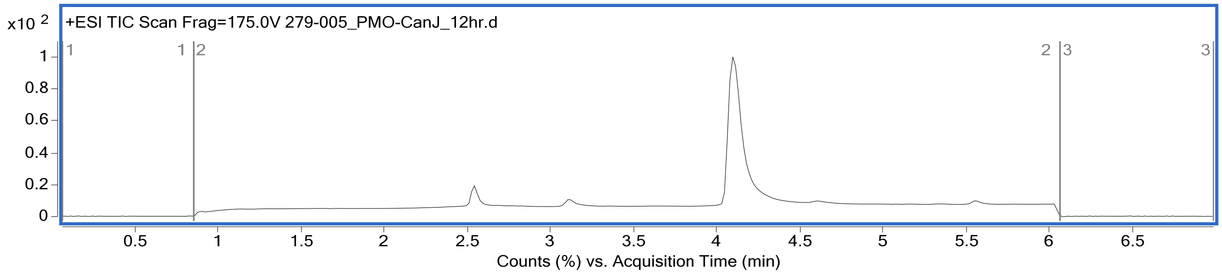

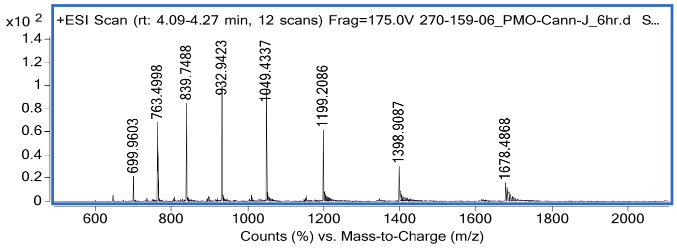

Spectral window of 4.09-4.27 min, observed mass calculated from computational deconvolution and confirmed by the $[M+8H]^{8+}$ ion, 1049.4337

**PMO-D-CXP2**
Mass Expected: 8506.32
Mass Observed: 8506.32
Peptide sequence: 5azido-Gly-asn-phe-lys-gln-nap-his-ala-Gly-dab-arg-lys-trp-lys-lys

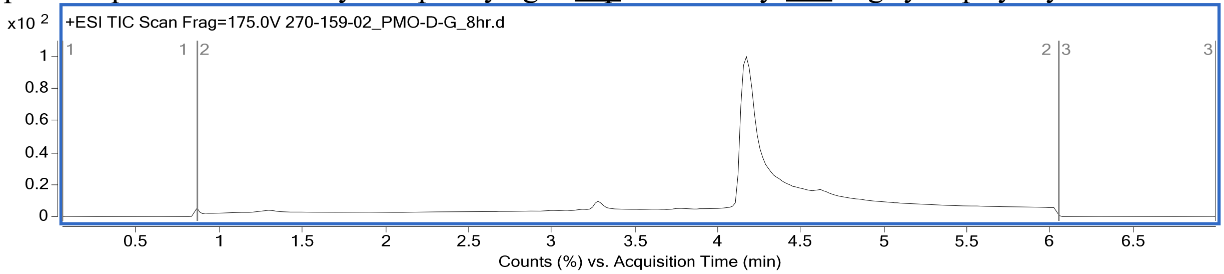

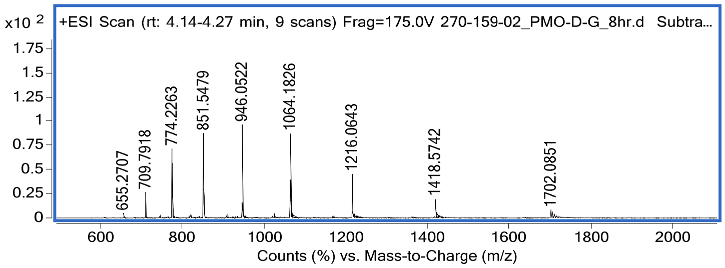

Spectral window of 4.14-4.27 min, observed mass calculated from computational deconvolution and confirmed by the $[M+8H]^{8+}$ ion, 1064.1826

**PMO-Scrambled-CXP2**
Mass Expected: 8506.32
Mass Observed: 8506.24
Peptide sequence: 5azido-Dab-Ala-Lys-Lys-Phe-Arg-Gly-His-Nap-Lys-Trp-Asn-Gln-Gly-Lys

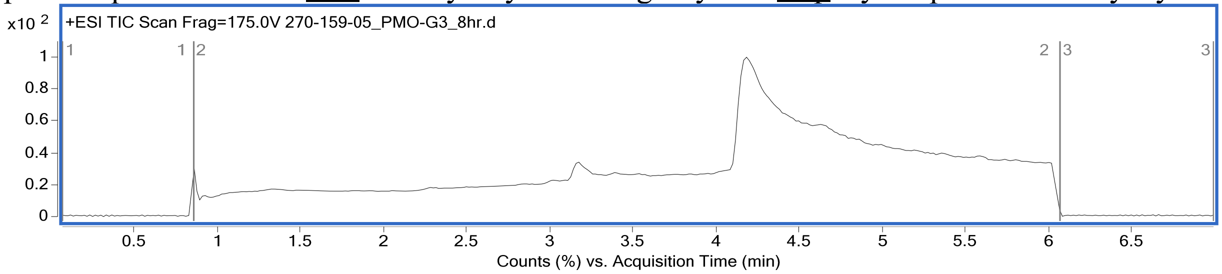

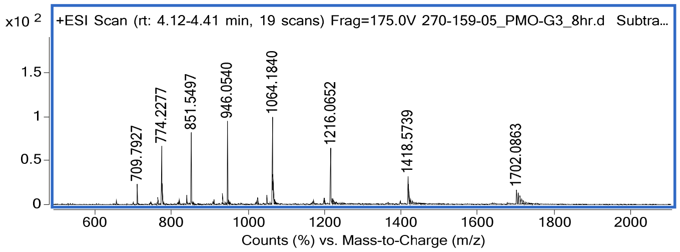

Spectral window of 4.12-4.41 min, observed mass calculated from computational deconvolution and confirmed by the $[M+8H]^{8+}$ ion, 1064.1840

**PMO-Canonical-CXP2**

Mass Expected: 8484.35

Mass Observed: 8483.40

Peptide sequence: 5azido-Gly-Asn-Phe-Lys-Gln-Phe-His-Ala-Gly-Lys-Arg-Lys-Trp-Lys-Lys

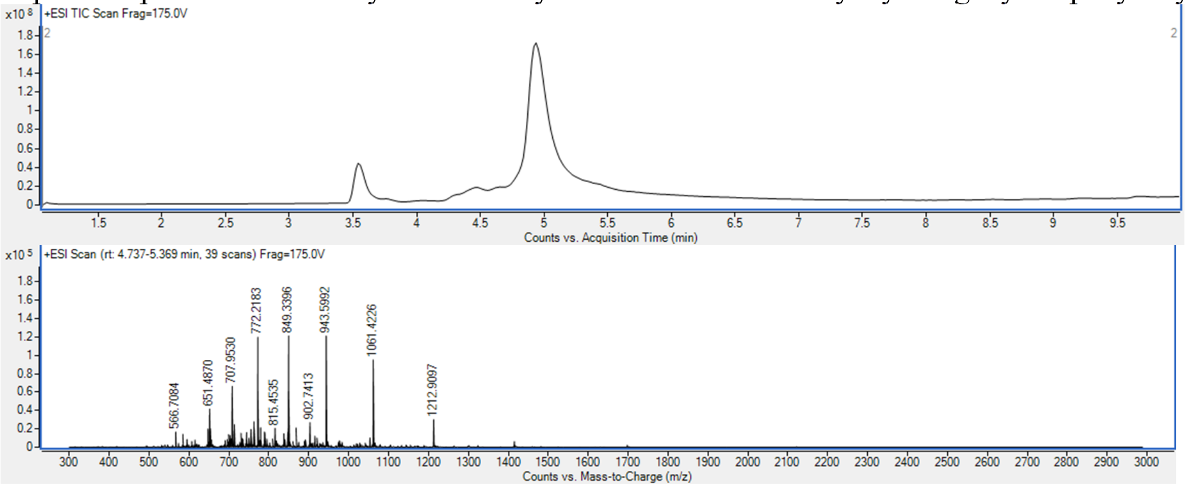

Spectral window of 4.74-5.36 min, observed mass calculated from computational deconvolution and confirmed by the $[M+8H]^{8+}$ ion, 1061.4226

**PMO-D-CXP3**

Mass Expected: 8311.09

Mass Observed: 8311.18

Peptide sequence: 5azido-Gly-Dab-Val-Ala-Arg-Asn-Asn-Dab-Thr-Lys-Gly-Lys-Trp-Lys-Lys

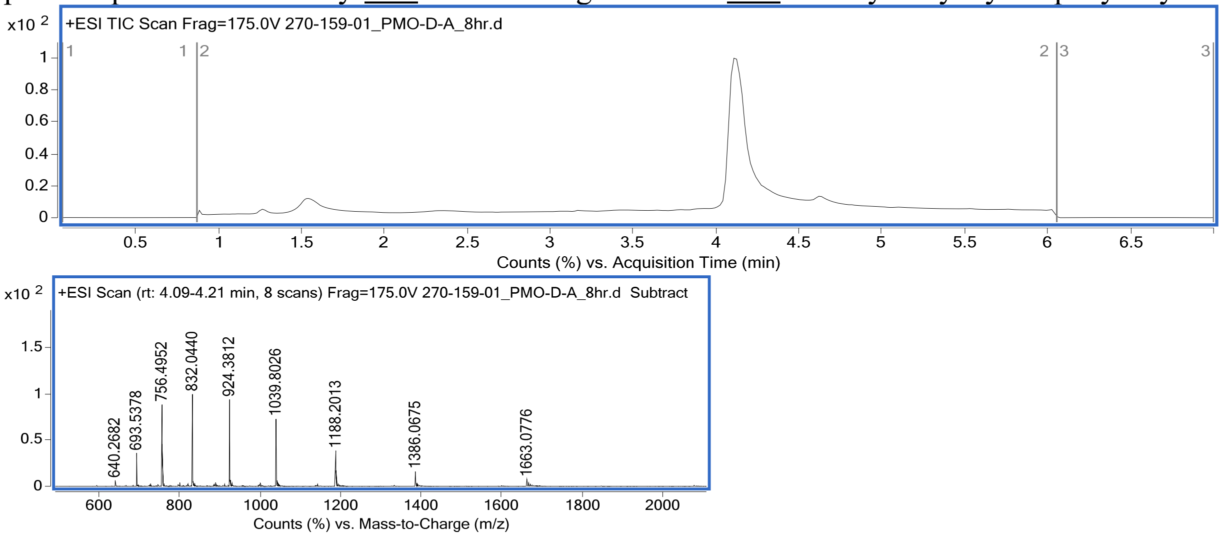

Spectral window of 4.09-4.21 min, observed mass calculated from computational deconvolution and confirmed by the $[M+8H]^{8+}$ ion, 1039.8026

**PMO-Scrambled-CXP3**
Mass Expected: 8311.09
Mass Observed: 8311.06
Peptide sequence: 5azido-Lys-Val-<u>Dab</u>-Lys-Asn-Lys-Ala-Trp-Arg-Asn-<u>Dab</u>-Gly-Thr-Lys-Gly

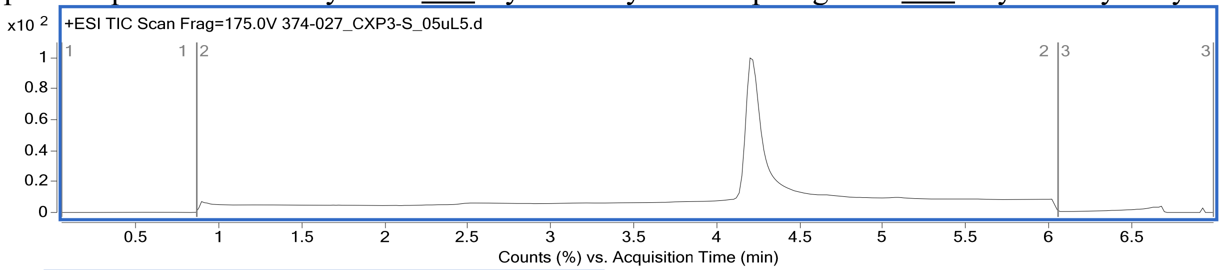

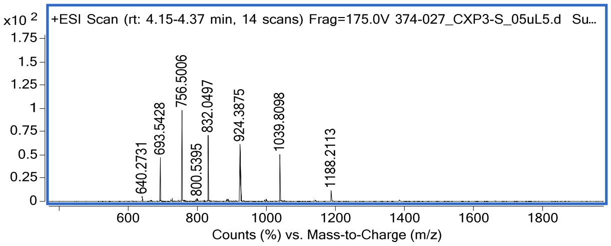

Spectral window of 4.15-4.37 min, observed mass calculated from computational deconvolution and confirmed by the $[M+8H]^{8+}$ ion, 1039.8098

**PMO-Canonical-CXP3**
Mass Expected: 8367.24
Mass Observed: 8367.26
Peptide sequence: 5azido-Gly-Lys-Val-Ala-Arg-Asn-Asn-Lys-Thr-Lys-Gly-Lys-Trp-Lys-Lys

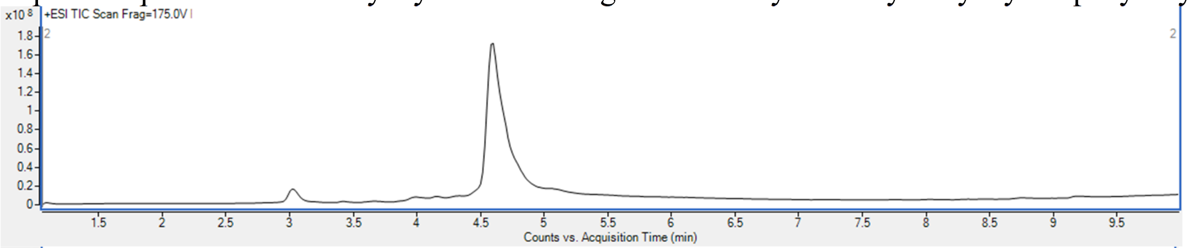

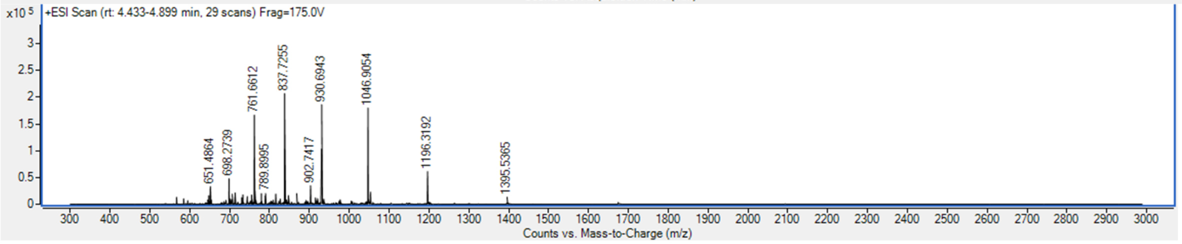

Spectral window of 4.43-4.90 min, observed mass calculated from computational deconvolution and confirmed by the $[M+8H]^{8+}$ ion, 1046.9054

**PMO-D-CXP4**

Mass Expected: 8385.29

Mass Observed: 8385.06

Peptide sequence: 5azido-Gly-cit-met-phe-Gly-pip-dab-dab-lys-ala-pro-lys-trp-lys-lys

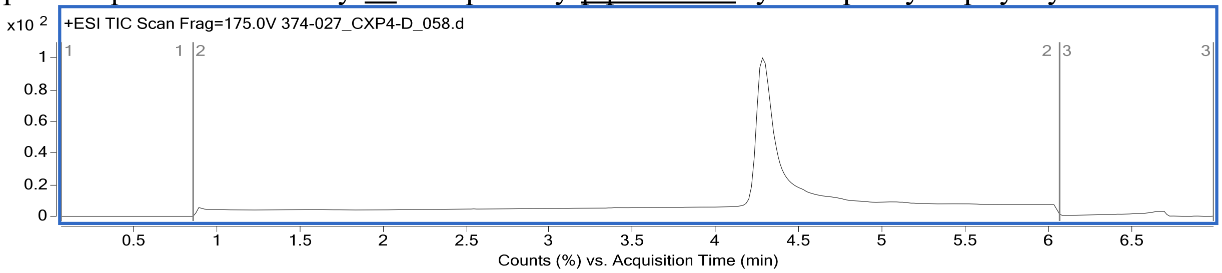

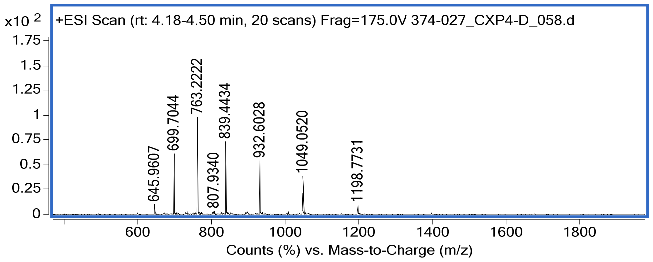

Spectral window of 4.18-4.50 min, observed mass calculated from computational deconvolution and confirmed by the $[M+8H]^{8+}$ ion, 1049.0520

**PMO-Scrambled-CXP4**

Mass Expected: 8385.29

Mass Observed: 8385.04

Peptide sequence: 5azido-Met-Lys-Cit-Ala-Pip-Trp-Lys-Phe-Dab-Gly-Lys-Dab-Gly-Pro-Lys

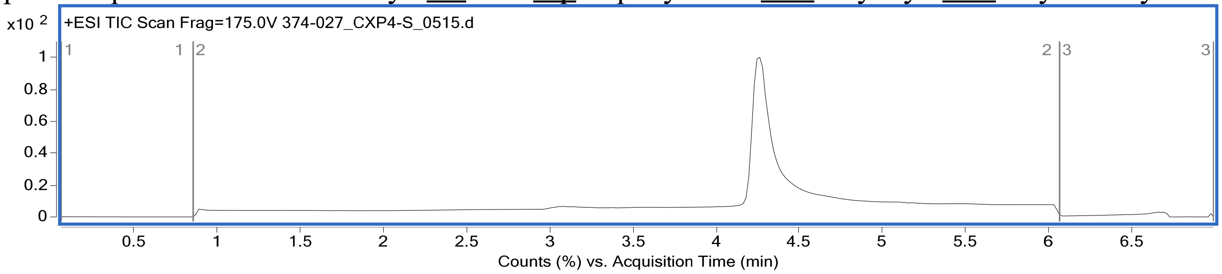

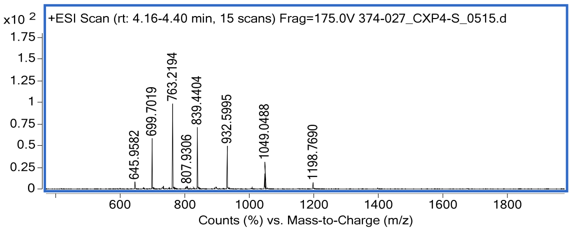

Spectral window of 4.16-4.40 min, observed mass calculated from computational deconvolution and confirmed by the $[M+8H]^{8+}$ ion, 1049.0488

**PMO-Canonical-CXP4**
Mass Expected: 8414.04
Mass Observed: 8414.12
Peptide sequence: 5azido-Gly-Gln-Met-Phe-Gly-Lys-Lys-Lys-Lys-Ala-Pro-Lys-Trp-Lys-Lys

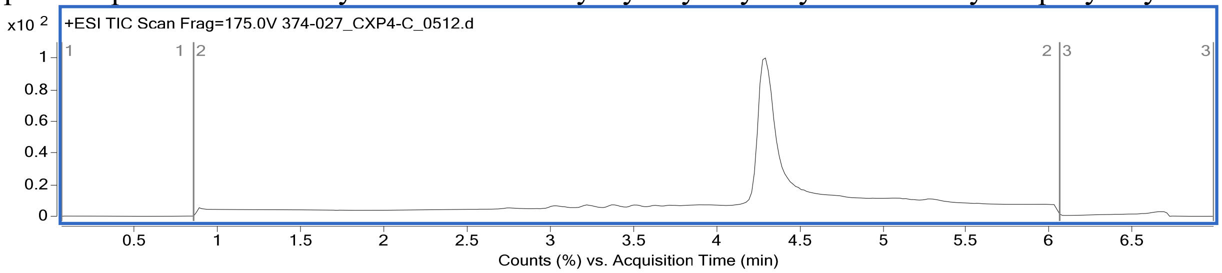

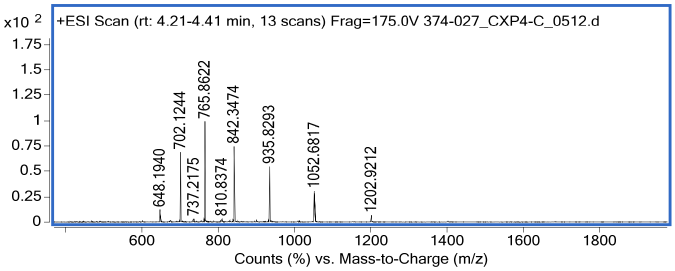

Spectral window of 4.21-4.41 min, observed mass calculated from computational deconvolution and confirmed by the $[M+8H]^{8+}$ ion, 1052.6817

**PMO-Penetratin**
Mass Expected: 8870.91
Mass Observed: 8870.79
Peptide sequence: 5azido-Arg-Gln-Ile-Lys-Ile-Trp-Phe-Gln-Asn-Arg-Arg-Met-Lys-Trp-Lys-Lys

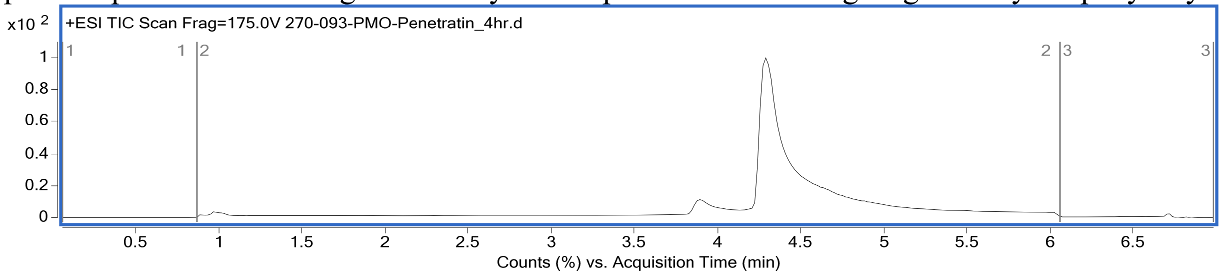

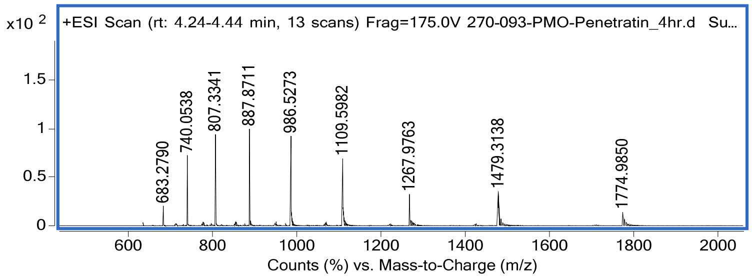

Spectral window of 4.24-4.44 min, observed mass calculated from computational deconvolution and confirmed by the $[M+8H]^{8+}$ ion, 1109.5982

**PMO-B-Peptide**
Mass Expected: 8259.1
Mass Observed: 8259.5
Peptide sequence: 5azido-Arg-Ahx-Arg-Arg-Bal-Arg-Arg-Ahx-Arg-Arg-Bal-Arg
Where Ahx represents Aminohexanoic acid and Bal represents Beta-alanine

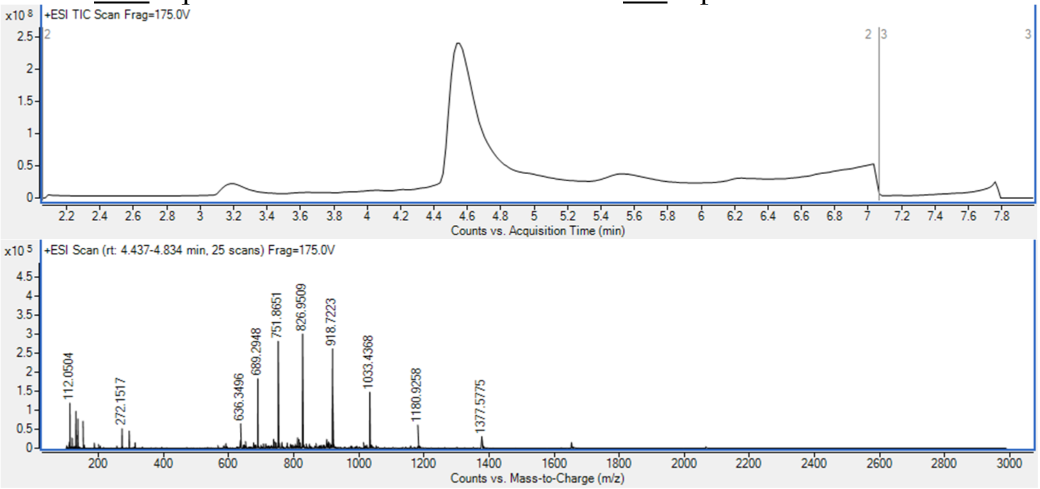

Spectral window of 4.44-4.83 min, observed mass calculated from computational deconvolution and confirmed by the $[M+8H]^{8+}$ ion, 1033.4368

**Example PMO-Library Fraction (3,000-member library, pool 10)**

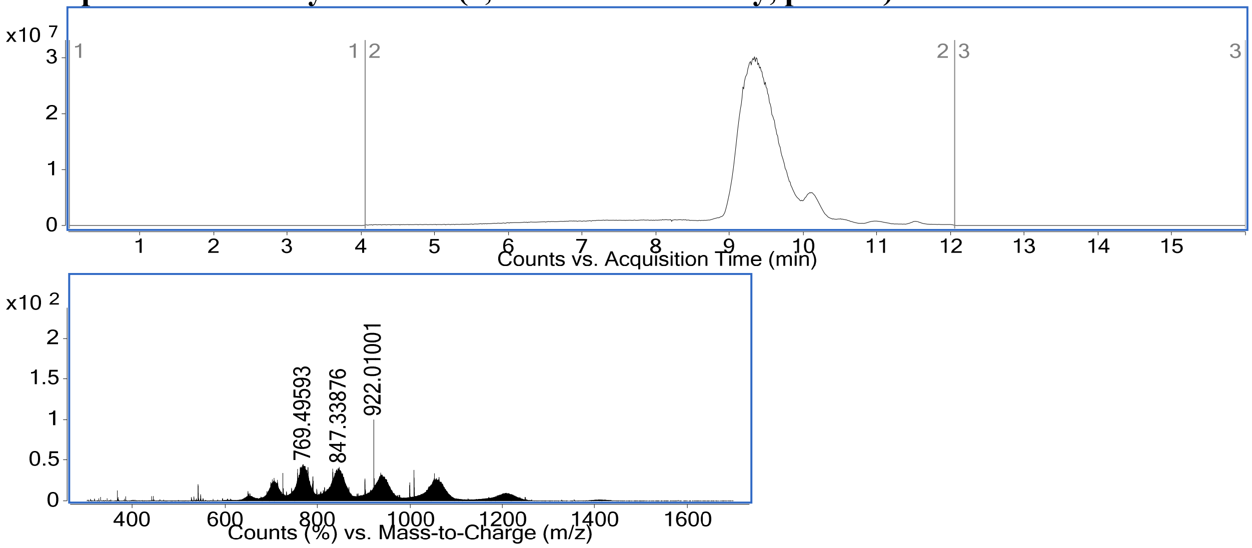

#### 4. Recovered peptide sequences from chromatographic resolution

**Library 1: 3,000-member library**

| Fraction Label | Fraction Retention Time (min) | Peptide sequence | Sequencing confidence score (ALC) | peptide mass (Da) |
|---|---|---|---|---|
| 01-1 | 01 - 10 | G(Pip)SDSANAGLSKWKK | 99 | 1697.90 |
| 01-1 | 01 - 10 | GFTNTH(Dab)LEDFKWKK | 99 | 1974.02 |
| 01-1 | 01 - 10 | GKQTFESLALNKWKK | 98 | 1901.06 |
| 01-1 | 01 - 10 | GSKFRNVSNEDKWKK | 98 | 1946.02 |
| 01-1 | 01 - 10 | GK(Dab)(Nap)DNELTSEKWKK | 98 | 1983.03 |
| 01-1 | 01 - 10 | G(Iph)(Dab)SNQN(Cit)(Cit)ETKWKK | 98 | 2129.94 |
| 01-1 | 01 - 10 | GGAG(Nap)DSNTHFKWKK | 97 | 1852.91 |
| 01-1 | 01 - 10 | GP(Pip)FLSENGQMKWKK | 97 | 1898.99 |
| 01-1 | 01 - 10 | GKTFVEPMGDKKWKK | 97 | 1902.03 |
| 01-1 | 01 - 10 | GLTKMGNRVDEKWKK | 97 | 1913.04 |
| 01-1 | 01 - 10 | GLNSHVNEV(Iph)GKWKK | 97 | 1991.90 |
| 01-1 | 01 - 10 | G(Dab)VGQ(Cit)M(Iph)GQEKWKK | 97 | 2028.90 |
| 01-1 | 01 - 10 | GQ(Dab)AQ(Iph)(Cit)EGMFKWKK | 97 | 2090.91 |
| 01-1 | 01 - 10 | G(Nap)TKE(Nap)EG(Cit)QLKWKK | 97 | 2106.12 |
| 01-1 | 01 - 10 | GVAKEQGNLLFKWKK | 96 | 1869.07 |
| 01-1 | 01 - 10 | G(Pal)G(Cit)SADANK(Iph)KWKK | 96 | 1990.88 |
| 01-1 | 01 - 10 | GERAQ(Iph)TEPDHKWKK | 96 | 2105.90 |
| 01-1 | 01 - 10 | GT(Iph)(Cit)NFEEHATKWKK | 96 | 2128.91 |
| 01-1 | 01 - 10 | GEVSHGTGMTPKWKK | 95 | 1765.90 |
| 01-1 | 01 - 10 | GQADGNFSA(Pip)QKWKK | 95 | 1813.93 |
| 01-1 | 01 - 10 | GLHT(Dab)DSDSPQKWKK | 95 | 1849.95 |
| 01-1 | 01 - 10 | GPLQKQ(Cit)ELAGKWKK | 95 | 1891.09 |
| 01-1 | 01 - 10 | GVGNA(Iph)(Pip)DPAVKWKK | 95 | 1891.87 |
| 01-1 | 01 - 10 | GGKS(Cit)DDKQSFKWKK | 95 | 1919.01 |
| 01-1 | 01 - 10 | GMT(Cit)DAEQV(Dab)(Nap)KWKK | 95 | 1998.03 |
| 01-1 | 01 - 10 | GMRDL(Cit)GS(Iph)GNKWKK | 95 | 2029.89 |
| 01-1 | 01 - 10 | GKTLF(Cit)APND(Iph)KWKK | 95 | 2085.98 |
| 01-1 | 01 - 10 | GNTF(Iph)NKMGQEKWKK | 95 | 2091.90 |
| 01-1 | 01 - 10 | GQQGNPG(Pip)EFTKWKK | 94 | 1853.96 |
| 01-1 | 01 - 10 | GQENQMKVPLQKWKK | 94 | 1965.07 |
| 01-1 | 01 - 10 | GLLKEEDR(Cit)(Cit)(Iph)KWKK | 94 | 2240.08 |

| | | | | |
|---|---|---|---|---|
| 01-1 | 01 - 10 | GLVVGSGDS(Pip)MKWKK | 93 | 1740.95 |
| 01-1 | 01 - 10 | GSDSKEMVDG(Pip)KWKK | 93 | 1843.94 |
| 01-1 | 01 - 10 | GKNTMDD(Cit)(Pip)T(Nap)KWKK | 93 | 2055.05 |
| 01-1 | 01 - 10 | GAGSVMD(Dab)PSNKWKK | 92 | 1727.89 |
| 01-1 | 01 - 10 | GSHALVLGEPGKWKK | 92 | 1729.97 |
| 01-1 | 01 - 10 | G(Cit)NNAGMLGDRKWKK | 92 | 1854.97 |
| 01-1 | 01 - 10 | GGEHV(Nap)PSG(Cit)QKWKK | 92 | 1915.00 |
| 01-1 | 01 - 10 | GL(Dab)APSLSTQGKWKK | 91 | 1723.98 |
| 01-1 | 01 - 10 | G(Cit)TL(Dab)A(Dab)AEEVKWKK | 91 | 1840.04 |
| 01-1 | 01 - 10 | G(Iph)P(Dab)VEGEND(Pip)KWKK | 91 | 2008.88 |
| 01-1 | 01 - 10 | G(Cit)RSMF(Cit)(Nap)GNEKWKK | 91 | 2102.07 |
| 01-1 | 01 - 10 | GQRDS(Cit)DDHP(Iph)KWKK | 91 | 2149.91 |
| 01-1 | 01 - 10 | GGSAERPLTS(Cit)KWKK | 90 | 1825.01 |
| 01-1 | 01 - 10 | GTSF(Dab)DTQFN(Iph)KWKK | 90 | 2082.89 |
| 01-1 | 01 - 10 | G(Iph)H(Cit)E(Cit)PMVPNKWKK | 90 | 2160.97 |
| 01-1 | 01 - 10 | GMDKPSVFTGLKWKK | 89 | 1845.01 |
| 01-1 | 01 - 10 | GFT(Iph)SGQNNTAKWKK | 89 | 1962.84 |
| 01-1 | 01 - 10 | GP(Nap)P(Iph)(Dab)NAAADKWKK | 89 | 1975.87 |
| 01-1 | 01 - 10 | GNKA(Iph)GVA(Cit)LEKWKK | 89 | 1981.95 |
| 01-1 | 01 - 10 | G(Pip)VNAPGGALEKWKK | 87 | 1703.96 |
| 01-1 | 01 - 10 | GAKKGETFAD(Cit)KWKK | 87 | 1874.03 |
| 01-1 | 01 - 10 | GPNRDMGQPFLKWKK | 87 | 1925.02 |
| 01-1 | 01 - 10 | G(Pip)RCL(Iph)EDGHFKWKK | 87 | 2125.93 |
| 01-1 | 01 - 10 | GFNHED(Cit)PPNGKWKK | 86 | 1933.97 |
| 01-1 | 01 - 10 | GQTED(Cit)E(Pip)VKVKWKK | 86 | 1981.09 |
| 01-1 | 01 - 10 | GVHENTPDKF(Nap)KWKK | 86 | 2034.06 |
| 01-1 | 01 - 10 | GDVAPH(Dab)PQGDKWKK | 85 | 1785.94 |
| 01-1 | 01 - 10 | GMVDAYGAVD(Pip)KWKK | 85 | 1816.94 |
| 01-1 | 01 - 10 | GVGHDGQTEFQKWKK | 85 | 1867.94 |
| 01-1 | 01 - 10 | GGLN(Cit)MDPTRGKWKK | 85 | 1868.00 |
| 01-1 | 01 - 10 | GTQKQANVQETKWKK | 85 | 1897.03 |
| 01-1 | 01 - 10 | GKCG(Cit)D(Cit)AKVLKWKK | 85 | 1898.08 |
| 01-1 | 01 - 10 | GQHDTMG(Iph)KPEKWKK | 85 | 2065.88 |
| 01-1 | 01 - 10 | GQRDS(Cit)DDSF(Iph)KWKK | 85 | 2149.89 |
| 01-1 | 01 - 10 | GV(Cit)HMETN(Iph)N(Cit)KWKK | 85 | 2181.95 |
| 01-1 | 01 - 10 | G(Pal)SGE(Dab)SPTQPKWKK | 84 | 1800.94 |
| 01-1 | 01 - 10 | GAFPRDDQ(Pip)LTKWKK | 84 | 1939.05 |
| 01-1 | 01 - 10 | G(Iph)KMNAGNDQAKWKK | 84 | 1971.84 |
| 01-1 | 01 - 10 | GSSDD(Cit)P(Iph)N(Dab)(Nap)KWKK | 84 | 2111.88 |

| | | | | |
|---|---|---|---|---|
| 01-1 | 01 - 10 | GKMSPNLNATEKWKK | 83 | 1854.99 |
| 01-1 | 01 - 10 | GKAKGETFAD(Cit)KWKK | 83 | 1874.03 |
| 01-1 | 01 - 10 | G(Pal)ES(Dab)DF(Cit)ANHKWKK | 83 | 1974.99 |
| 01-1 | 01 - 10 | GEP(Nap)KED(Pip)TT(Cit)KWKK | 83 | 2050.07 |
| 01-1 | 01 - 10 | GSSKEPS(Dab)QDVKWKK | 82 | 1826.98 |
| 01-1 | 01 - 10 | G(Cit)RTVQE(Dab)DAVKWKK | 82 | 1925.07 |
| 01-1 | 01 - 10 | GQTDPHN(Cit)L(Cit)FKWKK | 82 | 2036.08 |
| 01-1 | 01 - 10 | GPS(Nap)PDG(Dab)AVGKWKK | 81 | 1746.93 |
| 01-1 | 01 - 10 | GVQAF(Pip)LDGGPKWKK | 80 | 1779.99 |
| 01-1 | 01 - 10 | GSKLAGS(Cit)LKYKWKK | 80 | 1874.10 |
| 01-1 | 01 - 10 | GF(Cit)GLDT(Dab)(Pal)LCKWKK | 79 | 1924.03 |
| 01-1 | 01 - 10 | G(Cit)TN(Cit)SAQDRNKWKK | 79 | 1970.03 |
| 01-1 | 01 - 10 | GEFEAL(Cit)(Nap)A(Cit)LKWKK | 79 | 2054.12 |
| 01-1 | 01 - 10 | G(Pip)D(Iph)SELMQD(Dab)KWKK | 79 | 2086.89 |
| 01-1 | 01 - 10 | GVSTAGQTVMQKWKK | 78 | 1771.95 |
| 01-1 | 01 - 10 | GLDQDMPADKKKWKK | 78 | 1911.01 |
| 01-1 | 01 - 10 | GLTFAPDRVDEKWKK | 78 | 1913.03 |
| 01-1 | 01 - 10 | GA(Cit)SRDDQ(Pip)LTKWKK | 78 | 1939.05 |
| 01-1 | 01 - 10 | GGSAERLPTS(Cit)KRTG | 77 | 1697.89 |
| 01-1 | 01 - 10 | GTL(Nap)STVTAPQKWKK | 77 | 1865.03 |
| 01-1 | 01 - 10 | G(Cit)LP(Pal)GT(Pip)GKDKWKK | 77 | 1869.05 |
| 01-1 | 01 - 10 | G(Iph)SSADT(Dab)(Cit)ANKWKK | 77 | 1945.84 |
| 01-1 | 01 - 10 | GKATSLGEVGGKWKK | 76 | 1668.94 |
| 01-1 | 01 - 10 | GVSCAEKFGQVKWKK | 76 | 1817.97 |
| 01-1 | 01 - 10 | GSDGRMMVDG(Pip)KWKK | 75 | 1843.93 |
| 01-1 | 01 - 10 | GKNDWTNQLHLKWKK | 75 | 2019.09 |
| 01-1 | 01 - 10 | G(Iph)(Cit)LTVND(Cit)KEKWKK | 75 | 2156.01 |
| 01-1 | 01 - 10 | GNEFTGGSSEHKWKK | 74 | 1814.88 |
| 01-1 | 01 - 10 | GLKA(Pip)QQFA(Cit)VKWKK | 74 | 1938.14 |
| 01-1 | 01 - 10 | GLKVEPQFA(Cit)VKWKK | 73 | 1938.13 |
| 01-1 | 01 - 10 | GLVGVSGDNVMKWKK | 72 | 1740.95 |

| | | | | |
|---|---|---|---|---|
| 01-1 | 01 - 10 | GV(Dab)KGETFAD(Cit)KWKK | 72 | 1874.03 |
| 01-1 | 01 - 10 | G(Dab)FA(Pip)DNPE(Nap)LKWKK | 72 | 1979.05 |
| 01-1 | 01 - 10 | GFDLRDVPPQFKWKK | 72 | 1984.08 |
| 01-1 | 01 - 10 | GTLSLQG(Cit)SGNKWKK | 71 | 1783.98 |
| 01-1 | 01 - 10 | GVSCRD(Cit)AKVLKWKK | 71 | 1898.08 |
| 01-1 | 01 - 10 | G(Iph)GGPNKDFNNKWKK | 71 | 1985.85 |
| 01-1 | 01 - 10 | GKQ(Pal)(Pip)CSLALNKWKK | 70 | 1901.06 |
| 01-1 | 01 - 10 | GF(Dab)QRDVPPQFKWKK | 70 | 1984.09 |
| 01-1 | 01 - 10 | GNTF(Iph)NKCTQEKWKK | 70 | 2107.89 |
| 02-1 | 11 - 15 | GL(Dab)ASPLSTQGKWKK | 97 | 1723.98 |
| 02-1 | 11 - 15 | GEKVPDAR(Dab)DTKWKK | 97 | 1881.03 |
| 02-1 | 11 - 15 | GNEEDF(Pip)NGRSKWKK | 96 | 1943.97 |
| 02-1 | 11 - 15 | GVLPGK(Dab)LSDLKWKK | 95 | 1792.08 |
| 02-1 | 11 - 15 | G(Dab)DMASNHTVTKWKK | 95 | 1825.94 |
| 02-1 | 11 - 15 | GLRDQDKRGDLKWKK | 95 | 1966.10 |
| 02-1 | 11 - 15 | G(Iph)KSQPSALQSKWKK | 95 | 1968.92 |
| 02-1 | 11 - 15 | GL(Dab)FPA(Dab)GDF(Cit)KWKK | 94 | 1874.04 |
| 02-1 | 11 - 15 | GATHRVDNMSQKWKK | 94 | 1908.99 |
| 02-1 | 11 - 15 | G(Nap)HAVGEENQHKWKK | 94 | 1967.99 |
| 02-1 | 11 - 15 | GLKQEND(Pip)FT(Nap)KWKK | 94 | 2068.10 |
| 02-1 | 11 - 15 | GAHGNEHE(Iph)(Cit)SKWKK | 93 | 2060.86 |
| 02-1 | 11 - 15 | GHE(Iph)LSAQQKEKWKK | 93 | 2092.95 |
| 02-1 | 11 - 15 | G(Cit)(Iph)DEKTGHMNKWKK | 93 | 2111.90 |
| 02-1 | 11 - 15 | GLTEQQGF(Cit)PRKWKK | 91 | 1983.09 |
| 02-1 | 11 - 15 | GKL(Cit)E(Dab)EFGNFKWKK | 91 | 1991.09 |
| 02-1 | 11 - 15 | G(Dab)DQEQT(Cit)(Cit)(Dab)(Iph)KWKK | 91 | 2157.97 |
| 02-1 | 11 - 15 | G(Nap)PSNFAESHVKWKK | 90 | 1934.99 |
| 02-1 | 11 - 15 | GGGNSEHKE(Iph)GKWKK | 90 | 1937.81 |
| 02-1 | 11 - 15 | GQKHTTDP(Cit)P(Cit)KWKK | 90 | 1988.08 |
| 02-1 | 11 - 15 | G(Iph)KT(Cit)QQTGGTKWKK | 90 | 2000.92 |
| 02-1 | 11 - 15 | GVERQGN(Iph)P(Dab)DKWKK | 90 | 2037.91 |

| | | | | |
|---|---|---|---|---|
| 02-1 | 11 - 15 | G(Pip)NPEFFHFQEKWKK | 90 | 2071.05 |
| 02-1 | 11 - 15 | GGPDQAHE(Cit)SHKWKK | 89 | 1884.95 |
| 02-1 | 11 - 15 | GHTSSRDPLMMKWKK | 89 | 1924.99 |
| 02-1 | 11 - 15 | G(Dab)NHANHQQMDKWKK | 89 | 1944.96 |
| 02-1 | 11 - 15 | GP(Dab)(Dab)PT(Iph)(Cit)ADAKWKK | 89 | 1951.90 |
| 02-1 | 11 - 15 | G(Iph)PSVNSE(Dab)PTKWKK | 89 | 1953.87 |
| 02-1 | 11 - 15 | GPTLAPSFRTPKWKK | 88 | 1837.05 |
| 02-1 | 11 - 15 | GDSTLPSG(Pip)T(Iph)KWKK | 87 | 1926.86 |
| 02-1 | 11 - 15 | GQMSK(Cit)EHSLEKWKK | 87 | 1996.04 |
| 02-1 | 11 - 15 | G(Cit)VN(Pip)ATGGGAKWKK | 86 | 1679.93 |
| 02-1 | 11 - 15 | GTE(Cit)VGPPAPHKWKK | 85 | 1811.99 |
| 02-1 | 11 - 15 | G(Cit)(Cit)AKQE(Dab)DAVKWKK | 85 | 1925.07 |
| 02-1 | 11 - 15 | G(Dab)PSQ(Cit)MSGNGKWKK | 83 | 1784.92 |
| 02-1 | 11 - 15 | G(Cit)ESSVNA(Cit)L(Dab)KWKK | 83 | 1884.04 |
| 02-1 | 11 - 15 | GQHPLSKPD(Cit)MKWKK | 83 | 1960.06 |
| 02-1 | 11 - 15 | GPGVAKGANTGKWKK | 82 | 1621.92 |
| 02-1 | 11 - 15 | G(Cit)APQ(Cit)(Iph)HQQSKWKK | 82 | 2132.96 |
| 02-1 | 11 - 15 | GGNGHNFF(Cit)SPKWKK | 81 | 1883.97 |
| 02-1 | 11 - 15 | GEF(Cit)FVNKG(Cit)TKWKK | 81 | 2006.10 |
| 02-1 | 11 - 15 | GPPSLQKTS(Nap)PKWKK | 80 | 1902.06 |
| 02-1 | 11 - 15 | GFNHED(Cit)PPNGKWKK | 80 | 1933.97 |
| 02-1 | 11 - 15 | GVFGSQGQHATKWKK | 79 | 1781.94 |
| 02-1 | 11 - 15 | G(Pip)DDGA(Iph)NALHKWKK | 79 | 1961.85 |
| 02-1 | 11 - 15 | G(Cit)DTESFHSA(Iph)KWKK | 79 | 2073.87 |
| 02-1 | 11 - 15 | GKTFVNENGDKKWKK | 78 | 1902.02 |
| 02-1 | 11 - 15 | GKQGE(Iph)AKSG(Dab)KWKK | 77 | 1927.90 |
| 02-1 | 11 - 15 | GN(Dab)TNVRNDLFKWKK | 76 | 1943.06 |
| 02-1 | 11 - 15 | GVMKQGPPAPHKWKK | 74 | 1812.01 |
| 02-1 | 11 - 15 | GG(Dab)VTHNAPS(Nap)KWKK | 74 | 1829.98 |
| 02-1 | 11 - 15 | G(Dab)AS(Pip)(Dab)(Cit)QPG(Nap)KWKK | 74 | 1890.05 |
| 02-1 | 11 - 15 | GMHEVET(Dab)NNAKWKK | 74 | 1894.96 |

| | | | | |
|---|---|---|---|---|
| 02-1 | 11 - 15 | GAQFN(Iph)GVGAEKWKK | 74 | 1915.83 |
| 02-1 | 11 - 15 | G(Pip)LAMGSG(Dab)GVKWKK | 73 | 1667.94 |
| 02-1 | 11 - 15 | GESLAGFKD(Dab)TKWKK | 73 | 1817.99 |
| 02-1 | 11 - 15 | GEGS(Nap)GV(Iph)SAGKWKK | 73 | 1883.80 |
| 02-1 | 11 - 15 | GDNKS(Cit)SEVK(Nap)KWKK | 73 | 2011.08 |
| 02-1 | 11 - 15 | G(Pip)QFKPKNFVDKWKK | 72 | 1999.13 |
| 02-1 | 11 - 15 | G(Nap)SPMET(Dab)PMAKWKK | 71 | 1910.96 |
| 02-1 | 11 - 15 | GNEMP(Dab)DH(Nap)LVKWKK | 71 | 2002.04 |
| 02-1 | 11 - 15 | GAEFQTTPG(Dab)KKWKK | 70 | 1829.01 |
| 02-1 | 11 - 15 | GKEFSPVMAMPKWKK | 70 | 1887.00 |
| 02-1 | 11 - 15 | GP(Pal)QAESPQ(Dab)FKWKK | 70 | 1902.00 |
| 02-1 | 11 - 15 | GGVGE(Cit)(Nap)ESTMKWKK | 70 | 1913.96 |
| 02-1 | 11 - 15 | GVPTPT(Iph)(Cit)ADAKWKK | 70 | 1951.89 |
| 02-1 | 11 - 15 | GS(Nap)(Cit)EQGSFG(Nap)KWKK | 70 | 2013.00 |
| 03-1 | 16 - 18 | GVGMVSK(Dab)GDGKWKK | 73 | 1699.93 |
| 03-1 | 16 - 18 | G(Dab)NA(Dab)SPVSAEKWKK | 94 | 1724.94 |
| 03-1 | 16 - 18 | GNMGAGQGRTTKWKK | 95 | 1742.91 |
| 03-1 | 16 - 18 | GTAAAAP(Cit)P(Dab)FKWKK | 88 | 1752.99 |
| 03-1 | 16 - 18 | GDGV(Dab)EFGRAGKWKK | 71 | 1757.94 |
| 03-1 | 16 - 18 | GL(Cit)TGAGLHPTKWKK | 74 | 1774.01 |
| 03-1 | 16 - 18 | GALVATHAHTEKWKK | 92 | 1799.99 |
| 03-1 | 16 - 18 | GPKAFFAVDGVKWKK | 86 | 1801.02 |
| 03-1 | 16 - 18 | GEKDGP(Pip)STAMKWKK | 72 | 1811.95 |
| 03-1 | 16 - 18 | G(Dab)(Cit)DPG(Pip)STAMKWKK | 91 | 1811.96 |
| 03-1 | 16 - 18 | GTRDGP(Pip)STAMKWKK | 83 | 1811.96 |
| 03-1 | 16 - 18 | GGLGN(Cit)K(Dab)G(Cit)DKWKK | 96 | 1825.02 |
| 03-1 | 16 - 18 | GSVQ(Dab)(Cit)(Pip)TSGDKWKK | 86 | 1826.99 |
| 03-1 | 16 - 18 | G(Pip)AELTQPHVGKWKK | 92 | 1828.02 |
| 03-1 | 16 - 18 | GNTS(Dab)FTGSQHKWKK | 76 | 1828.94 |
| 03-1 | 16 - 18 | G(Dab)(Nap)TGG(Dab)LDKVKWKK | 84 | 1837.05 |
| 03-1 | 16 - 18 | GLSLFSTP(Dab)PMKWKK | 87 | 1843.03 |
| 03-1 | 16 - 18 | GDGTRRNDG(Dab)LKWKK | 70 | 1854.01 |

| | | | | |
|---|---|---|---|---|
| 03-1 | 16 - 18 | GVP(Dab)SKE(Cit)SLPKWKK | 81 | 1864.08 |
| 03-1 | 16 - 18 | G(Iph)SSEVGTLGVKWKK | 85 | 1871.85 |
| 03-1 | 16 - 18 | GQHLKE(Cit)GAVPKWKK | 95 | 1886.08 |
| 03-1 | 16 - 18 | GVGHDSR(Nap)GSMKWKK | 92 | 1892.96 |
| 03-1 | 16 - 18 | GGMVG(Iph)G(Pip)QPVKWKK | 93 | 1893.87 |
| 03-1 | 16 - 18 | GGFEKSS(Pip)FESKWKK | 96 | 1893.99 |
| 03-1 | 16 - 18 | GKTTVEAHG(Cit)FKWKK | 97 | 1897.04 |
| 03-1 | 16 - 18 | G(Pip)EEASNSFL(Pip)KWKK | 89 | 1899.01 |
| 03-1 | 16 - 18 | GPVSQL(Pip)(Iph)GTGKWKK | 93 | 1907.90 |
| 03-1 | 16 - 18 | G(Cit)VQ(Cit)TFTGGHKWKK | 85 | 1911.03 |
| 03-1 | 16 - 18 | GG(Dab)FQLSEPERKWKK | 93 | 1913.04 |
| 03-1 | 16 - 18 | GHTSSRPDFPMKWKK | 71 | 1924.98 |
| 03-1 | 16 - 18 | GLAFHNQHSEPKWKK | 94 | 1930.01 |
| 03-1 | 16 - 18 | GQELDTMSM(Pip)TKWKK | 72 | 1931.97 |
| 03-1 | 16 - 18 | GVGDA(Iph)PPLKLKWKK | 89 | 1932.96 |
| 03-1 | 16 - 18 | G(Cit)NLPVQRGVFKWKK | 87 | 1937.12 |
| 03-1 | 16 - 18 | G(Nap)(Dab)GGNLNDRFKWKK | 95 | 1940.03 |
| 03-1 | 16 - 18 | GQ(Iph)SSTQP(Dab)VAKWKK | 94 | 1940.89 |
| 03-1 | 16 - 18 | GEHSPT(Pip)(Cit)PQLKWKK | 71 | 1942.06 |
| 03-1 | 16 - 18 | GDVTNVRNDLFKWKK | 71 | 1943.05 |
| 03-1 | 16 - 18 | G(Dab)NTNVRNDLFKWKK | 93 | 1943.06 |
| 03-1 | 16 - 18 | GQF(Iph)EQ(Dab)TGGGKWKK | 80 | 1946.84 |
| 03-1 | 16 - 18 | G(Cit)TGS(Cit)(Dab)(Nap)SFSKWKK | 91 | 1947.02 |
| 03-1 | 16 - 18 | GTPQPNMSFFKKWKK | 78 | 1947.03 |
| 03-1 | 16 - 18 | G(Cit)(Dab)SL(Cit)TQEKSKWKK | 77 | 1957.10 |
| 03-1 | 16 - 18 | GAGD(Dab)LDQ(Iph)P(Pip)KWKK | 76 | 1964.89 |
| 03-1 | 16 - 18 | G(Dab)FTP(Cit)TNNHEKWKK | 93 | 1967.02 |
| 03-1 | 16 - 18 | GKE(Pip)A(Nap)TPPN(Cit)KWKK | 86 | 1987.09 |
| 03-1 | 16 - 18 | G(Cit)VFEHFGMNNKWKK | 80 | 2002.01 |
| 03-1 | 16 - 18 | G(Pip)(Cit)AV(Cit)T(Dab)(Nap)LDKWKK | 97 | 2006.13 |
| 03-1 | 16 - 18 | GQEAFTHQEMHKWKK | 81 | 2007.98 |
| 03-1 | 16 - 18 | GPQQQ(Cit)(Cit)(Cit)TGKKWKK | 85 | 2008.12 |
| 03-1 | 16 - 18 | G(Pal)R(Pip)G(Cit)TNFD(Pip)KWKK | 74 | 2017.09 |
| 03-1 | 16 - 18 | G(Pip)FLT(Cit)TNFD(Pip)KWKK | 98 | 2017.10 |
| 03-1 | 16 - 18 | GPQE(Cit)RAFEPHKWKK | 90 | 2018.07 |
| 03-1 | 16 - 18 | G(Cit)RVRNG(Iph)SG(Dab)KWKK | 72 | 2025.97 |
| 03-1 | 16 - 18 | GEKT(Iph)EQ(Dab)TNGKWKK | 98 | 2029.90 |
| 03-1 | 16 - 18 | GSLTERPS(Iph)E(Dab)KWKK | 77 | 2041.93 |
| 03-1 | 16 - 18 | GPND(Dab)LDQ(Iph)P(Pip)KWKK | 96 | 2047.92 |

| | | | | |
|---|---|---|---|---|
| 03-1 | 16 - 18 | GTRPTQ(Iph)SVD(Pip)KWKK | 87 | 2052.95 |
| 03-1 | 16 - 18 | GE(Nap)SET(Nap)PKSMKWKK | 72 | 2053.02 |
| 03-1 | 16 - 18 | GDKD(Nap)(Cit)S(Pip)RNDKWKK | 76 | 2080.07 |
| 03-1 | 16 - 18 | G(Cit)(Cit)RSDKNQ(Cit)LKWKK | 87 | 2082.17 |
| 03-1 | 16 - 18 | GRDSNLL(Iph)E(Pal)NKWKK | 70 | 2131.96 |
| 03-1 | 16 - 18 | G(Dab)DQEQT(Cit)(Cit)(Dab)(Iph)KWKK | 86 | 2157.97 |
| 03-1 | 16 - 18 | GHAT(Iph)TDR(Cit)(Cit)QKWKK | 82 | 2165.98 |
| 04-1 | 19 - 21 | GPD(Iph)N(Dab)LRGGNKWKK | 98 | 1965.89 |
| 04-1 | 19 - 21 | GP(Pip)(Cit)LEA(Cit)RLNKWKK | 98 | 2003.17 |
| 04-1 | 19 - 21 | GFVDA(Pip)GGLGHKWKK | 97 | 1748.96 |
| 04-1 | 19 - 21 | G(Nap)TKMDSFQHTKWKK | 97 | 2042.03 |
| 04-1 | 19 - 21 | GL(Iph)PTGGKLPGKWKK | 96 | 1862.92 |
| 04-1 | 19 - 21 | GMTEHV(Cit)S(Dab)LNKWKK | 96 | 1938.04 |
| 04-1 | 19 - 21 | GDFQVEPLHGHKWKK | 95 | 1929.01 |
| 04-1 | 19 - 21 | GGHEHVDH(Iph)NSKWKK | 95 | 2054.85 |
| 04-1 | 19 - 21 | GEAVMKA(Dab)PATKWKK | 94 | 1767.99 |
| 04-1 | 19 - 21 | GVTDLGH(Iph)HPVKWKK | 93 | 1997.92 |
| 04-1 | 19 - 21 | GND(Dab)SL(Nap)NRN(Cit)KWKK | 93 | 2037.08 |
| 04-1 | 19 - 21 | GTERN(Nap)FTSE(Pip)KWKK | 93 | 2057.06 |
| 04-1 | 19 - 21 | GLHQ(Nap)NS(Iph)NMPKWKK | 93 | 2160.93 |
| 04-1 | 19 - 21 | GG(Pip)QMEAGKVPKWKK | 92 | 1792.99 |
| 04-1 | 19 - 21 | G(Cit)T(Cit)(Cit)DLRKSLKWKK | 92 | 2054.20 |
| 04-1 | 19 - 21 | GNSGGGH(Dab)V(Nap)EKWKK | 91 | 1803.93 |
| 04-1 | 19 - 21 | GALMEK(Dab)(Cit)TPNKWKK | 90 | 1911.06 |
| 04-1 | 19 - 21 | GSQGRS(Nap)HQGEKWKK | 90 | 1932.98 |
| 04-1 | 19 - 21 | GL(Cit)NERSP(Iph)E(Dab)KWKK | 89 | 2124.98 |
| 04-1 | 19 - 21 | GEGQ(Dab)TNHTMAKWKK | 88 | 1838.93 |
| 04-1 | 19 - 21 | G(Iph)GS(Pip)GT(Dab)EPMKWKK | 88 | 1927.84 |
| 04-1 | 19 - 21 | GA(Cit)H(Iph)PFDSP(Dab)KWKK | 87 | 2050.91 |
| 04-1 | 19 - 21 | GNGAT(Nap)LLP(Dab)VKWKK | 85 | 1832.06 |
| 04-1 | 19 - 21 | G(Pip)EPLGKFQ(Pip)(Iph)KWKK | 85 | 2094.02 |
| 04-1 | 19 - 21 | GLHQ(Nap)NS(Iph)NDEKWKK | 85 | 2176.91 |
| 04-1 | 19 - 21 | GAFPKVR(Iph)VDVKWKK | 84 | 2054.02 |
| 04-1 | 19 - 21 | GNPHVQEL(Cit)VKKWKK | 83 | 1971.13 |
| 04-1 | 19 - 21 | GLVEPTMEQRKKWKK | 83 | 1981.10 |
| 04-1 | 19 - 21 | G(Iph)DLEHPV(Dab)TLKWKK | 83 | 2046.97 |
| 04-1 | 19 - 21 | GNQEHPVMNGKKWKK | 82 | 1904.00 |
| 04-1 | 19 - 21 | GV(Nap)(Cit)TKLF(Cit)SPKWKK | 82 | 2053.17 |
| 04-1 | 19 - 21 | G(Cit)FVEHFGMNNKWKK | 80 | 2002.01 |

| | | | | |
|---|---|---|---|---|
| 04-1 | 19 - 21 | GAFV(Pip)VR(Iph)VDVKWKK | 80 | 2054.02 |
| 04-1 | 19 - 21 | GAFPVKR(Iph)VDVKWKK | 79 | 2054.02 |
| 04-1 | 19 - 21 | GAFD(Cit)S(Pip)QAK(Iph)KWKK | 79 | 2072.96 |
| 04-1 | 19 - 21 | G(Cit)HSN(Nap)NE(Cit)N(Dab)KWKK | 79 | 2076.05 |
| 04-1 | 19 - 21 | G(Cit)(Dab)PQM(Dab)VQPLKWKK | 78 | 1920.10 |
| 04-1 | 19 - 21 | GAS(Cit)VDF(Dab)(Dab)(Cit)(Nap)KWKK | 78 | 2000.09 |
| 04-1 | 19 - 21 | GEAFLNHH(Cit)QLKWKK | 78 | 2016.09 |
| 04-1 | 19 - 21 | GN(Iph)VSPMQ(Pip)GGKWKK | 77 | 1938.85 |
| 04-1 | 19 - 21 | GNFT(Cit)DHP(Cit)KAKWKK | 77 | 1994.07 |
| 04-1 | 19 - 21 | G(Iph)VSVPQR(Cit)QSKWKK | 77 | 2080.99 |
| 04-1 | 19 - 21 | GADKK(Dab)R(Iph)VDVKWKK | 76 | 2054.02 |
| 04-1 | 19 - 21 | GMTEHVFP(Dab)LNKWKK | 75 | 1938.04 |
| 04-1 | 19 - 21 | GAFV(Pip)VR(Iph)VDVKWKK | 75 | 2054.02 |
| 04-1 | 19 - 21 | GMTTSPG(Cit)GPRKWKK | 74 | 1810.97 |
| 04-1 | 19 - 21 | GLVETPMEQ(Dab)GKWKK | 74 | 1853.99 |
| 04-1 | 19 - 21 | GMLANDHMKGMKWKK | 74 | 1897.96 |
| 04-1 | 19 - 21 | GTSMT(Dab)FFTNKKWKK | 74 | 1927.03 |
| 04-1 | 19 - 21 | GMDSL(Nap)NRNG(Dab)KWKK | 74 | 1954.01 |
| 04-1 | 19 - 21 | G(Dab)GAD(Cit)(Iph)HHGLKWKK | 73 | 1986.89 |
| 04-1 | 19 - 21 | GLRDG(Cit)GS(Dab)GKKWKK | 72 | 1797.02 |
| 04-1 | 19 - 21 | G(Iph)LSGPKGAAGKWKK | 70 | 1780.84 |
| 04-1 | 19 - 21 | GE(Iph)GG(Dab)LRGGNKWKK | 70 | 1882.86 |
| 04-1 | 19 - 21 | GNFSATFFTNKKWKK | 70 | 1927.02 |
| 04-1 | 19 - 21 | GANP(Iph)PFDPS(Dab)KWKK | 70 | 1967.87 |
| 05-1 | 22 - 24 | GLA(Pip)(Cit)VTTPLKKWKK | 98 | 1876.15 |
| 05-1 | 22 - 24 | GVVLPAAPP(Pal)GKWKK | 94 | 1719.01 |
| 05-1 | 22 - 24 | GVRL(Dab)FVQPL(Cit)KWKK | 92 | 1979.21 |
| 05-1 | 22 - 24 | GPTNLQHNKN(Iph)KWKK | 92 | 2088.96 |
| 05-1 | 22 - 24 | G(Iph)GTHFE(Dab)LNGKWKK | 91 | 1997.89 |
| 05-1 | 22 - 24 | GN(Cit)EGP(Cit)(Dab)LKFKWKK | 89 | 1969.11 |
| 05-1 | 22 - 24 | GSP(Nap)LSFTPKPKWKK | 88 | 1921.07 |
| 05-1 | 22 - 24 | GKQHLSLLDT(Dab)KWKK | 87 | 1905.11 |
| 05-1 | 22 - 24 | GLLSKPLRT(Cit)FKWKK | 86 | 1982.21 |
| 05-1 | 22 - 24 | G(Cit)AQA(Cit)SKVV(Iph)KWKK | 85 | 2040.00 |
| 05-1 | 22 - 24 | GS(Pip)AVPNLAKLKWKK | 82 | 1789.08 |
| 05-1 | 22 - 24 | GQP(Nap)GA(Dab)PAERKWKK | 81 | 1873.02 |
| 05-1 | 22 - 24 | GVVSSKKAFG(Pal)KWKK | 80 | 1821.05 |
| 05-1 | 22 - 24 | GVEMSDQP(Iph)(Pip)RKWKK | 78 | 2110.94 |
| 05-1 | 22 - 24 | GPADQQSGQRKKWKK | 77 | 1865.01 |

| | | | | |
|---|---|---|---|---|
| 05-1 | 22 - 24 | GVVSRVAARNTKWKK | 76 | 1823.08 |
| 05-1 | 22 - 24 | GTG(Nap)HHPATFGKWKK | 76 | 1871.97 |
| 05-1 | 22 - 24 | GL(Pal)LPLAAS(Pip)NKWKK | 74 | 1823.07 |
| 05-1 | 22 - 24 | GGLASKKASHQKWKK | 73 | 1777.02 |
| 05-1 | 22 - 24 | G(Pip)GKVVGPPT(Pal)KWKK | 73 | 1779.04 |
| 05-1 | 22 - 24 | G(Pip)ALNMNL(Dab)MLKWKK | 72 | 1896.07 |
| 05-1 | 22 - 24 | G(Nap)FT(Cit)DHP(Cit)KAKWKK | 71 | 2077.11 |
| 05-1 | 22 - 24 | GALR(Pip)VFPPG(Pal)KWKK | 68 | 1881.10 |
| 06-1 | 25 - 28 | G(Nap)M(Dab)MGSATKAKWKK | 98 | 1843.97 |
| 06-1 | 25 - 28 | GRFSGRMLTSSKWKK | 98 | 1892.03 |
| 06-1 | 25 - 28 | GNNRSTS(Cit)GHFKWKK | 98 | 1927.00 |
| 06-1 | 25 - 28 | GLLLRNHNHDGKWKK | 98 | 1939.08 |
| 06-1 | 25 - 28 | GPS(Dab)FDFRNNKKWKK | 98 | 1975.07 |
| 06-1 | 25 - 28 | GKVNHELQF(Dab)NKWKK | 98 | 1979.10 |
| 06-1 | 25 - 28 | GKFNR(Nap)HTDTVKWKK | 98 | 2065.11 |
| 06-1 | 25 - 28 | GK(Iph)KTGEREFNKWKK | 98 | 2131.99 |
| 06-1 | 25 - 28 | GKAAFGMR(Cit)AMKWKK | 97 | 1890.03 |
| 06-1 | 25 - 28 | G(Cit)GARFS(Cit)HVGKWKK | 97 | 1895.05 |
| 06-1 | 25 - 28 | GRSGNSPTS(Dab)(Iph)KWKK | 97 | 1928.86 |
| 06-1 | 25 - 28 | GHSMETH(Cit)GTHKWKK | 97 | 1943.96 |
| 06-1 | 25 - 28 | GT(Nap)SHGNA(Cit)(Cit)HKWKK | 97 | 1985.02 |
| 06-1 | 25 - 28 | GMHS(Iph)LNSHNGKWKK | 97 | 2019.85 |
| 06-1 | 25 - 28 | G(Iph)RSND(Cit)N(Iph)FKKWKK | 97 | 2333.90 |
| 06-1 | 25 - 28 | GR(Cit)MGAKNGAAKWKK | 96 | 1782.99 |
| 06-1 | 25 - 28 | GLGRGQSNVARKWKK | 96 | 1808.04 |
| 06-1 | 25 - 28 | GHNNG(Iph)Q(Cit)E(Iph)RKWKK | 96 | 2307.85 |
| 06-1 | 25 - 28 | GLMNVNKG(Dab)G(Cit)KWKK | 95 | 1840.04 |
| 06-1 | 25 - 28 | G(Dab)EAQHGQL(Cit)HKWKK | 95 | 1927.04 |
| 06-1 | 25 - 28 | GKS(Iph)GAQRTPSKWKK | 95 | 1954.91 |
| 06-1 | 25 - 28 | GPLLHGTGVR(Cit)KWKK | 94 | 1857.10 |
| 06-1 | 25 - 28 | G(Cit)LSKATHKENKWKK | 94 | 1935.09 |
| 06-1 | 25 - 28 | GLLSKLPRT(Cit)FKWKK | 92 | 1982.21 |
| 06-1 | 25 - 28 | GSL(Iph)S(Dab)MKTP(Cit)KWKK | 92 | 2043.97 |
| 06-1 | 25 - 28 | G(Nap)KDSNQK(Dab)(Cit)QKWKK | 92 | 2052.11 |
| 06-1 | 25 - 28 | G(Iph)(Iph)ETQQHA(Dab)EKWKK | 92 | 2238.81 |
| 06-1 | 25 - 28 | GSRSHP(Nap)AGMNKWKK | 91 | 1903.97 |
| 06-1 | 25 - 28 | G(Pip)(Nap)AG(Cit)GNVH(Cit)KWKK | 91 | 1942.05 |
| 06-1 | 25 - 28 | GQKSKDQQRTVKWKK | 91 | 1968.11 |
| 06-1 | 25 - 28 | GPE(Pip)P(Nap)FQRTGKWKK | 91 | 2005.08 |

| | | | | |
|---|---|---|---|---|
| 06-1 | 25 - 28 | GFVSE(Cit)(Cit)(Iph)RRAKWKK | 91 | 2202.06 |
| 06-1 | 25 - 28 | G(Pip)LKFNS(Cit)GGGKWKK | 90 | 1813.02 |
| 06-1 | 25 - 28 | GKDTV(Dab)TQRN(Cit)KWKK | 90 | 1969.11 |
| 06-1 | 25 - 28 | GH(Nap)(Cit)T(Nap)PGNTKKWKK | 89 | 2056.09 |
| 06-1 | 25 - 28 | GGEKLGRE(Dab)AGKWKK | 88 | 1767.00 |
| 06-1 | 25 - 28 | GKGSLLM(Pal)AQ(Dab)KWKK | 88 | 1846.05 |
| 06-1 | 25 - 28 | GN(Iph)RGGLGKG(Dab)KWKK | 88 | 1881.91 |
| 06-1 | 25 - 28 | GVHPLNGLKKEKWKK | 88 | 1885.12 |
| 06-1 | 25 - 28 | GVATQSRAVHFKWKK | 87 | 1866.05 |
| 06-1 | 25 - 28 | GK(Pip)NG(Cit)(Nap)F(Cit)PVKWKK | 87 | 2049.15 |
| 06-1 | 25 - 28 | GTRP(Cit)DE(Iph)RTRKWKK | 87 | 2211.04 |
| 06-1 | 25 - 28 | GNM(Dab)MGSATKAKWKK | 86 | 1760.93 |
| 06-1 | 25 - 28 | GMLSGQ(Dab)FV(Pip)FKWKK | 86 | 1905.06 |
| 06-1 | 25 - 28 | GGFMQ(Iph)KGVHTKWKK | 86 | 2027.92 |
| 06-1 | 25 - 28 | GHTG(Cit)GGGVH(Cit)KWKK | 85 | 1785.96 |
| 06-1 | 25 - 28 | GKKRGGHNHDGKWKK | 85 | 1856.01 |
| 06-1 | 25 - 28 | GALAPGH(Iph)VM(Dab)KWKK | 85 | 1918.90 |
| 06-1 | 25 - 28 | G(Iph)KEQDAKHSVKWKK | 85 | 2064.95 |
| 06-1 | 25 - 28 | GFA(Cit)DNGPGR(Dab)KWKK | 84 | 1840.99 |
| 06-1 | 25 - 28 | GT(Iph)G(Dab)PARPFSKWKK | 84 | 1955.91 |
| 06-1 | 25 - 28 | GSRL(Cit)GSSH(Pal)(Cit)KWKK | 83 | 1956.07 |
| 06-1 | 25 - 28 | GTF(Dab)MSF(Cit)GAHKWKK | 82 | 1904.99 |
| 06-1 | 25 - 28 | GNDK(Dab)(Nap)T(Dab)NNQKWKK | 82 | 1981.04 |
| 06-1 | 25 - 28 | GPGPRPERFF(Nap)KWKK | 82 | 2050.12 |
| 06-1 | 25 - 28 | GVH(Iph)(Cit)KSSN(Iph)(Cit)KWKK | 82 | 2281.90 |
| 06-1 | 25 - 28 | GP(Dab)GAMAQLGKKWKK | 81 | 1722.98 |
| 06-1 | 25 - 28 | GKNFGDSQF(Dab)(Dab)KWKK | 81 | 1893.01 |
| 06-1 | 25 - 28 | GDQAQQTSF(Dab)GKWKK | 80 | 1831.94 |
| 06-1 | 25 - 28 | GPATLQAGH(Nap)AKWKK | 78 | 1812.99 |
| 06-1 | 25 - 28 | GNPFGLVKRMGKWKK | 78 | 1869.07 |
| 06-1 | 25 - 28 | GQF(Nap)SFPTQRSKWKK | 76 | 2045.07 |
| 06-1 | 25 - 28 | GARRSTGGLGTKWKK | 75 | 1725.99 |
| 06-1 | 25 - 28 | GSKTSGVAVHFKWKK | 75 | 1783.00 |
| 06-1 | 25 - 28 | GNNRSTSDVHFKWKK | 75 | 1926.99 |
| 06-1 | 25 - 28 | GQF(Nap)S(Cit)STQRSKWKK | 75 | 2045.07 |
| 06-1 | 25 - 28 | G(Dab)(Dab)(Iph)S(Dab)MKTP(Cit)KWKK | 74 | 2043.98 |
| 06-1 | 25 - 28 | GTKG(Dab)LT(Nap)RAQKWKK | 73 | 1922.11 |
| 06-1 | 25 - 28 | GK(Dab)T(Dab)DHNHPGKWKK | 71 | 1856.00 |
| 06-1 | 25 - 28 | GRSDTAGHKSLKWKK | 70 | 1822.01 |

| | | | | |
|---|---|---|---|---|
| 06-1 | 25 - 28 | G(Cit)TNDRE(Dab)RA(Cit)KWKK | 70 | 2026.10 |
| 06-1 | 25 - 28 | GLGG(Cit)(Pip)AGH(Nap)AKWKK | 68 | 1813.00 |
| 06-1 | 25 - 28 | GRSD(Dab)LMSKFPKWKK | 68 | 1931.07 |
| 06-1 | 25 - 28 | G(Pal)(Cit)LQVS(Iph)(Dab)GGKWKK | 68 | 1988.93 |
| 06-1 | 25 - 28 | G(Iph)HK(Iph)SSMGSPKWKK | 68 | 2126.77 |
| 06-1 | 25 - 28 | GFTGAG(Dab)FKPFKWKK | 67 | 1822.02 |
| 06-1 | 25 - 28 | GFAG(Dab)DNGL(Nap)(Dab)KWKK | 67 | 1840.98 |
| 06-1 | 25 - 28 | GG(Pal)G(Dab)LFTNKFKWKK | 67 | 1882.05 |
| 06-1 | 25 - 28 | G(Iph)A(Nap)(Dab)(Dab)LMGVEKWKK | 67 | 2039.94 |
| 06-1 | 25 - 28 | G(Dab)E(Cit)TPNH(Iph)ERKWKK | 67 | 2162.97 |
| 07-1 | 29 - 32 | G(Dab)ATK(Cit)(Cit)AHLGKWKK | 99 | 1862.09 |
| 07-1 | 29 - 32 | GSKSVNLH(Dab)MSKWKK | 98 | 1853.02 |
| 07-1 | 29 - 32 | GKKK(Iph)NEE(Nap)GVKWKK | 98 | 2152.02 |
| 07-1 | 29 - 32 | G(Iph)SNGHSTF(Pip)(Iph)KWKK | 98 | 2171.78 |
| 07-1 | 29 - 32 | GNKEFG(Pip)SQRLKWKK | 97 | 1955.10 |
| 07-1 | 29 - 32 | GS(Dab)A(Iph)LKRETLKWKK | 97 | 2041.02 |
| 07-1 | 29 - 32 | GLTLNND(Dab)T(Dab)RKWKK | 96 | 1897.08 |
| 07-1 | 29 - 32 | GLFNFSG(Pip)FG(Dab)KWKK | 93 | 1865.02 |
| 07-1 | 29 - 32 | GATSAMTKREHKWKK | 93 | 1882.01 |
| 07-1 | 29 - 32 | GQ(Nap)(Cit)(Dab)DSNS(Dab)RKWKK | 93 | 2011.06 |
| 07-1 | 29 - 32 | GFEQ(Dab)KF(Iph)GQKKWKK | 93 | 2135.01 |
| 07-1 | 29 - 32 | GPKLDNMNRRLKWKK | 92 | 2007.14 |
| 07-1 | 29 - 32 | G(Iph)HNATNTFLRKWKK | 92 | 2096.97 |
| 07-1 | 29 - 32 | GQKAKLAPGKPKWKK | 91 | 1788.10 |
| 07-1 | 29 - 32 | G(Cit)AHEGTH(Cit)F(Dab)KWKK | 91 | 1963.04 |
| 07-1 | 29 - 32 | G(Dab)SSK(Iph)D(Dab)TREKWKK | 90 | 2045.94 |
| 07-1 | 29 - 32 | G(Pip)QSAASPLR(Dab)KWKK | 89 | 1806.05 |
| 07-1 | 29 - 32 | GTGFAPKV(Iph)N(Dab)KWKK | 89 | 1956.93 |
| 07-1 | 29 - 32 | GG(Pip)RTP(Nap)TPT(Nap)KWKK | 89 | 2000.09 |
| 07-1 | 29 - 32 | GNGSD(Nap)(Pip)GRM(Dab)KWKK | 88 | 1909.98 |
| 07-1 | 29 - 32 | G(Iph)F(Pip)PVDHVAHKWKK | 88 | 2070.96 |
| 07-1 | 29 - 32 | G(Dab)RAT(Cit)(Iph)GQEHKWKK | 88 | 2078.95 |
| 07-1 | 29 - 32 | GGSSMGGSRVRKWKK | 86 | 1743.94 |
| 07-1 | 29 - 32 | GKKK(Iph)NEEQPQKWKK | 86 | 2152.02 |
| 07-1 | 29 - 32 | GPSTPQGA(Cit)HRKWKK | 85 | 1858.02 |
| 07-1 | 29 - 32 | GKR(Iph)VNSQEH(Cit)KWKK | 84 | 2178.02 |
| 07-1 | 29 - 32 | GF(Pal)(Dab)(Dab)HF(Iph)GQKKWKK | 82 | 2135.00 |
| 07-1 | 29 - 32 | G(Cit)QAFNSLGR(Dab)KWKK | 80 | 1900.07 |
| 07-1 | 29 - 32 | GSAGEKH(Cit)(Iph)QHKWKK | 78 | 2073.93 |

| | | | | |
|---|---|---|---|---|
| 07-1 | 29 - 32 | GE(Iph)HG(Dab)HA(Cit)N(Cit)KWKK | 78 | 2101.93 |
| 07-1 | 29 - 32 | G(Pip)(Nap)FQQTSF(Dab)GKWKK | 77 | 1988.05 |
| 07-1 | 29 - 32 | G(Pip)VGH(Pip)QE(Dab)F(Iph)KWKK | 77 | 2091.98 |
| 07-1 | 29 - 32 | GPE(Pip)DPNRP(Iph)(Pip)KWKK | 77 | 2099.97 |
| 07-1 | 29 - 32 | GTGRHGMSEPRKWKK | 76 | 1877.99 |
| 07-1 | 29 - 32 | GG(Dab)(Cit)HGMSEPRKWKK | 74 | 1877.99 |
| 07-1 | 29 - 32 | GVV(Nap)KREV(Dab)VQKWKK | 72 | 2004.19 |
| 07-1 | 29 - 32 | GSPHAFDS(Iph)HRKWKK | 72 | 2076.90 |
| 07-1 | 29 - 32 | GGG(Iph)RGGLGK(Cit)KWKK | 71 | 1881.91 |
| 07-1 | 29 - 32 | GS(Cit)V(Cit)V(Iph)(Cit)QKRKWKK | 71 | 2211.12 |
| 07-1 | 29 - 32 | GR(Dab)AKLAGPKPKWKK | 70 | 1788.11 |
| 07-1 | 29 - 32 | GPSP(Dab)EGA(Cit)HRKWKK | 70 | 1858.02 |
| 07-1 | 29 - 32 | GQDKAG(Cit)THRFKWKK | 69 | 1967.07 |
| 07-1 | 29 - 32 | GLTQVND(Dab)T(Dab)RKWKK | 66 | 1897.08 |
| 07-1 | 29 - 32 | GHAKEVQKEKLKWKK | 65 | 1960.15 |
| 08-1 | 33 - 37 | GRNVPASAGH(Dab)KWKK | 97 | 1758.99 |
| 08-1 | 33 - 37 | GTTTENVHP(Dab)(Dab)KWKK | 94 | 1849.01 |
| 08-1 | 33 - 37 | GP(Cit)(Dab)Q(Dab)THPANKWKK | 94 | 1872.03 |
| 08-1 | 33 - 37 | GNTHF(Dab)TRSQ(Cit)KWKK | 94 | 1998.08 |
| 08-1 | 33 - 37 | GP(Cit)(Dab)Q(Dab)THAPNKWKK | 91 | 1872.03 |
| 08-1 | 33 - 37 | GGHSGG(Cit)HP(Cit)HKWKK | 85 | 1849.97 |
| 08-1 | 33 - 37 | GPN(Cit)(Dab)AGGRERKWKK | 83 | 1864.04 |
| 08-1 | 33 - 37 | GVVSSKKAFG(Pal)KWKK | 78 | 1821.05 |
| 08-1 | 33 - 37 | G(Cit)KLS(Cit)VDN(Dab)AKWKK | 78 | 1911.09 |
| 08-1 | 33 - 37 | GGR(Pip)DSSGGRKKWKK | 73 | 1796.00 |
| 08-1 | 33 - 37 | GVVARAAA(Pal)(Pip)PKWKK | 72 | 1779.05 |
| 08-1 | 33 - 37 | GSKVVGTQRS(Pip)KWKK | 72 | 1838.08 |
| 08-1 | 33 - 37 | G(Cit)SHMQSQHKSKWKK | 72 | 1977.02 |
| 08-1 | 33 - 37 | GKAPRAAPLT(Pal)KWKK | 71 | 1823.08 |
| 08-1 | 33 - 37 | GVVSVRAAVMPKWKK | 70 | 1779.05 |
| 08-1 | 33 - 37 | GVVFLPVALADKWKK | 68 | 1794.07 |
| 08-1 | 33 - 37 | GAKPRAAPLT(Pal)KWKK | 67 | 1823.08 |
| 08-1 | 33 - 37 | G(Cit)RLV(Cit)SRV(Pal)TKWKK | 67 | 2043.21 |
| 08-1 | 33 - 37 | GPRNSSSGGRKKWKK | 66 | 1796.00 |
| 08-1 | 33 - 37 | G(Pip)A(Dab)KPASR(Dab)DKWKK | 65 | 1821.06 |
| 09-1 | 38 - 45 | G(Dab)KSMVTF(Dab)HVKWKK | 98 | 1899.08 |

| | | | | | |
|---|---|---|---|---|---|
| 09-1 | 38 - 45 | GRSGHK(Nap)(Dab)(Cit)ELKWKK | 97 | 2031.14 | |
| 09-1 | 38 - 45 | GNEHHT(Nap)RFKNKWKK | 97 | 2130.11 | |
| 09-1 | 38 - 45 | G(Nap)HTQRE(Dab)LTKKWKK | 97 | 2060.15 | |
| 09-1 | 38 - 45 | GVKMRNEFLH(Dab)KWKK | 97 | 2024.14 | |
| 09-1 | 38 - 45 | GPRF(Iph)NKSLRAKWKK | 97 | 2112.05 | |
| 09-1 | 38 - 45 | G(Dab)TQHLTS(Dab)LKKWKK | 96 | 1878.11 | |
| 09-1 | 38 - 45 | GHQHV(Dab)(Iph)TQKNKWKK | 96 | 2114.99 | |
| 09-1 | 38 - 45 | GFQPDH(Dab)TT(Dab)(Dab)KWKK | 95 | 1896.02 | |
| 09-1 | 38 - 45 | GKPLFKHVR(Cit)SKWKK | 95 | 2019.21 | |
| 09-1 | 38 - 45 | GA(Nap)AKVTTFKRKWKK | 95 | 1969.15 | |
| 09-1 | 38 - 45 | GPKHQTKP(Iph)HNKWKK | 94 | 2110.00 | |
| 09-1 | 38 - 45 | G(Pip)DKTQ(Iph)VR(Dab)QKWKK | 94 | 2124.04 | |
| 09-1 | 38 - 45 | G(Pal)N(Iph)(Dab)GVRKNNKWKK | 94 | 2072.98 | |
| 09-1 | 38 - 45 | G(Dab)(Dab)HNFK(Nap)STQKWKK | 94 | 2009.09 | |
| 09-1 | 38 - 45 | GRTGS(Iph)P(Pip)(Dab)VFKWKK | 94 | 2012.97 | |
| 09-1 | 38 - 45 | G(Dab)(Dab)HLVGLQARKWKK | 93 | 1844.11 | |
| 09-1 | 38 - 45 | GKARS(Cit)MLKKTKWKK | 92 | 1970.18 | |
| 09-1 | 38 - 45 | GAHVE(Dab)VP(Pip)KPKWKK | 92 | 1853.09 | |
| **09-1** | **38-45** | **G(Cit)RSKDVTKHHKWKK** | **92** | **2015.14** | **CXP0** |
| 09-1 | 38 - 45 | GPRKMQPQ(Dab)AHKWKK | 91 | 1943.09 | |
| 09-1 | 38 - 45 | G(Dab)ETRMQ(Iph)KKNKWKK | 91 | 2158.02 | |
| 09-1 | 38 - 45 | GG(Nap)SH(Pip)HKSPVKWKK | 91 | 1922.05 | |
| 09-1 | 38 - 45 | GPTDRNK(Dab)K(Nap)(Iph)KWKK | 91 | 2179.05 | |
| 09-1 | 38 - 45 | G(Dab)MNDLVRAHRKWKK | 90 | 1962.10 | |
| 09-1 | 38 - 45 | GK(Iph)FKTERHVMKWKK | 90 | 2199.05 | |
| 09-1 | 38 - 45 | G(Pip)RELP(Dab)(Pip)NATKWKK | 89 | 1903.10 | |
| 09-1 | 38 - 45 | G(Dab)GFPEARFH(Dab)KWKK | 88 | 1911.05 | |
| 09-1 | 38 - 45 | GHGSNAPA(Nap)RRKWKK | 87 | 1913.04 | |
| 09-1 | 38 - 45 | GMRH(Nap)RTQ(Cit)KDKWKK | 87 | 2176.17 | |
| 09-1 | 38 - 45 | GKTGS(Iph)P(Pip)(Dab)VFKWKK | 87 | 1984.96 | |
| 09-1 | 38 - 45 | GS(Dab)QM(Iph)HL(Dab)RDKWKK | 86 | 2109.97 | |

| | | | | |
|---|---|---|---|---|
| 09-1 | 38 - 45 | G(Dab)(Dab)A(Iph)T(Dab)(Iph)QHDKWKK | 86 | 2167.82 |
| 09-1 | 38 - 45 | GGREGPK(Pip)HVTKWKK | 85 | 1857.06 |
| 09-1 | 38 - 45 | GT(Dab)ANRP(Cit)HKMKWKK | 84 | 1962.10 |
| 09-1 | 38 - 45 | GVQHPKANHLRKWKK | 84 | 1950.13 |
| 09-1 | 38 - 45 | GAV(Dab)A(Pip)GNR(Iph)(Nap)KWKK | 83 | 2033.97 |
| 09-1 | 38 - 45 | GGFLGFP(Dab)HKFKWKK | 83 | 1900.07 |
| 09-1 | 38 - 45 | GT(Dab)ANRQ(Pip)HKMKWKK | 82 | 1962.10 |
| 09-1 | 38 - 45 | GVVKRH(Iph)A(Nap)DKKWKK | 82 | 2173.07 |
| 09-1 | 38 - 45 | GTN(Iph)(Cit)KAALRKKWKK | 81 | 2082.06 |
| 09-1 | 38 - 45 | GSAN(Dab)(Iph)RKGA(Nap)KWKK | 81 | 2023.95 |
| 09-1 | 38 - 45 | G(Dab)(Iph)A(Pip)TQ(Iph)PRSKWKK | 79 | 2181.87 |
| 09-1 | 38 - 45 | GHGQQE(Pip)LVS(Pip)KWKK | 78 | 1900.05 |
| 09-1 | 38 - 45 | GH(Pip)TD(Cit)(Dab)(Dab)RFEKWKK | 77 | 2038.11 |
| 09-1 | 38 - 45 | G(Iph)REEG(Dab)VRK(Dab)KWKK | 77 | 2097.04 |
| 09-1 | 38 - 45 | GTHGV(Iph)LG(Cit)RHKWKK | 77 | 2056.98 |
| 09-1 | 38 - 45 | GHG(Nap)ST(Pip)LVS(Pip)KWKK | 77 | 1900.06 |
| 09-1 | 38 - 45 | GHS(Iph)M(Pip)Q(Cit)FRAKWKK | 77 | 2183.00 |
| 09-1 | 38 - 45 | G(Nap)L(Pip)(Dab)F(Dab)NNGGKWKK | 77 | 1895.04 |
| 09-1 | 38 - 45 | G(Dab)KSM(Dab)(Dab)F(Dab)HVKWKK | 76 | 1899.09 |
| 09-1 | 38 - 45 | GRTG(Cit)LG(Nap)(Dab)(Cit)RKWKK | 76 | 2021.17 |
| 09-1 | 38 - 45 | G(Pal)STH(Pip)SLVGSKWKK | 75 | 1811.99 |
| 09-1 | 38 - 45 | G(Dab)HKHTPMP(Cit)NKWKK | 74 | 1969.07 |
| 09-1 | 38 - 45 | GTVHNFK(Nap)STQKWKK | 74 | 2009.07 |
| 09-1 | 38 - 45 | G(Iph)(Dab)RSLLKHMDKWKK | 74 | 2123.02 |
| 10-1 | 46 - 60 | GHL(Iph)(Dab)GAGHTRKWKK | 98 | 1971.93 |
| 10-1 | 46 - 60 | GHA(Iph)NRSRFKAKWKK | 98 | 2110.01 |
| 10-1 | 46 - 60 | G(Pip)RD(Iph)NFKRGGKWKK | 97 | 2098.99 |
| 10-1 | 46 - 60 | GHSGFMVG(Dab)RRKWKK | 97 | 1897.05 |
| 10-1 | 46 - 60 | GDEHL(Cit)HKR(Iph)KKWKK | 97 | 2243.08 |
| 10-1 | 46 - 60 | GRVSTRAQ(Cit)K(Dab)KWKK | 96 | 1953.16 |
| 10-1 | 46 - 60 | GV(Cit)FNFK(Cit)(Dab)(Dab)KKWKK | 96 | 2047.21 |

| | | | | | |
|---|---|---|---|---|---|
| **10-1** | **46 - 60** | **GNFKQ(Nap)HAG(Dab)RKWKK** | **96** | **2005.10** | **CXP2** |
| 10-1 | 46 - 60 | GPQRTRKLVM(Pip)KWKK | 95 | 2005.20 | |
| 10-1 | 46 - 60 | G(Pip)FALHVRV(Iph)KKWKK | 95 | 2119.10 | |
| 10-1 | 46 - 60 | GSH(Cit)HNMRKKPKWKK | 94 | 2042.13 | |
| **10-1** | **46 - 60** | **G(Dab)PR(Dab)LHS(Dab)ATKWKK** | **94** | **1832.08** | **CXP5** |
| 10-1 | 46 - 60 | GVEFKSH(Cit)RRRKWKK | 94 | 2122.22 | |
| **10-1** | **46 - 60** | **G(Cit)MFG(Pip)(Dab)(Dab)KAPKWKK** | **94** | **1884.08** | **CXP4** |
| 10-1 | 46 - 60 | GALSRAA(Dab)HR(Nap)KWKK | 94 | 1929.11 | |
| **10-1** | **46 - 60** | **GKQKTS(Iph)GRG(Pip)KWKK** | **93** | **2010.99** | **CXP1** |
| **10-1** | **46 - 60** | **GVQH(Iph)R(Dab)KEKNKWKK** | **92** | **2162.06** | **CXP7** |
| 10-1 | 46 - 60 | GAGRAKA(Iph)KRTKWKK | 92 | 1982.01 | |
| 10-1 | 46 - 60 | GTFKQSGH(Dab)(Iph)(Pip)KWKK | 92 | 2053.96 | |
| 10-1 | 46 - 60 | GKQR(Dab)(Dab)E(Iph)(Iph)FFKWKK | 92 | 2350.96 | |
| 10-1 | 46 - 60 | GKRPGSF(Cit)RR(Iph)KWKK | 91 | 2184.09 | |
| 10-1 | 46 - 60 | GPPS(Iph)R(Dab)KEKNKWKK | 90 | 2079.01 | |
| 10-1 | 46 - 60 | GNRQAL(Dab)RRNTKWKK | 90 | 1979.15 | |
| 10-1 | 46 - 60 | G(Cit)K(Iph)RK(Iph)(Pip)RDPKWKK | 90 | 2379.04 | |
| 10-1 | 46 - 60 | GVHSPR(Iph)HV(Dab)(Iph)KWKK | 90 | 2227.91 | |
| 10-1 | 46 - 60 | GKHP(Iph)RKKA(Dab)(Nap)KWKK | 88 | 2185.12 | |
| 10-1 | 46 - 60 | GGVKHHG(Iph)L(Iph)HKWKK | 88 | 2180.87 | |
| 10-1 | 46 - 60 | G(Pip)V(Dab)LRKPPM(Pip)KWKK | 86 | 1943.19 | |
| 10-1 | 46 - 60 | GTH(Iph)R(Cit)A(Cit)R(Nap)KKWKK | 86 | 2303.13 | |
| 10-1 | 46 - 60 | GRH(Cit)RMSTRASKWKK (+13.03) | 85 | 2022.14 | |
| 10-1 | 46 - 60 | GGKNHQA(Dab)G(Iph)RKWKK | 83 | 1990.94 | |
| 10-1 | 46 - 60 | GRHR(Pal)RF(Cit)DKLKWKK | 83 | 2183.26 | |
| 10-1 | 46 - 60 | GETKPR(Iph)K(Dab)(Iph)FKWKK | 83 | 2301.97 | |
| 10-1 | 46 - 60 | GGMA(Dab)P(Cit)R(Dab)T(Dab)KWKK | 82 | 1840.05 | |
| 10-1 | 46 - 60 | GKP(Cit)N(Dab)(Iph)RRNQKWKK | 82 | 2193.08 | |
| 10-1 | 46 - 60 | G(Cit)(Pip)NQFRGR(Nap)KKWKK | 81 | 2136.21 | |
| 10-1 | 46 - 60 | GRRVLLHF(Dab)VRKWKK | 81 | 2046.27 | |
| **10-1** | **46 - 60** | **GP(Pip)G(Dab)K(Dab)SV(Iph)(Nap)KWKK** | **80** | **2034.00** | **CXP6** |
| 10-1 | 46 - 60 | GHKSK(Iph)LLPERKWKK | 80 | 2131.08 | |
| 10-1 | 46 - 60 | GAN(Dab)F(Cit)THKHHKWKK | 79 | 1999.09 | |

| | | | | |
|---|---|---|---|---|
| 10-1 | 46 - 60 | G(Pal)HPRRG(Pip)TPNKWKK | 78 | 1959.09 |
| 10-1 | 46 - 60 | GT(Pip)H(Iph)R(Dab)KEKNKWKK | 78 | 2162.06 |
| 10-1 | 46 - 60 | G(Dab)SMV(Dab)HA(Cit)RPKWKK | 77 | 1905.07 |
| 10-1 | 46 - 60 | GRRLH(Iph)FTPPRKWKK | 76 | 2203.10 |
| 10-1 | 46 - 60 | G(Nap)(Dab)AHGS(Iph)RF(Dab)KWKK | 75 | 2094.97 |
| 10-1 | 46 - 60 | GHHTN(Dab)Q(Iph)(Iph)KRKWKK | 72 | 2316.93 |
| 10-1 | 46 - 60 | G(Nap)EAPPH(Pip)KVRKWKK | 72 | 2007.14 |

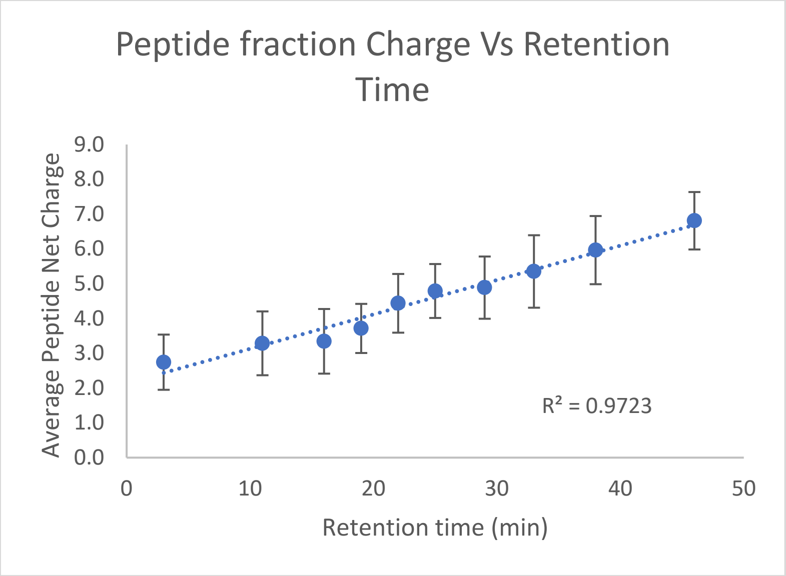

| Pool | Start time | Average Net charge |
|---|---|---|
| 01 | 3 | 2.7 |
| 02 | 11 | 3.3 |
| 03 | 16 | 3.3 |
| 04 | 19 | 3.7 |
| 05 | 22 | 4.4 |
| 06 | 25 | 4.8 |
| 07 | 29 | 4.9 |
| 08 | 33 | 5.4 |
| 09 | 38 | 6.0 |
| 10 | 46 | 6.8 |

**Library 2: 3,000-member library**

| Fraction Label | Fraction Retention Time (min) | | Sequencing confidence score (ALC) | peptide mass (Da) |
|---|---|---|---|---|
| 01-2 | 1-8 | GNAADSGGDSSKWKK | 96 | 1630.7812 |
| 01-2 | 1-8 | G(Pip)(Cit)DSEN(Cit)TGDKWKK | 96 | 1927.9614 |
| 01-2 | 1-8 | GPQSSTHADDLKWKK | 95 | 1820.9282 |
| 01-2 | 1-8 | G(Dab)QDEENLGQGKWKK | 95 | 1839.9341 |
| 01-2 | 1-8 | GASDKDS(Nap)DQDKWKK | 95 | 1927.9175 |
| 01-2 | 1-8 | GGDDE(Cit)VEFEPKWKK | 95 | 1943.949 |
| 01-2 | 1-8 | G(Nap)(Cit)NGNTNEFDKWKK | 95 | 2014.9761 |
| 01-2 | 1-8 | GV(Cit)GDAASQTVKWKK | 93 | 1754.9541 |
| 01-2 | 1-8 | GEQVSGKLDTAKWKK | 93 | 1797.9849 |
| 01-2 | 1-8 | GVEDEENLGQGKWKK | 93 | 1839.9229 |
| 01-2 | 1-8 | G(Cit)PQNADTSGFKWKK | 93 | 1843.9443 |
| 01-2 | 1-8 | GADTNEVPLQFKWKK | 93 | 1884.0007 |
| 01-2 | 1-8 | G(Pip)DS(Cit)VQ(Cit)EAVKWKK | 93 | 1938.0549 |
| 01-2 | 1-8 | GK(Cit)DDDE(Pip)MMAKWKK | 93 | 1987.9722 |
| 01-2 | 1-8 | GRENDSAGTGGKWKK | 92 | 1713.866 |
| 01-2 | 1-8 | GQEQVAGVADEKWKK | 92 | 1795.9329 |
| 01-2 | 1-8 | G(Dab)D(Pip)SDGGDQQKWKK | 92 | 1797.8872 |
| 01-2 | 1-8 | GVAQV(Cit)GPQMEKWKK | 92 | 1866.0046 |
| 01-2 | 1-8 | GGAT(Cit)MMD(Cit)QLKWKK | 92 | 1930.9983 |
| 01-2 | 1-8 | GEAFTGKDDPAKWKK | 91 | 1800.927 |
| 01-2 | 1-8 | GHFQEALASDPKWKK | 91 | 1864.9697 |
| 01-2 | 1-8 | GVDSNSFV(Cit)DSKWKK | 91 | 1876.9546 |
| 01-2 | 1-8 | GNDTPFLEDSPKWKK | 91 | 1884.9482 |
| 01-2 | 1-8 | GS(Cit)EEE(Dab)QVLVKWKK | 91 | 1940.0593 |
| 01-2 | 1-8 | GDETGPGFSYPKWKK | 90 | 1819.9006 |
| 01-2 | 1-8 | GAMNGNQNDDMKWKK | 90 | 1859.8518 |
| 01-2 | 1-8 | GGATVDNATGVKWKK | 89 | 1654.8904 |
| 01-2 | 1-8 | GGFEDSPLNMGKWKK | 89 | 1816.9043 |
| 01-2 | 1-8 | GVVEEPT(Cit)EQQKWKK | 89 | 1966.0386 |
| 01-2 | 1-8 | GG(Cit)MAEEEPQLKWKK | 88 | 1910.9785 |
| 01-2 | 1-8 | GETQNEPLQLNKWKK | 88 | 1936.0278 |

| | | | | |
|---|---|---|---|---|
| 01-2 | 1-8 | GGA(Nap)VTEAGGGKWKK | 87 | 1665.874 |
| 01-2 | 1-8 | GQSPT(Iph)ADEQPKWKK | 87 | 1995.8452 |
| 01-2 | 1-8 | GEKS(Cit)N(Cit)TDENKWKK | 87 | 2001.0142 |
| 01-2 | 1-8 | GQAVP(Cit)(Cit)GDMVKWKK | 86 | 1881.0156 |
| 01-2 | 1-8 | GGGD(Nap)(Pal)LEDSPKWKK | 86 | 1884.927 |
| 01-2 | 1-8 | GVAENADTSGFKWKK | 85 | 1760.8958 |
| 01-2 | 1-8 | GPEDDSGQASNKWKK | 85 | 1769.8445 |
| 01-2 | 1-8 | GQFRPLEGHMDKWKK | 85 | 1980.0264 |
| 01-2 | 1-8 | GLSDSEN(Cit)TGDKWKK | 84 | 1844.9131 |
| 01-2 | 1-8 | GHAEGTDFDVFKWKK | 84 | 1887.938 |
| 01-2 | 1-8 | GEVE(Dab)QSGSDEKWKK | 83 | 1829.9021 |
| 01-2 | 1-8 | GN(Cit)NGNTNEMMKWKK | 83 | 1931.9207 |
| 01-2 | 1-8 | GFQF(Cit)TPALEDKWKK | 83 | 1975.043 |
| 01-2 | 1-8 | GGNVSNANNSFKWKK | 82 | 1773.9023 |
| 01-2 | 1-8 | GMSNGNQNDDMKWKK | 82 | 1875.8469 |
| 01-2 | 1-8 | GRE(Cit)DEDQ(Cit)ELKWKK | 82 | 2098.0669 |
| 01-2 | 1-8 | GDSF(Dab)SVAD(Pip)QKWKK | 81 | 1844.9646 |
| 01-2 | 1-8 | GPLDNAGGRDDKWKK | 80 | 1779.9128 |
| 01-2 | 1-8 | GPFASSSLVETKWKK | 80 | 1787.9683 |
| 01-2 | 1-8 | GA(Nap)AGDDAEEMKWKK | 80 | 1855.8674 |
| 01-2 | 1-8 | GQAVP(Cit)DVDMVKWKK | 80 | 1881.0044 |
| 01-2 | 1-8 | GSDLQQQ(Cit)HADKWKK | 80 | 1948.998 |
| 01-2 | 1-8 | GGA(Pal)H(Iph)ADEQPKWKK | 80 | 1995.8352 |
| 01-2 | 1-8 | GEL(Dab)DDET(Iph)VDKWKK | 80 | 2058.866 |
| 01-2 | 1-8 | GGTGDTHEELTKWKK | 79 | 1809.9121 |
| 01-2 | 1-8 | G(Dab)GSQVA(Nap)DA(Cit)KWKK | 79 | 1851.9856 |
| 01-2 | 1-8 | GTQ(Iph)GVGQNEDKWKK | 79 | 1970.8247 |
| 01-2 | 1-8 | GQAVP(Cit)(Cit)GMETKWKK | 78 | 1897.0107 |
| 01-2 | 1-8 | G(Cit)DPGLFLE(Cit)QKWKK | 78 | 1983.0803 |
| 01-2 | 1-8 | GQDD(Cit)FGAVLNKWKK | 77 | 1885.9912 |
| 01-2 | 1-8 | GTTPSFQNMTDKWKK | 77 | 1891.9363 |
| 01-2 | 1-8 | GGMSR(Cit)FAEDEKWKK | 77 | 1948.969 |
| 01-2 | 1-8 | GM(Cit)PEDEAEEFKWKK | 77 | 2003.9524 |
| 01-2 | 1-8 | GGTGDTHE(Cit)SVKWKK | 76 | 1809.9236 |
| 01-2 | 1-8 | GS(Nap)QEALASDPKWKK | 76 | 1864.9583 |
| 01-2 | 1-8 | GAR(Iph)EAPD(Cit)GGKWKK | 76 | 1952.8618 |
| 01-2 | 1-8 | GFGLVDGATRDKWKK | 75 | 1800.9746 |
| 01-2 | 1-8 | GTAG(Iph)RGQNEDKWKK | 75 | 1970.8359 |
| 01-2 | 1-8 | GFQYLEPALEDKWKK | 75 | 1975.0315 |

| | | | | |
|---|---|---|---|---|
| 01-2 | 1-8 | G(Cit)DDVLGKDDLKWKK | 74 | 1897.0171 |
| 01-2 | 1-8 | G(Pip)SVSFFVNDNKWKK | 74 | 1905.001 |
| 01-2 | 1-8 | GM(Pal)NYY(Cit)SGGEKWKK | 74 | 1975.9475 |
| 01-2 | 1-8 | GQVG(Iph)QNDNNDKWKK | 74 | 2026.8259 |
| 01-2 | 1-8 | GKE(Iph)SVEDEQSKWKK | 74 | 2073.8767 |
| 01-2 | 1-8 | GQT(Cit)FDDVEGLKWKK | 73 | 1931.0015 |
| 01-2 | 1-8 | G(Dab)HDQNDNEE(Iph)KWKK | 73 | 2123.842 |
| 01-2 | 1-8 | GHFAGED(Dab)DDAKWKK | 72 | 1826.8811 |
| 01-2 | 1-8 | GA(Nap)AGDDAQNMKWKK | 72 | 1839.8838 |
| 01-2 | 1-8 | GVAQV(Cit)EDNAEKWKK | 72 | 1881.981 |
| 01-2 | 1-8 | GSDLE(Pip)E(Cit)HADKWKK | 72 | 1948.9868 |
| 01-2 | 1-8 | GLETAGDESDEKWKK | 71 | 1815.8752 |
| 01-2 | 1-8 | GVDG(Dab)(Dab)DENFMKWKK | 71 | 1876.9365 |
| 01-2 | 1-8 | GQNLQENE(Cit)GGKWKK | 71 | 1895.9714 |
| 01-2 | 1-8 | GGDG(Dab)VTMTGLKWKK | 70 | 1700.9143 |
| 01-2 | 1-8 | GAAP(Nap)GA(Cit)EGDKWKK | 70 | 1791.9167 |
| 01-2 | 1-8 | GQETG(Dab)SPFE(Nap)KWKK | 70 | 1941.9849 |
| 01-2 | 1-8 | GTAG(Iph)S(Pip)QNEDKWKK | 70 | 1970.8247 |
| 01-2 | 1-8 | GK(Cit)DDDE(Pip)YADKWKK | 70 | 2003.9814 |
| 02-2 | 8-11 | GQLTNMNNQQVKWKK | 70 | 1940.0164 |
| 02-2 | 8-11 | GAEMFR(Cit)NEPNKWKK | 70 | 2015.0273 |
| 02-2 | 8-11 | GE(Cit)HLGGTE(Dab)EKWKK | 71 | 1878.9814 |
| 02-2 | 8-11 | GMDSDK(Cit)ENGVKWKK | 71 | 1901.9531 |
| 02-2 | 8-11 | G(Iph)KGPANNDMYKWKK | 71 | 2032.8591 |
| 02-2 | 8-11 | GVEAVNTSG(Cit)KWKK | 72 | 1830.9854 |
| 02-2 | 8-11 | GSNGLFVVGAGKWKK | 73 | 1670.937 |
| 02-2 | 8-11 | GP(Dab)QFNQEEDFKWKK | 73 | 2003.9966 |
| 02-2 | 8-11 | GPN(Cit)K(Cit)TE(Iph)QNKWKK | 73 | 2167.9888 |
| 02-2 | 8-11 | GTTFS(Cit)NQ(Pip)MNKWKK | 74 | 1976.0164 |
| 02-2 | 8-11 | G(Pip)EPSSQ(Cit)AN(Iph)KWKK | 74 | 2038.8987 |
| 02-2 | 8-11 | GPESEAGGVVTKWKK | 75 | 1695.9055 |
| 02-2 | 8-11 | GLVEG(Dab)ENTSSKWKK | 75 | 1785.9485 |
| 02-2 | 8-11 | G(Pip)EPSENL(Cit)(Iph)NKWKK | 75 | 2108.9404 |
| 02-2 | 8-11 | GTRPASMNDVVKWKK | 76 | 1839.989 |
| 02-2 | 8-11 | GQKNNNDLPLSKWKK | 76 | 1893.0334 |
| 02-2 | 8-11 | GL(Cit)(Dab)QGHMQDQKWKK | 77 | 1964.0276 |
| 02-2 | 8-11 | GDDH(Dab)VSDHDSKWKK | 78 | 1876.8928 |
| 02-2 | 8-11 | GVEK(Cit)SGFANQKWKK | 78 | 1887.0227 |
| 02-2 | 8-11 | G(Iph)GQPTVTTAGKWKK | 79 | 1854.8391 |

| | | | | |
|---|---|---|---|---|
| 02-2 | 8-11 | GHRH(Cit)SNAD(Dab)GKWKK | 79 | 1900.9995 |
| 02-2 | 8-11 | GTDNV(Iph)SD(Cit)E(Dab)KWKK | 79 | 2059.8726 |
| 02-2 | 8-11 | G(Iph)ANGGGAGATKWKK | 80 | 1698.7239 |
| 02-2 | 8-11 | GMKG(Cit)AN(Cit)(Cit)GGKWKK | 80 | 1856.0066 |
| 02-2 | 8-11 | GASLGSTTV(Cit)(Cit)KWKK | 81 | 1800.012 |
| 02-2 | 8-11 | GFKSESDREFQKWKK | 81 | 2023.0388 |
| 02-2 | 8-11 | GAE(Dab)LGSNQGGKWKK | 82 | 1682.8965 |
| 02-2 | 8-11 | GAKDNSGFVD(Cit)KWKK | 82 | 1859.9756 |
| 02-2 | 8-11 | GGSTDTRFRDEKWKK | 82 | 1933.9871 |
| 02-2 | 8-11 | GT(Cit)GKQDLRDEKWKK | 82 | 1969.0608 |
| 02-2 | 8-11 | GDD(Dab)EQ(Dab)LDT(Iph)KWKK | 82 | 2058.8772 |
| 02-2 | 8-11 | GALADRKPADSKWKK | 83 | 1794.0012 |
| 02-2 | 8-11 | G(Dab)QTS(Cit)P(Cit)NQEKWKK | 83 | 1968.0403 |
| 02-2 | 8-11 | GVDDD(Iph)DTEKHKWKK | 83 | 2096.8564 |
| 02-2 | 8-11 | GGTGDTHEELTKWKK | 84 | 1809.9121 |
| 02-2 | 8-11 | GTNTNQQASDEKWKK | 84 | 1857.9082 |
| 02-2 | 8-11 | GVQPG(Dab)(Cit)ND(Cit)FKWKK | 84 | 1941.0447 |
| 02-2 | 8-11 | GRNTPG(Cit)TEFQKWKK | 84 | 1957.0396 |
| 02-2 | 8-11 | GGETDTHHMGPKWKK | 85 | 1831.8901 |
| 02-2 | 8-11 | GEFNETHEDSHKWKK | 85 | 1994.9348 |
| 02-2 | 8-11 | GTNTNQQASNNKWKK | 86 | 1841.9246 |
| 02-2 | 8-11 | G(Cit)PQSEANVVTKWKK | 86 | 1852.0068 |
| 02-2 | 8-11 | GAYELDA(Cit)D(Dab)PKWKK | 86 | 1900.9907 |
| 02-2 | 8-11 | G(Iph)(Pip)EPEGLEAAKWKK | 87 | 1964.8757 |
| 02-2 | 8-11 | GMKANPSNE(Iph)TKWKK | 87 | 2014.8696 |
| 02-2 | 8-11 | GNRGESEPGTNKWKK | 88 | 1810.9187 |
| 02-2 | 8-11 | GQ(Cit)HE(Cit)T(Iph)AGQKWKK | 88 | 2107.9314 |
| 02-2 | 8-11 | GLAGNDG(Dab)DV(Cit)KWKK | 89 | 1767.9492 |
| 02-2 | 8-11 | GGNVSNANNSFKWKK | 89 | 1773.9023 |
| 02-2 | 8-11 | G(Pip)DNSGFVD(Dab)GKWKK | 89 | 1786.9229 |
| 02-2 | 8-11 | GLTDN(Dab)QANQGKWKK | 89 | 1810.9551 |
| 02-2 | 8-11 | GAANPRGGDQ(Iph)KWKK | 89 | 1908.8357 |
| 02-2 | 8-11 | G(Iph)(Pip)EPEGLN(Dab)GKWKK | 89 | 1964.887 |
| 02-2 | 8-11 | GSDSHSFVAGTKWKK | 90 | 1757.8962 |
| 02-2 | 8-11 | GV(Cit)TNDEKSNSKWKK | 90 | 1900.9868 |
| 02-2 | 8-11 | GAKAND(Cit)GVP(Iph)KWKK | 90 | 1951.9031 |
| 02-2 | 8-11 | G(Pip)RDNSGFVD(Cit)KWKK | 91 | 1943.0239 |
| 02-2 | 8-11 | GQTEGGASDK(Cit)KWKK | 92 | 1799.9392 |
| 02-2 | 8-11 | GDVSVQL(Iph)DAGKWKK | 92 | 1926.8601 |

| | | | | |
|---|---|---|---|---|
| 02-2 | 8-11 | GNKVQMMDHS(Cit)KWKK | 92 | 1997.02 |
| 02-2 | 8-11 | GSSLQHNLSDAKWKK | 93 | 1821.9597 |
| 02-2 | 8-11 | GF(Cit)SFQGNVNTKWKK | 93 | 1921.0071 |
| 02-2 | 8-11 | GHQTDPPTFQLKWKK | 93 | 1934.0276 |
| 02-2 | 8-11 | G(Pip)SKDNTA(Cit)(Cit)DKWKK | 93 | 1941.0295 |
| 02-2 | 8-11 | GP(Dab)QFNQT(Cit)DMKWKK | 93 | 1988.0164 |
| 02-2 | 8-11 | GTTEVNQLPSVKWKK | 94 | 1838.0164 |
| 02-2 | 8-11 | G(Dab)QTAEANLHDKWKK | 95 | 1848.9707 |
| 02-2 | 8-11 | GVGQHDQFD(Pip)SKWKK | 96 | 1908.9707 |
| 02-2 | 8-11 | G(Dab)(Iph)STQGNESAKWKK | 96 | 1916.8142 |
| 03-2 | 11-14 | GPGADGTARQGKWKK | 84 | 1679.8967 |
| 03-2 | 11-14 | GAE(Dab)LGSNQGGKWKK | 87 | 1682.8965 |
| 03-2 | 11-14 | GQKATQGPGSTKWKK | 90 | 1724.9434 |
| 03-2 | 11-14 | GASAVEGNSNKKWKK | 92 | 1726.9226 |
| 03-2 | 11-14 | G(Pal)PPSHMLGGGKWKK | 73 | 1750.9202 |
| 03-2 | 11-14 | GVTDKAGDHTGKWKK | 82 | 1750.9226 |
| 03-2 | 11-14 | GVSEPE(Dab)GETGKWKK | 78 | 1754.9062 |
| 03-2 | 11-14 | GTQS(Dab)TNGTGFKWKK | 91 | 1762.9226 |
| 03-2 | 11-14 | G(Pip)NTSLAAPNNKWKK | 80 | 1777.97 |
| 03-2 | 11-14 | GDPSLVGQRVGKWKK | 94 | 1778.0063 |
| 03-2 | 11-14 | GKSSGVHV(Cit)GDKWKK | 80 | 1792.981 |
| 03-2 | 11-14 | GMGANEKLM(Dab)GKWKK | 86 | 1800.9604 |
| 03-2 | 11-14 | G(Dab)TLGMDDMGHKWKK | 92 | 1826.9033 |
| 03-2 | 11-14 | G(Dab)AMELDQT(Dab)AKWKK | 95 | 1828.9729 |
| 03-2 | 11-14 | GM(Dab)QESGGEKLKWKK | 80 | 1828.9729 |
| 03-2 | 11-14 | GPHSES(Dab)QLSPKWKK | 79 | 1831.9805 |
| 03-2 | 11-14 | G(Dab)LDPSHMLGNKWKK | 87 | 1833.9785 |
| 03-2 | 11-14 | GG(Dab)QPDPFQA(Pip)KWKK | 84 | 1835.9907 |
| 03-2 | 11-14 | GTSKEGKELSNKWKK | 95 | 1843.0063 |
| 03-2 | 11-14 | GMFGVDDTRSAKWKK | 85 | 1848.9417 |
| 03-2 | 11-14 | GVETAEANLHDKWKK | 85 | 1848.9595 |
| 03-2 | 11-14 | G(Dab)QTAEANLHDKWKK | 96 | 1848.9707 |
| 03-2 | 11-14 | GLNNG(Pip)AEDDKKWKK | 82 | 1851.9705 |
| 03-2 | 11-14 | G(Dab)VSL(Cit)PF(Cit)AGKWKK | 84 | 1855.0693 |
| 03-2 | 11-14 | GKASL(Cit)PF(Cit)AGKWKK | 74 | 1855.0696 |
| 03-2 | 11-14 | G(Pip)E(Pip)GE(Cit)FGGNKWKK | 73 | 1868.9758 |
| 03-2 | 11-14 | GAD(Pal)(Nap)(Pip)ALTTKWKK | 82 | 1870.0002 |
| 03-2 | 11-14 | GG(Iph)GLSGRSSEKWKK | 73 | 1872.8245 |
| 03-2 | 11-14 | GFVEF(Dab)ATVVDKWKK | 74 | 1877.031 |

| | | | | |
|---|---|---|---|---|
| 03-2 | 11-14 | G(Nap)LGNAN(Cit)D(Dab)AKWKK | 90 | 1878.9966 |
| 03-2 | 11-14 | GTHSM(Cit)PS(Cit)SAKWKK | 75 | 1881.9746 |
| 03-2 | 11-14 | GV(Cit)AQHHGVDNKWKK | 93 | 1883.998 |
| 03-2 | 11-14 | GKADR(Cit)MDAAVKWKK | 89 | 1884.0266 |
| 03-2 | 11-14 | GSESGGN(Iph)KPSKWKK | 93 | 1885.8083 |
| 03-2 | 11-14 | GKPPNQQLPG(Cit)KWKK | 79 | 1886.0752 |
| 03-2 | 11-14 | GFDTNQ(Nap)(Dab)TGGKWKK | 72 | 1886.9539 |
| 03-2 | 11-14 | GEVK(Cit)SGFANQKWKK | 78 | 1887.0229 |
| 03-2 | 11-14 | G(Pal)GFADS(Cit)SLHKWKK | 73 | 1888.981 |
| 03-2 | 11-14 | G(Pal)SNQNDPSM(Dab)KWKK | 72 | 1890.927 |
| 03-2 | 11-14 | GHENGHTTDS(Pal)KWKK | 91 | 1895.9138 |
| 03-2 | 11-14 | GST(Cit)LQENKGLKWKK | 95 | 1897.0647 |
| 03-2 | 11-14 | G(Dab)EDFGAEDHKKWKK | 71 | 1897.9546 |
| 03-2 | 11-14 | GRQLGMDDMGHKWKK | 97 | 1909.9517 |
| 03-2 | 11-14 | GTT(Iph)LNSNAAPKWKK | 90 | 1911.8604 |
| 03-2 | 11-14 | GPTNEDAEHHLKWKK | 86 | 1912.9656 |
| 03-2 | 11-14 | G(Iph)SGPSSE(Dab)QVKWKK | 82 | 1913.8396 |
| 03-2 | 11-14 | GFKEES(Dab)QLSPKWKK | 91 | 1915.0427 |
| 03-2 | 11-14 | GVA(Pip)E(Cit)PNFLVKWKK | 82 | 1922.1003 |
| 03-2 | 11-14 | GAKNDAQEN(Nap)SKWKK | 90 | 1923.9702 |
| 03-2 | 11-14 | GAKNDAQENHFKWKK | 92 | 1923.9817 |
| 03-2 | 11-14 | GRFQGVESP(Nap)GKWKK | 76 | 1924.022 |
| 03-2 | 11-14 | GN(Cit)KEGKELSNKWKK | 93 | 1926.0549 |
| 03-2 | 11-14 | G(Dab)T(Cit)DNLQDDVKWKK | 73 | 1927.0024 |
| 03-2 | 11-14 | GLHTDVFEKTSKWKK | 95 | 1927.0427 |
| 03-2 | 11-14 | GHGGSEHEFM(Pal)KWKK | 72 | 1928.9216 |
| 03-2 | 11-14 | GSKPDF(Pip)TDDKKWKK | 75 | 1929.0222 |
| 03-2 | 11-14 | GSFFHEMNSPSKWKK | 75 | 1932.9417 |
| 03-2 | 11-14 | G(Nap)(Dab)DKDEHNGAKWKK | 76 | 1932.9707 |
| 03-2 | 11-14 | GESST(Dab)GSM(Iph)NKWKK | 83 | 1935.791 |
| 03-2 | 11-14 | GMD(Nap)RGNSVVDKWKK | 74 | 1939.9839 |
| 03-2 | 11-14 | GLEF(Nap)DKGEG(Dab)KWKK | 90 | 1942.0212 |
| 03-2 | 11-14 | GMSSKPQL(Cit)LMKWKK | 71 | 1942.0757 |
| 03-2 | 11-14 | G(Iph)RSAEAASNNKWKK | 86 | 1942.8411 |
| 03-2 | 11-14 | GGF(Dab)HGEDFM(Cit)KWKK | 89 | 1946.9685 |
| 03-2 | 11-14 | GLEPKSLHMQDKWKK | 93 | 1948.0464 |
| 03-2 | 11-14 | GLE(Pip)VSLHMQDKWKK | 70 | 1948.0466 |
| 03-2 | 11-14 | GE(Cit)(Dab)KPES(Cit)QAKWKK | 84 | 1953.0657 |
| 03-2 | 11-14 | GVET(Dab)TPG(Cit)A(Iph)KWKK | 90 | 1954.9026 |

| | | | | |
|---|---|---|---|---|
| 03-2 | 11-14 | GRFQGVEN(Pip)DDKWKK | 73 | 1956.0078 |
| 03-2 | 11-14 | G(Iph)RSAEAASDEKWKK | 88 | 1958.8247 |
| 03-2 | 11-14 | G(Pip)STE(Cit)A(Nap)QGHKWKK | 76 | 1960.0181 |
| 03-2 | 11-14 | GKMSTE(Dab)(Cit)VEMKWKK | 94 | 1962.0293 |
| 03-2 | 11-14 | GMHEK(Cit)LVLEGKWKK | 91 | 1963.094 |
| 03-2 | 11-14 | GESSS(Nap)HAFHDKWKK | 88 | 1963.9441 |
| 03-2 | 11-14 | GSFDT(Dab)TEF(Nap)AKWKK | 80 | 1964.9895 |
| 03-2 | 11-14 | G(Dab)VK(Nap)NNEEGQKWKK | 85 | 1965.0332 |
| 03-2 | 11-14 | GN(Cit)V(Cit)(Dab)PT(Cit)NVKWKK | 79 | 1965.1135 |
| 03-2 | 11-14 | GS(Cit)ETEAEEHKKWKK | 93 | 1966.9973 |
| 03-2 | 11-14 | GFPETEAEEHKKWKK | 76 | 1967.0012 |
| 03-2 | 11-14 | GHSLD(Dab)NQT(Cit)FKWKK | 71 | 1969.0396 |
| 03-2 | 11-14 | G(Dab)(Cit)AFEGP(Iph)LGKWKK | 72 | 1970.9128 |
| 03-2 | 11-14 | G(Dab)F(Iph)DGQVVSPKWKK | 89 | 1971.8967 |
| 03-2 | 11-14 | GNDHHVA(Cit)FPQKWKK | 85 | 1972.0293 |
| 03-2 | 11-14 | GN(Dab)G(Iph)TSPS(Cit)MKWKK | 93 | 1973.8542 |
| 03-2 | 11-14 | GM(Dab)TEDK(Iph)A(Dab)GKWKK | 91 | 1974.8748 |
| 03-2 | 11-14 | GTTFS(Cit)NQKENKWKK | 75 | 1976.0342 |
| 03-2 | 11-14 | GPSRNDFPQHEKWKK | 80 | 1977.0081 |
| 03-2 | 11-14 | GQNG(Nap)G(Cit)KEPFKWKK | 77 | 1981.0435 |
| 03-2 | 11-14 | GTFTKDNEHMNKWKK | 96 | 1986.9846 |
| 03-2 | 11-14 | GRE(Iph)LNSNAAPKWKK | 83 | 1994.9087 |
| 03-2 | 11-14 | GTS(Nap)KQDAERLKWKK | 75 | 1995.0801 |
| 03-2 | 11-14 | GKGD(Cit)NDSG(Iph)(Pip)KWKK | 71 | 1998.8674 |
| 03-2 | 11-14 | GGND(Pip)GP(Iph)QRDKWKK | 87 | 2007.8677 |
| 03-2 | 11-14 | GRQ(Cit)DNLQDDVKWKK | 94 | 2010.0508 |
| 03-2 | 11-14 | GFES(Iph)GFKDNGKWKK | 81 | 2023.8552 |
| 03-2 | 11-14 | GTEMT(Pip)VG(Iph)(Pip)DKWKK | 73 | 2027.8901 |
| 03-2 | 11-14 | G(Iph)NRSSEQD(Dab)AKWKK | 72 | 2029.873 |
| 03-2 | 11-14 | GQDGV(Nap)(Cit)(Dab)(Cit)EMKWKK | 80 | 2040.0476 |
| 03-2 | 11-14 | GK(Nap)ADMS(Dab)(Cit)E(Cit)KWKK | 84 | 2042.0632 |
| 03-2 | 11-14 | G(Nap)(Cit)HG(Cit)PVKEDKWKK | 84 | 2043.0916 |
| 03-2 | 11-14 | G(Iph)NMSKEQD(Dab)AKWKK | 73 | 2045.8755 |
| 03-2 | 11-14 | GEPRGVV(Iph)NHDKWKK | 77 | 2045.9197 |
| 03-2 | 11-14 | G(Iph)DSNQFSVQTKWKK | 72 | 2048.8718 |
| 03-2 | 11-14 | G(Pip)VT(Cit)TN(Cit)PA(Iph)KWKK | 76 | 2065.9824 |
| 03-2 | 11-14 | GNFN(Iph)PN(Cit)VGEKWKK | 86 | 2070.9038 |
| 03-2 | 11-14 | GVE(Iph)(Cit)PN(Pip)(Cit)GPKWKK | 70 | 2075.9668 |
| 03-2 | 11-14 | GMTRH(Iph)AD(Cit)DGKWKK | 74 | 2082.8821 |

| | | | | |
|---|---|---|---|---|
| 03-2 | 11-14 | GQ(Cit)HE(Cit)T(Iph)AGQKWKK | 88 | 2107.9314 |
| 03-2 | 11-14 | GLK(Cit)(Nap)D(Cit)EE(Dab)MKWKK | 86 | 2126.1206 |
| 03-2 | 11-14 | GR(Cit)SE(Nap)REMEPKWKK | 74 | 2138.0955 |
| 03-2 | 11-14 | GKDTN(Iph)PN(Cit)REKWKK | 84 | 2153.9731 |
| 03-2 | 11-14 | GFK(Iph)QETHE(Pip)QKWKK | 85 | 2195.9878 |
| 04-2 | 14-16 | GPTNQGVQLGAKWKK | 83 | 1734.9641 |
| 04-2 | 14-16 | GPASS(Dab)VSDQ(Dab)KWKK | 73 | 1740.9382 |
| 04-2 | 14-16 | GA(Dab)SDAG(Dab)(Iph)GGKWKK | 93 | 1757.761 |
| 04-2 | 14-16 | GTDLSTNGTGFKWKK | 74 | 1762.9114 |
| 04-2 | 14-16 | GTS(Cit)ATNGTGFKWKK | 79 | 1762.9229 |
| 04-2 | 14-16 | GPDPPPQDDGTKWKK | 71 | 1788.8906 |
| 04-2 | 14-16 | GKSSGKG(Cit)CA(Pal)KWKK | 75 | 1792.9631 |
| 04-2 | 14-16 | GSKSGVHVVDDKWKK | 94 | 1792.9697 |
| 04-2 | 14-16 | GKSSGVHV(Dab)NDKWKK | 79 | 1792.9807 |
| 04-2 | 14-16 | GKSSGVHV(Cit)GDKWKK | 98 | 1792.981 |
| 04-2 | 14-16 | GL(Dab)DN(Dab)APCPNKWKK | 80 | 1793.947 |
| 04-2 | 14-16 | GTAVED(Pip)QTTGKWKK | 84 | 1797.9487 |
| 04-2 | 14-16 | GGA(Dab)QWQAGT(Pal)KWKK | 87 | 1816.9597 |
| 04-2 | 14-16 | GAGHSELLKQSKWKK | 75 | 1820.0168 |
| 04-2 | 14-16 | GPDRGMSMGRAKWKK | 80 | 1827.946 |
| 04-2 | 14-16 | GHQNHG(Cit)PGGEKWKK | 84 | 1839.9355 |
| 04-2 | 14-16 | GTHGNAHNNMPKWKK | 79 | 1842.9172 |
| 04-2 | 14-16 | GTHGNAHNENVKWKK | 91 | 1842.9351 |
| 04-2 | 14-16 | GASKQQEVG(Cit)DKWKK | 70 | 1868.9971 |
| 04-2 | 14-16 | GASRDLEVTPMKWKK | 71 | 1869.0044 |
| 04-2 | 14-16 | GASR(Dab)QEVTPMKWKK | 81 | 1869.0154 |
| 04-2 | 14-16 | GPAHS(Pip)(Nap)NDGTKWKK | 92 | 1871.9543 |
| 04-2 | 14-16 | G(Nap)(Dab)GQT(Dab)LPSEKWKK | 80 | 1879.0215 |
| 04-2 | 14-16 | GTLDANQMEHAKWKK | 75 | 1879.9475 |
| 04-2 | 14-16 | GDESKQ(Pip)SSA(Cit)KWKK | 71 | 1884.9919 |
| 04-2 | 14-16 | GLNLPANAHQ(Cit)KWKK | 77 | 1885.0547 |
| 04-2 | 14-16 | GMT(Pip)TANEDENKWKK | 78 | 1900.9214 |

| | | | | |
|---|---|---|---|---|
| 04-2 | 14-16 | GTMLNANEDENKWKK | 77 | 1900.9214 |
| 04-2 | 14-16 | GMT(Pip)TANEDENKWKK | 76 | 1900.9214 |
| 04-2 | 14-16 | GTMLNANEDENKWKK | 76 | 1900.9214 |
| 04-2 | 14-16 | G(Dab)SQEDDKTEVKWKK | 86 | 1900.9756 |
| 04-2 | 14-16 | GVSEEDDKTQ(Dab)KWKK | 83 | 1900.9756 |
| 04-2 | 14-16 | GS(Dab)QEDDKTEVKWKK | 81 | 1900.9756 |
| 04-2 | 14-16 | GDLSEDDKTQ(Dab)KWKK | 71 | 1900.9756 |
| 04-2 | 14-16 | G(Dab)SQEDDKTQ(Dab)KWKK | 91 | 1900.9868 |
| 04-2 | 14-16 | GTTDCH(Pal)HHTGKWKK | 77 | 1906.9121 |
| 04-2 | 14-16 | GF(Dab)DEKEVDNQKWKK | 86 | 1974.0071 |
| 04-2 | 14-16 | GF(Dab)DEKE(Dab)NNQKWKK | 96 | 1974.0183 |
| 04-2 | 14-16 | GKGD(Cit)NHT(Cit)NQKWKK | 79 | 1978.0359 |
| 04-2 | 14-16 | GR(Iph)(Cit)GANAKADKWKK | 78 | 1982.9202 |
| 04-2 | 14-16 | GEWKSNEHSCNKWKK | 77 | 1983.9485 |
| 04-2 | 14-16 | GKT(Iph)DAHKGTDKWKK | 96 | 1995.8928 |
| 04-2 | 14-16 | G(Dab)SDH(Iph)QTTMAKWKK | 91 | 2013.8491 |
| 04-2 | 14-16 | G(Dab)SDH(Iph)QTTTTKWKK | 97 | 2013.8669 |
| 04-2 | 14-16 | GNAHVQ(Dab)DP(Iph)LKWKK | 74 | 2016.9294 |
| 04-2 | 14-16 | GHSLTDATHN(Iph)KWKK | 76 | 2018.8723 |
| 04-2 | 14-16 | GNP(Pip)TDATHN(Iph)KWKK | 73 | 2018.8726 |
| 04-2 | 14-16 | GQTKSE(Dab)H(Dab)M(Iph)KWKK | 76 | 2083.9387 |
| 04-2 | 14-16 | GWENKDT(Cit)GG(Iph)KWKK | 76 | 2086.8987 |
| 04-2 | 14-16 | GNFHNKDT(Cit)N(Iph)KWKK | 96 | 2169.947 |
| 05-2 | 16-18 | GTLTDVTSGVGKWKK | 72 | 1699.937 |
| 05-2 | 16-18 | GT(Pip)S(Dab)TEAG(Dab)SKWKK | 79 | 1728.9385 |
| 05-2 | 16-18 | GT(Cit)AC(Dab)GAEGHKWKK | 90 | 1752.8955 |
| 05-2 | 16-18 | G(Pip)AGKTFGGENKWKK | 89 | 1756.9485 |
| 05-2 | 16-18 | GGRRSAGMESAKWKK | 96 | 1771.9377 |
| 05-2 | 16-18 | GQL(Cit)DVTSGVGKWKK | 82 | 1782.9854 |
| 05-2 | 16-18 | GS(Dab)STAE(Dab)V(Cit)NKWKK | 93 | 1814.9863 |
| 05-2 | 16-18 | GFTDSP(Dab)FAGKKWKK | 86 | 1819.9846 |

| | | | | |
|---|---|---|---|---|
| 05-2 | 16-18 | GFTSPQSFAGKKWKK | 72 | 1819.9846 |
| 05-2 | 16-18 | GDMTA(Dab)AY(Dab)DSKWKK | 86 | 1823.9102 |
| 05-2 | 16-18 | GKEPDSG(Pip)DTKKWKK | 83 | 1852.9907 |
| 05-2 | 16-18 | G(Dab)(Cit)PDSG(Pip)MGRKWKK | 81 | 1852.9956 |
| 05-2 | 16-18 | GLNKDDGK(Dab)DVKWKK | 94 | 1854.0225 |
| 05-2 | 16-18 | GVVAKSYANQ(Pip)KWKK | 73 | 1856.0532 |
| 05-2 | 16-18 | GQHKLC(Nap)GAAVKWKK | 74 | 1874.0249 |
| 05-2 | 16-18 | GKAEHKTDVAEKWKK | 80 | 1878.0225 |
| 05-2 | 16-18 | G(Dab)TSD(Nap)(Pip)(Cit)NGGKWKK | 71 | 1880.9758 |
| 05-2 | 16-18 | GGEKPTNNRPQKWKK | 74 | 1891.0288 |
| 05-2 | 16-18 | GDPQMV(Dab)G(Cit)NKKWKK | 89 | 1896.0266 |
| 05-2 | 16-18 | GRNSTCP(Dab)V(Cit)NKWKK | 87 | 1898.0171 |
| 05-2 | 16-18 | GRNSTAE(Dab)V(Cit)NKWKK | 90 | 1898.0347 |
| 05-2 | 16-18 | GHEVG(Cit)SSRNNKWKK | 70 | 1906.9988 |
| 05-2 | 16-18 | GAS(Cit)FMQAHQSKWKK | 81 | 1913.9795 |
| 05-2 | 16-18 | GAS(Cit)FC(Pip)THQSKWKK | 81 | 1913.9797 |
| 05-2 | 16-18 | GAS(Cit)(Pal)C(Pip)(Dab)HQSKWKK | 80 | 1913.9907 |
| 05-2 | 16-18 | GAS(Cit)FET(Dab)HQSKWKK | 87 | 1913.9973 |
| 05-2 | 16-18 | GNT(Pal)HANGH(Nap)AKWKK | 70 | 1916.9658 |
| 05-2 | 16-18 | GNTRCPNGH(Nap)AKWKK | 81 | 1916.9692 |
| 05-2 | 16-18 | GVD(Cit)EANGH(Nap)AKWKK | 86 | 1916.9758 |
| 05-2 | 16-18 | GNTREANGH(Nap)AKWKK | 90 | 1916.9871 |
| 05-2 | 16-18 | GNGGEQH(Nap)SS(Cit)KWKK | 88 | 1919.9502 |
| 05-2 | 16-18 | G(Cit)K(Cit)S(Cit)A(Cit)MGGKWKK | 74 | 1929.0596 |
| 05-2 | 16-18 | GFDKTRDNKSAKWKK | 85 | 1932.0442 |
| 05-2 | 16-18 | GVQAMSRVE(Cit)HKWKK | 81 | 1964.064 |
| 05-2 | 16-18 | GVQ(Dab)NN(Cit)TC(Pip)(Nap)KWKK | 84 | 2009.0532 |
| 05-2 | 16-18 | GVQ(Dab)NKDEE(Dab)(Nap)KWKK | 77 | 2009.0593 |
| 05-2 | 16-18 | GVQ(Dab)NN(Cit)TE(Dab)(Nap)KWKK | 89 | 2009.0708 |
| 05-2 | 16-18 | GHTER(Iph)LQGNAKWKK | 80 | 2048.9307 |
| 05-2 | 16-18 | GNPRERDFEKDKWKK | 79 | 2056.0715 |

| | | | | |
|---|---|---|---|---|
| 05-2 | 16-18 | G(Nap)SE(Cit)DQHTTHKWKK | 79 | 2059.0137 |
| 05-2 | 16-18 | GQTKSE(Dab)Q(Pip)N(Iph)KWKK | 81 | 2083.9565 |
| 05-2 | 16-18 | GKS(Pip)SN(Cit)TE(Iph)QKWKK | 85 | 2099.9514 |
| 05-2 | 16-18 | GKSRGN(Cit)TE(Iph)QKWKK | 95 | 2099.9626 |
| 05-2 | 16-18 | GTEDKE(Iph)T(Cit)RGKWKK | 79 | 2115.9463 |
| 06-2 | 18-21 | GPSERVSGVGVKWKK | 92 | 1736.9797 |
| 06-2 | 18-21 | GPS(Cit)QVSGVGVKWKK | 73 | 1736.9800 |
| 06-2 | 18-21 | GASDGFP(Pip)QVGKWKK | 70 | 1753.9377 |
| 06-2 | 18-21 | GANGQTN(Dab)T(Dab)TKWKK | 83 | 1756.9443 |
| 06-2 | 18-21 | GEVGNQG(Pal)GKAKWKK | 73 | 1757.9436 |
| 06-2 | 18-21 | G(Dab)PALNM(Dab)EAAKWKK | 74 | 1766.9727 |
| 06-2 | 18-21 | GGHDSVYAAHAKWKK | 73 | 1777.9124 |
| 06-2 | 18-21 | G(Cit)VEA(Dab)SVTAPKWKK | 82 | 1781.0061 |
| 06-2 | 18-21 | GHVN(Dab)GLDALTKWKK | 83 | 1790.0063 |
| 06-2 | 18-21 | GHVN(Dab)GLDKG(Dab)KWKK | 95 | 1790.0176 |
| 06-2 | 18-21 | GN(Pip)SS(Dab)NGVQLKWKK | 80 | 1794.9966 |
| 06-2 | 18-21 | GPALE(Dab)T(Cit)GMAKWKK | 70 | 1796.9832 |
| 06-2 | 18-21 | GRPGTSFTGVQKWKK | 78 | 1799.9907 |
| 06-2 | 18-21 | GSRFAGSAF(Pip)SKWKK | 85 | 1805.9802 |
| 06-2 | 18-21 | GRFAANPA(Dab)MSKWKK | 86 | 1814.9839 |
| 06-2 | 18-21 | GQ(Dab)EVEVGT(Dab)LKWKK | 81 | 1825.0322 |
| 06-2 | 18-21 | GK(Pip)LVVACHKGKWKK | 75 | 1831.0879 |
| 06-2 | 18-21 | G(Pip)(Cit)SAGDMPH(Dab)KWKK | 71 | 1847.9690 |
| 06-2 | 18-21 | GSF(Dab)NAHV(Cit)NGKWKK | 90 | 1852.9922 |
| 06-2 | 18-21 | GVVAKSFSNP(Cit)KWKK | 84 | 1856.0532 |
| 06-2 | 18-21 | GGQKT(Dab)PAD(Nap)PKWKK | 87 | 1861.0110 |
| 06-2 | 18-21 | GG(Pip)AFANT(Cit)(Pip)QKWKK | 91 | 1868.0283 |
| 06-2 | 18-21 | GNSSLEASHMFKWKK | 73 | 1872.9417 |
| 06-2 | 18-21 | GRVVNRPPATQKWKK | 70 | 1888.1021 |
| 06-2 | 18-21 | GTLNS(Dab)NH(Cit)VDKWKK | 85 | 1907.0239 |
| 06-2 | 18-21 | G(Pip)TLGK(Cit)(Cit)TSLKWKK | 95 | 1910.1328 |

| | | | | |
|---|---|---|---|---|
| 06-2 | 18-21 | G(Cit)TA(Dab)TM(Cit)HMGKWKK | 72 | 1912.9990 |
| 06-2 | 18-21 | GNLNRQNFTTGKWKK | 85 | 1915.0288 |
| 06-2 | 18-21 | G(Pip)TNRQNFTTGKWKK | 85 | 1915.0291 |
| 06-2 | 18-21 | GKH(Nap)ADKSANPKWKK | 75 | 1915.0327 |
| 06-2 | 18-21 | GARNRQNFTTGKWKK | 94 | 1915.0400 |
| 06-2 | 18-21 | GV(Dab)QTHD(Cit)(Dab)NPKWKK | 92 | 1918.0398 |
| 06-2 | 18-21 | G(Cit)TSSF(Pip)KMANKWKK | 84 | 1919.0312 |
| 06-2 | 18-21 | G(Cit)TSSF(Pip)KDT(Dab)KWKK | 87 | 1919.0491 |
| 06-2 | 18-21 | GNHN(Dab)PMTREAKWKK | 88 | 1920.0012 |
| 06-2 | 18-21 | G(Pip)FQFTTASHDKWKK | 72 | 1929.9963 |
| 06-2 | 18-21 | G(Pip)Q(Cit)HTTASHDKWKK | 89 | 1930.0034 |
| 06-2 | 18-21 | GNEN(Dab)L(Pip)TEAYKWKK | 89 | 1930.0173 |
| 06-2 | 18-21 | GSS(Pip)(Pip)F(Cit)TNSQKWKK | 74 | 1930.0288 |
| 06-2 | 18-21 | GKGTHEL(Cit)ENVKWKK | 82 | 1934.0601 |
| 06-2 | 18-21 | GKLSG(Pip)(Nap)(Cit)ANNKWKK | 83 | 1934.0752 |
| 06-2 | 18-21 | GRL(Nap)SRTNGAQKWKK | 74 | 1950.0813 |
| 06-2 | 18-21 | GT(Cit)(Cit)RPMKGTVKWKK | 75 | 1954.1162 |
| 06-2 | 18-21 | GKLNSH(Nap)VDPTKWKK | 88 | 1958.0637 |
| 06-2 | 18-21 | G(Dab)(Iph)GGQNEKLTKWKK | 77 | 1969.9136 |
| 06-2 | 18-21 | G(Dab)(Iph)GGQN(Cit)(Dab)LTKWKK | 95 | 1969.9248 |
| 06-2 | 18-21 | G(Dab)EKAFN(Cit)QHTKWKK | 93 | 1982.0710 |
| 06-2 | 18-21 | GTQKAFN(Cit)QHTKWKK | 86 | 1982.0713 |
| 06-2 | 18-21 | G(Nap)TSD(Dab)TDRRSKWKK | 70 | 1985.0342 |
| 06-2 | 18-21 | GH(Cit)VPTNKRNDKWKK | 84 | 1988.0930 |
| 06-2 | 18-21 | GVSELFLKT(Cit)(Cit)KWKK | 70 | 2001.1636 |
| 06-2 | 18-21 | G(Cit)F(Dab)QASFNGF(Iph)KWKK | 85 | 2050.9138 |
| 06-2 | 18-21 | GN(Dab)GPKK(Cit)VEF(Iph)KWKK | 75 | 2052.0032 |
| 06-2 | 18-21 | GQ(Cit)NV(Dab)HS(Iph)C(Pip)KWKK | 80 | 2093.9343 |
| 06-2 | 18-21 | GFEG(Cit)GR(Iph)(Dab)QEKWKK | 85 | 2102.9412 |
| 06-2 | 18-21 | GNQQH(Pal)(Iph)Q(Cit)G(Pip)KWKK | 78 | 2165.9634 |
| 06-2 | 18-21 | GNQQRE(Iph)Q(Cit)G(Pip)KWKK | 79 | 2165.9844 |

| | | | | |
|---|---|---|---|---|
| 07-2 | 21-24 | GGTTGLGGNRSKWKK | 70 | 1669.9126 |
| 07-2 | 21-24 | GEGL(Dab)(Pip)(Dab)SPASKWKK | 79 | 1736.9797 |
| 07-2 | 21-24 | GGGQPLANTH(Dab)KWKK | 83 | 1744.9597 |
| 07-2 | 21-24 | GHVN(Dab)GLDK(Dab)GKWKK | 73 | 1790.0176 |
| 07-2 | 21-24 | GE(Dab)NTVPE(Dab)GVKWKK | 82 | 1794.9854 |
| 07-2 | 21-24 | GKFGAFE(Dab)EAVKWKK | 93 | 1848.0159 |
| 07-2 | 21-24 | GKFGAFE(Dab)QA(Dab)KWKK | 96 | 1848.0271 |
| 07-2 | 21-24 | GKPDVNAYNGKKWKK | 87 | 1856.0168 |
| 07-2 | 21-24 | GVSGRNFGVFQKWKK | 83 | 1861.0225 |
| 07-2 | 21-24 | GVS(Dab)LTCGRFQKWKK | 72 | 1861.0256 |
| 07-2 | 21-24 | GKP(Dab)DLTG(Pip)(Pip)MKWKK | 90 | 1864.0618 |
| 07-2 | 21-24 | GKP(Dab)DLTG(Pip)KEKWKK | 89 | 1864.0796 |
| 07-2 | 21-24 | GTKENSHM(Dab)LGKWKK | 94 | 1867 |
| 07-2 | 21-24 | GQA(Cit)VPKVG(Cit)DKWKK | 79 | 1878.0701 |
| 07-2 | 21-24 | GSPFH(Cit)S(Dab)DGKKWKK | 77 | 1882.0076 |
| 07-2 | 21-24 | GGVSKNG(Cit)WEKKWKK | 75 | 1912.0544 |
| 07-2 | 21-24 | GTTP(Pip)TLFE(Dab)KKWKK | 87 | 1913.0999 |
| 07-2 | 21-24 | G(Dab)(Pip)DRQNFTTGKWKK | 82 | 1915.0291 |
| 07-2 | 21-24 | GNKMATE(Dab)(Cit)SHKWKK | 71 | 1925.0166 |
| 07-2 | 21-24 | GNNNK(Cit)ESQ(Dab)(Dab)KWKK | 79 | 1941.0405 |
| 07-2 | 21-24 | GKDNQ(Cit)HQNGVKWKK | 86 | 1947.03 |
| 07-2 | 21-24 | GREP(Pip)TLFE(Dab)KKWKK | 73 | 1996.1482 |
| 07-2 | 21-24 | GQ(Cit)P(Pip)TLFE(Dab)KKWKK | 85 | 1996.1484 |
| 07-2 | 21-24 | GNL(Pip)HQA(Nap)TD(Cit)KWKK | 72 | 2029.0759 |
| 07-2 | 21-24 | GPNG(Cit)Q(Cit)FS(Nap)KKWKK | 87 | 2039.0967 |
| 07-2 | 21-24 | G(Dab)WREDRDRGVKWKK | 82 | 2039.1038 |
| 07-2 | 21-24 | GPNG(Cit)Q(Cit)FHFKKWKK | 87 | 2039.1079 |
| 07-2 | 21-24 | GA(Dab)D(Iph)KRGDEKKWKK | 85 | 2041.9458 |
| 07-2 | 21-24 | GQ(Cit)DVVHS(Iph)QTKWKK | 71 | 2093.9409 |
| 07-2 | 21-24 | GQ(Cit)NV(Dab)HS(Iph)QTKWKK | 75 | 2093.9519 |
| 07-2 | 21-24 | GDTHHHGDKQ(Iph)KWKK | 88 | 2097.8894 |

| | | | | | |
|---|---|---|---|---|---|
| 07-2 | 21-24 | GAEDL(Cit)HVTH(Iph)KWKK | 88 | 2101.946 | |
| 08-2 | 24-28 | GDAT(Cit)S(Dab)GAR(Dab)KWKK | 70 | 1784.9871 | |
| 08-2 | 24-28 | GPSSHGVP(Pip)AYKWKK | 88 | 1790.9692 | |
| 08-2 | 24-28 | GV(Dab)PGF(Dab)VQNHKWKK | 81 | 1848.0383 | |
| 08-2 | 24-28 | G(Dab)TNSTVTLHKKWKK | 98 | 1851.0591 | |
| 08-2 | 24-28 | G(Dab)(Dab)DSTVTLHKKWKK | 75 | 1851.0591 | |
| 08-2 | 24-28 | GAGKP(Pip)(Cit)AKQMKWKK | 72 | 1864.073 | |
| 08-2 | 24-28 | GFRGKVSFTSSKWKK | 80 | 1866.0376 | |
| 08-2 | 24-28 | GHGHG(Nap)STSELKWKK | 79 | 1871.9543 | |
| 08-2 | 24-28 | G(Dab)CQG(Nap)STSKNKWKK | 75 | 1871.9575 | |
| 08-2 | 24-28 | GCER(Nap)STSKGGKWKK | 80 | 1871.9578 | |
| 08-2 | 24-28 | GELNGVY(Dab)TMTKWKK | 72 | 1877.9934 | |
| 08-2 | 24-28 | GNKNGVHKDDTKWKK | 84 | 1877.9973 | |
| 08-2 | 24-28 | GNKNGVY(Dab)TMTKWKK | 94 | 1878.0046 | |
| 08-2 | 24-28 | G(Pip)VTRSNLQPQKWKK | 77 | 1919.0967 | |
| 08-2 | 24-28 | GKPTRSNVLG(Nap)KWKK | 75 | 1919.1006 | |
| 08-2 | 24-28 | GKPTRSNPRVNKWKK | 80 | 1919.1077 | |
| 08-2 | 24-28 | GG(Dab)KEFNTTR(Pip)KWKK | 73 | 1929.0811 | |
| 08-2 | 24-28 | G(Cit)(Cit)(Cit)GQEVGV(Pip)KWKK | 74 | 1936.0869 | |
| 08-2 | 24-28 | G(Cit)(Dab)(Pip)NVELNHSKWKK | 83 | 1946.0713 | |
| 08-2 | 24-28 | G(Cit)(Dab)(Pip)N(Cit)ALNHSKWKK | 71 | 1946.0825 | |
| 08-2 | 24-28 | G(Nap)(Dab)HS(Cit)(Dab)S(Dab)DPKWK K | 83 | 1947.0339 | |
| 08-2 | 24-28 | G(Pip)NKTEL(Cit)NQMKWKK | 81 | 2011.0898 | |
| 08-2 | 24-28 | GKDKKTFRDQTKWKK | 87 | 2017.1333 | |
| 08-2 | 24-28 | GR(Cit)W(Pip)TQF(Dab)G(Dab)KWKK | 75 | 2028.1396 | |
| 08-2 | 24-28 | GQSKE(Dab)Q(Nap)(Cit)THKWKK | 81 | 2062.0972 | |
| 09-2 | 28-34 | G(Nap)AEPC(Cit)ELYVKWKK | 64 | 2028.0403 | |
| **10-2** | **34-60** | **G(Dab)VARNN(Dab)TKGKWKK** | **94** | **1810.0549** | **CXP3** |
| 10-2 | 34-60 | G(Pip)TSDKA(Pip)KA(Dab)KWKK | 74 | 1823.0642 | |
| 10-2 | 34-60 | GQRGA(Dab)YSGHTKWKK | 86 | 1826.9763 | |
| 10-2 | 34-60 | GY(Pip)AALLLTYTKWKK | 73 | 1905.0989 | |

| | | | | | |
|---|---|---|---|---|---|
| 10-2 | 34-60 | **GNFNNRS(Dab)GHKKWKK** | **94** | **1924.0405** | **CXP8** |
| 10-2 | 34-60 | **GARSEQHH(Nap)S(Dab)KWKK** | **90** | **1999.04** | **CXP9** |

**Library 3: 12,000-member library**

| Fraction Label | Fraction Retention Time (min) | | Sequencing confidence score (ALC) | peptide mass (Da) |
|---|---|---|---|---|
| 01-3 | 1-6 | GGVGQDQTGATKWKK | 71 | 1684 |
| 01-3 | 1-6 | GAQT(Dab)GAALDSKWKK | 79 | 1684 |
| 01-3 | 1-6 | GAGGVNFGDDPKWKK | 87 | 1699 |
| 01-3 | 1-6 | GTLPGVNEPAGKWKK | 92 | 1705 |
| 01-3 | 1-6 | GTTPDVNALSGKWKK | 92 | 1725 |
| 01-3 | 1-6 | GVSSGNLNSQAKWKK | 81 | 1727 |
| 01-3 | 1-6 | GFSQSSGGALEKWKK | 86 | 1733 |
| 01-3 | 1-6 | GSGVVMSAGFEKWKK | 90 | 1734 |
| 01-3 | 1-6 | GTTGETFEVGGKWKK | 91 | 1748 |
| 01-3 | 1-6 | GQANNAVAVPNKWKK | 97 | 1748 |
| 01-3 | 1-6 | GVASE(Dab)LGDNTKWKK | 88 | 1756 |
| 01-3 | 1-6 | GASSSTN(Cit)DNGKWKK | 77 | 1760 |
| 01-3 | 1-6 | GFVGTEQGLPAKWKK | 94 | 1769 |
| 01-3 | 1-6 | GNGAFGETEPVKWKK | 92 | 1771 |
| 01-3 | 1-6 | GNVAADQSTMSKWKK | 78 | 1774 |
| 01-3 | 1-6 | GVLSPT(Cit)GDADKWKK | 80 | 1782 |
| 01-3 | 1-6 | GSVAADPNDFPKWKK | 96 | 1783 |
| 01-3 | 1-6 | GGGAFDTKLLDKWKK | 90 | 1787 |
| 01-3 | 1-6 | GMQAVVGTAFDKWKK | 73 | 1789 |
| 01-3 | 1-6 | GSG(Cit)HDVAPEAKWKK | 79 | 1790 |
| 01-3 | 1-6 | GT(Cit)(Cit)AGPAALEKWKK | 77 | 1794 |
| 01-3 | 1-6 | GVVNMGVDTDPKWKK | 76 | 1797 |
| 01-3 | 1-6 | G(Dab)AT(Cit)ADVADEKWKK | 88 | 1799 |
| 01-3 | 1-6 | GKEAGNADMNTKWKK | 82 | 1801 |
| 01-3 | 1-6 | GLA(Cit)DTSTATNKWKK | 76 | 1801 |
| 01-3 | 1-6 | GGGNGFQPFNDKWKK | 97 | 1803 |
| 01-3 | 1-6 | GPVVETEAPLVKWKK | 82 | 1804 |
| 01-3 | 1-6 | GTDPLVEAPTNKWKK | 97 | 1807 |

| | | | | |
|---|---|---|---|---|
| 01-3 | 1-6 | GRPEDVNALSGKWKK | 83 | 1808 |
| 01-3 | 1-6 | GAQ(Dab)D(Cit)TGPESKWKK | 73 | 1812 |
| 01-3 | 1-6 | GK(Cit)SSG(Cit)PDASKWKK | 83 | 1813 |
| 01-3 | 1-6 | GTNPA(Cit)FLDAGKWKK | 83 | 1813 |
| 01-3 | 1-6 | G(Cit)VHAEQSDAGKWKK | 78 | 1821 |
| 01-3 | 1-6 | GGLGQFSTFSEKWKK | 74 | 1823 |
| 01-3 | 1-6 | GVQPDSLDLSVKWKK | 91 | 1823 |
| 01-3 | 1-6 | GSTVTAL(Cit)DPNKWKK | 71 | 1825 |
| 01-3 | 1-6 | G(Dab)GEQVQVDTVKWKK | 80 | 1825 |
| 01-3 | 1-6 | GESPPLPTQ(Cit)GKWKK | 93 | 1833 |
| 01-3 | 1-6 | GDE(Nap)PDVATGSKWKK | 85 | 1838 |
| 01-3 | 1-6 | GNEVVAPSLFNKWKK | 86 | 1840 |
| 01-3 | 1-6 | GADQGVE(Cit)TDVKWKK | 79 | 1841 |
| 01-3 | 1-6 | GNRGNVSSEDNKWKK | 80 | 1842 |
| 01-3 | 1-6 | G(Dab)SDDDNLQADKWKK | 82 | 1843 |
| 01-3 | 1-6 | GVNQNQTTSSNKWKK | 80 | 1843 |
| 01-3 | 1-6 | GLKAENDADMSKWKK | 82 | 1844 |
| 01-3 | 1-6 | GTDSTSQFQPSKWKK | 86 | 1848 |
| 01-3 | 1-6 | GHG(Cit)ENGEDGEKWKK | 94 | 1851 |
| 01-3 | 1-6 | G(Pip)EPAEVNDTTKWKK | 74 | 1852 |
| 01-3 | 1-6 | GDRGGEQ(Cit)LGLKWKK | 75 | 1852 |
| 01-3 | 1-6 | GDLLLPDSQDSKWKK | 92 | 1853 |
| 01-3 | 1-6 | GSNLLLSPEMVKWKK | 87 | 1853 |
| 01-3 | 1-6 | GGDKGTN(Nap)EDAKWKK | 92 | 1854 |
| 01-3 | 1-6 | GLATESENNPEKWKK | 86 | 1854 |
| 01-3 | 1-6 | G(Nap)VAADQSTMSKWKK | 75 | 1857 |
| 01-3 | 1-6 | GTPSFQLDDGEKWKK | 89 | 1859 |
| 01-3 | 1-6 | GEHAGGE(Nap)TTDKWKK | 86 | 1864 |
| 01-3 | 1-6 | GA(Dab)EQTENDDPKWKK | 79 | 1869 |
| 01-3 | 1-6 | GDQF(Cit)VETPGAKWKK | 89 | 1871 |
| 01-3 | 1-6 | GPSEDLNTTMNKWKK | 93 | 1872 |
| 01-3 | 1-6 | GKDGLDGV(Cit)FNKWKK | 89 | 1872 |
| 01-3 | 1-6 | GFTNSELTSPQKWKK | 97 | 1874 |
| 01-3 | 1-6 | GG(Dab)TQLAFTE(Cit)KWKK | 86 | 1874 |
| 01-3 | 1-6 | GHEVDQPDPSTKWKK | 77 | 1875 |
| 01-3 | 1-6 | GLNDLPTAQKEKWKK | 86 | 1879 |
| 01-3 | 1-6 | GTAENEVPVD(Cit)KWKK | 76 | 1881 |
| 01-3 | 1-6 | GDAQHDNEDSTKWKK | 88 | 1882 |
| 01-3 | 1-6 | GRPSDPQDMEGKWKK | 81 | 1882 |

| | | | | |
|---|---|---|---|---|
| 01-3 | 1-6 | GPTDLQTDLNDKWKK | 77 | 1882 |
| 01-3 | 1-6 | GEQSV(Cit)VVGQEKWKK | 85 | 1882 |
| 01-3 | 1-6 | GVAMQDQANNFKWKK | 95 | 1888 |
| 01-3 | 1-6 | GSD(Cit)SKQDTLSKWKK | 85 | 1888 |
| 01-3 | 1-6 | GPPANV(Nap)NESLKWKK | 84 | 1888 |
| 01-3 | 1-6 | GAQ(Nap)LVEAPTNKWKK | 83 | 1890 |
| 01-3 | 1-6 | GQVQENLAVNEKWKK | 84 | 1894 |
| 01-3 | 1-6 | GS(Nap)(Dab)PANNTEDKWKK | 72 | 1895 |
| 01-3 | 1-6 | GS(Nap)(Dab)PANNTEDKWKK | 80 | 1895 |
| 01-3 | 1-6 | GV(Cit)GD(Cit)PESLNKWKK | 83 | 1895 |
| 01-3 | 1-6 | GSKVL(Cit)VESNDKWKK | 90 | 1898 |
| 01-3 | 1-6 | GTAFG(Nap)D(Cit)VNAKWKK | 91 | 1899 |
| 01-3 | 1-6 | GTTSNNET(Cit)LNKWKK | 89 | 1901 |
| 01-3 | 1-6 | GMT(Iph)VSTLDGGKWKK | 93 | 1904 |
| 01-3 | 1-6 | GHPNS(Iph)GSGDDKWKK | 83 | 1909 |
| 01-3 | 1-6 | G(Cit)MAVASPD(Nap)NKWKK | 88 | 1909 |
| 01-3 | 1-6 | GE(Dab)E(Cit)TDG(Cit)PVKWKK | 93 | 1911 |
| 01-3 | 1-6 | GTTSLLVQE(Cit)LKWKK | 80 | 1911 |
| 01-3 | 1-6 | GVSKLFDHDNSKWKK | 85 | 1912 |
| 01-3 | 1-6 | GNAEPVFL(Cit)DTKWKK | 77 | 1913 |
| 01-3 | 1-6 | GAGMQ(Cit)Q(Cit)TDVKWKK | 92 | 1914 |
| 01-3 | 1-6 | GTQGSES(Cit)TT(Nap)KWKK | 74 | 1915 |
| 01-3 | 1-6 | GKQD(Dab)DM(Cit)DTGKWKK | 94 | 1917 |
| 01-3 | 1-6 | GVMKDPVFSDEKWKK | 87 | 1917 |
| 01-3 | 1-6 | GQLTG(Pip)EPED(Cit)KWKK | 79 | 1922 |
| 01-3 | 1-6 | G(Nap)RGNVSSEDNKWKK | 75 | 1925 |
| 01-3 | 1-6 | GTHFDDNLQADKWKK | 84 | 1926 |
| 01-3 | 1-6 | GTTRN(Cit)GQEDTKWKK | 79 | 1929 |
| 01-3 | 1-6 | GQQAFDP(Cit)NFGKWKK | 94 | 1931 |
| 01-3 | 1-6 | GQ(Pip)KSAFLMDDKWKK | 78 | 1931 |
| 01-3 | 1-6 | GVEDATQ(Cit)EQLKWKK | 78 | 1940 |
| 01-3 | 1-6 | GAVMVKEQN(Cit)DKWKK | 92 | 1941 |
| 01-3 | 1-6 | GANQ(Iph)LGDAETKWKK | 77 | 1942 |
| 01-3 | 1-6 | GV(Dab)VV(Iph)G(Dab)DDDKWKK | 75 | 1942 |
| 01-3 | 1-6 | GSDFQSQLDEQKWKK | 84 | 1947 |
| 01-3 | 1-6 | GNTPDQT(Iph)GVPKWKK | 94 | 1952 |
| 01-3 | 1-6 | GVEAS(Iph)PNQNAKWKK | 90 | 1953 |
| 01-3 | 1-6 | GF(Cit)LNDHEAMAKWKK | 79 | 1955 |
| 01-3 | 1-6 | GQTGLLG(Iph)ENTKWKK | 78 | 1956 |

| | | | | |
|---|---|---|---|---|
| 01-3 | 1-6 | GNVTS(Iph)GNE(Cit)GKWKK | 82 | 1958 |
| 01-3 | 1-6 | G(Cit)RMDDAADT(Cit)KWKK | 75 | 1959 |
| 01-3 | 1-6 | GT(Cit)FQAPDNNFKWKK | 81 | 1961 |
| 01-3 | 1-6 | GTQP(Nap)PTTQQDKWKK | 97 | 1963 |
| 01-3 | 1-6 | GQQQNEVPVD(Cit)KWKK | 81 | 1964 |
| 01-3 | 1-6 | GSEQE(Cit)KNDPDKWKK | 94 | 1969 |
| 01-3 | 1-6 | GVQ(Pip)NLTLDFFKWKK | 82 | 1973 |
| 01-3 | 1-6 | GDDHS(Nap)PNAD(Cit)KWKK | 79 | 1975 |
| 01-3 | 1-6 | GHDVNASPV(Iph)DKWKK | 77 | 1977 |
| 01-3 | 1-6 | GN(Iph)SLPPA(Cit)EAKWKK | 88 | 1979 |
| 01-3 | 1-6 | GADKSQFFRDDKWKK | 73 | 1979 |
| 01-3 | 1-6 | GDTG(Cit)(Cit)(Nap)EPTVKWKK | 84 | 1980 |
| 01-3 | 1-6 | GFQQLP(Cit)EVNTKWKK | 88 | 1983 |
| 01-3 | 1-6 | G(Cit)QSNNET(Cit)LNKWKK | 73 | 1984 |
| 01-3 | 1-6 | GASSE(Nap)NE(Cit)EDKWKK | 83 | 1985 |
| 01-3 | 1-6 | GVTMGQ(Iph)DPQSKWKK | 90 | 1986 |
| 01-3 | 1-6 | GSA(Cit)(Iph)VSTLDNKWKK | 75 | 1987 |
| 01-3 | 1-6 | GET(Cit)A(Nap)FDDANKWKK | 90 | 1987 |
| 01-3 | 1-6 | GTGLSQ(Iph)AFDQKWKK | 95 | 1990 |
| 01-3 | 1-6 | GSF(Nap)EQDAEFAKWKK | 95 | 1991 |
| 01-3 | 1-6 | GPNSHTD(Iph)GENKWKK | 84 | 1994 |
| 01-3 | 1-6 | G(Pip)TDQ(Cit)Q(Cit)TDVKWKK | 75 | 1997 |
| 01-3 | 1-6 | G(Pip)EDPTSQA(Iph)TKWKK | 79 | 1998 |
| 01-3 | 1-6 | GASN(Iph)TDNF(Cit)GKWKK | 80 | 2006 |
| 01-3 | 1-6 | G(Cit)DQTSDAVP(Iph)KWKK | 89 | 2013 |
| 01-3 | 1-6 | GEPTPSQNSM(Iph)KWKK | 82 | 2014 |
| 01-3 | 1-6 | G(Cit)LTF(Cit)KEPDTKWKK | 72 | 2015 |
| 01-3 | 1-6 | GLVEN(Cit)HQQQDKWKK | 80 | 2018 |
| 01-3 | 1-6 | GHNDNE(Nap)TNDLKWKK | 79 | 2019 |
| 01-3 | 1-6 | GQQL(Cit)DT(Cit)P(Nap)GKWKK | 94 | 2020 |
| 01-3 | 1-6 | GAK(Nap)QDPT(Cit)MEKWKK | 90 | 2024 |
| 01-3 | 1-6 | GKPGVV(Iph)(Cit)DDDKWKK | 85 | 2025 |
| 01-3 | 1-6 | G(Cit)PDQVPE(Iph)LGKWKK | 94 | 2035 |
| 01-3 | 1-6 | GMEQ(Nap)LTDQPEKWKK | 74 | 2038 |
| 01-3 | 1-6 | GTDHT(Iph)DPTNDKWKK | 77 | 2039 |
| 01-3 | 1-6 | G(Cit)LNRNTE(Cit)DLKWKK | 87 | 2039 |
| 01-3 | 1-6 | GNNP(Cit)SVQD(Iph)SKWKK | 82 | 2041 |
| 01-3 | 1-6 | GP(Nap)TESRNDFEKWKK | 74 | 2042 |
| 01-3 | 1-6 | GF(Cit)VPTNF(Cit)EEKWKK | 71 | 2047 |

| | | | | |
|---|---|---|---|---|
| 01-3 | 1-6 | GKD(Iph)ANDTFSQKWKK | 89 | 2049 |
| 01-3 | 1-6 | GPVQE(Iph)DPPEQKWKK | 79 | 2062 |
| 01-3 | 1-6 | GG(Iph)(Cit)SQETD(Cit)LKWKK | 85 | 2087 |
| 01-3 | 1-6 | GADP(Iph)RQNEQLKWKK | 90 | 2094 |
| 01-3 | 1-6 | GRDEF(Cit)N(Cit)EFDKWKK | 79 | 2136 |
| 02-3 | 6-10 | GP(Dab)GFGGAEAAKWKK | 97 | 1627 |
| 02-3 | 6-10 | GTFVA(Dab)FGDGGKWKK | 72 | 1721 |
| 02-3 | 6-10 | GADLDVNALSGKWKK | 84 | 1725 |
| 02-3 | 6-10 | GD(Cit)SG(Dab)GFGSNKWKK | 77 | 1748 |
| 02-3 | 6-10 | GVSMGVAFTSTKWKK | 72 | 1750 |
| 02-3 | 6-10 | GALAQFANLGVKWKK | 75 | 1754 |
| 02-3 | 6-10 | GNQSQPTTGTAKWKK | 88 | 1755 |
| 02-3 | 6-10 | GPNT(Dab)NFAGADKWKK | 77 | 1757 |
| 02-3 | 6-10 | GFVGTEQGLPAKWKK | 74 | 1769 |
| 02-3 | 6-10 | GDP(Pip)GATHGEDKWKK | 77 | 1775 |
| 02-3 | 6-10 | GSNQHGDQLGAKWKK | 81 | 1777 |
| 02-3 | 6-10 | GHNSDNV(Dab)GADKWKK | 94 | 1779 |
| 02-3 | 6-10 | GTLVSP(Nap)GSATKWKK | 73 | 1780 |
| 02-3 | 6-10 | GVLVRSAGGD(Cit)KWKK | 86 | 1781 |
| 02-3 | 6-10 | GM(Dab)FGNPSADTKWKK | 91 | 1790 |
| 02-3 | 6-10 | GSKTSSAEQVNKWKK | 92 | 1801 |
| 02-3 | 6-10 | G(Pal)ALSAQSVLNKWKK | 72 | 1801 |
| 02-3 | 6-10 | GRPEDVNALSGKWKK | 88 | 1808 |
| 02-3 | 6-10 | GRSVVDLLSNGKWKK | 94 | 1810 |
| 02-3 | 6-10 | GK(Cit)SSGQ(Pip)DASKWKK | 94 | 1813 |
| 02-3 | 6-10 | GMSKTDSQVPAKWKK | 93 | 1814 |
| 02-3 | 6-10 | GPKPQAVTTPEKWKK | 87 | 1818 |
| 02-3 | 6-10 | G(Nap)SAP(Cit)GL(Cit)GGKWKK | 89 | 1820 |
| 02-3 | 6-10 | GSA(Nap)S(Cit)GFGSNKWKK | 96 | 1831 |
| 02-3 | 6-10 | G(Dab)GAVQNGF(Cit)MKWKK | 93 | 1831 |
| 02-3 | 6-10 | GGHVSEVP(Cit)GFKWKK | 75 | 1836 |
| 02-3 | 6-10 | G(Cit)NNSQVSGA(Cit)KWKK | 78 | 1841 |
| 02-3 | 6-10 | GVNQNQTTSSNKWKK | 94 | 1843 |
| 02-3 | 6-10 | GSKETESNTVTKWKK | 92 | 1846 |
| 02-3 | 6-10 | GPNSHNGEA(Cit)NKWKK | 82 | 1847 |
| 02-3 | 6-10 | GDG(Cit)(Pip)VT(Cit)VAPKWKK | 77 | 1849 |
| 02-3 | 6-10 | GVGPQ(Dab)(Cit)EQGQKWKK | 71 | 1850 |
| 02-3 | 6-10 | GGA(Cit)DAREQPTKWKK | 77 | 1852 |
| 02-3 | 6-10 | GDRGGEQ(Cit)LGLKWKK | 94 | 1852 |

| | | | | |
|---|---|---|---|---|
| 02-3 | 6-10 | GKDQGVG(Cit)DDLKWKK | 82 | 1854 |
| 02-3 | 6-10 | GG(Dab)(Iph)DVGGG(Cit)NKWKK | 84 | 1856 |
| 02-3 | 6-10 | GNFS(Cit)SAN(Dab)PDKWKK | 79 | 1859 |
| 02-3 | 6-10 | GDQ(Cit)S(Dab)NFVAAKWKK | 83 | 1859 |
| 02-3 | 6-10 | GNG(Dab)VNFEEFGKWKK | 84 | 1863 |
| 02-3 | 6-10 | GS(Dab)SVFDQMSLKWKK | 96 | 1864 |
| 02-3 | 6-10 | GKESPPQT(Cit)AVKWKK | 85 | 1864 |
| 02-3 | 6-10 | GQPESENRSPAKWKK | 78 | 1865 |
| 02-3 | 6-10 | GK(Cit)EANNEPSAKWKK | 81 | 1867 |
| 02-3 | 6-10 | GGTKNVQDPNFKWKK | 87 | 1870 |
| 02-3 | 6-10 | G(Dab)ATA(Cit)SLE(Cit)NKWKK | 96 | 1870 |
| 02-3 | 6-10 | G(Dab)ATA(Cit)SLE(Cit)NKWKK | 96 | 1870 |
| 02-3 | 6-10 | GKDGLDGV(Cit)FNKWKK | 96 | 1872 |
| 02-3 | 6-10 | GVLTNTNSKMDKWKK | 89 | 1873 |
| 02-3 | 6-10 | GKEGT(Cit)NTFLGKWKK | 86 | 1874 |
| 02-3 | 6-10 | GFFP(Dab)ENAQTAKWKK | 79 | 1875 |
| 02-3 | 6-10 | GPTHAAEKEDMKWKK | 89 | 1879 |
| 02-3 | 6-10 | GLNDLPTAQKEKWKK | 76 | 1879 |
| 02-3 | 6-10 | GP(Cit)GQTPTQS(Cit)KWKK | 77 | 1880 |
| 02-3 | 6-10 | GFNE(Pip)AVVPTEKWKK | 88 | 1882 |
| 02-3 | 6-10 | G(Cit)ST(Cit)GKDPNTKWKK | 81 | 1884 |
| 02-3 | 6-10 | G(Pal)VLVQNSNQSKWKK | 75 | 1887 |
| 02-3 | 6-10 | GVGHDPQA(Cit)V(Cit)KWKK | 84 | 1887 |
| 02-3 | 6-10 | GAQ(Nap)LVEAPTNKWKK | 75 | 1890 |
| 02-3 | 6-10 | GA(Cit)DKLDPFALKWKK | 80 | 1897 |
| 02-3 | 6-10 | GND(Dab)HAAFES(Cit)KWKK | 78 | 1898 |
| 02-3 | 6-10 | G(Iph)AQA(Cit)SAGTLKWKK | 92 | 1899 |
| 02-3 | 6-10 | G(Cit)NGGDTSM(Nap)NKWKK | 85 | 1900 |
| 02-3 | 6-10 | GST(Dab)LFDPN(Cit)VKWKK | 84 | 1900 |
| 02-3 | 6-10 | GAFTL(Cit)SLNGLWWKK | 75 | 1901 |
| 02-3 | 6-10 | GNE(Cit)PVPDQKAKWKK | 84 | 1905 |
| 02-3 | 6-10 | GGVVFQQ(Nap)QPGKWKK | 89 | 1907 |
| 02-3 | 6-10 | G(Cit)RAGPQPDLFKWKK | 87 | 1908 |
| 02-3 | 6-10 | GNRSVMLDNNPKWKK | 83 | 1910 |
| 02-3 | 6-10 | GKPSVFMLEPNKWKK | 84 | 1912 |
| 02-3 | 6-10 | GNHQS(Cit)QEALSKWKK | 76 | 1921 |
| 02-3 | 6-10 | GLLV(Cit)(Cit)TSQTPKWKK | 78 | 1923 |
| 02-3 | 6-10 | G(Nap)RG(Pip)SSSEDNKWKK | 84 | 1925 |
| 02-3 | 6-10 | GN(Cit)FQ(Dab)DPSLTKWKK | 82 | 1929 |

| | | | | |
|---|---|---|---|---|
| 02-3 | 6-10 | GNA(Pal)FDTKLLDKWKK | 92 | 1935 |
| 02-3 | 6-10 | GNNA(Cit)(Nap)Q(Dab)GLDKWKK | 90 | 1936 |
| 02-3 | 6-10 | GTKTVNQEGD(Nap)KWKK | 85 | 1939 |
| 02-3 | 6-10 | GLQLTE(Cit)VDTNKWKK | 81 | 1940 |
| 02-3 | 6-10 | GV(Cit)GHH(Pip)D(Cit)TSKWKK | 72 | 1943 |
| 02-3 | 6-10 | GEAP(Nap)DKGNFQKWKK | 72 | 1953 |
| 02-3 | 6-10 | GKLATMSD(Nap)NQKWKK | 97 | 1955 |
| 02-3 | 6-10 | GTNLGERD(Cit)S(Cit)KWKK | 72 | 1956 |
| 02-3 | 6-10 | GV(Dab)GDNSTQL(Iph)KWKK | 79 | 1957 |
| 02-3 | 6-10 | GST(Cit)KG(Iph)DQAAKWKK | 81 | 1958 |
| 02-3 | 6-10 | GAKTD(Iph)TPTMAKWKK | 74 | 1959 |
| 02-3 | 6-10 | GNFF(Pip)(Cit)ATDNVKWKK | 86 | 1961 |
| 02-3 | 6-10 | GVKEDPFLRDPKWKK | 91 | 1966 |
| 02-3 | 6-10 | GR(Cit)V(Cit)MLDNSAKWKK | 83 | 1970 |
| 02-3 | 6-10 | GS(Pal)GMP(Iph)KDTAKWKK | 77 | 1978 |
| 02-3 | 6-10 | GRVFLNPPL(Cit)DKWKK | 92 | 1978 |
| 02-3 | 6-10 | GVSQSQFFRDDKWKK | 83 | 1979 |
| 02-3 | 6-10 | GRG(Cit)DP(Iph)AEGVKWKK | 88 | 1981 |
| 02-3 | 6-10 | GA(Cit)HLMQFPETKWKK | 79 | 1981 |
| 02-3 | 6-10 | GAEAARV(Iph)LDNKWKK | 79 | 1982 |
| 02-3 | 6-10 | G(Cit)N(Dab)LFDNP(Cit)VKWKK | 81 | 1983 |
| 02-3 | 6-10 | G(Cit)(Dab)GE(Dab)DL(Pal)(Pal)FKWKK | 75 | 1984 |
| 02-3 | 6-10 | GD(Dab)N(Nap)AMKS(Cit)DKWKK | 72 | 1985 |
| 02-3 | 6-10 | G(Nap)EP(Cit)VPDQKAKWKK | 78 | 1988 |
| 02-3 | 6-10 | G(Cit)KLFEFSSVEKWKK | 95 | 1993 |
| 02-3 | 6-10 | G(Nap)KNESSTA(Nap)DKWKK | 80 | 1996 |
| 02-3 | 6-10 | GPTNHDPSP(Iph)NKWKK | 72 | 2002 |
| 02-3 | 6-10 | GQTQELRRNVDKWKK | 72 | 2009 |
| 02-3 | 6-10 | GVV(Cit)VP(Iph)(Cit)GTTKWKK | 97 | 2010 |
| 02-3 | 6-10 | G(Nap)T(Cit)GPQAFQ(Cit)KWKK | 89 | 2010 |
| 02-3 | 6-10 | G(Nap)(Cit)FQ(Dab)DPSLTKWKK | 86 | 2012 |
| 02-3 | 6-10 | GKPSD(Iph)TNTNNKWKK | 85 | 2014 |
| 02-3 | 6-10 | GL(Cit)(Iph)VQATPPLKWKK | 77 | 2019 |
| 02-3 | 6-10 | GAESVEREQ(Pip)(Nap)KWKK | 80 | 2021 |
| 02-3 | 6-10 | GHVTQEE(Cit)LLFKWKK | 92 | 2023 |
| 02-3 | 6-10 | G(Iph)G(Cit)GRVV(Cit)DSKWKK | 81 | 2027 |
| 02-3 | 6-10 | GEGKTVD(Iph)LPFKWKK | 76 | 2029 |
| 02-3 | 6-10 | GA(Iph)VSRNDMTNKWKK | 84 | 2031 |
| 02-3 | 6-10 | GVPDR(Cit)TL(Cit)(Cit)NKWKK | 91 | 2036 |

| | | | | |
|---|---|---|---|---|
| 02-3 | 6-10 | GQHDSFQG(Cit)(Cit)(Cit)KWKK | 92 | 2040 |
| 02-3 | 6-10 | GVRQD(Iph)TPTMAKWKK | 89 | 2042 |
| 02-3 | 6-10 | GEFVNN(Iph)KDTAKWKK | 88 | 2061 |
| 02-3 | 6-10 | GPG(Cit)NRV(Iph)LDNKWKK | 89 | 2065 |
| 02-3 | 6-10 | GADP(Iph)RQNEQLKWKK | 88 | 2094 |
| 02-3 | 6-10 | GQDQEHV(Cit)F(Nap)PKWKK | 78 | 2104 |
| 02-3 | 6-10 | GKQQVQT(Iph)QDNKWKK | 90 | 2112 |
| 02-3 | 6-10 | GTTEP(Pal)Q(Cit)(Iph)QLKWKK | 78 | 2145 |
| 04-3 | 13-15 | GEKVGGAGHAGKWKK | 72 | 1633 |
| 04-3 | 13-15 | G(Pip)SGLFPGSGAKWKK | 88 | 1669 |
| 04-3 | 13-15 | GRAAGALT(Dab)VDKWKK | 92 | 1724 |
| 04-3 | 13-15 | GVGHNGKVDVGKWKK | 98 | 1732 |
| 04-3 | 13-15 | GFS(Dab)TDAGTGHKWKK | 77 | 1743 |
| 04-3 | 13-15 | GGMGAD(Nap)GVEAKWKK | 83 | 1754 |
| 04-3 | 13-15 | GN(Pip)PVVGGFPTKWKK | 80 | 1764 |
| 04-3 | 13-15 | GS(Cit)GPGNTTKVKWKK | 87 | 1768 |
| 04-3 | 13-15 | GAFVLAKSQGVKWKK | 79 | 1770 |
| 04-3 | 13-15 | GFTGTPG(Cit)GNLKWKK | 72 | 1771 |
| 04-3 | 13-15 | GSTSPSVTKLNKWKK | 91 | 1784 |
| 04-3 | 13-15 | G(Dab)GGLNKMAMDKWKK | 72 | 1787 |
| 04-3 | 13-15 | GA(Dab)VN(Nap)GAV(Cit)GKWKK | 86 | 1792 |
| 04-3 | 13-15 | GAR(Pip)DNGSSPNKWKK | 71 | 1794 |
| 04-3 | 13-15 | GE(Pip)SEFGGPQAKWKK | 76 | 1798 |
| 04-3 | 13-15 | G(Dab)S(Nap)APSGKDPKWKK | 81 | 1806 |
| 04-3 | 13-15 | G(Cit)SSNATHGQVKWKK | 90 | 1808 |
| 04-3 | 13-15 | GVESHGKLVTPKWKK | 89 | 1817 |
| 04-3 | 13-15 | GANNNKKS(Cit)AAKWKK | 82 | 1825 |
| 04-3 | 13-15 | GAALVAP(Cit)QHLKWKK | 75 | 1827 |
| 04-3 | 13-15 | GD(Pip)APQ(Cit)HVGAKWKK | 88 | 1828 |
| 04-3 | 13-15 | GSKG(Dab)FLKVDSKWKK | 82 | 1831 |
| 04-3 | 13-15 | GH(Cit)PTAA(Dab)AQEKWKK | 93 | 1832 |
| 04-3 | 13-15 | G(Pip)GD(Cit)EAPKALKWKK | 92 | 1834 |
| 04-3 | 13-15 | GH(Dab)TASLTNQDKWKK | 89 | 1837 |
| 04-3 | 13-15 | GKGGFHKSLNVKWKK | 80 | 1837 |
| 04-3 | 13-15 | G(Pal)NGHAANNRSKWKK | 72 | 1839 |
| 04-3 | 13-15 | GAL(Iph)SKGA(Cit)GGKWKK | 73 | 1841 |
| 04-3 | 13-15 | G(Pip)FSG(Cit)A(Cit)PTAKWKK | 78 | 1841 |
| 04-3 | 13-15 | GNA(Dab)LQTEHNAKWKK | 80 | 1848 |
| 04-3 | 13-15 | G(Dab)(Pip)PLSDGL(Cit)NKWKK | 87 | 1849 |

| | | | | |
|---|---|---|---|---|
| 04-3 | 13-15 | GHVTSREVATTKWKK | 98 | 1851 |
| 04-3 | 13-15 | GGFKG(Nap)G(Cit)GNLKWKK | 96 | 1854 |
| 04-3 | 13-15 | GPVHGHFNEAPKWKK | 94 | 1855 |
| 04-3 | 13-15 | GQGLVFRLGQSKWKK | 77 | 1855 |
| 04-3 | 13-15 | GNS(Dab)FVVELPTKWKK | 88 | 1856 |
| 04-3 | 13-15 | GVG(Iph)D(Dab)EAAATKWKK | 83 | 1857 |
| 04-3 | 13-15 | G(Dab)GTVLTMRDNKWKK | 89 | 1857 |
| 04-3 | 13-15 | GVVPHVADNQKKWKK | 89 | 1857 |
| 04-3 | 13-15 | GHEDAH(Dab)VENGKWKK | 85 | 1858 |
| 04-3 | 13-15 | GSA(Pal)VA(Cit)N(Dab)PFKWKK | 77 | 1861 |
| 04-3 | 13-15 | GVH(Cit)ATARDSTKWKK | 91 | 1865 |
| 04-3 | 13-15 | GRDSPSVTKLNKWKK | 93 | 1867 |
| 04-3 | 13-15 | GP(Iph)LSSAKGNAKWKK | 74 | 1868 |
| 04-3 | 13-15 | GRSAVRPFGDLKWKK | 91 | 1868 |
| 04-3 | 13-15 | GAALAQFRFVTKWKK | 91 | 1874 |
| 04-3 | 13-15 | GHNSMSHEVPSKWKK | 72 | 1875 |
| 04-3 | 13-15 | G(Dab)QDTFRSLTGKWKK | 94 | 1875 |
| 04-3 | 13-15 | G(Dab)FDTG(Pip)NVTFKWKK | 77 | 1877 |
| 04-3 | 13-15 | GVM(Dab)EDPPRPSKWKK | 84 | 1878 |
| 04-3 | 13-15 | GQ(Pip)(Dab)GSNTS(Nap)LKWKK | 73 | 1880 |
| 04-3 | 13-15 | GTH(Dab)SEFNPQAKWKK | 90 | 1881 |
| 04-3 | 13-15 | GKAQ(Cit)DG(Dab)NLQKWKK | 85 | 1881 |
| 04-3 | 13-15 | GNRVLKGMGEEKWKK | 95 | 1883 |
| 04-3 | 13-15 | GLP(Pal)KATQNTNKWKK | 77 | 1885 |
| 04-3 | 13-15 | GTDNSHHALNQKWKK | 85 | 1887 |
| 04-3 | 13-15 | GLVFEPTFA(Dab)NKWKK | 78 | 1888 |
| 04-3 | 13-15 | GRN(Nap)APSGKDPKWKK | 89 | 1889 |
| 04-3 | 13-15 | GMDAHSTSAG(Iph)KWKK | 72 | 1900 |
| 04-3 | 13-15 | GKEEPPHSFSPKWKK | 83 | 1905 |
| 04-3 | 13-15 | G(Cit)TGHDTFGQHKWKK | 82 | 1907 |
| 04-3 | 13-15 | GNTGS(Nap)N(Nap)GQSKWKK | 75 | 1909 |
| 04-3 | 13-15 | GE(Pip)P(Nap)GGNDFTKWKK | 79 | 1910 |
| 04-3 | 13-15 | GVFEES(Pip)GETHKWKK | 89 | 1911 |
| 04-3 | 13-15 | GAFLRG(Cit)N(Cit)GDKWKK | 73 | 1914 |
| 04-3 | 13-15 | GKT(Cit)PSGRDFVKWKK | 87 | 1914 |
| 04-3 | 13-15 | GAFQPSK(Cit)AMQKWKK | 76 | 1915 |
| 04-3 | 13-15 | GFTTVQKQDKAKWKK | 92 | 1916 |
| 04-3 | 13-15 | GMKNEDGNQM(Dab)KWKK | 90 | 1917 |
| 04-3 | 13-15 | GHTNADNTF(Dab)FKWKK | 95 | 1917 |

| | | | | |
|---|---|---|---|---|
| 04-3 | 13-15 | G(Dab)PEHV(Nap)GFANKWKK | 74 | 1918 |
| 04-3 | 13-15 | GPAVVVFHN(Cit)QKWKK | 87 | 1918 |
| 04-3 | 13-15 | G(Dab)VSTEHEQQNKWKK | 91 | 1922 |
| 04-3 | 13-15 | GNFTDEQRAHGKWKK | 83 | 1925 |
| 04-3 | 13-15 | GAG(Pip)HMDFN(Cit)(Dab)KWKK | 73 | 1925 |
| 04-3 | 13-15 | GVAFHTDMNLKKWKK | 84 | 1926 |
| 04-3 | 13-15 | GPTKTRSADF(Cit)KWKK | 91 | 1930 |
| 04-3 | 13-15 | GPEQQDKTAEHKWKK | 74 | 1933 |
| 04-3 | 13-15 | GHLEGQQFGDRKWKK | 89 | 1937 |
| 04-3 | 13-15 | GQPLLEHHEALKWKK | 74 | 1937 |
| 04-3 | 13-15 | GMT(Dab)PTMHQLEKWKK | 79 | 1938 |
| 04-3 | 13-15 | GFARE(Cit)DHVSAKWKK | 97 | 1939 |
| 04-3 | 13-15 | GAQ(Dab)THFPQFDKWKK | 87 | 1941 |
| 04-3 | 13-15 | GREAS(Cit)SKTQEKWKK | 91 | 1943 |
| 04-3 | 13-15 | GELFVA(Cit)N(Dab)PFKWKK | 73 | 1944 |
| 04-3 | 13-15 | GN(Nap)G(Pip)SDPREPKWKK | 88 | 1945 |
| 04-3 | 13-15 | GQ(Dab)AN(Nap)PTMDKKWKK | 92 | 1952 |
| 04-3 | 13-15 | G(Cit)AQHPPG(Iph)GNKWKK | 77 | 1958 |
| 04-3 | 13-15 | GDPVHG(Iph)KEPGKWKK | 71 | 1959 |
| 04-3 | 13-15 | GP(Nap)PAENEETKKWKK | 79 | 1962 |
| 04-3 | 13-15 | GR(Iph)GAANNLQTKWKK | 83 | 1968 |
| 04-3 | 13-15 | G(Cit)RDA(Cit)PNHVPKWKK | 85 | 1970 |
| 04-3 | 13-15 | GK(Pal)TGGSHG(Iph)(Cit)KWKK | 79 | 1972 |
| 04-3 | 13-15 | G(Cit)PNHQELGDRKWKK | 88 | 1973 |
| 04-3 | 13-15 | GHFN(Nap)DHGVSNKWKK | 90 | 1974 |
| 04-3 | 13-15 | GESQKDFTRTNKWKK | 88 | 1976 |
| 04-3 | 13-15 | GAHSQ(Nap)DPFKTKWKK | 81 | 1978 |
| 04-3 | 13-15 | GN(Dab)DAHSTQS(Iph)KWKK | 95 | 1983 |
| 04-3 | 13-15 | GHHSQDNAE(Nap)VKWKK | 72 | 1984 |
| 04-3 | 13-15 | GVTREL(Cit)FVD(Dab)KWKK | 92 | 1986 |
| 04-3 | 13-15 | G(Iph)SNNGSTNRNKWKK | 71 | 1987 |
| 04-3 | 13-15 | GFDQNDHVFKSKWKK | 92 | 1987 |
| 04-3 | 13-15 | GSDSVKTD(Iph)N(Dab)KWKK | 83 | 1989 |
| 04-3 | 13-15 | GKVDG(Iph)(Pal)GNFSKWKK | 77 | 1995 |
| 04-3 | 13-15 | GQDKSFNQQQKKWKK | 82 | 2001 |
| 04-3 | 13-15 | GGRNED(Nap)(Dab)(Cit)NPKWKK | 82 | 2006 |
| 04-3 | 13-15 | GFVHPDMRNQLKWKK | 89 | 2007 |
| 04-3 | 13-15 | GDG(Iph)PSVF(Nap)PAKWKK | 76 | 2010 |
| 04-3 | 13-15 | G(Cit)RVFDSKP(Cit)VKWKK | 81 | 2012 |

| | | | | |
|---|---|---|---|---|
| 04-3 | 13-15 | GVG(Cit)N(Iph)(Dab)TDQVKWKK | 88 | 2013 |
| 04-3 | 13-15 | GFEKD(Iph)TGVPVKWKK | 84 | 2015 |
| 04-3 | 13-15 | GFSTTQS(Iph)Q(Dab)TKWKK | 73 | 2023 |
| 04-3 | 13-15 | G(Cit)D(Cit)AAA(Iph)NLLKWKK | 77 | 2025 |
| 04-3 | 13-15 | G(Dab)GP(Cit)TS(Cit)(Iph)NNKWKK | 75 | 2027 |
| 04-3 | 13-15 | GNP(Iph)QVQHGQTKWKK | 82 | 2032 |
| 04-3 | 13-15 | GPPSNNA(Nap)(Iph)SQKWKK | 87 | 2035 |
| 04-3 | 13-15 | GD(Pal)FLVEDDHRKWKK | 79 | 2044 |
| 04-3 | 13-15 | G(Dab)DT(Iph)AFDHPQKWKK | 81 | 2054 |
| 04-3 | 13-15 | GFHGQV(Iph)PMTNKWKK | 79 | 2054 |
| 04-3 | 13-15 | GRS(Cit)A(Iph)QQNAVKWKK | 84 | 2054 |
| 04-3 | 13-15 | GG(Cit)A(Pip)DN(Pip)(Iph)FTKWKK | 75 | 2057 |
| 04-3 | 13-15 | GKAQ(Nap)(Nap)MEPNPKWKK | 74 | 2059 |
| 04-3 | 13-15 | GG(Cit)SSSH(Nap)N(Iph)PKWKK | 74 | 2063 |
| 04-3 | 13-15 | GKPS(Cit)DH(Iph)VAMKWKK | 88 | 2065 |
| 04-3 | 13-15 | GLEERNK(Cit)NDRKWKK | 80 | 2081 |
| 04-3 | 13-15 | GP(Cit)EAE(Cit)LHQ(Nap)KWKK | 77 | 2085 |
| 04-3 | 13-15 | GAVRPEHNE(Iph)LKWKK | 73 | 2088 |
| 04-3 | 13-15 | GS(Cit)HTEE(Cit)RA(Nap)KWKK | 85 | 2091 |
| 04-3 | 13-15 | GNF(Cit)(Dab)ES(Iph)Q(Dab)TKWKK | 85 | 2106 |
| 04-3 | 13-15 | G(Iph)SEFQRQSTTKWKK | 88 | 2107 |
| 04-3 | 13-15 | GLNHNNA(Nap)(Iph)SQKWKK | 82 | 2118 |
| 04-3 | 13-15 | GVVT(Cit)(Dab)(Iph)NT(Cit)(Cit)KWKK | 92 | 2128 |
| 04-3 | 13-15 | GQSLK(Cit)HD(Iph)L(Cit)KWKK | 77 | 2178 |
| 04-3 | 13-15 | G(Iph)QQHVS(Cit)S(Nap)QKWKK | 71 | 2191 |
| 04-3 | 13-15 | G(Iph)TTHSQ(Iph)(Cit)PTKWKK | 75 | 2225 |
| 05-3 | 15-17 | GSNGAKPSPTPKWKK | 76 | 1706 |
| 05-3 | 15-17 | GERAARAPTGGKWKK | 83 | 1736 |
| 05-3 | 15-17 | GHSAGKSPPLNKWKK | 93 | 1758 |
| 05-3 | 15-17 | GA(Pip)(Dab)HFSHGVGKWKK | 72 | 1788 |
| 05-3 | 15-17 | GHFGAKPPSTPKWKK | 85 | 1789 |
| 05-3 | 15-17 | GG(Cit)PGAVLDPRKWKK | 81 | 1789 |
| 05-3 | 15-17 | GVGERPA(Dab)LQAKWKK | 88 | 1791 |
| 05-3 | 15-17 | GKAAQSHV(Cit)SGKWKK | 81 | 1792 |
| 05-3 | 15-17 | G(Dab)SQPSEG(Dab)S(Cit)KWKK | 85 | 1799 |
| 05-3 | 15-17 | GPGAL(Nap)SHEGSKWKK | 81 | 1802 |
| 05-3 | 15-17 | GQVTRTGAGHEKWKK | 74 | 1806 |
| 05-3 | 15-17 | GVDST(Dab)KTA(Pip)DKWKK | 87 | 1813 |
| 05-3 | 15-17 | GLHSGAVLDPRKWKK | 73 | 1815 |

| | | | | |
|---|---|---|---|---|
| 05-3 | 15-17 | G(Dab)HGT(Cit)PTGQLKWKK | 86 | 1818 |
| 05-3 | 15-17 | GAVA(Cit)SSRKAEKWKK | 84 | 1826 |
| 05-3 | 15-17 | GVGFNFRAGLVKWKK | 71 | 1830 |
| 05-3 | 15-17 | GS(Dab)SEVGFNKNKWKK | 95 | 1832 |
| 05-3 | 15-17 | GTNELN(Dab)AANFKWKK | 83 | 1844 |
| 05-3 | 15-17 | GNGT(Dab)(Dab)PQFETKWKK | 84 | 1844 |
| 05-3 | 15-17 | GGARKPQQLVVKWKK | 79 | 1846 |
| 05-3 | 15-17 | G(Nap)(Pip)PVVGGS(Cit)TKWKK | 81 | 1847 |
| 05-3 | 15-17 | G(Dab)VKFQPPANVKWKK | 83 | 1850 |
| 05-3 | 15-17 | GPGLQDSPNQRKWKK | 77 | 1862 |
| 05-3 | 15-17 | GFL(Pip)(Dab)SSFSSNKWKK | 74 | 1865 |
| 05-3 | 15-17 | GRDSPSVTKLNKWKK | 90 | 1867 |
| 05-3 | 15-17 | G(Pip)P(Dab)GFGGDA(Iph)KWKK | 80 | 1870 |
| 05-3 | 15-17 | GALDVEHLPNNKWKK | 74 | 1872 |
| 05-3 | 15-17 | G(Nap)SFSN(Dab)G(Dab)SQKWKK | 72 | 1874 |
| 05-3 | 15-17 | GKTSQV(Cit)HVGNKWKK | 92 | 1877 |
| 05-3 | 15-17 | GTVDTPRNVKPKWKK | 77 | 1877 |
| 05-3 | 15-17 | GF(Dab)HVMPAQTPKWKK | 78 | 1878 |
| 05-3 | 15-17 | GVDT(Dab)P(Nap)TAEVKWKK | 88 | 1879 |
| 05-3 | 15-17 | GAHSTF(Cit)(Cit)G(Pip)AKWKK | 84 | 1881 |
| 05-3 | 15-17 | GFAGVN(Nap)A(Cit)(Dab)VKWKK | 85 | 1882 |
| 05-3 | 15-17 | GDFPKAATLRNKWKK | 87 | 1883 |
| 05-3 | 15-17 | GFRFGGHETPPKWKK | 94 | 1895 |
| 05-3 | 15-17 | GFGVSFRVKVDKWKK | 82 | 1904 |
| 05-3 | 15-17 | GGDP(Nap)PFELKGKWKK | 83 | 1907 |
| 05-3 | 15-17 | GAE(Dab)KAN(Cit)(Nap)SSKWKK | 81 | 1911 |
| 05-3 | 15-17 | G(Pal)(Pip)AGFEHKSLKWKK | 79 | 1913 |
| 05-3 | 15-17 | G(Dab)(Cit)DSQLS(Cit)V(Dab)KWKK | 82 | 1913 |
| 05-3 | 15-17 | GRNSEVGFNKNKWKK | 96 | 1915 |
| 05-3 | 15-17 | G(Pal)SLS(Dab)PTEFHKWKK | 86 | 1916 |
| 05-3 | 15-17 | GRSTAPQMNDFKWKK | 73 | 1917 |
| 05-3 | 15-17 | GG(Nap)T(Dab)(Dab)PQFETKWKK | 92 | 1927 |
| 05-3 | 15-17 | GLNDT(Dab)(Pal)KQLVKWKK | 76 | 1929 |
| 05-3 | 15-17 | GA(Nap)HPSAFEPKKWKK | 89 | 1931 |
| 05-3 | 15-17 | GFH(Dab)FVVELTPKWKK | 94 | 1939 |
| 05-3 | 15-17 | G(Nap)Q(Nap)HA(Dab)DVGAKWKK | 87 | 1942 |
| 05-3 | 15-17 | GV(Cit)QQQQ(Dab)VPNKWKK | 82 | 1948 |
| 05-3 | 15-17 | G(Pip)SSTFAEF(Cit)KKWKK | 91 | 1950 |
| 05-3 | 15-17 | GPQTNQRKESLKWKK | 85 | 1951 |

| | | | | |
|---|---|---|---|---|
| 05-3 | 15-17 | GG(Pal)GDKL(Cit)(Cit)QKKWKK | 78 | 1958 |
| 05-3 | 15-17 | GAH(Cit)(Cit)(Cit)(Dab)TPTLKWKK | 87 | 1961 |
| 05-3 | 15-17 | G(Cit)DASQFKN(Dab)FKWKK | 86 | 1964 |
| 05-3 | 15-17 | G(Cit)PKAES(Nap)(Dab)MPKWKK | 80 | 1964 |
| 05-3 | 15-17 | GF(Iph)TQPGTVGHKWKK | 84 | 1967 |
| 05-3 | 15-17 | GHFGE(Dab)(Nap)TTNNKWKK | 97 | 1967 |
| 05-3 | 15-17 | GRFN(Dab)ESNLLMKWKK | 96 | 1974 |
| 05-3 | 15-17 | GFE(Nap)PGQHGQQKWKK | 75 | 1975 |
| 05-3 | 15-17 | GAFSDHA(Iph)GFTKWKK | 75 | 1976 |
| 05-3 | 15-17 | GASLGHD(Iph)VQKKWKK | 87 | 1978 |
| 05-3 | 15-17 | GHKE(Cit)PAFVDKKWKK | 75 | 1978 |
| 05-3 | 15-17 | GDPRQTKVD(Pip)FKWKK | 73 | 1982 |
| 05-3 | 15-17 | GKGAGG(Iph)F(Cit)EHKWKK | 84 | 1983 |
| 05-3 | 15-17 | GNHT(Nap)PFELKGKWKK | 90 | 1990 |
| 05-3 | 15-17 | GPNAF(Nap)(Cit)VKPLKWKK | 76 | 1990 |
| 05-3 | 15-17 | GNRSQSG(Iph)KEAKWKK | 96 | 2000 |
| 05-3 | 15-17 | G(Pal)A(Nap)(Cit)PNGP(Cit)HKWKK | 74 | 2002 |
| 05-3 | 15-17 | G(Pal)PRF(Pip)(Iph)TAGGKWKK | 72 | 2003 |
| 05-3 | 15-17 | GR(Nap)SDLNPSHMKWKK | 79 | 2004 |
| 05-3 | 15-17 | GAHA(Iph)KMLPSEKWKK | 76 | 2007 |
| 05-3 | 15-17 | GQRETMF(Dab)S(Cit)TKWKK | 86 | 2007 |
| 05-3 | 15-17 | GPTP(Cit)(Nap)FSEKPKWKK | 79 | 2007 |
| 05-3 | 15-17 | GQGVQD(Iph)VNPKKWKK | 82 | 2008 |
| 05-3 | 15-17 | GFK(Iph)MGTPFGTKWKK | 84 | 2009 |
| 05-3 | 15-17 | GARTSESN(Iph)(Dab)QKWKK | 93 | 2016 |
| 05-3 | 15-17 | GRFS(Dab)(Cit)D(Iph)PGGKWKK | 88 | 2016 |
| 05-3 | 15-17 | GATASDK(Iph)RS(Cit)KWKK | 74 | 2016 |
| 05-3 | 15-17 | GSQDRFS(Dab)PG(Iph)KWKK | 79 | 2017 |
| 05-3 | 15-17 | GKKT(Iph)GA(Cit)AFDKWKK | 83 | 2018 |
| 05-3 | 15-17 | GK(Cit)SQNLQH(Cit)VKWKK | 83 | 2018 |
| 05-3 | 15-17 | GGKQSNKLE(Iph)PKWKK | 87 | 2024 |
| 05-3 | 15-17 | GA(Cit)LF(Nap)RQGNNKWKK | 72 | 2024 |
| 05-3 | 15-17 | GA(Pip)EDAPR(Iph)AFKWKK | 78 | 2026 |
| 05-3 | 15-17 | GRSFA(Nap)DQRLVKWKK | 84 | 2039 |
| 05-3 | 15-17 | GMRPLHEN(Nap)PTKWKK | 79 | 2042 |
| 05-3 | 15-17 | G(Cit)VQPSDRVH(Nap)KWKK | 86 | 2042 |
| 05-3 | 15-17 | GQES(Dab)MNN(Dab)T(Iph)KWKK | 85 | 2047 |
| 05-3 | 15-17 | GAT(Iph)TEKENSFKWKK | 73 | 2050 |
| 05-3 | 15-17 | GHAAF(Iph)TRDG(Cit)KWKK | 98 | 2055 |

| | | | | |
|---|---|---|---|---|
| 05-3 | 15-17 | GVLFTNSHG(Iph)(Cit)KWKK | 95 | 2055 |
| 05-3 | 15-17 | GTN(Dab)TPKEML(Iph)KWKK | 78 | 2057 |
| 05-3 | 15-17 | G(Iph)(Cit)HKNGEPVVKWKK | 89 | 2060 |
| 05-3 | 15-17 | GSTDHVS(Dab)(Nap)P(Iph)KWKK | 78 | 2063 |
| 05-3 | 15-17 | G(Dab)N(Iph)PTEHNFSKWKK | 93 | 2069 |
| 05-3 | 15-17 | GKNSDTV(Iph)LNRKWKK | 97 | 2070 |
| 05-3 | 15-17 | GRLVFD(Dab)AE(Iph)VKWKK | 86 | 2072 |
| 05-3 | 15-17 | GDV(Nap)RQRMTVQKWKK | 82 | 2080 |
| 05-3 | 15-17 | GLDQARSR(Cit)(Nap)MKWKK | 78 | 2081 |
| 05-3 | 15-17 | GPTK(Dab)QD(Iph)LFLKWKK | 84 | 2085 |
| 05-3 | 15-17 | GHQL(Cit)(Nap)FSEV(Pip)KWKK | 81 | 2090 |
| 05-3 | 15-17 | GPNRQD(Iph)VNPKKWKK | 87 | 2091 |
| 05-3 | 15-17 | G(Iph)HQSSVF(Nap)PAKWKK | 91 | 2093 |
| 05-3 | 15-17 | GSFDK(Pip)HTDT(Iph)KWKK | 82 | 2100 |
| 05-3 | 15-17 | G(Cit)(Iph)QL(Cit)FAS(Dab)PKWKK | 84 | 2100 |
| 05-3 | 15-17 | GT(Cit)(Dab)NFG(Iph)LDRKWKK | 93 | 2103 |
| 05-3 | 15-17 | G(Iph)KNTSN(Pal)HELKWKK | 84 | 2114 |
| 05-3 | 15-17 | GA(Iph)(Cit)RRTDMPSKWKK | 77 | 2114 |
| 05-3 | 15-17 | GVHSVP(Iph)F(Cit)EKKWKK | 84 | 2123 |
| 05-3 | 15-17 | G(Iph)QNPNAP(Cit)(Cit)RKWKK | 72 | 2134 |
| 05-3 | 15-17 | GKL(Iph)G(Iph)QQNGTKWKK | 88 | 2142 |
| 05-3 | 15-17 | GV(Iph)G(Dab)HQTPD(Iph)KWKK | 82 | 2150 |
| 05-3 | 15-17 | G(Cit)KT(Nap)DHSF(Nap)QKWKK | 88 | 2164 |
| 05-3 | 15-17 | GT(Iph)R(Cit)LKDPQFKWKK | 86 | 2185 |
| 06-3 | 17-20 | G(Dab)SPVSGGSVKKWKK | 90 | 1668 |
| 06-3 | 17-20 | G(Dab)HPSNGG(Cit)GAKWKK | 80 | 1704 |
| 06-3 | 17-20 | GVTPKTALGKGKWKK | 89 | 1722 |
| 06-3 | 17-20 | GGKNAVLT(Dab)VAKWKK | 90 | 1723 |
| 06-3 | 17-20 | GAVVSRAVS(Dab)SKWKK | 82 | 1726 |
| 06-3 | 17-20 | GASFSHSA(Dab)AVKWKK | 75 | 1727 |
| 06-3 | 17-20 | GGG(Dab)SFATKLTKWKK | 94 | 1732 |
| 06-3 | 17-20 | G(Dab)SVPAFNVG(Dab)KWKK | 72 | 1741 |
| 06-3 | 17-20 | G(Dab)(Dab)(Pal)(Dab)TGGLSNKWKK | 88 | 1747 |
| 06-3 | 17-20 | GR(Nap)(Dab)GVSGTGAKWKK | 72 | 1752 |
| 06-3 | 17-20 | GAHSLGTSALRKWKK | 82 | 1763 |
| 06-3 | 17-20 | GD(Pip)(Pip)GGHSTSTKWKK | 76 | 1764 |
| 06-3 | 17-20 | GVM(Dab)AARLASTKWKK | 82 | 1770 |
| 06-3 | 17-20 | GS(Dab)QAGG(Nap)(Dab)NTKWKK | 76 | 1782 |
| 06-3 | 17-20 | GNG(Dab)GNHKVPNKWKK | 77 | 1787 |

| | | | | |
|---|---|---|---|---|
| 06-3 | 17-20 | G(Dab)NPSAGLFTHKWKK | 88 | 1794 |
| 06-3 | 17-20 | GP(Dab)FVVQGT(Dab)PKWKK | 87 | 1795 |
| 06-3 | 17-20 | GA(Nap)TE(Dab)GGFA(Dab)KWKK | 87 | 1800 |
| 06-3 | 17-20 | GSGRG(Cit)GLHNVKWKK | 80 | 1804 |
| 06-3 | 17-20 | GVMP(Pip)VHGTSTKWKK | 91 | 1805 |
| 06-3 | 17-20 | GLVASMGHTNKKWKK | 74 | 1808 |
| 06-3 | 17-20 | GNK(Dab)EKGLLGTKWKK | 75 | 1810 |
| 06-3 | 17-20 | GAV(Pip)(Dab)PVHDMAKWKK | 81 | 1816 |
| 06-3 | 17-20 | G(Dab)AQHTTHESGKWKK | 86 | 1818 |
| 06-3 | 17-20 | GRNAPVFNVG(Dab)KWKK | 80 | 1824 |
| 06-3 | 17-20 | GFA(Dab)L(Dab)TNLSNKWKK | 93 | 1830 |
| 06-3 | 17-20 | GLFSFKGQ(Dab)GPKWKK | 87 | 1831 |
| 06-3 | 17-20 | GL(Cit)(Dab)PSKVPPSKWKK | 77 | 1832 |
| 06-3 | 17-20 | GHHEETGGVSHKWKK | 71 | 1840 |
| 06-3 | 17-20 | GS(Cit)TGHF(Dab)TLAKWKK | 83 | 1841 |
| 06-3 | 17-20 | GGMGSKNN(Dab)(Nap)SKWKK | 83 | 1842 |
| 06-3 | 17-20 | G(Dab)KTTPRSGVFKWKK | 71 | 1843 |
| 06-3 | 17-20 | GG(Dab)NKA(Nap)PKMGKWKK | 75 | 1850 |
| 06-3 | 17-20 | GSGKRDEAQGRKWKK | 75 | 1854 |
| 06-3 | 17-20 | GVLTP(Pip)N(Dab)VQLKWKK | 82 | 1860 |
| 06-3 | 17-20 | GTAAAK(Iph)G(Dab)SHKWKK | 77 | 1866 |
| 06-3 | 17-20 | GAG(Nap)S(Pip)VKTTMKWKK | 73 | 1868 |
| 06-3 | 17-20 | GFTSSGPQQRLKWKK | 75 | 1871 |
| 06-3 | 17-20 | GVQN(Dab)QF(Dab)AD(Dab)KWKK | 90 | 1872 |
| 06-3 | 17-20 | GVSKF(Cit)ATKANKWKK | 91 | 1873 |
| 06-3 | 17-20 | GTGHGVHVFRDKWKK | 87 | 1875 |
| 06-3 | 17-20 | GKVNVSNTLPRKWKK | 79 | 1878 |
| 06-3 | 17-20 | G(Dab)ESSQHNKAEKWKK | 93 | 1880 |
| 06-3 | 17-20 | GN(Cit)(Pip)(Dab)NVFGPTKWKK | 82 | 1882 |
| 06-3 | 17-20 | GQ(Dab)AFMGPRA(Cit)KWKK | 94 | 1885 |
| 06-3 | 17-20 | GKGPGPE(Nap)NSRKWKK | 77 | 1889 |
| 06-3 | 17-20 | GKLLPMGSQHQKWKK | 93 | 1889 |
| 06-3 | 17-20 | GHLSNS(Cit)QKLGKWKK | 88 | 1891 |
| 06-3 | 17-20 | GSHLTVL(Cit)G(Pip)MKWKK | 89 | 1891 |
| 06-3 | 17-20 | GQNFPLTTHVSKWKK | 73 | 1894 |
| 06-3 | 17-20 | GGSTAKA(Iph)SLHKWKK | 72 | 1895 |
| 06-3 | 17-20 | G(Pip)AGAPE(Cit)FNRKWKK | 82 | 1895 |
| 06-3 | 17-20 | GKVARQTLSNQKWKK | 77 | 1895 |
| 06-3 | 17-20 | GTG(Pip)HNHVEVEKWKK | 90 | 1898 |

| | | | | |
|---|---|---|---|---|
| 06-3 | 17-20 | G(Dab)NLLSTMTRNKWKK | 83 | 1900 |
| 06-3 | 17-20 | GKPFRSGQNSMKWKK | 88 | 1902 |
| 06-3 | 17-20 | GFTRAKVVFSPKWKK | 92 | 1902 |
| 06-3 | 17-20 | GKTNSFSQQ(Dab)NKWKK | 88 | 1904 |
| 06-3 | 17-20 | GPV(Cit)SARPNKQKWKK | 83 | 1904 |
| 06-3 | 17-20 | GNMLVAKA(Nap)(Dab)NKWKK | 75 | 1908 |
| 06-3 | 17-20 | GAFK(Nap)QRGG(Dab)PKWKK | 73 | 1908 |
| 06-3 | 17-20 | GHDPDTVPKHLKWKK | 88 | 1909 |
| 06-3 | 17-20 | GLFDGGRGLK(Nap)KWKK | 82 | 1910 |
| 06-3 | 17-20 | G(Dab)TNA(Dab)NA(Iph)PTKWKK | 86 | 1912 |
| 06-3 | 17-20 | GAGENFG(Cit)HKFKWKK | 78 | 1914 |
| 06-3 | 17-20 | GKQMLLM(Dab)A(Dab)MKWKK | 77 | 1916 |
| 06-3 | 17-20 | G(Cit)NHSRVVNANKWKK | 80 | 1918 |
| 06-3 | 17-20 | GQTTTNEALHRKWKK | 78 | 1921 |
| 06-3 | 17-20 | GFGSVVKK(Cit)T(Pal)KWKK | 82 | 1921 |
| 06-3 | 17-20 | GSKV(Nap)LEFPG(Dab)KWKK | 79 | 1924 |
| 06-3 | 17-20 | GVSKSA(Nap)L(Cit)KSKWKK | 73 | 1924 |
| 06-3 | 17-20 | GVRT(Cit)G(Cit)(Pip)GVFKWKK | 77 | 1926 |
| 06-3 | 17-20 | GLTNFHSPNHLKWKK | 79 | 1930 |
| 06-3 | 17-20 | G(Iph)KAAAPPAFHKWKK | 89 | 1933 |
| 06-3 | 17-20 | GEFSGKQNRSMKWKK | 76 | 1934 |
| 06-3 | 17-20 | GQHHTSNVPHEKWKK | 91 | 1936 |
| 06-3 | 17-20 | GLMNGRFTHNPKWKK | 80 | 1937 |
| 06-3 | 17-20 | G(Cit)VFTE(Nap)GGVKKWKK | 75 | 1941 |
| 06-3 | 17-20 | GL(Dab)QS(Cit)RTFQGKWKK | 76 | 1944 |
| 06-3 | 17-20 | GTLHRNQN(Cit)GVKWKK | 94 | 1946 |
| 06-3 | 17-20 | G(Dab)AF(Cit)FNQKQGKWKK | 91 | 1947 |
| 06-3 | 17-20 | G(Dab)AF(Cit)FNQKQGKWKK | 89 | 1947 |
| 06-3 | 17-20 | GNPQVP(Dab)VG(Iph)NKWKK | 77 | 1948 |
| 06-3 | 17-20 | GVRNEEH(Dab)SNLKWKK | 88 | 1948 |
| 06-3 | 17-20 | GPKG(Nap)(Nap)QGVAFKWKK | 90 | 1948 |
| 06-3 | 17-20 | G(Pal)ARP(Cit)QQGHTKWKK | 84 | 1950 |
| 06-3 | 17-20 | GVM(Dab)ARTVNN(Nap)KWKK | 82 | 1952 |
| 06-3 | 17-20 | GKSQL(Cit)S(Cit)VARKWKK | 77 | 1953 |
| 06-3 | 17-20 | GERSTLHDSKMKWKK | 87 | 1954 |
| 06-3 | 17-20 | GF(Pip)QVPPRFTSKWKK | 75 | 1955 |
| 06-3 | 17-20 | GQAETRHAFF(Dab)KWKK | 72 | 1957 |
| 06-3 | 17-20 | G(Dab)NHGLVPAQ(Iph)KWKK | 74 | 1959 |
| 06-3 | 17-20 | G(Dab)GTKANEF(Nap)FKWKK | 87 | 1961 |

| | | | | |
|---|---|---|---|---|
| 06-3 | 17-20 | G(Nap)(Dab)T(Dab)P(Nap)TAEVKWKK | 92 | 1962 |
| 06-3 | 17-20 | GGAHLTTN(Dab)Q(Iph)KWKK | 85 | 1965 |
| 06-3 | 17-20 | GG(Dab)PFF(Nap)H(Cit)TGKWKK | 89 | 1967 |
| 06-3 | 17-20 | GQVG(Iph)STAPK(Cit)KWKK | 72 | 1968 |
| 06-3 | 17-20 | GVVGVKSQ(Iph)Q(Dab)KWKK | 90 | 1968 |
| 06-3 | 17-20 | GM(Nap)TKLSHVPNKWKK | 92 | 1974 |
| 06-3 | 17-20 | GG(Dab)(Cit)E(Cit)RFVMAKWKK | 91 | 1974 |
| 06-3 | 17-20 | GKSS(Nap)TQNPMHKWKK | 89 | 1977 |
| 06-3 | 17-20 | G(Dab)PEP(Nap)FTTSRKWKK | 91 | 1982 |
| 06-3 | 17-20 | GKGAGG(Iph)F(Cit)EHKWKK | 82 | 1983 |
| 06-3 | 17-20 | GVGLQS(Cit)KA(Iph)TKWKK | 80 | 1984 |
| 06-3 | 17-20 | GS(Nap)SKRQLMTSKWKK | 77 | 1985 |
| 06-3 | 17-20 | GAL(Pip)(Nap)QHPDHTKWKK | 82 | 1992 |
| 06-3 | 17-20 | GA(Dab)VE(Iph)(Dab)QLLPKWKK | 73 | 1993 |
| 06-3 | 17-20 | GQFLLTEAQHRKWKK | 74 | 1993 |
| 06-3 | 17-20 | G(Iph)SPPN(Cit)FAANKWKK | 80 | 1998 |
| 06-3 | 17-20 | G(Cit)TNN(Nap)LVGEFKWKK | 74 | 1998 |
| 06-3 | 17-20 | GAS(Iph)TSFFEVPKWKK | 79 | 2008 |
| 06-3 | 17-20 | GAS(Iph)TSFFQ(Dab)PKWKK | 90 | 2008 |
| 06-3 | 17-20 | G(Pip)LT(Dab)(Iph)NVVVNKWKK | 81 | 2008 |
| 06-3 | 17-20 | GQSS(Iph)NRSTHAKWKK | 77 | 2011 |
| 06-3 | 17-20 | GNR(Cit)LHSFFPSKWKK | 77 | 2012 |
| 06-3 | 17-20 | GS(Cit)FFFTGFPRKWKK | 74 | 2013 |
| 06-3 | 17-20 | GH(Nap)DMRHQAATKWKK | 84 | 2014 |
| 06-3 | 17-20 | GKGGTPPNK(Iph)(Nap)KWKK | 89 | 2019 |
| 06-3 | 17-20 | G(Cit)ATK(Dab)FT(Cit)(Cit)MKWKK | 85 | 2020 |
| 06-3 | 17-20 | GGVSKHEQV(Iph)NKWKK | 73 | 2021 |
| 06-3 | 17-20 | GNTPL(Iph)KD(Dab)PLKWKK | 72 | 2021 |
| 06-3 | 17-20 | G(Pip)FNS(Nap)PLENKKWKK | 88 | 2022 |
| 06-3 | 17-20 | GQ(Dab)Q(Cit)STSGH(Iph)KWKK | 93 | 2025 |
| 06-3 | 17-20 | G(Cit)HQVP(Dab)VG(Iph)NKWKK | 83 | 2031 |
| 06-3 | 17-20 | GPPHT(Iph)GKT(Cit)LKWKK | 87 | 2031 |
| 06-3 | 17-20 | GNTAEHK(Nap)ERVKWKK | 72 | 2031 |
| 06-3 | 17-20 | G(Dab)(Nap)EP(Nap)TTNRAKWKK | 89 | 2033 |
| 06-3 | 17-20 | GLNFRNK(Cit)GFMKWKK | 89 | 2034 |
| 06-3 | 17-20 | G(Pip)(Iph)FVTFTNGTKWKK | 97 | 2036 |
| 06-3 | 17-20 | GND(Nap)SLQPFRNKWKK | 82 | 2038 |
| 06-3 | 17-20 | GRAFL(Nap)PAREMKWKK | 88 | 2038 |
| 06-3 | 17-20 | GG(Nap)ND(Cit)TNKMRKWKK | 76 | 2040 |

| | | | | |
|---|---|---|---|---|
| 06-3 | 17-20 | GVN(Nap)F(Nap)PLAQNKWKK | 71 | 2047 |
| 06-3 | 17-20 | GFT(Dab)LTTNLD(Iph)KWKK | 76 | 2048 |
| 06-3 | 17-20 | GHRHGASFQ(Iph)SKWKK | 72 | 2050 |
| 06-3 | 17-20 | GS(Iph)SSMMKRSNKWKK | 71 | 2051 |
| 06-3 | 17-20 | GNPR(Iph)STAPK(Cit)KWKK | 80 | 2051 |
| 06-3 | 17-20 | GTTT(Iph)KLHTNLKWKK | 75 | 2052 |
| 06-3 | 17-20 | GP(Dab)(Iph)(Dab)(Dab)H(Pal)(Cit)A(Dab) KWKK | 77 | 2053 |
| 06-3 | 17-20 | GTFK(Nap)SRELA(Cit)KWKK | 79 | 2056 |
| 06-3 | 17-20 | GFKV(Iph)HNGANFKWKK | 97 | 2057 |
| 06-3 | 17-20 | GL(Nap)RDPAGHA(Iph)KWKK | 74 | 2057 |
| 06-3 | 17-20 | GHT(Nap)E(Iph)G(Pip)LGTKWKK | 90 | 2061 |
| 06-3 | 17-20 | GS(Pal)D(Dab)(Iph)EVTKNKWKK | 71 | 2064 |
| 06-3 | 17-20 | GRALNQG(Nap)F(Nap)NKWKK | 72 | 2064 |
| 06-3 | 17-20 | GRVRGELQN(Iph)AKWKK | 77 | 2066 |
| 06-3 | 17-20 | GFL(Iph)NVNRAPNKWKK | 73 | 2068 |
| 06-3 | 17-20 | GNTMHQSH(Cit)(Nap)LKWKK | 79 | 2072 |
| 06-3 | 17-20 | G(Iph)KGSFGE(Cit)RNKWKK | 90 | 2075 |
| 06-3 | 17-20 | GRMDH(Cit)TQG(Pip)(Nap)KWKK | 76 | 2075 |
| 06-3 | 17-20 | GPRFDLF(Cit)QF(Dab)KWKK | 91 | 2077 |
| 06-3 | 17-20 | GPHFNRS(Cit)A(Nap)FKWKK | 90 | 2080 |
| 06-3 | 17-20 | GRGQNN(Nap)LREFKWKK | 90 | 2081 |
| 06-3 | 17-20 | GLKNEF(Pip)(Iph)TQGKWKK | 96 | 2086 |
| 06-3 | 17-20 | G(Nap)(Dab)STTRF(Cit)N(Cit)KWKK | 92 | 2087 |
| 06-3 | 17-20 | GFNEF(Cit)TT(Cit)HKKWKK | 87 | 2088 |
| 06-3 | 17-20 | GATFKQH(Iph)PQNKWKK | 74 | 2094 |
| 06-3 | 17-20 | GTGRS(Iph)HQ(Cit)QTKWKK | 78 | 2095 |
| 06-3 | 17-20 | GLNNKVHVT(Iph)FKWKK | 95 | 2095 |
| 06-3 | 17-20 | GMVSRQF(Cit)(Cit)K(Cit)KWKK | 85 | 2117 |
| 06-3 | 17-20 | GDKD(Iph)(Cit)QHVAKKWKK | 84 | 2121 |
| 06-3 | 17-20 | GKM(Pip)NL(Iph)ENPQKWKK | 90 | 2123 |
| 06-3 | 17-20 | GFTKHQSFQ(Iph)SKWKK | 84 | 2133 |
| 06-3 | 17-20 | GQ(Cit)T(Iph)KLHTNLKWKK | 90 | 2135 |
| 06-3 | 17-20 | GAAKLE(Cit)FQ(Iph)RKWKK | 74 | 2143 |
| 06-3 | 17-20 | GAHQ(Nap)(Iph)EVTKNKWKK | 88 | 2147 |
| 06-3 | 17-20 | G(Iph)(Iph)GVNHVPDNKWKK | 83 | 2148 |
| 06-3 | 17-20 | GRK(Nap)VN(Cit)SEN(Nap)KWKK | 95 | 2148 |
| 06-3 | 17-20 | GV(Iph)KNQQNR(Cit)SKWKK | 76 | 2154 |
| 06-3 | 17-20 | GELLG(Iph)(Iph)PARDKWKK | 80 | 2167 |

| | | | | |
|---|---|---|---|---|
| 06-3 | 17-20 | GVHN(Iph)LEN(Nap)(Dab)QKWKK | 75 | 2174 |
| 06-3 | 17-20 | GVQPLR(Iph)EQK(Cit)KWKK | 71 | 2178 |
| 06-3 | 17-20 | GTENTHG(Iph)SH(Iph)KWKK | 73 | 2179 |
| 06-3 | 17-20 | GEHT(Iph)GGQKQ(Iph)KWKK | 81 | 2181 |
| 06-3 | 17-20 | GSATE(Iph)NNH(Iph)LKWKK | 75 | 2182 |
| 06-3 | 17-20 | GV(Iph)GR(Iph)PDKQVKWKK | 80 | 2195 |
| 06-3 | 17-20 | GKQSK(Nap)NFNN(Iph)KWKK | 84 | 2200 |
| 06-3 | 17-20 | GL(Cit)NH(Iph)LQQRNKWKK | 90 | 2203 |
| 06-3 | 17-20 | G(Iph)KRD(Cit)PN(Iph)AGKWKK | 87 | 2211 |
| 06-3 | 17-20 | GRT(Nap)GNQ(Iph)F(Pal)QKWKK | 75 | 2219 |
| 06-3 | 17-20 | GTH(Iph)TVELQ(Iph)VKWKK | 85 | 2223 |
| 06-3 | 17-20 | GQPKL(Cit)EFQ(Iph)RKWKK | 72 | 2226 |
| 06-3 | 17-20 | GLANNR(Iph)DHV(Iph)KWKK | 71 | 2235 |
| 06-3 | 17-20 | GNQ(Dab)E(Iph)NNH(Iph)LKWKK | 93 | 2265 |
| 06-3 | 17-20 | G(Nap)(Pal)(Cit)(Cit)NLG(Cit)K(Iph)KWKK | 72 | 2271 |
| 06-3 | 17-20 | GVP(Nap)EER(Iph)RQ(Cit)KWKK | 80 | 2291 |
| 06-3 | 17-20 | GN(Pal)P(Dab)(Iph)VD(Iph)(Cit)RKWKK | 72 | 2302 |
| 07-3 | 20-23 | GVGGGSGVGV(Iph)KWKK | 81 | 1712 |
| 07-3 | 20-23 | GSGLKHGE(Dab)PAKWKK | 78 | 1746 |
| 07-3 | 20-23 | GLKG(Pal)NGVTAKKWKK | 71 | 1786 |
| 07-3 | 20-23 | GAKQEL(Dab)GPSKKWKK | 90 | 1808 |
| 07-3 | 20-23 | G(Dab)GHNQF(Pip)STGKWKK | 86 | 1824 |
| 07-3 | 20-23 | GQ(Dab)LEHNKPGGKWKK | 82 | 1830 |
| 07-3 | 20-23 | GLFSFKGQ(Dab)GPKWKK | 88 | 1831 |
| 07-3 | 20-23 | GDSS(Pip)VHNAHVKWKK | 74 | 1842 |
| 07-3 | 20-23 | GNGAFV(Cit)(Dab)TKTKWKK | 90 | 1845 |
| 07-3 | 20-23 | GPLNFLLSHVGKWKK | 82 | 1847 |
| 07-3 | 20-23 | GSNSVSEVKLHKWKK | 92 | 1850 |
| 07-3 | 20-23 | GSA(Dab)T(Dab)ELFPHKWKK | 87 | 1852 |
| 07-3 | 20-23 | GAFT(Dab)S(Nap)ARPGKWKK | 88 | 1854 |
| 07-3 | 20-23 | GAFVHLFRSGAKWKK | 73 | 1855 |
| 07-3 | 20-23 | GSAAN(Nap)SHKLSKWKK | 89 | 1862 |
| 07-3 | 20-23 | GHPKAGPTEFKKWKK | 73 | 1862 |
| 07-3 | 20-23 | GFVGPPNLLKKKWKK | 73 | 1863 |
| 07-3 | 20-23 | G(Dab)RLAEQPT(Dab)VKWKK | 86 | 1864 |
| 07-3 | 20-23 | GNNNT(Cit)T(Dab)KGNKWKK | 72 | 1870 |
| 07-3 | 20-23 | GLFGVQTKTTKKWKK | 96 | 1873 |
| 07-3 | 20-23 | GR(Dab)LNEVKSQGKWKK | 92 | 1881 |
| 07-3 | 20-23 | GLFRGKN(Nap)GGSKWKK | 91 | 1883 |

| | | | | |
|---|---|---|---|---|
| 07-3 | 20-23 | GSNSRVFVLGRKWKK | 92 | 1885 |
| 07-3 | 20-23 | GKEHGNVGNMRKWKK | 75 | 1892 |
| 07-3 | 20-23 | GKHA(Nap)GPDA(Cit)(Dab)KWKK | 75 | 1900 |
| 07-3 | 20-23 | GAFP(Pal)ANSRLKKWKK | 89 | 1902 |
| 07-3 | 20-23 | GVDVGKNTHERKWKK | 82 | 1905 |
| 07-3 | 20-23 | GDVSHVH(Dab)LFTKWKK | 93 | 1905 |
| 07-3 | 20-23 | GPPVDKRVM(Dab)LKWKK | 89 | 1905 |
| 07-3 | 20-23 | GEERAFHAV(Dab)VKWKK | 96 | 1908 |
| 07-3 | 20-23 | GS(Dab)GF(Iph)FT(Dab)AGKWKK | 75 | 1910 |
| 07-3 | 20-23 | G(Iph)(Dab)GDTGNGNRKWKK | 71 | 1914 |
| 07-3 | 20-23 | GFTKTFG(Dab)NHLKWKK | 98 | 1915 |
| 07-3 | 20-23 | GN(Cit)G(Pip)(Nap)KGG(Nap)GKWKK | 83 | 1917 |
| 07-3 | 20-23 | GNKAFKTESKNKWKK | 90 | 1917 |
| 07-3 | 20-23 | G(Nap)TA(Cit)AGLKTRKWKK | 82 | 1922 |
| 07-3 | 20-23 | GVKERAPKAFQKWKK | 83 | 1924 |
| 07-3 | 20-23 | G(Iph)G(Dab)(Cit)V(Dab)GGRTKWKK | 72 | 1927 |
| 07-3 | 20-23 | G(Dab)FSVRS(Pip)(Cit)DSKWKK | 77 | 1931 |
| 07-3 | 20-23 | GSFTRDKGR(Dab)DKWKK | 79 | 1932 |
| 07-3 | 20-23 | GHFSVSEVKLHKWKK | 96 | 1933 |
| 07-3 | 20-23 | GLRVGGKR(Nap)DSKWKK | 82 | 1935 |
| 07-3 | 20-23 | GTGFLLQHKDKKWKK | 94 | 1937 |
| 07-3 | 20-23 | GGARMRF(Dab)NTHKWKK | 81 | 1940 |
| 07-3 | 20-23 | G(Dab)G(Pal)G(Nap)R(Pip)Q(Dab)TKWKK | 75 | 1940 |
| 07-3 | 20-23 | GV(Dab)MEPSEFR(Dab)KWKK | 79 | 1945 |
| 07-3 | 20-23 | GVVHHEFNGQKKWKK | 94 | 1945 |
| 07-3 | 20-23 | GN(Nap)NT(Cit)T(Dab)KGNKWKK | 94 | 1953 |
| 07-3 | 20-23 | GSPFM(Nap)(Dab)QPAKKWKK | 79 | 1953 |
| 07-3 | 20-23 | GVDNNTDNFHKKWKK | 79 | 1954 |
| 07-3 | 20-23 | GHAKKNEMSFNKWKK | 83 | 1956 |
| 07-3 | 20-23 | GLKNG(Dab)MTF(Nap)(Dab)KWKK | 74 | 1958 |
| 07-3 | 20-23 | GVKDGPN(Iph)LHGKWKK | 85 | 1960 |
| 07-3 | 20-23 | GSNQ(Dab)QFKEVHKWKK | 84 | 1967 |
| 07-3 | 20-23 | G(Nap)SGARHTFFTKWKK | 80 | 1971 |
| 07-3 | 20-23 | G(Iph)EKLTFGVASKWKK | 78 | 1975 |
| 07-3 | 20-23 | G(Dab)EQN(Nap)LP(Dab)(Dab)QKWKK | 85 | 1976 |
| 07-3 | 20-23 | GPVP(Nap)AEFRL(Dab)KWKK | 87 | 1976 |
| 07-3 | 20-23 | GRHPFGDNNNRKWKK | 78 | 1977 |
| 07-3 | 20-23 | G(Pal)PS(Nap)(Pip)NRQGTKWKK | 87 | 1981 |
| 07-3 | 20-23 | GVP(Dab)VPE(Dab)K(Iph)VKWKK | 72 | 1991 |

| | | | | |
|---|---|---|---|---|
| 07-3 | 20-23 | GQ(Pip)AFPPE(Cit)HHKWKK | 71 | 1996 |
| 07-3 | 20-23 | GQRH(Nap)HNGGFPKWKK | 81 | 1997 |
| 07-3 | 20-23 | GNHD(Nap)K(Cit)VSGHKWKK | 91 | 1998 |
| 07-3 | 20-23 | GQNEHHHFGVFKWKK | 93 | 2002 |
| 07-3 | 20-23 | G(Dab)QNKKEP(Nap)LVKWKK | 83 | 2003 |
| 07-3 | 20-23 | GNFVF(Cit)STH(Dab)FKWKK | 93 | 2006 |
| 07-3 | 20-23 | GVFSNNHA(Pip)(Iph)AKWKK | 84 | 2009 |
| 07-3 | 20-23 | GKGVT(Iph)KRANLKWKK | 88 | 2010 |
| 07-3 | 20-23 | GNGSTKMNVH(Iph)KWKK | 71 | 2011 |
| 07-3 | 20-23 | G(Nap)QSQAPRNMHKWKK | 83 | 2016 |
| 07-3 | 20-23 | GAR(Dab)ATFFQG(Iph)KWKK | 81 | 2021 |
| 07-3 | 20-23 | G(Dab)SG(Nap)(Cit)(Iph)PGRAKWKK | 83 | 2022 |
| 07-3 | 20-23 | GKKS(Nap)HGFFPEKWKK | 91 | 2024 |
| 07-3 | 20-23 | GKKS(Nap)HG(Pal)MLEKWKK | 84 | 2025 |
| 07-3 | 20-23 | GF(Dab)A(Nap)PSPR(Cit)FKWKK | 79 | 2026 |
| 07-3 | 20-23 | GD(Iph)NKKLGT(Dab)QKWKK | 82 | 2027 |
| 07-3 | 20-23 | GTTNFRA(Nap)SFHKWKK | 82 | 2028 |
| 07-3 | 20-23 | G(Dab)KT(Iph)FGAGQ(Nap)KWKK | 71 | 2029 |
| 07-3 | 20-23 | G(Nap)KTNTHHKDTKWKK | 95 | 2029 |
| 07-3 | 20-23 | GKT(Dab)(Iph)VEPNHSKWKK | 88 | 2035 |
| 07-3 | 20-23 | G(Nap)N(Pip)DGNA(Nap)FKKWKK | 95 | 2036 |
| 07-3 | 20-23 | G(Dab)(Nap)NNTDNFHKKWKK | 96 | 2037 |
| 07-3 | 20-23 | G(Iph)S(Dab)(Dab)P(Iph)TASSKWKK | 96 | 2046 |
| 07-3 | 20-23 | GVG(Iph)HLHQVS(Cit)KWKK | 77 | 2057 |
| 07-3 | 20-23 | G(Pal)EGRNALA(Iph)RKWKK | 88 | 2058 |
| 07-3 | 20-23 | G(Pip)(Iph)(Pip)VVVHESNKWKK | 84 | 2059 |
| 07-3 | 20-23 | GPHG(Dab)G(Iph)(Dab)PV(Iph)KWKK | 73 | 2060 |
| 07-3 | 20-23 | GEHGGHK(Iph)PMFKWKK | 88 | 2063 |
| 07-3 | 20-23 | GTGNSR(Cit)HPN(Iph)KWKK | 84 | 2063 |
| 07-3 | 20-23 | GNGTNHR(Cit)A(Iph)NKWKK | 86 | 2064 |
| 07-3 | 20-23 | GP(Dab)AD(Iph)LKNK(Cit)KWKK | 90 | 2066 |
| 07-3 | 20-23 | GRFNKH(Cit)ESQLKKWK | 78 | 2066 |
| 07-3 | 20-23 | GDLNKAK(Cit)(Dab)V(Iph)KWKK | 73 | 2068 |
| 07-3 | 20-23 | G(Nap)DR(Iph)VNSG(Dab)TKWKK | 94 | 2069 |
| 07-3 | 20-23 | GKLFK(Iph)KVPADKWKK | 91 | 2069 |
| 07-3 | 20-23 | GNK(Dab)F(Cit)RTSD(Nap)KWKK | 88 | 2072 |
| 07-3 | 20-23 | GTRS(Dab)STDQR(Iph)KWKK | 89 | 2074 |
| 07-3 | 20-23 | G(Dab)(Iph)FATHPTFMKWKK | 92 | 2075 |
| 07-3 | 20-23 | GG(Iph)DLRNAH(Cit)NKWKK | 74 | 2077 |

| | | | | |
|---|---|---|---|---|
| 07-3 | 20-23 | GA(Pal)TGPNR(Iph)(Cit)HKWKK | 78 | 2081 |
| 07-3 | 20-23 | GDHR(Cit)GFRGG(Iph)KWKK | 81 | 2082 |
| 07-3 | 20-23 | GQLKTN(Iph)KTKVKWKK | 76 | 2083 |
| 07-3 | 20-23 | G(Nap)FSRERTKNVKWKK | 82 | 2084 |
| 07-3 | 20-23 | GRF(Cit)S(Iph)NPG(Dab)MKWKK | 95 | 2089 |
| 07-3 | 20-23 | G(Nap)GK(Iph)NK(Cit)PNGKWKK | 88 | 2092 |
| 07-3 | 20-23 | GTK(Cit)LSKHAE(Iph)KWKK | 83 | 2094 |
| 07-3 | 20-23 | GMALN(Cit)(Dab)(Nap)QR(Cit)KWKK | 79 | 2094 |
| 07-3 | 20-23 | GLSLFHQ(Dab)VQ(Iph)KWKK | 82 | 2095 |
| 07-3 | 20-23 | GFS(Iph)P(Pip)FSHA(Cit)KWKK | 85 | 2099 |
| 07-3 | 20-23 | GPNFK(Dab)KAD(Iph)(Cit)KWKK | 92 | 2100 |
| 07-3 | 20-23 | GQHNF(Nap)FVLNHKWKK | 85 | 2103 |
| 07-3 | 20-23 | G(Nap)TLP(Iph)KD(Dab)PLKWKK | 73 | 2104 |
| 07-3 | 20-23 | G(Cit)KDVVNR(Iph)GHKWKK | 91 | 2105 |
| 07-3 | 20-23 | GG(Iph)LRRTTPQ(Cit)KWKK | 80 | 2109 |
| 07-3 | 20-23 | GKGRA(Iph)FGQQ(Nap)KWKK | 94 | 2112 |
| 07-3 | 20-23 | GTRSR(Iph)VQFVTKWKK | 78 | 2117 |
| 07-3 | 20-23 | GNR(Cit)E(Cit)LHVQ(Cit)KWKK | 78 | 2117 |
| 07-3 | 20-23 | G(Nap)TH(Iph)EQAGTRKWKK | 90 | 2120 |
| 07-3 | 20-23 | G(Cit)GH(Cit)VRF(Cit)S(Nap)KWKK | 81 | 2121 |
| 07-3 | 20-23 | GRGF(Cit)(Iph)VHNLTKWKK | 94 | 2124 |
| 07-3 | 20-23 | GETVNF(Iph)NRSHKWKK | 78 | 2127 |
| 07-3 | 20-23 | GQHQET(Dab)PQH(Iph)KWKK | 83 | 2128 |
| 07-3 | 20-23 | G(Iph)KNT(Dab)G(Iph)FAPKWKK | 87 | 2131 |
| 07-3 | 20-23 | G(Cit)G(Dab)F(Iph)MFGH(Cit)KWKK | 85 | 2133 |
| 07-3 | 20-23 | GSLFTPS(Iph)(Cit)FRKWKK | 88 | 2135 |
| 07-3 | 20-23 | GTVHP(Iph)L(Dab)(Cit)(Nap)PKWKK | 80 | 2141 |
| 07-3 | 20-23 | G(Pip)G(Iph)SMKTP(Iph)VKWKK | 88 | 2142 |
| 07-3 | 20-23 | GS(Nap)(Cit)TN(Dab)FR(Cit)(Cit)KWKK | 86 | 2143 |
| 07-3 | 20-23 | GS(Cit)KLP(Nap)(Iph)P(Pip)TKWKK | 74 | 2146 |
| 07-3 | 20-23 | G(Nap)GTNHR(Cit)A(Iph)NKWKK | 75 | 2147 |
| 07-3 | 20-23 | G(Pal)RKQLTSS(Cit)(Iph)KWKK | 78 | 2148 |
| 07-3 | 20-23 | G(Nap)SSKMH(Iph)NQVKWKK | 92 | 2151 |
| 07-3 | 20-23 | GRL(Cit)(Dab)D(Cit)KPS(Iph)KWKK | 89 | 2153 |
| 07-3 | 20-23 | GKTSPD(Iph)(Iph)N(Dab)PKWKK | 72 | 2155 |
| 07-3 | 20-23 | GSN(Nap)T(Iph)(Pip)QPQHKWKK | 82 | 2158 |
| 07-3 | 20-23 | GPLSE(Iph)(Pip)(Iph)SHAKWKK | 96 | 2163 |
| 07-3 | 20-23 | GFSTR(Cit)GFRN(Iph)KWKK | 92 | 2165 |
| 07-3 | 20-23 | GRELGPR(Cit)(Iph)EKKWKK | 83 | 2165 |

| | | | | |
|---|---|---|---|---|
| 07-3 | 20-23 | GRQVP(Cit)(Iph)FLQ(Dab)KWKK | 85 | 2168 |
| 07-3 | 20-23 | GVPH(Nap)ARN(Cit)T(Iph)KWKK | 93 | 2172 |
| 07-3 | 20-23 | GTRR(Dab)(Iph)QEEQLKWKK | 88 | 2183 |
| 07-3 | 20-23 | GRSSLAR(Nap)FQ(Iph)KWKK | 76 | 2185 |
| 07-3 | 20-23 | GQAKES(Iph)QKA(Iph)KWKK | 73 | 2186 |
| 07-3 | 20-23 | GH(Nap)EAPRNQ(Nap)(Nap)KWKK | 79 | 2193 |
| 07-3 | 20-23 | GMNGNH(Cit)M(Cit)(Iph)RKWKK | 78 | 2197 |
| 07-3 | 20-23 | GHDKHANS(Iph)T(Iph)KWKK | 94 | 2206 |
| 07-3 | 20-23 | G(Iph)VEK(Cit)(Nap)NVNHKWKK | 83 | 2217 |
| 07-3 | 20-23 | G(Pip)(Cit)FTPS(Iph)(Cit)FRKWKK | 90 | 2218 |
| 07-3 | 20-23 | GLLQAQ(Iph)(Pal)K(Cit)(Nap)KWKK | 82 | 2226 |
| 07-3 | 20-23 | GV(Dab)(Nap)(Cit)(Cit)QLHT(Iph)KWKK | 78 | 2232 |
| 07-3 | 20-23 | GP(Cit)SQNH(Nap)(Cit)(Iph)KKWKK | 89 | 2245 |
| 07-3 | 20-23 | GFK(Iph)LKEQGQ(Iph)KWKK | 90 | 2274 |
| 07-3 | 20-23 | GFRDV(Iph)FHQ(Cit)FKWKK | 73 | 2276 |
| 07-3 | 20-23 | G(Dab)EH(Iph)(Dab)(Iph)(Cit)NHDKWKK | 87 | 2305 |
| 08-3 | 23-27 | G(Dab)GAS(Dab)TPSVHKWKK | 74 | 1706 |
| 08-3 | 23-27 | G(Pip)LSGTLH(Dab)SGKWKK | 85 | 1748 |
| 08-3 | 23-27 | GSKGAGPKMHSKWKK | 90 | 1750 |
| 08-3 | 23-27 | GGDLFALGLARKWKK | 74 | 1783 |
| 08-3 | 23-27 | GRESGHH(Dab)GASKWKK | 86 | 1788 |
| 08-3 | 23-27 | GNTVA(Dab)(Dab)(Dab)PS(Cit)KWKK | 72 | 1796 |
| 08-3 | 23-27 | GSGKQHM(Pip)GLSKWKK | 81 | 1821 |
| 08-3 | 23-27 | GG(Pip)VNHHVASQKWKK | 76 | 1825 |
| 08-3 | 23-27 | GALL(Nap)SHARGGKWKK | 74 | 1829 |
| 08-3 | 23-27 | GQ(Dab)(Nap)KA(Dab)PAGNKWKK | 83 | 1833 |
| 08-3 | 23-27 | GP(Dab)(Dab)HPLNFGTKWKK | 83 | 1833 |
| 08-3 | 23-27 | GAVGH(Nap)FGSTNKWKK | 90 | 1837 |
| 08-3 | 23-27 | G(Dab)KGHKASDS(Cit)KWKK | 88 | 1837 |
| 08-3 | 23-27 | GGSSTVRNTK(Cit)KWKK | 80 | 1857 |
| 08-3 | 23-27 | G(Dab)GFSQEKRG(Dab)KWKK | 81 | 1859 |
| 08-3 | 23-27 | GKVG(Dab)G(Cit)NHS(Cit)KWKK | 80 | 1863 |
| 08-3 | 23-27 | G(Dab)QGVNFGHTRKWKK | 94 | 1866 |
| 08-3 | 23-27 | G(Pip)VV(Cit)HNGHGLKWKK | 93 | 1866 |
| 08-3 | 23-27 | GLF(Pip)GVTKSHTKWKK | 98 | 1866 |
| 08-3 | 23-27 | GLRFAGNSSHKKWKK | 80 | 1867 |
| 08-3 | 23-27 | GKNKHGPVFQSKWKK | 81 | 1892 |
| 08-3 | 23-27 | GVDSKGHHETHKWKK | 86 | 1897 |
| 08-3 | 23-27 | GRNFTLMK(Pip)GGKWKK | 74 | 1900 |

| | | | | |
|---|---|---|---|---|
| 08-3 | 23-27 | GNFHARV(Cit)HGGKWKK | 87 | 1902 |
| 08-3 | 23-27 | G(Iph)A(Dab)LFKTSGGKWKK | 91 | 1904 |
| 08-3 | 23-27 | GSLRLPSEKKPKWKK | 77 | 1905 |
| 08-3 | 23-27 | GTVPGSLAT(Iph)HKWKK | 88 | 1906 |
| 08-3 | 23-27 | G(Dab)(Dab)SFTLTNFKKWKK | 90 | 1908 |
| 08-3 | 23-27 | GH(Pip)(Dab)(Iph)SGGG(Dab)FKWKK | 90 | 1911 |
| 08-3 | 23-27 | GGAN(Dab)HHAFL(Nap)KWKK | 75 | 1914 |
| 08-3 | 23-27 | GQGG(Dab)HHAFL(Nap)KWKK | 83 | 1914 |
| 08-3 | 23-27 | GFTKTFG(Dab)NHLKWKK | 97 | 1915 |
| 08-3 | 23-27 | GRVM(Nap)HAGTP(Dab)KWKK | 79 | 1916 |
| 08-3 | 23-27 | G(Pip)NLTFFKN(Dab)GKWKK | 71 | 1917 |
| 08-3 | 23-27 | GKPQTFFKN(Dab)GKWKK | 92 | 1917 |
| 08-3 | 23-27 | G(Pal)NKFGLALHKKWKK | 98 | 1926 |
| 08-3 | 23-27 | GFHA(Nap)KAGG(Cit)HKWKK | 79 | 1929 |
| 08-3 | 23-27 | GGNASEFLRERKWKK | 73 | 1929 |
| 08-3 | 23-27 | G(Pip)L(Cit)V(Dab)LSVQHKWKK | 71 | 1929 |
| 08-3 | 23-27 | GEA(Dab)TAM(Pip)HH(Cit)KWKK | 71 | 1930 |
| 08-3 | 23-27 | GPG(Pip)(Iph)AKGVQEKWKK | 80 | 1935 |
| 08-3 | 23-27 | GT(Nap)FKAHLDVGKWKK | 82 | 1935 |
| 08-3 | 23-27 | GSLSKFFVEK(Dab)KWKK | 88 | 1935 |
| 08-3 | 23-27 | G(Nap)LSLKS(Dab)RDAKWKK | 79 | 1937 |
| 08-3 | 23-27 | GEVSGK(Cit)VFHKKWKK | 95 | 1938 |
| 08-3 | 23-27 | G(Iph)(Dab)S(Dab)GA(Nap)GLNKWKK | 88 | 1939 |
| 08-3 | 23-27 | GKFPLEAN(Dab)RNKWKK | 87 | 1939 |
| 08-3 | 23-27 | GSNN(Dab)VRNTK(Cit)KWKK | 93 | 1940 |
| 08-3 | 23-27 | G(Iph)(Dab)LAGF(Dab)LVTKWKK | 90 | 1944 |
| 08-3 | 23-27 | GLETNTHHKDTKWKK | 88 | 1946 |
| 08-3 | 23-27 | GDKPPS(Nap)PAHRKWKK | 78 | 1952 |
| 08-3 | 23-27 | GKA(Pal)SQHNSFHKWKK | 72 | 1954 |
| 08-3 | 23-27 | GPLHSQHNSFHKWKK | 85 | 1954 |
| 08-3 | 23-27 | GQQSKLF(Cit)HA(Dab)KWKK | 89 | 1966 |
| 08-3 | 23-27 | G(Dab)STN(Nap)RS(Dab)NFKWKK | 95 | 1973 |
| 08-3 | 23-27 | GSVAQQHFHRLKWKK | 74 | 1973 |
| 08-3 | 23-27 | GGKLLTH(Cit)SL(Nap)KWKK | 87 | 1973 |
| 08-3 | 23-27 | GVHK(Cit)LNV(Dab)T(Cit)KWKK | 93 | 1975 |
| 08-3 | 23-27 | GT(Cit)AFLVHRKTKWKK | 79 | 1980 |
| 08-3 | 23-27 | GDPKMKKTHFVKWKK | 75 | 1981 |
| 08-3 | 23-27 | GKVPG(Cit)NQFKRKWKK | 94 | 1981 |
| 08-3 | 23-27 | GNKS(Dab)AL(Iph)RGNKWKK | 93 | 1983 |

| | | | | |
|---|---|---|---|---|
| 08-3 | 23-27 | GSRFQERV(Dab)NVKWKK | 91 | 1985 |
| 08-3 | 23-27 | GQPM(Dab)FKTNLRKWKK | 82 | 1985 |
| 08-3 | 23-27 | G(Dab)PTAQ(Dab)AKF(Iph)KWKK | 77 | 1986 |
| 08-3 | 23-27 | GTV(Dab)HSLAT(Iph)HKWKK | 90 | 1989 |
| 08-3 | 23-27 | GV(Nap)RVDKATPRKWKK | 83 | 1989 |
| 08-3 | 23-27 | GV(Nap)RVDKAPTRKWKK | 89 | 1989 |
| 08-3 | 23-27 | G(Pip)QN(Dab)(Nap)KVTLNKWKK | 77 | 1990 |
| 08-3 | 23-27 | GRGHFFLVF(Dab)QKWKK | 82 | 2001 |
| 08-3 | 23-27 | GLGQA(Nap)ARHF(Cit)KWKK | 77 | 2004 |
| 08-3 | 23-27 | G(Cit)KSH(Dab)(Cit)AS(Nap)NKWKK | 73 | 2005 |
| 08-3 | 23-27 | G(Cit)KSH(Dab)(Cit)AS(Nap)NKWKK | 95 | 2005 |
| 08-3 | 23-27 | GHGRKD(Nap)QALHKWKK | 90 | 2009 |
| 08-3 | 23-27 | GKGVT(Iph)KRANLKWKK | 71 | 2010 |
| 08-3 | 23-27 | GK(Nap)HADQLSHKKWKK | 77 | 2011 |
| 08-3 | 23-27 | G(Pip)(Cit)AAKFQLFRKWKK | 87 | 2014 |
| 08-3 | 23-27 | GRKQ(Dab)(Dab)G(Iph)SESKWKK | 83 | 2015 |
| 08-3 | 23-27 | GHTN(Cit)(Cit)SLHKNKWKK | 94 | 2015 |
| 08-3 | 23-27 | GFQ(Dab)H(Cit)LGH(Cit)NKWKK | 87 | 2017 |
| 08-3 | 23-27 | GNQH(Dab)(Nap)QHVLPKWKK | 75 | 2020 |
| 08-3 | 23-27 | G(Iph)(Pip)AGP(Iph)RAAGKWKK | 80 | 2022 |
| 08-3 | 23-27 | G(Dab)RA(Pal)GL(Iph)FSTKWKK | 91 | 2023 |
| 08-3 | 23-27 | GKP(Dab)(Cit)HHE(Pip)QNKWKK | 72 | 2023 |
| 08-3 | 23-27 | GRKEMVNEHPHKWKK | 93 | 2027 |
| 08-3 | 23-27 | GHQQKAPARA(Iph)KWKK | 73 | 2030 |
| 08-3 | 23-27 | GS(Nap)VHHHQQPQKWKK | 87 | 2045 |
| 08-3 | 23-27 | GFSQ(Pal)TR(Dab)V(Nap)LKWKK | 82 | 2046 |
| 08-3 | 23-27 | GKFA(Cit)FTHSFRKWKK | 97 | 2048 |
| 08-3 | 23-27 | G(Nap)S(Nap)GH(Cit)NPHVKWKK | 84 | 2049 |
| 08-3 | 23-27 | G(Cit)HFAT(Nap)QR(Dab)SKWKK | 73 | 2051 |
| 08-3 | 23-27 | GN(Iph)SETNLKARKWKK | 82 | 2056 |
| 08-3 | 23-27 | GRLKPTH(Cit)SL(Nap)KWKK | 93 | 2056 |
| 08-3 | 23-27 | GANR(Nap)APEKK(Nap)KWKK | 90 | 2058 |
| 08-3 | 23-27 | GR(Nap)TEPAKPR(Cit)KWKK | 89 | 2059 |
| 08-3 | 23-27 | G(Pal)AQG(Dab)FRL(Iph)PKWKK | 91 | 2060 |
| 08-3 | 23-27 | GE(Iph)KTKTS(Dab)S(Cit)KWKK | 90 | 2061 |
| 08-3 | 23-27 | GPKAFRGQH(Iph)TKWKK | 97 | 2065 |
| 08-3 | 23-27 | GT(Dab)(Dab)FR(Nap)N(Cit)TKKWKK | 84 | 2071 |
| 08-3 | 23-27 | G(Nap)PNN(Iph)GHNT(Dab)KWKK | 81 | 2074 |
| 08-3 | 23-27 | GNSESH(Iph)HP(Dab)FKWKK | 85 | 2078 |

| | | | | |
|---|---|---|---|---|
| 08-3 | 23-27 | G(Nap)KQ(Dab)GSK(Iph)LPKWKK | 87 | 2078 |
| 08-3 | 23-27 | GFNAPKL(Iph)TRLKWKK | 83 | 2083 |
| 08-3 | 23-27 | GPR(Pal)S(Dab)A(Iph)RLNKWKK | 71 | 2085 |
| 08-3 | 23-27 | G(Pal)EPT(Iph)(Dab)PVKFKWKK | 72 | 2089 |
| 08-3 | 23-27 | GK(Pal)S(Dab)NS(Iph)(Cit)VKKWKK | 80 | 2091 |
| 08-3 | 23-27 | GGMLTKERV(Iph)HKWKK | 84 | 2094 |
| 08-3 | 23-27 | GKVVR(Dab)P(Cit)(Iph)TNKWKK | 92 | 2094 |
| 08-3 | 23-27 | GHDRKKMT(Cit)FQKWKK | 96 | 2098 |
| 08-3 | 23-27 | GHSHH(Iph)PHPVEKWKK | 72 | 2100 |
| 08-3 | 23-27 | G(Dab)(Iph)AGG(Dab)FRT(Iph)KWKK | 76 | 2105 |
| 08-3 | 23-27 | GNRG(Nap)(Cit)(Iph)PGRAKWKK | 79 | 2105 |
| 08-3 | 23-27 | GKQ(Nap)NAT(Dab)P(Iph)KKWKK | 77 | 2107 |
| 08-3 | 23-27 | GR(Nap)SMHTF(Cit)TKKWKK | 71 | 2112 |
| 08-3 | 23-27 | GTKR(Dab)(Cit)GDE(Iph)KKWKK | 85 | 2114 |
| 08-3 | 23-27 | GNQNLK(Iph)LQHVKWKK | 76 | 2117 |
| 08-3 | 23-27 | G(Pip)KSGAF(Iph)(Iph)EGKWKK | 94 | 2118 |
| 08-3 | 23-27 | G(Nap)(Cit)SGS(Pip)RR(Nap)QKWKK | 83 | 2118 |
| 08-3 | 23-27 | GLT(Iph)(Iph)(Pal)N(Dab)GLGKWKK | 79 | 2119 |
| 08-3 | 23-27 | GFR(Pip)AEG(Iph)P(Nap)TKWKK | 88 | 2124 |
| 08-3 | 23-27 | GAKLQQKKT(Iph)(Cit)KWKK | 76 | 2125 |
| 08-3 | 23-27 | GA(Iph)FNNRKAS(Nap)KWKK | 87 | 2128 |
| 08-3 | 23-27 | GHKLFPQGRM(Iph)KWKK | 87 | 2137 |
| 08-3 | 23-27 | G(Nap)(Iph)FQEARGPNKWKK | 78 | 2139 |
| 08-3 | 23-27 | GFVQ(Dab)G(Iph)(Dab)PV(Iph)KWKK | 75 | 2143 |
| 08-3 | 23-27 | GQ(Iph)RTGGQ(Cit)(Nap)(Pip)KWKK | 83 | 2150 |
| 08-3 | 23-27 | G(Cit)MK(Nap)AGRD(Iph)TKWKK | 75 | 2156 |
| 08-3 | 23-27 | GRVKREESPH(Iph)KWKK | 91 | 2161 |
| 08-3 | 23-27 | GP(Iph)VN(Nap)RAFDKKWKK | 72 | 2167 |
| 08-3 | 23-27 | GFHKGFQ(Cit)HS(Iph)KWKK | 92 | 2168 |
| 08-3 | 23-27 | GPEGT(Nap)DKFR(Iph)KWKK | 80 | 2170 |
| 08-3 | 23-27 | GL(Pip)(Iph)ST(Nap)LHKEKWKK | 83 | 2174 |
| 08-3 | 23-27 | G(Iph)(Dab)M(Iph)GTSQEKKWKK | 71 | 2177 |
| 08-3 | 23-27 | GASVR(Iph)HGKE(Iph)KWKK | 77 | 2180 |
| 08-3 | 23-27 | GPE(Iph)P(Iph)GRSFVKWKK | 76 | 2185 |
| 08-3 | 23-27 | GRG(Iph)LVTP(Iph)QKKWKK | 88 | 2195 |
| 08-3 | 23-27 | G(Iph)G(Dab)(Cit)(Iph)ESHGRKWKK | 71 | 2196 |
| 08-3 | 23-27 | G(Nap)EKSPKNRM(Iph)KWKK | 79 | 2210 |
| 08-3 | 23-27 | G(Iph)(Iph)SKETRALNKWKK | 91 | 2215 |
| 08-3 | 23-27 | GFSRV(Cit)(Iph)LERQKWKK | 90 | 2215 |

| | | | | |
|---|---|---|---|---|
| 08-3 | 23-27 | GTRSRF(Iph)(Nap)D(Pip)TKWKK | 80 | 2229 |
| 08-3 | 23-27 | G(Iph)(Dab)QA(Iph)NQRAFKWKK | 97 | 2231 |
| 08-3 | 23-27 | GH(Pal)H(Iph)FPTR(Cit)NKWKK | 77 | 2237 |
| 08-3 | 23-27 | G(Nap)VR(Iph)TNNQR(Pal)KWKK | 75 | 2256 |
| 08-3 | 23-27 | GE(Dab)VS(Cit)N(Dab)(Iph)R(Iph)KWKK | 77 | 2258 |
| 08-3 | 23-27 | GPDR(Iph)GFK(Iph)QKKWKK | 77 | 2272 |
| 08-3 | 23-27 | G(Iph)(Iph)QRTAQNFNKWKK | 72 | 2275 |
| 08-3 | 23-27 | GN(Dab)GS(Iph)HT(Iph)P(Iph)KWKK | 88 | 2282 |
| 08-3 | 23-27 | G(Cit)E(Dab)N(Iph)SQK(Iph)HKWKK | 80 | 2296 |
| 08-3 | 23-27 | G(Iph)R(Iph)VR(Cit)(Cit)LQSKWKK | 90 | 2369 |
| 09-3 | 27-35 | GDVKGGGAKFHKWKK | 87 | 1766 |
| 09-3 | 27-35 | GAQ(Nap)VGG(Dab)E(Dab)VKWKK | 80 | 1807 |
| 09-3 | 27-35 | GVAG(Dab)GV(Nap)FEPKWKK | 74 | 1823 |
| 09-3 | 27-35 | GPRAGRLSPARKWKK | 85 | 1831 |
| 09-3 | 27-35 | G(Dab)KSAPKVQKTKWKK | 82 | 1837 |
| 09-3 | 27-35 | GVNDKQVGMTLKWKK | 85 | 1855 |
| 09-3 | 27-35 | G(Dab)VKQ(Dab)ANNHPKWKK | 83 | 1858 |
| 09-3 | 27-35 | GLSTGVHKPNRKWKK | 72 | 1859 |
| 09-3 | 27-35 | GHAHTQH(Dab)LGLKWKK | 96 | 1864 |
| 09-3 | 27-35 | G(Dab)NQKSSHKTPKWKK | 84 | 1877 |
| 09-3 | 27-35 | G(Nap)TVA(Dab)(Dab)(Dab)PS(Cit)KWKK | 90 | 1879 |
| 09-3 | 27-35 | GKARPPVGFKHKWKK | 84 | 1887 |
| 09-3 | 27-35 | GTSGLRLH(Dab)(Nap)GKWKK | 84 | 1888 |
| 09-3 | 27-35 | GHQA(Dab)MPQAKKKWKK | 93 | 1889 |
| 09-3 | 27-35 | GGAKK(Cit)RNSEPKWKK | 89 | 1894 |
| 09-3 | 27-35 | GNKFGR(Dab)LTD(Dab)KWKK | 80 | 1901 |
| 09-3 | 27-35 | GLTHG(Dab)(Nap)RA(Dab)VKWKK | 75 | 1901 |
| 09-3 | 27-35 | GLASA(Nap)HSKKNKWKK | 86 | 1903 |
| 09-3 | 27-35 | GVRSLVTQGRHKWKK | 87 | 1903 |
| 09-3 | 27-35 | GARHRGAMQVKKWKK | 81 | 1904 |
| 09-3 | 27-35 | GFLKKPVVGL(Cit)KWKK | 85 | 1908 |
| 09-3 | 27-35 | GSRRNLKGHSQKWKK | 96 | 1933 |
| 09-3 | 27-35 | GAV(Iph)A(Dab)QART(Dab)KWKK | 93 | 1940 |
| 09-3 | 27-35 | GPVQKEKSF(Dab)HKWKK | 91 | 1950 |
| 09-3 | 27-35 | G(Nap)MSKSLPSR(Dab)KWKK | 75 | 1953 |
| 09-3 | 27-35 | GHREKHANFVAKWKK | 91 | 1959 |
| 09-3 | 27-35 | GHR(Dab)VP(Iph)(Dab)(Dab)AGKWKK | 74 | 1960 |
| 09-3 | 27-35 | GSRKKQHLLVTKWKK | 86 | 1960 |
| 09-3 | 27-35 | GAVN(Cit)ATRLRRKWKK | 84 | 1964 |

| | | | | |
|---|---|---|---|---|
| 09-3 | 27-35 | G(Iph)VAVKSNP(Dab)KKWKK | 86 | 1966 |
| 09-3 | 27-35 | GF(Pal)GV(Dab)GF(Dab)(Iph)AKWKK | 72 | 1969 |
| 09-3 | 27-35 | GKG(Cit)TK(Nap)AKHTKWKK | 98 | 1975 |
| 09-3 | 27-35 | G(Pip)RRPLPSARFKWKK | 90 | 1976 |
| 09-3 | 27-35 | GRKLKPNKTELKWKK | 90 | 1977 |
| 09-3 | 27-35 | GKD(Nap)QAARRSTKWKK | 73 | 1980 |
| 09-3 | 27-35 | G(Nap)KADKTTLKKKWKK | 98 | 1980 |
| 09-3 | 27-35 | GL(Pip)NFT(Cit)SK(Pip)NKWKK | 91 | 1983 |
| 09-3 | 27-35 | GKQLHFQT(Dab)HPKWKK | 91 | 1986 |
| 09-3 | 27-35 | GTAE(Cit)VF(Cit)Q(Dab)KKWKK | 75 | 1987 |
| 09-3 | 27-35 | GHPS(Nap)STHF(Dab)KKWKK | 89 | 1988 |
| 09-3 | 27-35 | GQSKKKPVRL(Cit)KWKK | 94 | 1991 |
| 09-3 | 27-35 | GKNKGVR(Dab)FFFKWKK | 86 | 1993 |
| 09-3 | 27-35 | GVF(Dab)(Dab)(Iph)SGN(Dab)FKWKK | 92 | 1994 |
| 09-3 | 27-35 | GAA(Iph)PKMG(Dab)(Nap)(Dab)KWKK | 75 | 1995 |
| 09-3 | 27-35 | GDHTPVV(Cit)RHKKWKK | 90 | 1996 |
| 09-3 | 27-35 | G(Iph)NSSASPR(Cit)VKWKK | 79 | 1998 |
| 09-3 | 27-35 | GVQRMHQTKPKKWKK | 74 | 2003 |
| 09-3 | 27-35 | GAVGEF(Nap)LFFKKWKK | 87 | 2005 |
| 09-3 | 27-35 | GREKQ(Dab)PTNQRKWKK | 86 | 2007 |
| 09-3 | 27-35 | GSPHGRTKAH(Iph)KWKK | 88 | 2014 |
| 09-3 | 27-35 | GRQFL(Dab)PDRSRKWKK | 86 | 2025 |
| 09-3 | 27-35 | GDAL(Iph)GKDTKRKWKK | 85 | 2027 |
| 09-3 | 27-35 | GKQEFN(Dab)LF(Cit)VKWKK | 87 | 2032 |
| 09-3 | 27-35 | GNG(Pip)VA(Iph)TFFKKWKK | 91 | 2033 |
| 09-3 | 27-35 | G(Iph)LS(Dab)QPH(Dab)(Cit)AKWKK | 77 | 2033 |
| 09-3 | 27-35 | GPQGN(Nap)F(Nap)VRAKWKK | 77 | 2033 |
| 09-3 | 27-35 | GSHG(Nap)LQRFSRKWKK | 90 | 2035 |
| 09-3 | 27-35 | G(Pal)LL(Pip)(Pip)QFLNHKWKK | 79 | 2035 |
| 09-3 | 27-35 | G(Dab)(Iph)KSDVG(Cit)KKKWKK | 84 | 2042 |
| 09-3 | 27-35 | G(Pip)Q(Cit)SHVFHKQKWKK | 80 | 2044 |
| 09-3 | 27-35 | G(Pip)GH(Iph)TGRKELKWKK | 76 | 2047 |
| 09-3 | 27-35 | G(Cit)KSPT(Nap)SRNRKWKK | 91 | 2050 |
| 09-3 | 27-35 | GFHPKFQLKVRKWKK | 77 | 2050 |
| 09-3 | 27-35 | G(Iph)SRAPALREKKWKK | 80 | 2051 |
| 09-3 | 27-35 | GRH(Cit)SD(Dab)PF(Cit)KKWKK | 92 | 2051 |
| 09-3 | 27-35 | GRHHKA(Cit)(Nap)DPSKWKK | 84 | 2052 |
| 09-3 | 27-35 | GRG(Nap)SKN(Cit)LHHKWKK | 98 | 2053 |
| 09-3 | 27-35 | GVFHLL(Dab)(Nap)KP(Cit)KWKK | 72 | 2058 |

| | | | | |
|---|---|---|---|---|
| 09-3 | 27-35 | GESRTVRRFFLKWKK | 90 | 2061 |
| 09-3 | 27-35 | GVNVGDRFH(Iph)(Dab)KWKK | 83 | 2067 |
| 09-3 | 27-35 | GLT(Dab)S(Cit)SS(Iph)RHKWKK | 95 | 2068 |
| 09-3 | 27-35 | GV(Cit)S(Nap)(Dab)F(Cit)Q(Dab)KKWKK | 89 | 2070 |
| 09-3 | 27-35 | GKRDNQ(Dab)VF(Cit)RKWKK | 91 | 2070 |
| 09-3 | 27-35 | G(Dab)ERSR(Dab)QF(Nap)TKWKK | 86 | 2071 |
| 09-3 | 27-35 | GHPTT(Nap)RDRLKKWKK | 76 | 2071 |
| 09-3 | 27-35 | GK(Cit)(Dab)P(Nap)L(Dab)VL(Nap)KWKK | 73 | 2071 |
| 09-3 | 27-35 | GHSKARHGQ(Iph)QKWKK | 90 | 2072 |
| 09-3 | 27-35 | GQ(Iph)(Dab)A(Nap)T(Dab)PVHKWKK | 78 | 2073 |
| 09-3 | 27-35 | GL(Nap)SSHRHQNFKWKK | 90 | 2073 |
| 09-3 | 27-35 | GNL(Nap)(Nap)GFKPRVKWKK | 77 | 2075 |
| 09-3 | 27-35 | GV(Iph)NNP(Dab)LL(Cit)KKWKK | 83 | 2078 |
| 09-3 | 27-35 | GPENPRSTR(Iph)(Dab)KWKK | 75 | 2080 |
| 09-3 | 27-35 | G(Pip)(Cit)HLRKE(Cit)LPKWKK | 89 | 2083 |
| 09-3 | 27-35 | GRLLSTQ(Iph)RS(Dab)KWKK | 77 | 2084 |
| 09-3 | 27-35 | GFL(Iph)SDHVGRHKWKK | 88 | 2091 |
| 09-3 | 27-35 | G(Iph)ASVRSKSH(Nap)KWKK | 92 | 2092 |
| 09-3 | 27-35 | GH(Iph)TSNARR(Cit)AKWKK | 89 | 2093 |
| 09-3 | 27-35 | GNS(Nap)KRFQ(Cit)(Dab)LKWKK | 84 | 2097 |
| 09-3 | 27-35 | G(Iph)MFGGQR(Cit)(Dab)QKWKK | 83 | 2104 |
| 09-3 | 27-35 | G(Nap)KNLE(Iph)GPK(Dab)KWKK | 76 | 2106 |
| 09-3 | 27-35 | GV(Nap)(Dab)QTN(Cit)RFHKWKK | 98 | 2106 |
| 09-3 | 27-35 | GVK(Iph)KH(Cit)SRGDKWKK | 84 | 2107 |
| 09-3 | 27-35 | G(Dab)R(Dab)(Nap)P(Iph)SNNTKWKK | 86 | 2109 |
| 09-3 | 27-35 | GSRG(Iph)NLFRTHKWKK | 83 | 2111 |
| 09-3 | 27-35 | GHQ(Dab)N(Nap)F(Nap)VRAKWKK | 78 | 2116 |
| 09-3 | 27-35 | GLHF(Dab)(Cit)(Cit)LREHKWKK | 74 | 2116 |
| 09-3 | 27-35 | GPNA(Cit)TR(Iph)KHNKWKK | 80 | 2118 |
| 09-3 | 27-35 | G(Pal)RPQDR(Dab)QK(Nap)KWKK | 76 | 2123 |
| 09-3 | 27-35 | G(Pal)NSHSFT(Iph)HKKWKK | 80 | 2129 |
| 09-3 | 27-35 | GRNKFNK(Iph)ELGKWKK | 94 | 2129 |
| 09-3 | 27-35 | GQ(Cit)LHV(Iph)KP(Pip)TKWKK | 84 | 2129 |
| 09-3 | 27-35 | G(Pal)F(Iph)(Nap)GVFSNTKWKK | 80 | 2140 |
| 09-3 | 27-35 | G(Pal)R(Dab)DF(Pip)ATH(Iph)KWKK | 76 | 2144 |
| 09-3 | 27-35 | G(Iph)NKRVHLQLLKWKK | 94 | 2144 |
| 09-3 | 27-35 | GNSQL(Iph)KERQKKWKK | 83 | 2154 |
| 09-3 | 27-35 | GHH(Nap)RQVNTK(Nap)KWKK | 97 | 2164 |
| 09-3 | 27-35 | G(Iph)G(Dab)RQTQ(Dab)R(Nap)KWKK | 83 | 2166 |

| | | | | |
|---|---|---|---|---|
| 09-3 | 27-35 | GRRVF(Nap)GN(Dab)T(Iph)KWKK | 84 | 2170 |
| 09-3 | 27-35 | G(Iph)NRST(Nap)LHKLKWKK | 87 | 2189 |
| 09-3 | 27-35 | GML(Cit)(Iph)(Dab)(Nap)(Cit)P(Dab)VKWK K | 77 | 2194 |
| 09-3 | 27-35 | GRHANQQK(Iph)L(Nap)KWKK | 92 | 2215 |
| 09-3 | 27-35 | GHH(Iph)S(Cit)HK(Cit)NQKWKK | 96 | 2225 |
| 09-3 | 27-35 | G(Dab)KS(Iph)DH(Cit)(Dab)(Iph)PKWKK | 79 | 2237 |
| 09-3 | 27-35 | GQL(Pal)(Pal)EF(Iph)RKTKWKK | 87 | 2241 |
| 09-3 | 27-35 | GVK(Nap)(Dab)(Iph)KRFDEKWKK | 77 | 2242 |
| 09-3 | 27-35 | GK(Iph)KRGQL(Iph)FSKWKK | 74 | 2260 |
| 09-3 | 27-35 | GRTHG(Cit)(Dab)FV(Iph)(Iph)KWKK | 85 | 2270 |
| 09-3 | 27-35 | GLQ(Iph)SS(Dab)(Cit)R(Iph)HKWKK | 87 | 2281 |
| 09-3 | 27-35 | GV(Nap)RP(Nap)R(Cit)NA(Iph)KWKK | 77 | 2287 |
| 09-3 | 27-35 | GRDRV(Iph)KRP(Iph)PKWKK | 88 | 2320 |
| 09-3 | 27-35 | GR(Iph)HKQ(Iph)(Dab)ESFKWKK | 71 | 2328 |
| 10-3 | 35-55 | G(Dab)VARNN(Dab)TKGKWKK | 91 | 1810 |
| 10-3 | 35-55 | GRPGAPQTK(Dab)HKWKK | 72 | 1842 |
| 10-3 | 35-55 | GV(Dab)LTG(Dab)ARFHKWKK | 87 | 1851 |
| 10-3 | 35-55 | GNARHP(Dab)PARSKWKK | 79 | 1856 |
| 10-3 | 35-55 | GASNGKL(Dab)K(Dab)(Nap)KWKK | 78 | 1865 |
| 10-3 | 35-55 | GVFKG(Pip)RSVT(Dab)KWKK | 96 | 1870 |
| 10-3 | 35-55 | GSTAALH(Dab)A(Nap)RKWKK | 76 | 1874 |
| 10-3 | 35-55 | GVGTAHRQN(Dab)RKWKK | 89 | 1889 |
| 10-3 | 35-55 | GPVN(Cit)(Dab)K(Cit)VAPKWKK | 77 | 1889 |
| 10-3 | 35-55 | GN(Cit)KKF(Dab)G(Dab)(Pip)AKWKK | 82 | 1898 |
| 10-3 | 35-55 | GRTP(Pip)GAAFRRKWKK | 93 | 1908 |
| 10-3 | 35-55 | GFNK(Dab)KVKKPAKWKK | 93 | 1910 |
| 10-3 | 35-55 | GS(Pip)(Nap)G(Dab)SHNH(Dab)KWKK | 71 | 1912 |
| 10-3 | 35-55 | GAG(Nap)SHVTKHHKWKK | 79 | 1921 |
| 10-3 | 35-55 | GAAKKDHHPRNKWKK | 87 | 1924 |
| 10-3 | 35-55 | G(Iph)(Dab)SQVKGKG(Dab)KWKK | 95 | 1927 |
| 10-3 | 35-55 | GVEFSHKKKKGKWKK | 94 | 1938 |
| 10-3 | 35-55 | G(Dab)TARRNSFKNKWKK | 72 | 1944 |
| 10-3 | 35-55 | G(Dab)KSTFRL(Dab)NKKWKK | 97 | 1944 |
| 10-3 | 35-55 | GNQ(Dab)ALH(Dab)A(Nap)RKWKK | 84 | 1957 |
| 10-3 | 35-55 | GG(Pip)VKG(Pip)H(Iph)SVKWKK | 71 | 1959 |
| 10-3 | 35-55 | GG(Pip)VKGH(Pip)(Iph)VSKWKK | 82 | 1959 |
| 10-3 | 35-55 | G(Dab)VAKET(Dab)K(Nap)HKWKK | 79 | 1960 |
| 10-3 | 35-55 | GKTNRAHQAHRKWKK | 83 | 1969 |

| | | | | |
|---|---|---|---|---|
| 10-3 | 35-55 | G(Pip)GAKT(Dab)FFQ(Nap)KWKK | 90 | 1972 |
| 10-3 | 35-55 | GNA(Dab)(Nap)VPN(Dab)RRKWKK | 79 | 1974 |
| 10-3 | 35-55 | GPHQNSHKHHLKWKK | 96 | 1985 |
| 10-3 | 35-55 | G(Iph)(Dab)ALGA(Cit)(Dab)HHKWKK | 85 | 1986 |
| 10-3 | 35-55 | G(Dab)G(Iph)L(Dab)ADH(Dab)RKWKK | 75 | 1992 |
| 10-3 | 35-55 | GTG(Iph)DKGHARKKWKK | 84 | 1993 |
| 10-3 | 35-55 | G(Pip)RSGRRDS(Cit)HKWKK | 96 | 2004 |
| 10-3 | 35-55 | GLK(Pal)KRNGKHEKWKK | 78 | 2008 |
| 10-3 | 35-55 | GREHKQKSP(Cit)(Dab)KWKK | 88 | 2017 |
| 10-3 | 35-55 | GKLTRKHELLEKWKK | 82 | 2017 |
| 10-3 | 35-55 | GLKKHRNF(Dab)NLKWKK | 97 | 2020 |
| 10-3 | 35-55 | GDV(Pip)PGKRSN(Iph)KWKK | 79 | 2022 |
| 10-3 | 35-55 | GVFKN(Dab)N(Nap)PR(Dab)KWKK | 83 | 2022 |
| 10-3 | 35-55 | G(Dab)(Nap)(Nap)ATV(Dab)SFRKWKK | 84 | 2025 |
| 10-3 | 35-55 | GQ(Cit)Q(Dab)KP(Dab)FRNKWKK | 81 | 2025 |
| 10-3 | 35-55 | GKPQFHSFRK(Dab)KWKK | 93 | 2025 |
| 10-3 | 35-55 | GRASLKF(Dab)(Cit)(Nap)(Dab)KWKK | 94 | 2026 |
| 10-3 | 35-55 | GQRARRNSFKNKWKK | 88 | 2027 |
| 10-3 | 35-55 | GP(Cit)(Nap)Q(Dab)AKRKTKWKK | 82 | 2033 |
| 10-3 | 35-55 | GNTE(Dab)VG(Dab)(Iph)HRKWKK | 90 | 2036 |
| 10-3 | 35-55 | GKRKLQKQRFGKWKK | 90 | 2039 |
| 10-3 | 35-55 | GKNL(Iph)VNHHTGKWKK | 86 | 2043 |
| 10-3 | 35-55 | GTTRNNHKR(Nap)AKWKK | 87 | 2045 |
| 10-3 | 35-55 | GAKQKT(Dab)FFQ(Nap)KWKK | 85 | 2045 |
| 10-3 | 35-55 | GRRNNLHQAHRKWKK | 98 | 2052 |
| 10-3 | 35-55 | GVF(Pip)GA(Iph)RFGRKWKK | 95 | 2059 |
| 10-3 | 35-55 | GDKGVHRKLS(Iph)KWKK | 87 | 2063 |
| 10-3 | 35-55 | GGF(Iph)(Pip)G(Dab)LERHKWKK | 80 | 2065 |
| 10-3 | 35-55 | GQAHARRF(Iph)(Dab)GKWKK | 90 | 2066 |
| 10-3 | 35-55 | GRRDDKHHL(Dab)(Pal)KWKK | 85 | 2075 |
| 10-3 | 35-55 | GV(Iph)THFGHARKKWKK | 93 | 2076 |
| 10-3 | 35-55 | GLKQFSF(Nap)(Dab)KHKWKK | 91 | 2082 |
| 10-3 | 35-55 | G(Nap)(Iph)P(Pip)NPAGKRKWKK | 92 | 2086 |
| 10-3 | 35-55 | G(Iph)TSKHARVNRKWKK | 94 | 2092 |
| 10-3 | 35-55 | GGK(Cit)SAP(Nap)(Iph)(Dab)RKWKK | 81 | 2093 |
| 10-3 | 35-55 | G(Iph)HKH(Dab)P(Pip)DPNKWKK | 88 | 2094 |
| 10-3 | 35-55 | G(Dab)(Dab)KGAP(Iph)R(Nap)FKWKK | 72 | 2096 |
| 10-3 | 35-55 | GRSRNKGVQ(Iph)KKWKK | 97 | 2096 |
| 10-3 | 35-55 | GSGR(Cit)KF(Pip)(Dab)(Iph)TKWKK | 82 | 2102 |

| | | | | |
|---|---|---|---|---|
| 10-3 | 35-55 | GSGR(Cit)KF(Pip)(Dab)(Iph)TKWKK | 82 | 2102 |
| 10-3 | 35-55 | GARAH(Iph)KFRGHKWKK | 81 | 2103 |
| 10-3 | 35-55 | G(Dab)(Nap)(Pip)KPGRSN(Iph)KWKK | 84 | 2105 |
| 10-3 | 35-55 | GGG(Iph)(Pip)(Iph)Q(Dab)TE(Dab)KWKK | 74 | 2114 |
| 10-3 | 35-55 | G(Nap)ETAKG(Dab)(Iph)HRKWKK | 80 | 2119 |
| 10-3 | 35-55 | G(Pip)RQKT(Dab)FFQ(Nap)KWKK | 96 | 2128 |
| 10-3 | 35-55 | GNL(Dab)(Dab)R(Nap)AL(Iph)QKWKK | 78 | 2135 |
| 10-3 | 35-55 | GK(Iph)P(Dab)FKH(Cit)AKKWKK | 93 | 2136 |
| 10-3 | 35-55 | GS(Iph)L(Dab)S(Nap)(Nap)(Dab)L(Dab)KWKK | 82 | 2137 |
| 10-3 | 35-55 | G(Dab)GR(Iph)KKP(Iph)GVKWKK | 93 | 2138 |
| 10-3 | 35-55 | GT(Iph)RRN(Cit)(Dab)PDVKWKK | 91 | 2138 |
| 10-3 | 35-55 | GKG(Nap)HKKREL(Nap)KWKK | 89 | 2140 |
| 10-3 | 35-55 | GHERHRPPRL(Nap)KWKK | 82 | 2145 |
| 10-3 | 35-55 | GTPQRHRKLS(Iph)KWKK | 89 | 2146 |
| 10-3 | 35-55 | G(Dab)(Dab)N(Cit)(Iph)NRVKFKWKK | 90 | 2158 |
| 10-3 | 35-55 | GQVG(Pip)(Dab)VK(Iph)(Iph)NKWKK | 78 | 2167 |
| 10-3 | 35-55 | G(Nap)GHRAPVRR(Iph)KWKK | 81 | 2169 |
| 10-3 | 35-55 | G(Iph)PNVRHFHSRKWKK | 90 | 2173 |
| 10-3 | 35-55 | GHSSRT(Iph)RHRLKWKK | 86 | 2173 |
| 10-3 | 35-55 | GSR(Iph)K(Pip)SHT(Nap)NKWKK | 89 | 2176 |
| 10-3 | 35-55 | GHF(Iph)HRKPVHTKWKK | 92 | 2182 |
| 10-3 | 35-55 | GKPHNKLHR(Iph)EKWKK | 83 | 2182 |
| 10-3 | 35-55 | GFKKD(Iph)KFRGHKWKK | 97 | 2186 |
| 10-3 | 35-55 | GNHQQ(Iph)RARHQKWKK | 91 | 2198 |
| 10-3 | 35-55 | GDR(Nap)AF(Dab)(Iph)RGRKWKK | 95 | 2198 |
| 10-3 | 35-55 | GKHPR(Iph)FNNRLKWKK | 82 | 2205 |
| 10-3 | 35-55 | GHRHSVRKHE(Iph)KWKK | 93 | 2209 |
| 10-3 | 35-55 | GKK(Iph)FKT(Nap)TQHKWKK | 96 | 2238 |
| 10-3 | 35-55 | GK(Iph)A(Iph)(Dab)ERKTNKWKK | 95 | 2243 |
| 10-3 | 35-55 | GHF(Pip)VTHRL(Nap)(Iph)KWKK | 82 | 2256 |
| 10-3 | 35-55 | G(Nap)K(Dab)(Iph)(Cit)RFTQ(Dab)KWKK | 96 | 2257 |
| 10-3 | 35-55 | GLRHHRN(Cit)(Iph)KQKWKK | 96 | 2269 |
| 10-3 | 35-55 | GF(Dab)R(Nap)QKK(Iph)QEKWKK | 90 | 2284 |
| 10-3 | 35-55 | G(Dab)HRS(Iph)LHDK(Iph)KWKK | 78 | 2289 |
| 10-3 | 35-55 | GR(Pip)(Iph)T(Iph)PEF(Dab)KKWKK | 89 | 2300 |
| 10-3 | 35-55 | GTQKRR(Cit)F(Iph)(Dab)(Nap)KWKK | 85 | 2313 |
| 10-3 | 35-55 | GKP(Iph)VKK(Pip)(Nap)(Iph)PKWKK | 89 | 2316 |

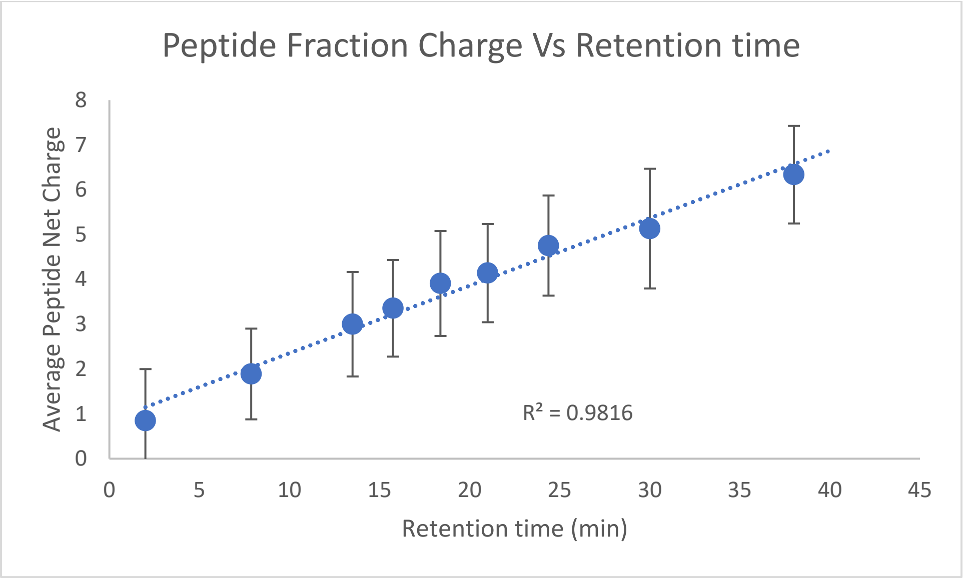

#### 5. References

(1) Simon, M. D.; Heider, P. L.; Adamo, A.; Vinogradov, A. A.; Mong, S. K.; Li, X.; Berger, T.; Policarpo, R. L.; Zhang, C.; Zou, Y.; Liao, X.; Spokoyny, A. M.; Jensen, K. F.; Pentelute, B. L. Rapid Flow-Based Peptide Synthesis. *ChemBioChem* **2014**, *15* (5), 713–720. https://doi.org/10.1002/CBIC.201300796.

(2) Schissel, C. K.; Farquhar, C. E.; Loas, A.; Malmberg, A. B.; Pentelute, B. L. In-Cell Penetration Selection-Mass Spectrometry Produces Noncanonical Peptides for Antisense Delivery. *ACS Chem bio* **2023**, *18* (3), 615–628. https://doi.org/10.1101/2022.04.13.488231.

(3) Vinogradov, A. A.; Gates, Z. P.; Zhang, C.; Quartararo, A. J.; Halloran, K. H.; Pentelute, B. L. Library Design-Facilitated High-Throughput Sequencing of Synthetic Peptide Libraries. *ACS Comb Sci* **2017**, *19* (11), 694–701. https://doi.org/10.1021/acscombsci.7b00109.
